# Mitochondrial Transplantation promotes protective effector and memory CD4^+^ T cell response during *Mycobacterium tuberculosis* infection and diminishes exhaustion and senescence in elderly CD4^+^ T cells

**DOI:** 10.1101/2024.01.24.577036

**Authors:** Colwyn A. Headley, Shalini Gautam, Angelica Olmo-Fontanez, Andreu Garcia-Vilanova, Varun Dwivedi, Alyssa Schami, Susan Weintraub, Philip S. Tsao, Jordi B. Torrelles, Joanne Turner

## Abstract

Tuberculosis (TB), caused by the bacterium *Mycobacterium tuberculosis* (*M.tb*), remains a significant health concern worldwide, especially in populations with weakened or compromised immune systems, such as the elderly. Proper adaptive immune function, particularly a CD4^+^ T cell response, is central to host immunity against *M.tb*. Chronic infections, such as *M.tb*, as well as aging promote T cell exhaustion and senescence, which can impair immune control and promote progression to TB disease. Mitochondrial dysfunction contributes to T cell dysfunction, both in aging and chronic infections and diseases. Mitochondrial perturbations can disrupt cellular metabolism, enhance oxidative stress, and impair T-cell signaling and effector functions. This study examined the impact of mitochondrial transplantation (mito-transfer) on CD4^+^ T cell differentiation and function using aged mouse models and human CD4^+^ T cells from elderly individuals. Our study revealed that mito-transfer in naïve CD4^+^ T cells promoted the generation of protective effector and memory CD4^+^ T cells during *M.tb* infection in mice. Further, mito-transfer enhanced the function of elderly human T cells by increasing their mitochondrial mass and modulating cytokine production, which in turn reduced exhaustion and senescence cell markers. Our results suggest that mito-transfer could be a novel strategy to reestablish aged CD4^+^ T cell function, potentially improving immune responses in the elderly and chronic TB patients, with a broader implication for other diseases where mitochondrial dysfunction is linked to T cell exhaustion and senescence.

## 1. Introduction

Tuberculosis (TB), primarily caused by *Mycobacterium tuberculosis* (*M.tb*) [1–4], remains a significant global health threat. It is the second leading cause of death by an infection and the 13^th^ leading cause of death worldwide [5]. *M.tb* infection is contracted through inhalation of aerosolized bacterial particles [1–4], and the ensuing immune response involves a complex interplay between host innate and adaptive immunity [1–4], with antigen presentation of *M.tb* antigens to T cells being a key element in the immune response against *M.tb* infection [1, 2, 6–13]. Several of these studies identified intrinsic defects in T cells as principal contributors toward increased bacterial burden in aged individuals infected with *M.tb* [14–21]. In this context, when splenic T cells from young *M.tb*-infected mice were adoptively transferred into irradiated and *M.tb*-infected mice of any age, the *M.tb* burden remained comparable across experimental groups [14]. However, transferring splenic T cells from older mice into irradiated young mice led to a significant increase in *M.tb* burden and dissemination, indicating that aging impacts T cell-mediated protection [14]. Therefore, understanding the impact of T cell dysfunction on TB in elderly populations is pivotal for the development of effective TB management and treatment strategies for this vulnerable group.

T cell exhaustion and senescence, the most characterized dysfunctional T cell end-states, are frequently observed in individuals with chronic infections and in the elderly [22–36]. T cell exhaustion, a result of persistent antigen exposure, leads to functional impairments such as altered transcriptomic signatures, decreased production of effector populations and cytokines, and diminished T cell proliferative potential [22–27]. For *M.tb* infection, exhausted CD4^+^ T cells exhibit increased expression of inhibitory receptors such as PD-1, LAG-3, and TIM-3, which negatively impact T cell activation and effector functions [26, 36]. T cell senescence refers to age-related decline in T cell function, which involves a reduced T cell repertoire, altered subset distribution, impaired functionality, and dysregulated signaling pathways [28–35]. Although exhaustion and senescence are distinct T cell fates, both contribute to a weakened immune response in the elderly. Senescence reduces the naive T cell pool and the few remaining naive T cells often exhibit intrinsic signaling defects, dampening their proper activation and differentiation. Those naive cells that do transition toward effector and memory lineages are often subjected to constant antigenic stress, driving them toward exhaustion. These combined factors result in an impaired ability to respond effectively to pathogens, including *M.tb*.

Any perturbation in mitochondrial function can disrupt T cell metabolism, interfere with calcium and redox signaling pathways, increase oxidative stress, and impair T cell signaling and effector functions [24, 29, 35, 37–43]. Mitochondrial dysfunction (mito-dysfunction) is a pivotal contributor to T cell exhaustion and senescence [24, 29, 35, 37–43] and can be characterized by reduced quality of mitochondrial DNA, decreased efficiency of the electron transport chain (ETC), irregular mitochondrial membrane potential (Δψm), and abnormal generation of reactive oxygen species (mitoROS) [24, 29, 35, 37–43]. Addressing mito-dysfunction could offer a therapeutic opportunity to enhance immune responses against TB [44]. This approach may also improve responses to anti-TB therapy, which usually involves a lengthy drug regimen, that becomes increasing hard to maintain in individuals such as the elderly due to their associated cytotoxicity [45].

Our previous studies indicated the potential of mitochondrial transplantation (mito-transfer) to improve the function of CD4^+^ T cells isolated from 18-22 month old (aged) mice conferring higher resistance against *M.tb* infection [44]. Mito-transfer is the process of delivering healthy/non-dysfunctional mitochondria to diseased or dysfunctional cells, tissue, or organs to improve functionality [46–51]. Our results demonstrated that the delivery of exogenous mitochondria to aged non-activated CD4^+^ T cells led to significant mitochondrial proteome alterations highlighted by improved aerobic metabolism and decreased cellular mitoROS. Additionally, mito-transferred aged CD4^+^ T cells showed improvement in activation-induced TCR-signaling kinetics, markers of IL-2 signaling (CD25), increased IL-2 production, and enhanced proliferation *ex vivo*. Importantly, we showed that adoptive transfer of mito-transferred naive aged CD4^+^ T cells to RAG-KO mice protected them from *M.tb* infection [44]. These findings underscored the potential of targeted mito-transfer in rescuing aged lymphocytes, with important implications for future immunological and therapeutic studies for chronic infections and diseases [44].

Herein, we assessed the impact of mito-transfer on CD4^+^ T cell differentiation during *M.tb* infection. Our data reveal that mito-transfer promoted the generation of protective effector and effector memory CD4^+^ T cells in aged mice during *M.tb* infection. Interestingly, we observed a concomitant reduction in T cell exhaustion, suggesting that mito-transfer may at least transiently alleviate some age-related effects on T cell function. Additionally, mito-transfer enhanced CD4^+^ T cell function in elderly individuals, evidenced by increased mitochondrial mass, modulated cytokine production, enhanced T cell activation, and reduced markers of exhaustion and senescence. These findings highlight the potential of targeting mitochondrial dysfunction, with mito-transfer as a mean to enhance immune responses and improve treatment outcomes in elderly individuals infected with *M.tb*, or potentially other chronic infections or diseases.

## 2. Methods

### Mice

Specific pathogen-free C57BL/6 young (2 to 4 months) and old (18 to 22 months) mice were obtained from the National Institutes of Health/National Institute on Aging (NIH/NIA), Charles River Laboratories (Wilmington, MA, USA), or The Jackson Laboratories (JAX, Bar Harber, ME, USA). TCR-KO (B6.129S2-Tcra^tm1Mom/J^, 6 weeks old) mice were obtained from JAX. Mice were housed in groups of 5 in individually sterile ventilated cages and fed a standard chow diet, *ad libitum*, for the duration of the study. All procedures were approved by the Texas Biomedical Research Insitutte (Texas Biomed) institutional laboratory animal care and use committee (IACUC) under protocol 1608 MU, and the biosafety committee protocol #17-006.

### Isolation of CD4^+^ T cells from mice and human PBMC

Young and old mice were humanely euthanized by CO_2_ uptake, and spleens surgically excised. Isolated spleens were individually placed in 6 well tissue culture plates containing 4 ml of chilled (4°C) complete DMEM and gently hand homogenized using sterile syringe stoppers. Spleen homogenate was passed through a 70 µm microfilter to remove large debris. The resulting single-cell suspensions were centrifuged for 5 min at 300 x *g* at 4°C and suspended in Gey’s lysis buffer (8 mM NH_4_Cl, 5 mM KHCO_3_) for 1 min to lyse erythrocytes, and then neutralized with an equivalent volume of complete DMEM. Splenocytes were washed and suspended in separation buffer (10 mM D-Glucose, 0.5 mM MgCl_2_, 4.5 mM KCl, 0.7 mM Na_2_HPO_4_, 1.3 mM NaH_2_PO_4_, 25 mM NaHCO_3_, 20 mM HEPES, 5 µM DETC, 25 µM deferoxamine) containing appropriate antibody-magnetic particles for negative selection (α-CD8, α-CD11b, α-CD109, α-CD24, α-CD45, α-CD49b, α-Ly-6G, α-ɣδ TCR, α-TER-119). The resulting population of CD4^+^ T cells (85-95% purity) was used for downstream experiments.

Human de-identified donor buffy coats were commercially obtained from Innovative Research (Novi, MI, USA). Human CD4^+^ T cells were purified via negative selection using available kits (Miltenyi Biotech, Auburn, CA), following the manufacturer’s protocol. The donors were majority black (8 black, 2 Caucasian), Male (7 male, 3 female), and ranged in age from 51 to 66 (mean age 60.5) **(Supp. Table 1).**

### Isolation of mouse embryonic fibroblasts

Mouse embryonic fibroblasts (MEFs) were isolated from 12 to 14 days post-coitum C57BL/6 mice as described [52]. Briefly, mouse embryos were aseptically isolated, minced, and digested in 0.05% trypsin and crude DNase solution for 30 min at 37°C, 5% CO_2_, in a humidified incubator. Primary MEFs were cultured in sterile T-150 flasks (Corning, NY, USA), in 50:50 F12/DMEM (Corning) supplemented with 10% FBS, 1% HEPES, 2% NEAA, 1 mM sodium pyruvate, 2 mM L-glutamine, and 100 µg/ml primocin. MEFs were sub-passaged as necessary, with MEFs from passages 2 to 3 being used for all experiments.

### Mitochondrial isolation and transfer

Mitochondria were isolated from donor mouse or human fibroblasts (MEFs and human neonatal dermal fibroblasts, respectively) and mito-transferred by centrifugation with CD4^+^ T cells using a published protocol [53]. Briefly, MEFs were homogenized in SHE buffer [250 mM sucrose, 20 mM HEPES, 2 mM EGTA, 10 mM KCl, 1.5 mM MgCl_2_, and 0.1% defatted bovine serum albumin (BSA)], containing complete Minitab protease inhibitor cocktail (Sigma Alrich, MO, US). Cell homogenates were centrifuged at 800 x *g* for 3 min at 4°C to remove cellular debris. The resulting supernatant was recovered and centrifuged at 10,000 x *g* for 5 min to pellet isolated mitochondria. CD4^+^ T cells and isolated mitochondria were suspended in 150 μL cold PBS in V-bottom 96 well plates, and plates were centrifuged at 1,400 x *g* for 5 min at 4°C. After centrifugation, cells were washed 2 to 3 times with cold PBS before further experimentation. Independent of mito-transfer, all experimental groups were centrifuged accordingly.

### Proteomics

Cell preparations and extracted mitochondria were lysed in a buffer containing 5% SDS/50 mM triethylammonium bicarbonate (TEAB) in the presence of protease and phosphatase inhibitors (Halt; Thermo Scientific) and nuclease (Pierce™ Universal Nuclease for Cell Lysis; Thermo Scientific). Aliquots normalized to 65 µg protein (EZQ™ Protein Quantitation Kit; Thermo Scientific) were reduced with tris(2-carboxyethyl) phosphine hydrochloride (TCEP), alkylated in the dark with iodoacetamide, and applied to S-Traps (mini; Protifi) for tryptic digestion (sequencing grade; Promega) in 50 mM TEAB. Peptides were eluted from the S-Traps with 0.2% formic acid in 50% aqueous acetonitrile and quantified using Pierce™ Quantitative Fluorometric Peptide Assay (Thermo Scientific).

On-line high-performance liquid chromatography (HPLC) separation was accomplished with an RSLC NANO HPLC system [Thermo Scientific/Dionex: column, PicoFrit™ (New Objective; 75 μm i.d.) packed to 15 cm with C18 adsorbent (Vydac; 218MS 5 μm, 300 Å)], using as mobile phase A, 0.5% acetic acid (HAc)/0.005% trifluoroacetic acid (TFA) in water; and as mobile phase B, 90% acetonitrile/0.5% HAc/0.005% TFA/9.5% water; gradient 3 to 42% B in 120 min; flow rate at 0.4 μl/min. Data-independent acquisition mass spectrometry (DIA-MS) was conducted on an Orbitrap Fusion Lumos mass spectrometer (Thermo Scientific). A pool was made of all of the HPLC fractions (experimental samples), and 2 µg peptide aliquots were analyzed using gas-phase fractionation and 4-*m/z* windows (120k resolution for precursor scans, 30k for product ion scans, all in the Orbitrap) to create a DIA-MS chromatogram library [54] by searching against a panhuman spectral library [55]. Subsequently, experimental samples were blocked by replicate and randomized within each replicate. Injections of 2 µg of peptides were employed. MS data for experimental samples were also acquired in the Orbitrap using 12-*m/z* windows (staggered; 120k resolution for precursor scans, 30k for product ion scans) and searched against the DIA-MS chromatogram library generated. Scaffold DIA (ver. 1.3.1; Proteome Software) was used for all DIA data processing.

### Bioinformatic Analysis

The MetaScape platform was utilized for gene annotation and discovery [56]. The proteomic libraries generated from CD4^+^ T cells isolated from old mice that received healthy mitochondria (OM) and un-manipulated CD4^+^ T cells from old mice (O) were filtered against previously published data sets associated with CD4^+^ T cell exhaustion [57] and cellular aging/senescence [58], by matching uniport ID to gene names/symbol. Resulting DEPs were considered statistically significant when the false discovery rate (FDR)-adjusted *p*-value was less than 0.05. Bioinformatic analysis on these significant DEPs via the MetaScape bioinformatics tool was conducted [56]. The analysis encompassed Pathway & Process Enrichment (PPE) which revealed enriched functional pathways, and Protein-Protein Interaction Enrichment (PPIE) which identified Protein-Protein interaction (PPI) networks and PPI network clusters via Molecular Complex Detection (MCODE) [56].

### T cell activation

For experiments requiring human CD4^+^ T cell activation, cells were incubated in complete RPMI or Immunocult T Cell expansion media supplemented with phorbol 12-myristate 13-acetate (PMA) and Ionomycin (20 ng/ml and 1 µg/ml respectively), αCD3/αCD28 (0.5 µg/ml and 0.05 µg/ml respectively), or CD3-CD28-CD2 activation beads (STEMCELL;25 ul/ 1×10^6^ cells) for specified times.

### Quantitation of intracellular reactive oxigen species (ROS)

For flow cytometry-based ROS experiments, CD4^+^ T cells were incubated in media containing MitoSOX (5 µM) for 30 min at 37°C, to measure mitochondrial and cytosolic ROS, respectively. Cells were washed twice with phosphate-buffered saline (PBS, pH 7.4) and stained with antibodies targeting surface proteins of interest (*i.e.* α-CD3, α-CD4, and α-CD69; 25 µg/ml) for 20 min before fixation in 2% paraformaldehyde. Flow cytometry-based ROS and other flow-based quantifications were performed on a BD-LSR II (BD Biosciences, NJ, USA), BD FACs Symphony, Cyan or CytoFLEX LX (Beckman Coulter, CA, USA) flow cytometers. Data were analyzed using FlowJo (FlowJo LLC-BD, NJ, USA) and GraphPad Prism ver. 9 (GraphPad Software, CA, USA).

### Adoptive transfer of naïve CD4^+^ T cells and infection of mice with *M.tb*

Naïve CD4^+^ T cells from old mice were isolated by negative magnetic bead selection. Non-manipulated naïve CD4^+^ T cells or mito-transferred naïve CD4^+^ T cells (∼ 2.0 x 10^6^ cells) were tail vein injected into TCR-KO mice (n= 4 to 5 mice per group) in a final volume of 200 µl. A control group of mice (n= 3 to 4 mice) were tail-vein injected with 200 µl of CD4^+^ T cells (∼ 2.0 x 10^6^ cells) isolated from young mice. Roughly 24 h after injections, mice were infected with *M.tb* (strain Erdman, ATCC no 35801), via low dose aerosol infection (∼100 colony forming units, CFU) [59]. The presence of colitis in TCR-KO mice adoptively transferred with naïve CD4^+^ T cells was scored in a blinded manner by a board-certified veterinary pathologist in the Pathology Core at Texas Biomedical Research Institute. No visible pathologies were obeserved.

### Quantitation of mitochondrial mass and mitochondrial viability/membrane potential

Cellular mitochondrial mass and potential were determined using commercially available probes MitoView Green (MTV-G; Biotium, CA, USA) and Mitotracker Deep Red (MTDR; Life Technologies, CA, USA), respectively. Antibody labeled (α-CD4) CD4^+^ T cells were fixed in 2% paraformaldehyde before MTV-G staining (50 nM) for 30 min at RT or CD4^+^ T cells were stained with MTDR (50 nM) for 20 min before fixation in 2% paraformaldehyde. MFI of MTV-G and MTDR^+^ CD4^+^ T cells was determined by flow cytometry.

### Quantitation of GSH

The total GSH detection assay was performed according to the manufacturer’s directions.

### Cytokine detection

Briefly, 2.5 x 10^5^ CD4^+^ T cells were cultured in 96 well tissue culture plates containing PMA/Ionomycin for 24 h or 72 h, after which media were collected and measured for the presence of cytokines IL-2, IFNɣ, IL-4, IL-5, IL-10, IL-17/IL-17A, TNF, and the chemokine CCL5/RANTES following the manufacturer’s protocol.

### Proliferation Assay

Human CD4^+^ T cell proliferation was determined by the Click-it EdU detection assay (Thermo Fisher) [60]. Briefly, 2.5 x 10^5^ CD4^+^ purified T cells were activated for 72 h in 600 µL RPMI containing 50 µM 5-Ethynyl-2’-deoxyuridine (EdU) and αCD28 (0.05 µg/ml), in sterile 48 well plates precoated with 400 µL of αCD3 (0.5 µg/ml in PBS), and in RPMI containing PMA/Ionomycin. EdU incorporation into DNA from proliferating cells was detected by a fluorescent bioconjugate and flow cytometry. The assay was performed per the manufacturer’s instructions.

### β-Galactosidase Staining

β-galactosidase staining was done using the commercially available kit CellEvent™ Senescence Green Flow Cytometry Assay (Invitrogen), following the manufacturer’s protocol.

## 3. Results

### 3.1 Mito-transfer impacts processes beyond the mitochondrial proteome, including T cell exhaustion

To investigate the impact of mito-transfer on T cell exhaustion, we first performed bioinformatic analysis on the proteomic library of CD4^+^ T cells from old mice, with or without mito-transfer, via the Metascape platform. A total of 578 differentially expressed proteins (DEPs) related to T cell exhaustion (Tex-DEPs) were uncovered. Of these DEPs, 295 DEPs were significantly higher and 283 DEPs were significantly lower in abundance in CD4^+^ T cells from old mice post mito-transfer **(Fig. 1. A, Supp. Table 2)**.

**Fig. 1.**
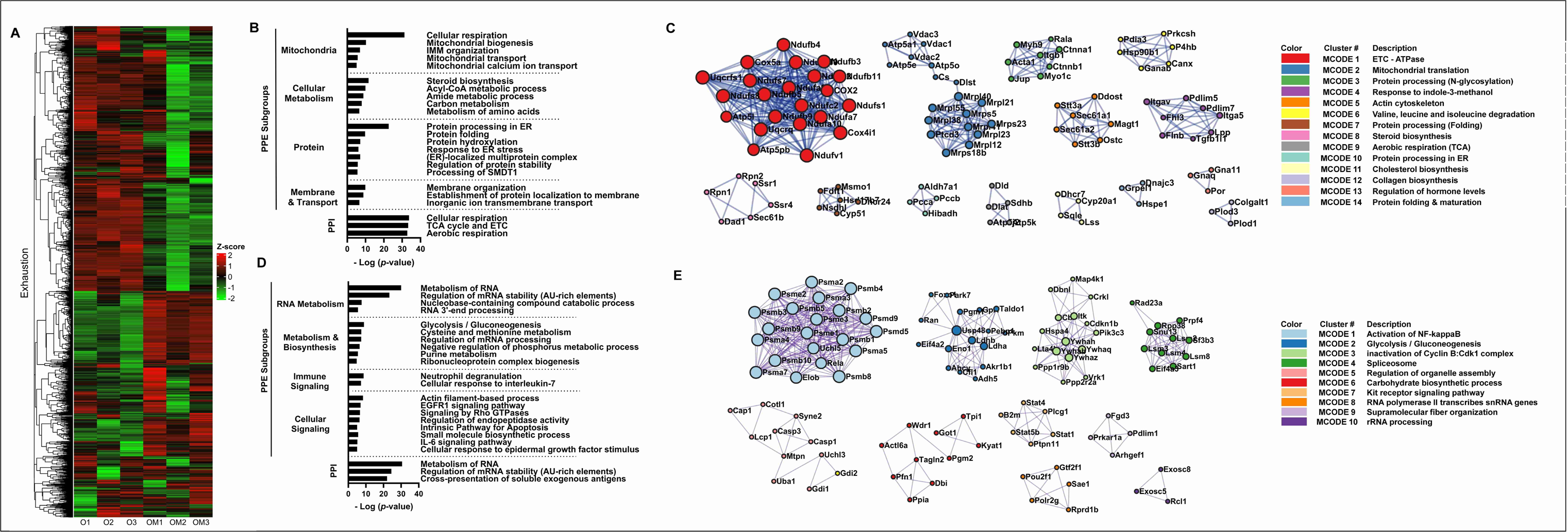
Mito-transfer modulates the expression of CD4^+^ T cell exhaustion-associated genes. **A)** Heatmap of proteins related to T cell exhaustion from CD4+ T cells from old mice with and without mito-transfer (4h). **B)** Pathway & Process Enrichment (PPE) analysis & Protein-Protein Interaction (PPI) networks, and **C)** Molecular Complex Detection **(MCODE)** clusters identified as significantly upregulated in CD4^+^ T cells from old mice after mito-transfer. **D)** Pathway & Process Enrichment (PPE) analysis & Protein-Protein Interaction (PPI) networks, and **E)** Molecular Complex Detection (MCODE) clusters identified as significantly downregulated in CD4^+^ T cells from old mice after mito-transfer. Bioinformatic Analysis was performed using the MetaScape analysis tool.

The PPEs of upregulated Tex-DEPs were manually parsed into four PPE sub-groups: mitochondria, cellular metabolism, proteins, and membrane & transport **(Fig. 1. B, Supp. Table 3)**. Within the mitochondria PPE subgroup, pathways associated with mitochondrial biogenesis, oxidative phosphorylation, and calcium transport were uncovered **(Fig. 1. B)**, suggesting increased mitochondrial activity and turnover in mito-transferred CD4^+^ T cells. Processes and pathways within the cellular metabolism PPE subgroup included steroid biosynthesis, acyl-CoA metabolism, amide metabolism, carbon metabolism, and metabolism of amino acids, and which are also interconnected to mitochondria **(Fig. 1. B)**. Within the Protein PPE subgroup, processes related to endoplasmic reticulum (ER) function including protein folding, post-translation hydroxylation modifications, and ER stress response, were identified **(Fig. 1. B)**. The membrane & transport PPE subgroup were primarily related to membrane assembly, and transmembrane ion transport **(Fig. 1. B)**. PPIE analysis further revealed 3 PPI networks (**Fig. 1. B, Supp. Table 4**) and 14 MCODE clusters (MCs) (**Fig. 1. C**). While the uncovered PPI networks were all directly related to aerobic respiration **(Fig. 1. B)**, the 14 MCODE clusters were manually binned into subgroups, namely mitochondria (MC-1, 2, 6 and 9), protein processing (MC-3, 7, 10, and 14), cell organization & signaling (cluster 5) and, lipid metabolism (MC-8 and 11) **(Fig. 1. C, Supp. Table 5).**

The PPEs of downregulated Tex-DEPs, were also manually parsed into four subgroups; RNA metabolism, cellular metabolism & biosynthesis, immune signaling, and cellular signaling **(Fig. 1. D, Supp. Table. 6**. Within the RNA metabolism PPE subgroup, processes related to mRNA stability and 3’-end processing were identified. **(Fig. 1. D)**. The processes within the metabolism and biosynthesis PPE subgroup were related to glycolysis/gluconeogenesis, cysteine and methionine metabolism, and pyruvate metabolism **(Fig. 1. D)**. In the Immune signaling PPE subgroup, neutrophil degranulation and IL-7 signaling were identified. While prominent processes in the cellular signaling PPE subgroup included Rho GTPases, EGFR1, IL-6, and apoptotic signaling **(Fig. 1. D)**. Regarding PPI network analysis, 2/3 downregulated networks pertained to RNA regulation and stability, with the third network pertaining to the cross-presentation of soluble exogenous antigens from endosomes **(Fig. 1. D, Supp. Table 7)**. The MCODE analysis of downregulated DEPs related to T cell exhaustion, identified 10 clusters that were manually sub-grouped into hypoxic signaling (MC 1), metabolic reprogramming/glycolysis (MC-2 and 6), cell cycle (MC-3 and 7), cell structure (MC-5 and 9), and RNA processing (MC-4,8 and 10) **(Fig. 1. E, Supp. Table 8).** Collectively, these data suggest that mito-transfer modulates the expression of proteins associated with T cell exhaustion, and drives reprogramming of CD4^+^ T cell bioenergetics, shifting from glycolysis towards oxidative phosphorylation.

### 3.2 Mito-transfer impacted aging/senescence-associated pathways in CD4^+^ T cells from old mice

We further examined the impact of mito-transfer on the expression of senescence associated proteins in CD4^+^ T cells from old mice. A total of 24 senescence DEPs (Sen-DEPs) were identified; 6 were significantly upregulated, and 18 were significantly downregulated **(Fig. 2. A, Supp. Table 9)**. The 6 upregulated Sen-DEPs (Atp5o, Lmna, Hspd1, Sumf1, Pycr1, and Dbn1) did not substantiate any bioinformatic insight on associated pathways. However, the 18 downregulated Sen-DEPs were associated with cellular processes related to cellular signaling and stress responses, ROS and DNA damage responses, metabolism and catabolic processes, immune responses, and cell death **(Fig. 2. B, Supp. Table 10)**. The PPI network analysis of downregulated Sen-DPEs revealed a dampening in IL-6 and IL-3 signaling **(Fig. 2. B, Supp. Table 11)**. While MCODE cluster analysis uncovered 2 clusters; a metabolic cluster related to glycolysis/gluconeogenesis, and a signal transduction cluster **(Fig. 2. C, Supp. Table 12)**. These findings also suggest a reprogramming of the T cell’s energy metabolism following mito-transfer, potentially shifting from glycolysis towards oxidative phosphorylation. The intracellular signaling cluster was associated with PDGF, Kit receptor signaling, and downstream signal transduction. PDGF is a pathway crucial for cell growth and division [61], and Kit receptor signaling is essential for cell survival, proliferation, and differentiation [62]. These data suggest that mito-transfer may impact signal transduction and cell communication, influencing T cell growth, proliferation, and immune responses.

**Fig. 2.**
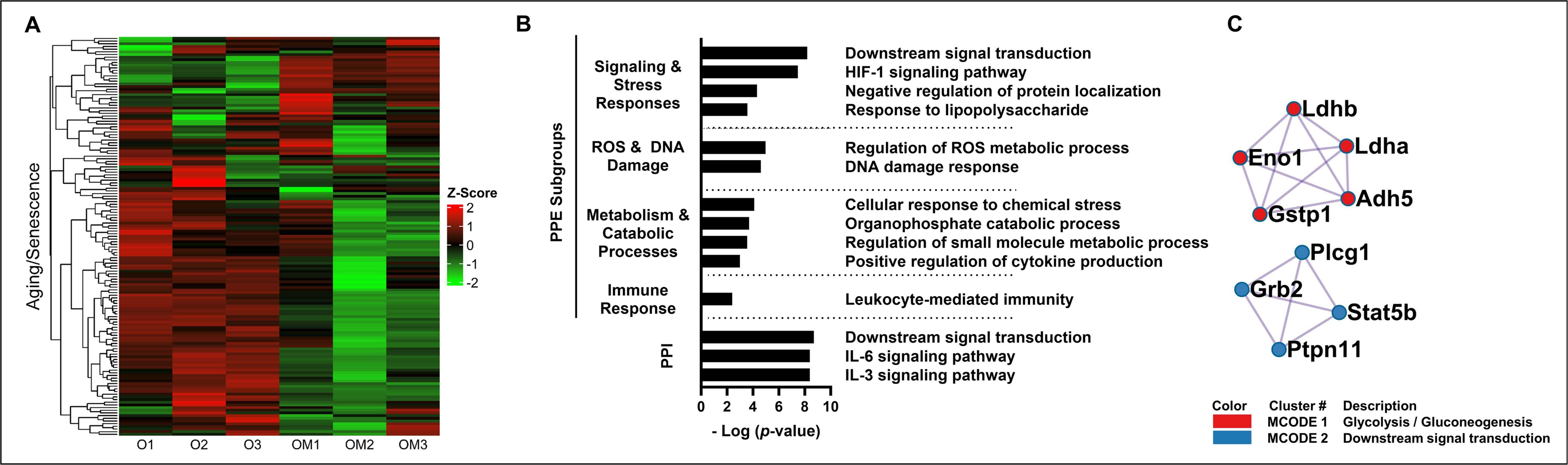
Mito-transfer modulates the expression of CD4^+^ T cell senescence-associated genes. **A)** Heatmap of proteins related to T cell aging/senescence from CD4+ T cells from old mice with and without mito-transfer (4h). **B)** Pathway & Process Enrichment (PPE) analysis & Protein-Protein Interaction (PPI) networks, and **C)** Molecular Complex Detection (MCODE) clusters identified as significantly downregulated in CD4^+^ T cells from old mice after mito-transfer. Bioinformatic Analysis was performed using the MetaScape analysis tool.

### 3.3 Mito-transfer in naïve CD4 T cells from old mice drives T cell differentiation

In our previous study, we demonstrated that mito-transferred naïve CD4^+^ T cells from old mice had improved control of influenza A virus (IAV) or *M.tb* infection in RAG-KO mice [44]. Specific to the *M.tb* experiments, in comparison to *M.tb* infected-RAG-KO that received either naïve CD4^+^ T cells from old mice or no naive CD4^+^ T cells, the mice that received mito-transferred naïve CD4^+^ T cells from old mice had the lowest *M.tb* burden in the lung and spleen, at 21 days post-infection [44]. Combined with these previous results, and after delineating the Tex-DEPs and Sen-DEPs pathways likely innervated by mito-transfer, we next aimed to understand the functional implications of these changes to CD4^+^ T cells during *M.tb* infection.

To achieve this, we used a TCR-KO adoptive transfer model of *M.tb* infection (i.e. TB-TKO), and evaluated the subpopulations of CD4^+^ T cells that developed in mice at 30 and 60 days post-infection, after receiving naive CD4^+^ T cells from old mice with (*test group; OM-TB-TKO*) or without (*old control; O-TB-TKO*) mito-transfer **[Figs. 3** (lung) and **4** (spleen)]. We additionally compared these groups against TCR-KO mice adoptively transferred with naïve CD4^+^ T cells from young mice (*young control; Y-TB-TKO*) **[Figs. 3** (lung) and **4** (spleen)]. Because the maximum survival of *M.tb-*infected TCR-KO mice is approximately 50 days [13, 63], the *M.tb*-infected TCR-KO mice were orally treated with antimycobacterial drugs isoniazid (INH; 50mg/kg/day) and rifampicin (RIF; 20mg/kg/day) from 30 to 60 days post-infection (*i.e.* to study endpoint). This timeframe reduces the *M.tb* burden sufficiently to extend survival, but does not fully eradicate a chronic infection [64]. In the context of *M.tb* infection, protective effector and effector memory CD4^+^ T cells are CD27^-^ [65, 66] with either PD1^+^ or KLRG1^+^ expression, while exhausted CD4^+^ T cells express TIM3^+^ [8, 36]. Therefore, we measured CD4^+^ T cell activation (CD69^+^, CD25^+^), generation of effector memory (CD27^-^, PD1^+^, KLRG1^+^), and exhaustion (TIM3^+^) **[Figs. 3** (lung) and **4** (spleen)].

**Fig. 3.**
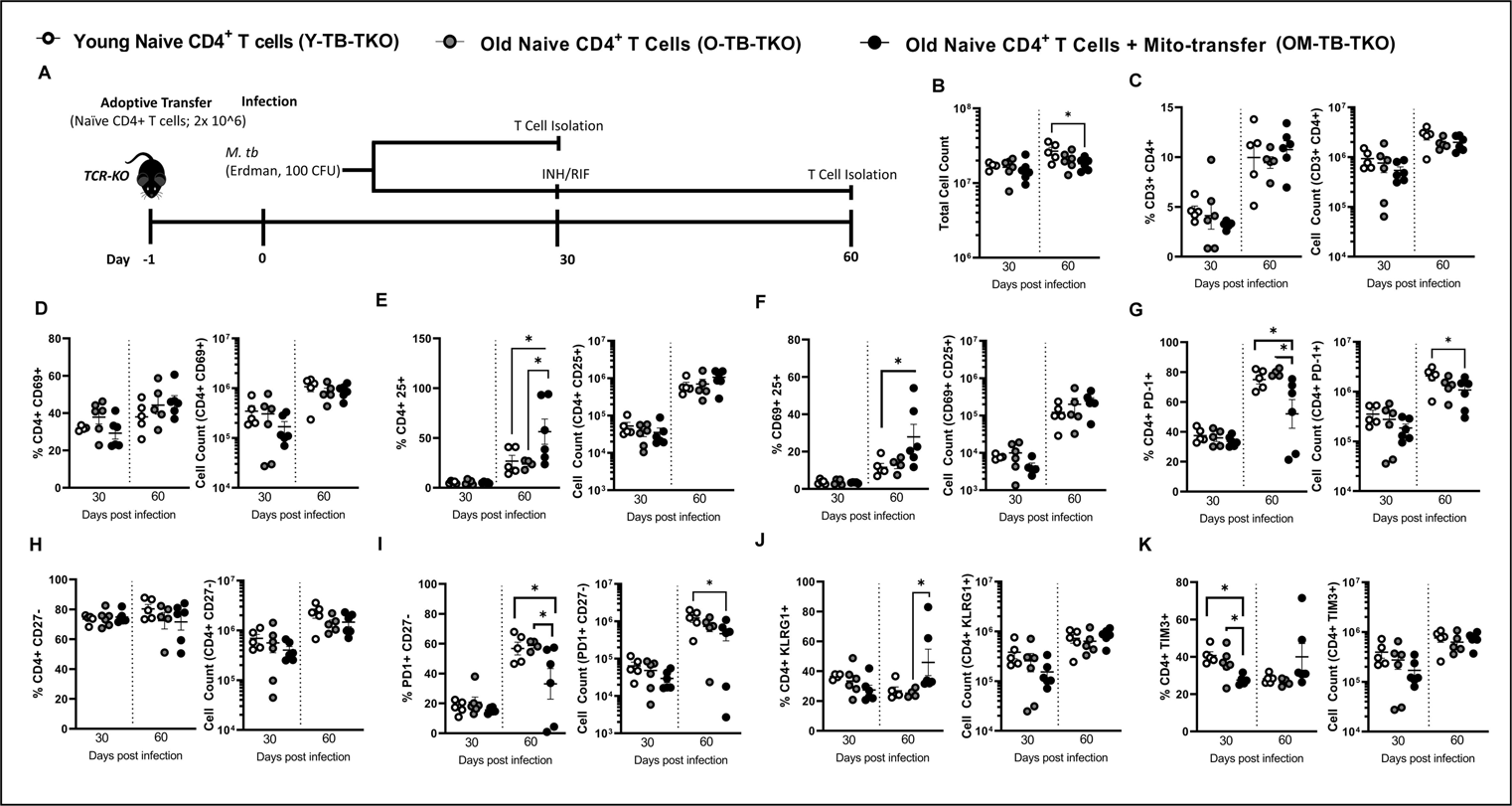
Mito-transfer in naïve CD4 T cells from old mice protects mice against pathogens & promotes protective T cell phenotypes in the lung. **A)** 2.0 x 10^6^ naïve CD4^+^ T cells from young mice and old mice with or without mito-transfer, were tail vein injected in TCR-KO mice, that were the subsequently infected *M.tb.* A subset of mice were euthanized at 30 days post-infection. Another subset of mice were treated with INH/RIF for an additional 30 days (up to 60 days post-infection). The absolute and percentage of various CD4^+^T cell subsets in the lung of *M.tb* infected mice were evaluated at both time points (30 and 60 days post-infection). **B)** Absolute cell count of the digested lung from TCR-KO mice infected with *M.tb*. The percents and absolute counts of **C)** CD3^+^CD4^+^ T cells, **D)** CD4^+^CD69^+^,E) CD4^+^CD25^+^, **F)** CD4^+^ CD69^+^ CD25^+^, **G)** CD4^+^ PD-1^+^, **H)** CD4^+^ CD27^-^, **I)** CD4^+^ PD-1^+^ CD27^-^, **J)** CD4^+^ KLRG1^+^, **K)** CD4^+^ TIM3^+^ subsets isolated from TCR-KO mice infected with *M.tb*, at 30 and 60 days post-infection. p< 0.05 = significant (*) using one-way-ANOVA.

At 30 days post-infection, there were no significant differences in the total lung cell numbers among Y-TB-TKO, O-TB-TKO and OM-TB-TKO experimental groups **(Fig. 3. B)**. Additionally, at 30 days post-infection, except for a lower frequency of TIM-3^+^ CD4^+^ T cells in the lung of OM-TB-TKO mice, no other significant differences in CD4^+^ T cell frequencies or absolute numbers were observed **(Fig. 3. C-K)**. However, at 60 days post-infection, the lung of OM-TB-TKO mice, contained significantly fewer cells **(Fig. 3. B),** but with higher frequencies of CD25^+^ CD4^+^ T cells **(Fig. 3. E)**, CD69^+^ CD25^+^ CD4^+^ T cells **(Fig. 3. F)**, KLRG1^+^ CD4^+^ T cells **(Fig. 3. J)**, and lower frequencies of PD1^+^ CD4^+^ T cells **(Fig. 3. G)**, and PD1^+^ CD27^-^ CD4^+^ T cells **(Fig. 3. I)** than Y-TB-TKO or O-TB-TKO mice. No other significant differences were noted among the groups of *M.tb-*infected TCR-KO mice.

The T cell composition in the spleen of TB-TKO mice were also assessed **(Fig. 4. A-K)**. At 30 days post-infection OM-TB-TKO and Y-TB-TKO mice had comparable frequencies of CD25^+^ CD4^+^T cells **(Fig. 4. E)**, which were both significantly higher than that of O-TB-TKO mice **(Fig. 4. E).** No other significant differences were noted among TB-TKO experimental groups at this timepoint **(Fig. 4. A-K)**. At 60 days post-infection, the spleen of OM-TB-TKO mice contained significantly lower frequencies of CD3^+^ CD4^+^ T cells **(Fig. 4. C)**, CD69^+^ CD4^+^ T cells **(Fig. 4. D)**, CD25^+^ CD69^+^ T cells **(Fig. 4. F)**, PD1^+^ CD27^-^ T cells **(Fig. 4. I)**, and TIM3^+^ CD4^+^ T cells **(Fig. 4. K)**, when compared against O-TB-TKO or Y-TB-TKO mice. These data collectively suggest that mito-transfer in naïve CD4^+^ T cells from old mice promotes memory T cell differentiation in *M.tb* infected TCR-KO mice.

**Fig. 4.**
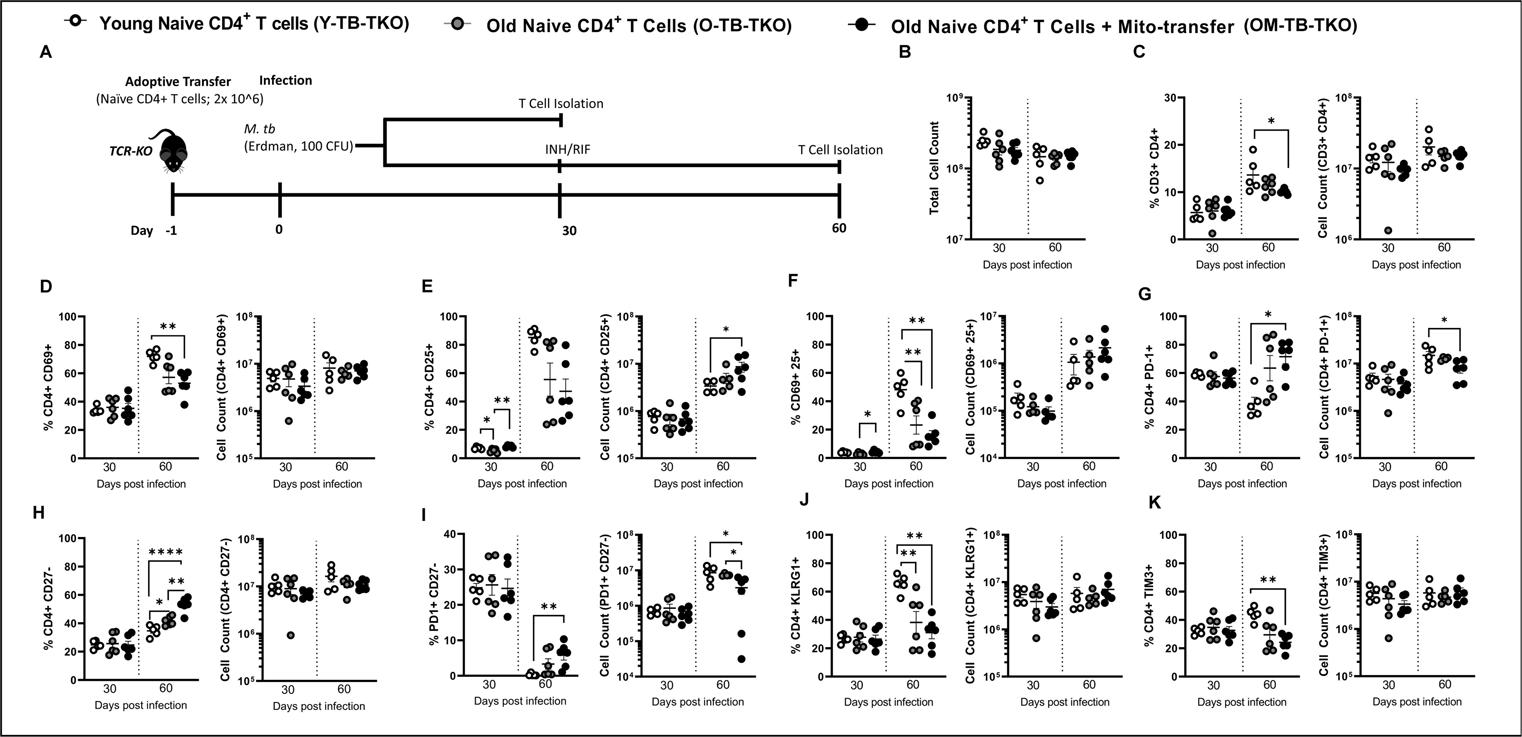
Mito-transfer in naïve CD4 T cells from old mice protects mice against pathogens & promotes protective T cell phenotypes in the spleen. **A)** 2.0 x 10^6^ naïve CD4^+^ T cells from young mice and old mice with or without mito-transfer, were tail vein injected in TCR-KO mice, that were the subsequently infected *M.tb.* A subset of mice were euthanized at 30 days post-infection. Another subset of mice were treated with INH/RIF for an additional 30 days (up to 60 days post-infection). The absolute and percentage of various CD4^+^T cell subsets in the spleen of *M.tb* infected mice were evaluated at both time points (30 and 60 days post-infection). **B)** Absolute cell count of the digested lung from TCR-KO mice infected with *M.tb*. The percents and absolute counts of **C)** CD3^+^CD4^+^ T cells, **D)** CD4^+^CD69^+^,E) CD4^+^CD25^+^, **F)** CD4^+^ CD69^+^ CD25^+^, **G)** CD4^+^ PD-1^+^, **H)** CD4^+^ CD27^-^, **I)** CD4^+^ PD-1^+^ CD27^-^, **J)** CD4^+^ KLRG1^+^, **K)** CD4^+^ TIM3^+^ subsets isolated from TCR-KO mice infected with *M.tb*, at 30 and 60 days post-infection. p< 0.05 = significant (*) using one-way-ANOVA.

### 3.4 Mito-transfer improves the function of elderly CD4^+^ T cells

We also determined whether mito-transfer could provide comparable improvements to human CD4^+^ T cells isolated from elderly individuals. Donor mitochondria isolated from primary human neonatal dermal fibroblasts were transplanted into CD4^+^ T cells from elderly individuals (mean age 60.5 years old), after which the relative mitochondrial mass and mitoROS were examined in both non-activated and activated CD4^+^ T cell cultures, by quantifying the MFI of MTV-G and MitoSOX, respectively.

Compared to non-manipulated elderly CD4^+^ T cells (i.e. *E-CD4*^+^ *T cells*), the relative MFI of MTV-G in elderly CD4^+^ T cells that received mito-transfer (i.e. *EM-CD4*^+^ *T cells)* significantly increased in both unstimulated and PMA/ionomycin stimulated conditions **(Fig. 5. A)**. These data indicate that mito-transfer increased the mitochondrial mass of CD4^+^ T cells from elderly individuals. Compared to E-CD4^+^ T cells, the relative MFI of MitoSOX in EM-CD4^+^ T cells significantly decreased under basal conditions, and increased under PMA/ionomycin stimulated conditions **(Fig. 5. B)**. These data indicate that mito-transfer decreased basal mitoROS and increased activation-induced mitoROS in elderly CD4^+^ T cells.

**Fig. 5.**
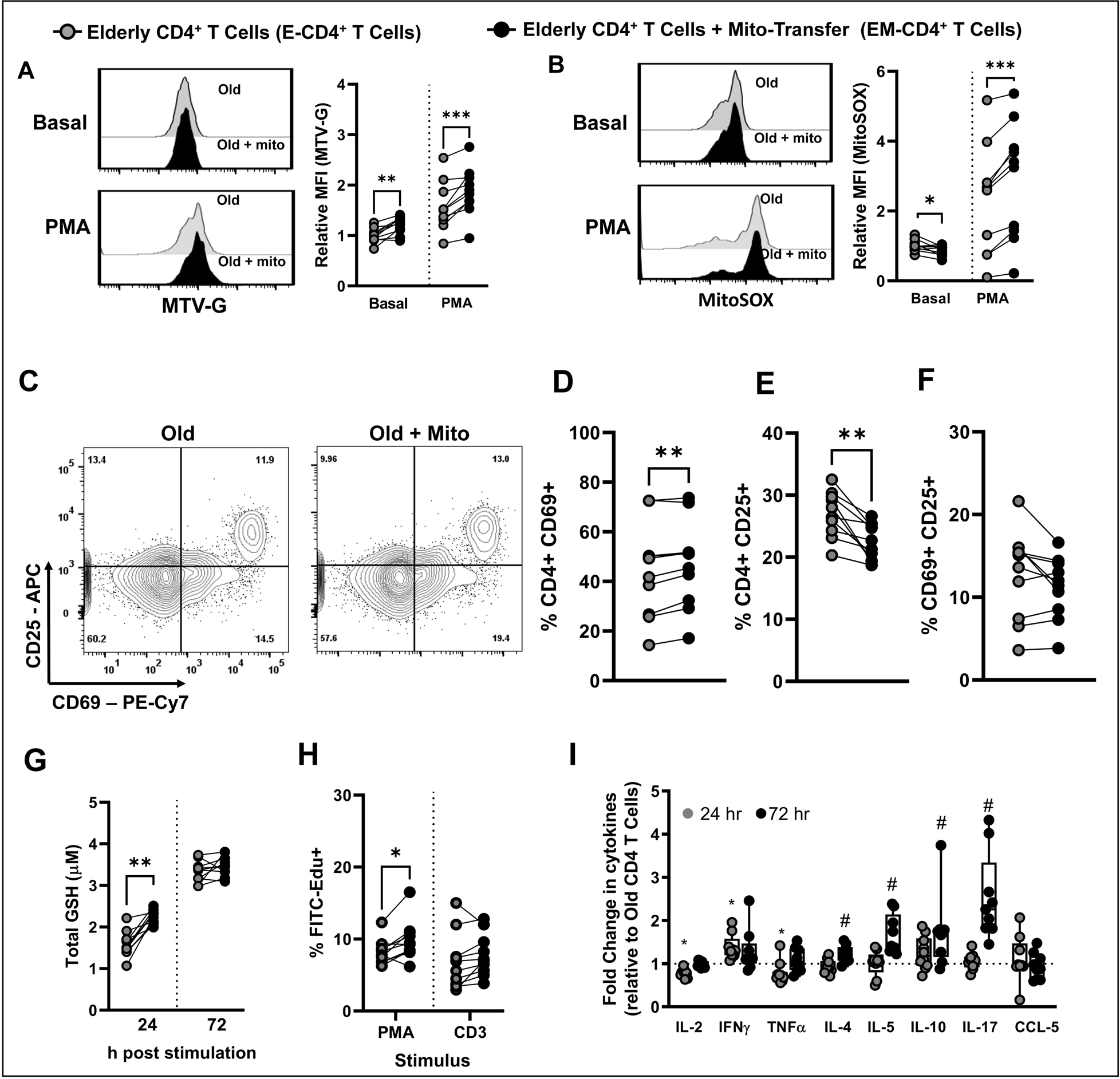
T cell activation and proliferation improved in CD4+ T cells from old humans after mito-transfer. Donor mitochondria isolated from primary human neonatal dermal fibroblasts were transplanted into CD4^+^ T cells from elderly humans. Flow cytometry histograms and the relative fold changes of **A)** MTV-G, **B)** MitoSOX stained, unstimulated, or PMA/Ionomycin stimulated human CD4^+^ T cells with or without mito-transfer. **C)** Contour plots of CD25 v CD69 expression on CD4^+^ T cells with or without mito-transfer, after PMA/Ionomycin stimulation (24h). The percentages of **D)** CD69^+^, **E)** CD25^+^, and **F)** CD69^+^CD25^+^ human CD4+ T cells after PMA/Ionomycin stimulation (24h). **G)** Total GSH produced by CD4^+^ T cells with or without mito-transfer, stimulation with PMA/ Ionomycin, at 24h and 72 h. **H)** The number of EdU+ cells in PMA/Ionomycin-stimulated human CD4^+^ T cell cultures with or without mito-transfer. **I**) The relative fold change in cytokine production of human CD4^+^ T cells with or without mito-transfer, after stimulation with PMA/Ionomycin for 24 h and 72 h. 9-10 biological replicates per group with p ≤ 0.05 = *, p ≤ 0.01 = **, or p ≤ 0.001 = *** using paired Student’s *t*-test.

Next, the impact of acute (24h stimulation) on E-CD4^+^ and EM-CD4^+^ T cells was examined. Compared to E-CD4^+^ T cells, EM-CD4^+^ T cells had higher frequencies (%) of CD69^+^ expression **(Fig. 5. C-D)**, suggesting that mito-transfer improved T cell activation. However, the % of CD25^+^ cells significantly decreased **(Fig. 5. E)**, and there were no differences in the frequency of CD69^+^ CD25^+^ cells between treatment groups **(Fig. 5. F).** The total glutathione (GSH) production from E-CD4^+^ and EM-CD4^+^ T cells was also compared at 24 and 72 h post stimulation; EM-CD4^+^ T cells had significantly higher GSH production after 24 h of stimulation, indicating an improved initial cellular antioxidant capacity (**Fig. 5. G**). The proliferative capacity of E-CD4^+^ and EM-CD4^+^ T cells were also briefly assessed by EdU-incorporation (**Fig. 5. H**). EM-CD4^+^ T cells had significantly higher % of FITC-EdU^+^ cells after stimulation with PMA/IONO, with a similar trend noted between CD3/CD28 stimulated-EM-CD4^+^ and E-CD4^+^ T cells (**Fig. 5. H**). Collectively, these data suggest that mito-transfer may modulate elderly CD4^+^ T cell proliferation.

The levels of soluble cytokines (IL-2, IFNɣ, TNF, IL-4, IL-5, IL-10, IL-17) and the chemokine CCL-5, in the culture media of activated E-CD4^+^ and EM-CD4^+^ T cells were measured after 24 h and 72 h of stimulation (**Fig. 5. I, Supp. Fig. 1.**). Relative to E-CD4^+^ T cells, the supernatant of EM-CD4^+^ T cells contained significantly lower amounts of IL-2, and TNF, and higher amounts of IFNɣ after 24 h stimulation with PMA/ionomycin (**Fig. 5. I, Supp. Fig. 1.**). No other differences were noted for the cytokines detected at 24 h post stimulation. After 72 h stimulation of PMA/ionomycin, the supernatant of EM-CD4^+^ T cells contained significantly higher amounts of IL-4, IL-5, IL-10, and IL-17 (**Fig. 5. I, Supp. Fig. 1.**). These data suggest that mito-transfer can alter cytokine production in elderly CD4^+^ T cells.

### 3.5 Mito-transfer reduced the expression of exhaustion and senescence markers on elderly CD4^+^ T cells

To further investigate the effects of mito-transfer on the functionality of elderly human CD4^+^ T cells, we evaluated their viability and proliferation following two distinct stimulation strategies: single stimulation (Single stim) and continuous stimulation (Cont. stim) **(Supp. Fig 2.)**. In the Single stim experiment CD4^+^ T cells were activated once via CD3/CD28 crosslinking, after which the cell viability and doubling rate were tracked for up to 14 days post-activation **(Supp. Fig 2. A-B).** Except for a noticeable divergence at the 10-day mark, there were no major differences in T cell proliferation rates **(Supp. Fig 2. A)**. However, the viability of E-CD4^+^ T cells decreased significantly after 7 days, with viable T cells from only 6 donors remaining by day 14 **(Supp. Fig 2. B).** In the *cont. stim* experiments a different trend was observed **(Supp. Fig. 2. C-D)** where the doubling rates, and the viability of both E-CD4^+^ and EM-CD4^+^ T cells remained comparable up to 14 days post-activation **(Supp. Fig. 2. C-D)**. These data suggest that mito-transfer may enhance the survivability of elderly CD4^+^ T cells during T cell activation and division, after initial encounter with cognate antigens.

Because viability does not exclude the presence of dysfunctional CD4^+^ T cells, the CD4^+^ T cells were phenotyped for markers broadly associated with T cell memory (CD45RO, CD62L), activation (CD69), exhaustion (PD1^+^, and co-expression of TIM3^+^, LAG3^+^), terminal differentiation (CD28, CD27, CD57, KLRG1), senescence (β-gal, and p16), proliferation (Ki67) and DNA damage (ɣH2AX), at 7 days after initial activation **(Fig. 6. A-N)**.

**Fig. 6.**
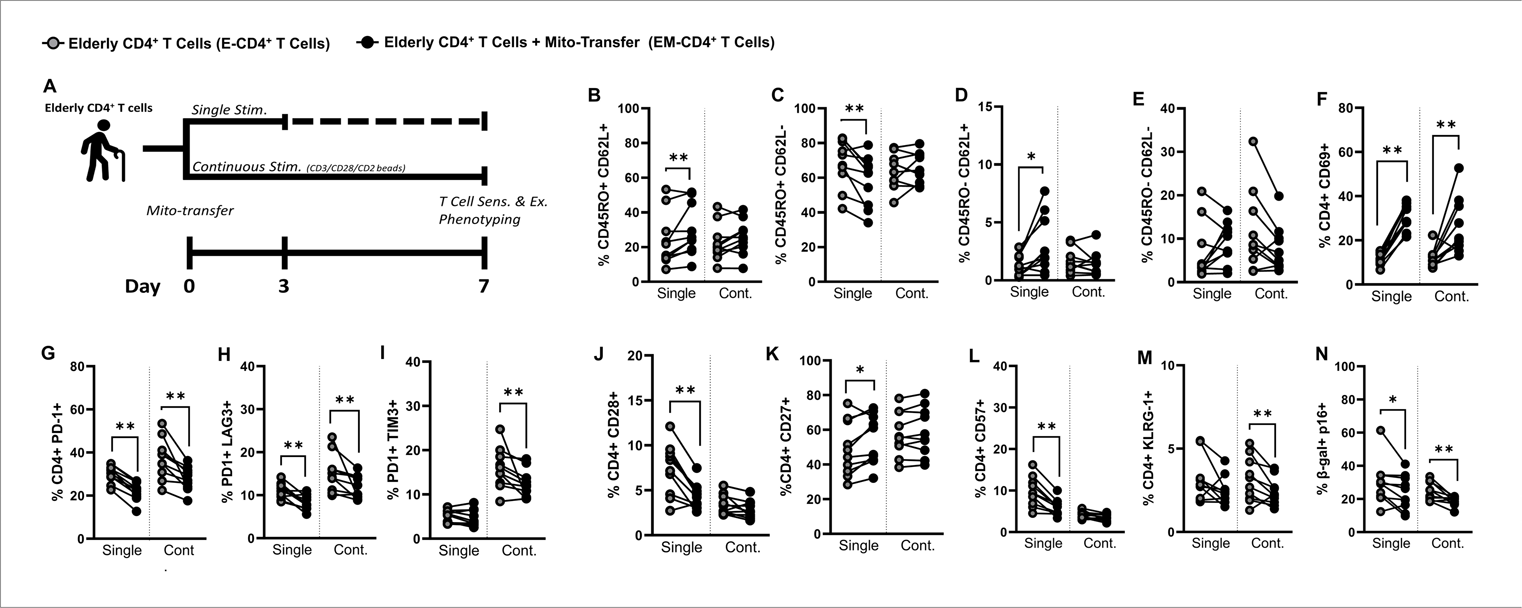
Mito-transfer phenotype and functionality recapitulated in aged human CD4 T cells. **A)** Elderly CD4 T cells with or without mito-transfer were activated via surface cross-linking (CD3/CD28). CD4 T cells either received a single (single) or continuous (cont.) stimulation. The percentages of **B)** CD45RO+ CD62L+, **C)** CD45RO+ CD62L-, **D)** CD45RO-CD62L-, **E)** CD45RO-CD62L-, **F)** CD69+, **G)** PD-1+, **H)** PD-1+ LAG3+, **I)** PD-1+ TIM3+, **J)** CD28+, **K)** CD27 +, **L)** CD57+, **M)** KLRG1+, and **N)** β-Gal-FITC, p16+ human CD4+ T cells at 7 days after intial activation with CD3/CD28/CD2. 10biological replicates per group, with p ≤ 0.05 = *, p ≤ 0.01 = **, using paired Student’s t-test.

In the experimental groups that received a single stimulation (***single stim***), compared to E-CD4^+^ T cells, EM-CD4^+^ T cells had significantly higher frequencies (%) of CD45RO^+^ CD62L^+^ cells **(Fig. 6. B)**, and lower % of CD45RO^+^ CD62L^-^ cells **(Fig. 6. C)**. Compared to E-CD4^+^ T cells, EM-CD4^+^ T cells had increased % of CD45RO^-^ CD62L^+^ cells **(Fig. 6. D),** but comparable % of CD45RO^-^ CD62L^-^ cells **(Fig. 6. E).** Collectively, these data suggest that mito-transfer modulated the expansion of memory T cells.

Compared to E-CD4^+^ T cells, EM-CD4^+^ T cells had a higher % of recently activated (CD69^+^) cells (**Fig. 6. F)**, and a lower frequency of exhausted (PD1^+^, PD1^+^ LAG3^+^) cells **(Fig. 6. G-I).** Regarding markers associated with terminal differentiation, compared to E-CD4^+^ T cells, EM-CD4^+^ T cells had a lower % of CD28^+^ cells **(Fig. 6. J)**, a higher % of CD27^+^ cells **(Fig. 6. K)**, comparable levels of CD28- CD27- cells **(Supp. Fig. 3. A)**, lower % of CD57^+^ cells **(Fig. 6. L)**, and comparable amounts of KLRG1^+^ cells **(Fig. 6. M)**. Regarding senescence, compared to E-CD4^+^ T cells, EM- CD4^+^ T cells had significantly lower % of β-gal^+^ cells **(Supp. Fig. 3. B)** and β-gal^+^ p16^+^ cells **(Fig. 6. N)**. E-CD4^+^ and EM-CD4^+^ T cells had comparable % of Ki67^+^ **(Supp. Fig. 3. D)** and ɣH2AX^+^ cells **(Supp. Fig. 3. E).**

In the Cont. stim experimental groups, at 7 days post initial stimulation, E-CD4^+^ T cells and EM- CD4^+^ T cells had comparable % of memory markers subsets (**Fig. 6. B-E).** Compared to E-CD4^+^ T cells, EM-CD4^+^ T cells had a higher % of recently activated CD69^+^ cells (**Fig. 6. F)**, and lower % of exhausted (PD1^+^, PD1^+^ LAG3^+^, PD1^+^ TIM3^+^) cells **(Fig. 6. G-I).** Regarding markers associated with terminal differentiation, except for lower % of KLRG1^+^ cells **(Fig. 6. M)**, all other marker subsets examined were at comparable levels **(Fig. 6. J-M)**. Regarding senescence, compared to E-CD4^+^ T cells, EM-CD4^+^ T cells had significantly lower % of β-gal^+^ **(Supp. Fig. 3. B)** and β-gal^+^ p16^+^ cells **(Fig. 6. N)**. However, there was no difference in the % of Ki67^+^ cells **(Supp. Fig. 3. D)** and ɣH2AX^+^ **(Supp. Fig. 3. E)** cells between E-CD4^+^ and EM-CD4^+^ T cells.

Since the viability of E-CD4^+^ and EM-CD4^+^ T cells remained comparable under continuous stimulation at 14 days, the cells were also phenotyped for exhaustion and senescence. Compared to E-CD4^+^ T cells, EM-CD4^+^ T cells had a higher % of CD45RO^+^ CD62L^+^ (**Supp. Fig. 3. F)** and a lower % of CD45RO^+^ CD62L^-^ cells **(Supp. Fig. 3. G)** after 14 days of continuous stimulation. Both E-CD4^+^ and EM-CD4^+^ T cells had negligible amounts of CD45RO^-^ CD62L^-^ (**Supp. Fig. 3. H)** and CD45RO^-^ CD62L^+^ cells **(Supp. Fig. 3. I)**. EM-CD4^+^ T cells still displayed higher levels of CD69^+^ **(Supp. Fig. 3. J)**, along with fewer exhausted **(Supp. Fig. 3. K-M)** and senescent **(Supp. Fig. 3. Q -U)** cells. These (Single and Cont. stim) data collectively indicate that mito-transfer significantly improved the viability and activation of elderly CD4^+^ T cells, and significantly reduced T cell exhaustion and senescence ex vivo.

## 4. Discussion

The elderly population is at heightened risk for both developing TB disease as well as having TB treatment complications [67–70]. Underlying immune dysfunctions in this demographic contribute to increased *M.tb* burden and mortality [14–17]. Both aging and chronic infections like TB are associated with increased T cell exhaustion and senescence, conditions that impair the generation of protective effector and memory responses [22, 24–26, 36, 71–74]. Central to T cell exhaustion and senescence is mito-dysfunction [24, 35, 39–43] and the potential of mito-transfer to mitigate aging-associated T cell dysfunction was investigated. First, we leveraged bioinformatics (MetaScape) to survey cellular pathways that change in aged CD4^+^ T cells after mito-transfer. MetaScape analysis revealed that prominent pathways associated with T cell exhaustion and senescence were downregulated in CD4^+^ T cells that received mito-transfer. Second, whether mito-transfer impacted the differentiation of aged naive CD4^+^ T cells in *M.tb* infected mice was assessed. Our data show that mito-transfer reduced CD4^+^ T cell exhaustion in *M.tb* infected mice. Lastly, the impact of mito-transfer in elderly human CD4^+^ T cells *was* examined (ex vivo) to evaluate the biological relevance of our findings for humans. Mito-transfer increased T cell activation and cytokine production in elderly CD4^+^ T cells and decreased the expression of markers associated with T cell exhaustion and senescence. Our collective data re-emphasize that targeting mitochondrial health has significant potential in boosting immune responses and improving chronic infections, like TB and others in the elderly.

### 4.1. Bioinformatic analysis

Bioinformatic analysis using MetaScape revealed a significant impact of mito-transfer on pathways linked to T cell exhaustion. Beyond the expected upregulation of pathways related to mitochondrial function, the upregulation of several indirect and unrelated pathways, particularly those associated with broader cellular metabolism (e.g., steroid, amino, and carbon metabolism) and protein processing (e.g., ER stress, protein hydroxylation) were identified. These findings align with our earlier studies that reported on the increased metabolic activity in mito-transferred aged CD4^+^ T cells. These data also suggest the induction of a metabolic reprogramming or stress response in CD4^+^ T cells following mito-transfer. Moreover, the concurrent downregulation of pathways associated with RNA metabolism, glycolysis, apoptosis, and immune responses, including to IL-7 signaling, suggests an energy shift from glycolysis to oxidative phosphorylation [75]. For instance, previous work shows that IL-7 signaling promotes glycolysis in T cells [75, 76]. IL-7 signaling is also associated with the expression of the anti-apoptotic protein Bcl-2 [77]. MCODE cluster analysis of downregulated Tex-DEPs also revealed a reduction in apoptotic signaling. Nevertheless, future studies are needed enumerate the full impact of mito-transfer on the IL-7 signaling axis. The MetaScape analysis also hinted at mito-transfer alleviating cellular senescence. The downregulated Sen-DEPs in mito-transferred CD4^+^ T from old mice, were largely related to cellular stress responses (hypoxia/HIF1α, DNA damage, and ROS), glycolytic metabolism, pro-inflammatory immune signaling, including IL-6, which are all well characterized hyperactive pathways in (pathogenic) senescent cells [78, 79]. These analyses foreshadowed that mito-transfer may shift cellular functionality including anti-senescence reprogramming, which was confirmed in the elderly human CD4^+^ T cell experiments.

### 4.2. *In vivo* mouse experiments on T cell differentiation

Our previous studies demonstrated improved control of *M.tb* and delayed mortality from IAV in RAG-KO mice receiving mito-transferred naïve CD4^+^ T cells. Having identified significant alterations in signaling pathways in mito-transfer in CD4^+^ T cells from old mice *ex vivo*, examining the *in vivo* implications of these changes was a pragmatic next step. In this study, T cell differentiation during *M.tb* infection was examined. A primary observation was the reduction of CD4^+^ T cell exhaustion markers during *M.tb* infection. Specifically, at 30 days post infection, a reduction in CD4^+^ TIM-3^+^ T cells was observed in the OM-TB-TKO group. At 60 days post-infection, these mice also showed increased frequencies of CD4^+^ T cells with activation (CD25^+^, CD69^+^ CD25^+^) and effector memory (KLRG1^+^) markers in the lungs. In the spleen of OM-TB-TKO and Y-TB-TKO mice, similar % of activated CD4^+^ T cells were observed. At the 60-day mark in the spleen OM-TB-TKO mice, there were significantly lower frequencies of activated (CD69^+^, CD25^+^CD69^+^), memory (PD1^+^CD27-), and exhausted (TIM3^+^) CD4^+^ T cells. Therefore, we observe an association between changes in senescence and exhaustion markers and the ability to control *M.tb* infection *in vivo* in mice.

### 4.2. Elderly Human CD4 T cells after mito-transfer

An early observation was the mitochondrial mass of elderly T cell significantly increased after mito-transfer, as evidenced by the relative MFI of MTV-G in elderly cells with and without mito-transfer. This increase was apparent in both unstimulated and stimulated conditions. Concurrently, mito-transfer appeared to modulate the production of mitoROS, with decreased mitoROS observed in unstimulated T cells, and an increase was noted in stimulated conditions, suggesting an improved response to activation signals. This is also in line with our data from mouse studies [44]. A significant improvement in T cell activation following mito-transfer, evidenced by increased frequency of CD69^+^ expression in mito-transferred elderly CD4^+^ T cells was also noted. The proliferative capacity and antioxidant defense of mito-transferred elderly CD4^+^ T cells also improved. Following mito-transfer, both the total glutathione (GSH) production and the frequency of EdU^+^ cells increased significantly, pointing towards a rejuvenated proliferative capability of elderly CD4^+^ T cells.

In reference to work by Callender *et al.,* CD4^+^ T cell senescence may be potentially dampened by increasing CD4 T cell mitochondrial mass and activity. Callender and team demonstrated that CD4^+^ T cells have higher mitochondrial mass than CD8^+^ T cells from the same individual. CD4^+^ T cells also exhibited greater metabolic versatility and retain higher functional capacities, leading to a slower rate of senescence, and this was attributed to increased mitochondrial mass [40]. Our data somewhat agrees with this notion, as we also observed a correlation between increased mitochondrial mass and improved functionality in CD4^+^ T cells [40]. After mito-transfer, there was a significant increase in the mitochondrial mass of elderly CD4^+^ T cells, reduced signs of exhaustion and senescence, as well as subsequent improvements in T cell activation and viability. Although we enhanced mitochondrial mass in CD4^+^ T cells, we would not anticipate functional improvements if dysfunctional mitochondria were transferred. Studies examining the impact of intercellular mito-transfer, modulated by tube-nano-tunnels [80], and engulfment of dysfunctional mitochondria [81], demonstrate that immune cell function worsened [80, 81]. This reinforces that the quality of donated mitochondria is equally important in the mitochondrial mass dependent plasticity of aged CD4^+^ T cells and other immune cells.

The cytokine data further suggest improvements in elderly T cell function. The levels of several soluble cytokines (IL-2, IL-4, IL-5, Il-10, IL-17, IFNɣ and TNFα) in the culture media of activated non-manipulated and mito-transferred elderly CD4^+^ T cells varied at different time points post-stimulation. Particularly, the increases in IL-4, IL-5, and IL-17, alongside alterations in IL-10, could reflect a complex immunomodulatory effect induced by mito-transfer. This might include a shift towards a Th2 response, potential regulation of inflammatory processes, and alterations in T cell activation and proliferation states. However, the experimental conditions, specifically the *in vitro* setting with non-specific PMA/Ionomycin stimulation, could have markedly influenced these outcomes. Such stimulation does not fully replicate the antigen-specific immune responses *in vivo* and may exaggerate certain pathways, leading to a cytokine profile that might not entirely represent physiological conditions. Therefore, we remain cautious when interpreting the consequences of these changes.

The reduction in naïve T cell proportions (CD45RO- CD62L+) and an increase in effector memory T cells (CD45RO+ CD62L-) after continuous stimulation in cells receiving young mitochondria suggest that these cells are possibly undergoing enhanced differentiation. However, the reduction in CD27 expression post-transplantation is indicative of a more complex effect, as CD27 is crucial for T cell survival and long-term memory. A potential interpretation is that while mito-transfer aids in the short-term activation and function of aged T cells, it may also prompt these cells towards a more differentiated and potentially terminal phenotype. This may be beneficial for immediate immune responses, but long-term impact of mito-transfer on maintenance of the T cell population warrants further investigation. Nevertheless, the decrease in exhaustion markers such as PD-1, LAG-3, and TIM-3, coupled with a reduction in senescence markers CD57 and KLRG1, supports the notion that mitochondrial transplantation confers a rejuvenated phenotype on aged T cells. This is further supported by the downregulation of P16 and SA-β-gal, markers typically upregulated in senescent cells [82, 83].

The increase in CD69 expression in mito-transferred cells suggests that cells have an enhanced activation profile, which could translate into improved T cell-mediated immune responses. This is significant given the common decline in T cell function with age. Furthermore, the data points to the potential of mitochondrial transplantation to mitigate the effects of continuous stimulation, which typically leads to T cell exhaustion. Such improvements could have significant implications for bolstering the immune system in the elderly, potentially improving responses to infections and vaccination. While these *in vitro* results are promising, the true physiological impact of mitochondrial transplantation must be validated *in vivo*. Further investigation is necessary to ascertain the longevity of these effects, the mechanisms underlying the observed changes, and whether such an approach could be feasibly translated into clinical therapies aimed at countering age-related immune decline.

In summary, this study re-emphasizes the pivotal role that mitochondrial health holds in the viability and performance of CD4^+^ T cells, and how it can be manipulated through mito-transfer to enhance their function, particularly in the elderly population. The approach of mitochondrial health and activity in these cells presents the potential for a slower senescence rate, decrease in T cell exhaustion, and possibly reprogramming their immune response towards less inflammation. In the context of chronic infections, our studies demonstrate proof of principle that mito-transfer, at minimum, can transiently reduce T cell exhaustion during active *M.tb* infection. Further, combinatorial use of mito-transfer and antimycobacterials may also lead to accelerated clearance of infections in the elderly. Therefore, the potential of mito-transfer to reduce immune exhaustion in other chronic conditions, like HIV/AIDS and cancer, warrants exploration. Ultimately, mito-transfer may lead to improved health and longevity for the elderly by bolstering their immune system’s response to infections and possibly other immune-related conditions.

Collectively, these findings suggest that mito-transfer may ameliorate some aspects of immunosenescence in elderly CD4^+^ T cells, potentially restoring a more ‘youthful’ immune functionality. These improvements could have significant implications for bolstering the immune system in the elderly, potentially improving responses to infections and vaccination. Nonetheless, while these *ex vivo* and *in vivo* results are promising, further investigation is necessary to ascertain the longevity of these effects, the mechanisms underlying the observed changes, and whether such an approach could be feasibly translated into clinical therapies aimed at countering age-related immune decline.

## Acknowledgments

We would like to acknowledge the thoughtful discussions with Drs. Larry S. Schlesinger, Douglas Green and James Stambulli which have strengthened this manuscript. We would also like to acknowledge Drs. Gourav Choudhury and Marcel Daadi for sharing their MEF isolation protocol.

## Author Contributions

This project was conceived by CAH & JT; CAH designed the experiments; CAH, SG, AOF, AV, VD, AS performed experiments; CAH & SW performed proteomic analysis, CAH performed the statistical analysis, CAH and JT wrote the manuscript; JT provided ongoing mentoring to refine the studies, CAH, PS, JBT and JT edited the manuscript.

## Data Availability

Authors declare that the data supporting the findings of this study are available within the paper and its supplementary information files. Reagents are available upon request.

---

**Supp. Fig. 1.**
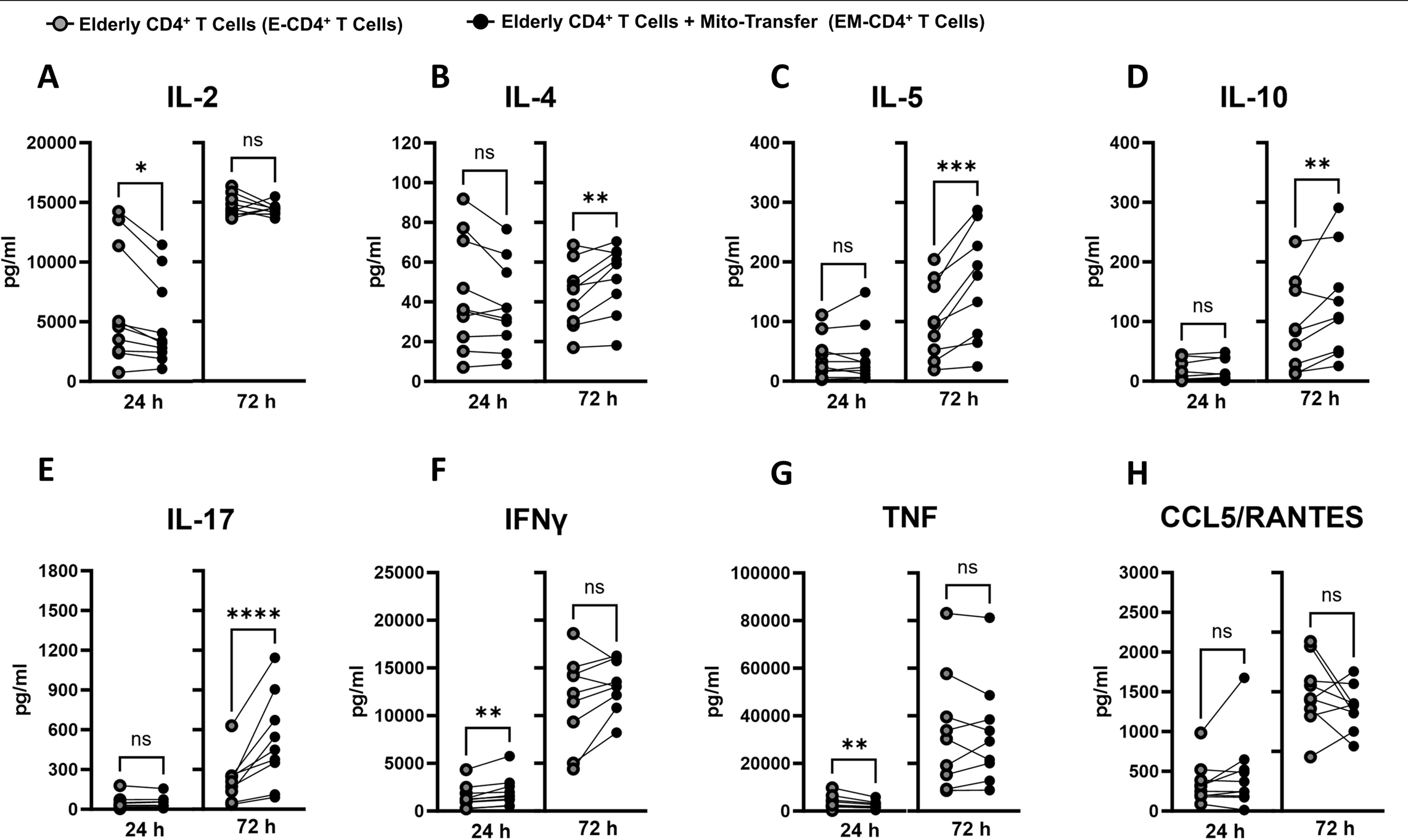
Mito-transfer alters cytokine production of CD4+ T cells in aged humans. CD4+ T cells from aged humans with or without mito-transfer were stimulated with PMA/Ionomycin. After 24h and 72h stimulation with PMA/Ionomycin, the supernatants were examined by Luminex array for cytokines produced. 9-10 biological replicates per group with p ≤ 0.05 = *, p ≤ 0.01 = **, or p ≤ 0.001 = *** using paired Student’s *t-*test.

**Supp. Fig 2.**
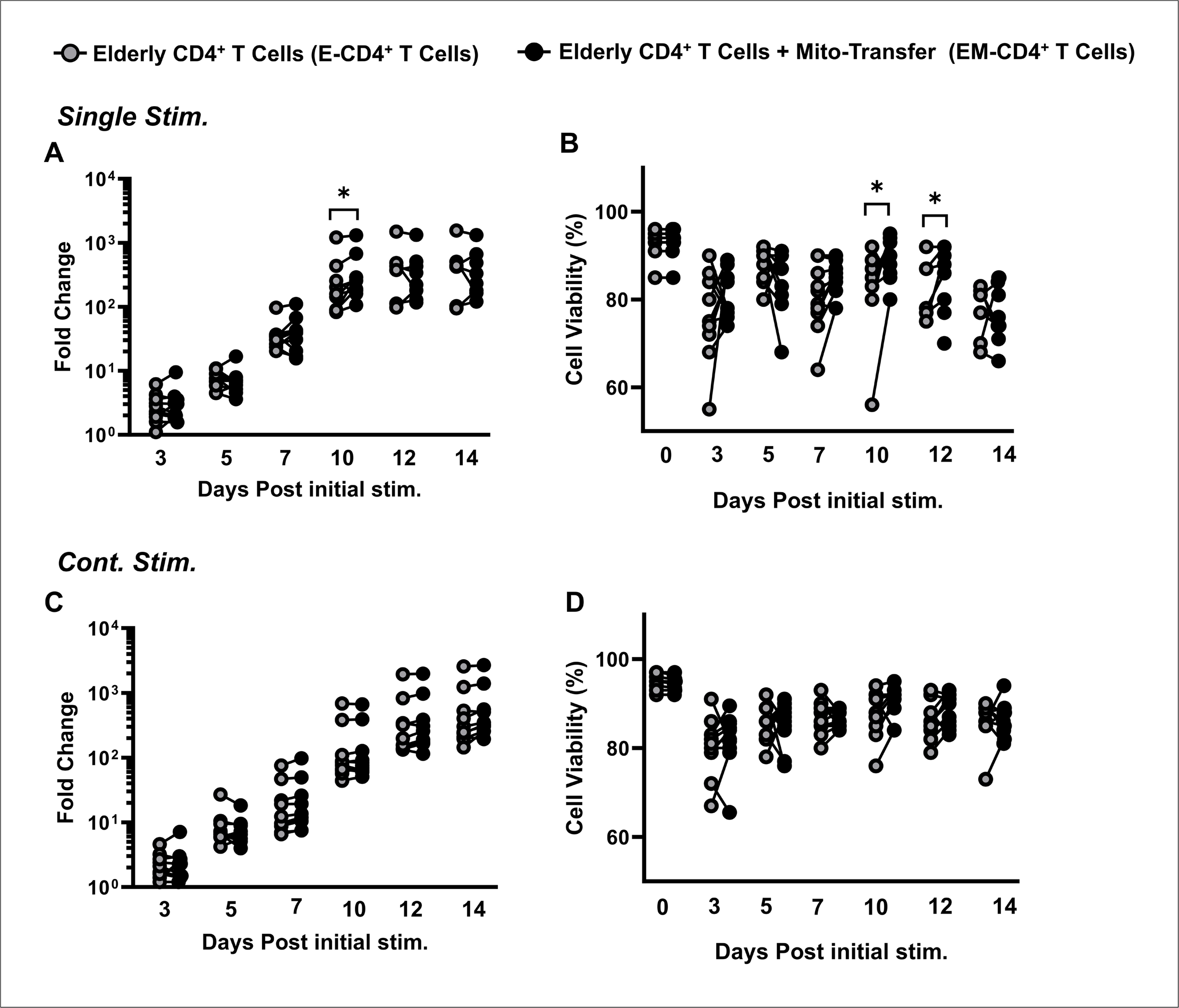
Impact of mito-transfer on elderly T cell proliferation and viability. Elderly CD4 T cells with or without mito-transfer were activated via surface cross-linking (CD3/CD28). CD4 T cells either received a single (single) or continuous (cont.) stimulation. **A)** Fold change and **B)** viability of CD4+ T cells with or without mito-transfer after single stimulation. **C)** Fold change and **D)** viability of CD4+ T cells with or without mito-transfer with continuous stimulation. 10 biological replicates per group, with p ≤ 0.05 = *, p ≤ 0.01 = **, using paired Student’s *t*-test.

**Supp Fig 3.**
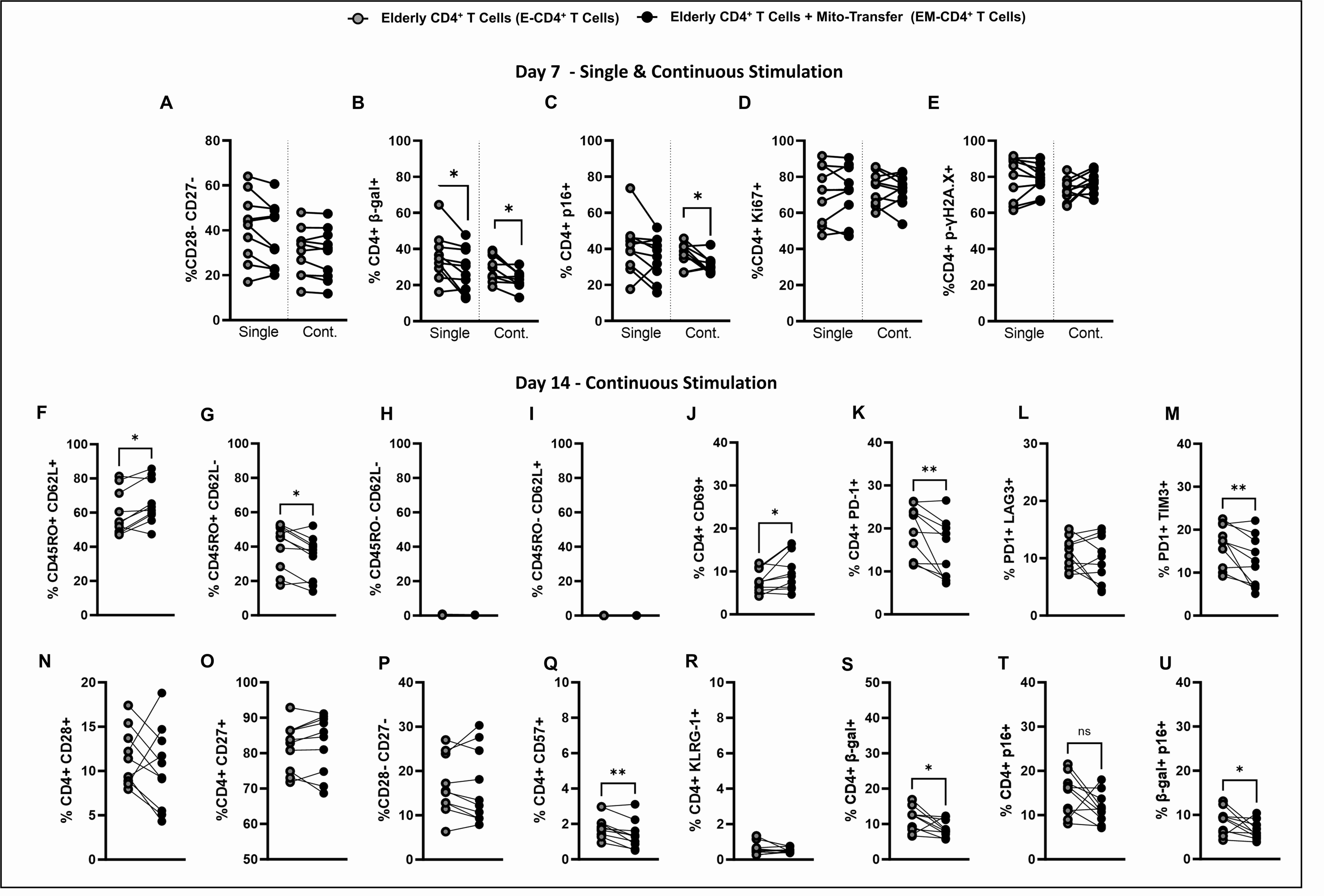
Mito-transfer improves T cell activation and reduces human T cell exhaustion and senescence. Donor mitochondria isolated from primary human neonatal dermal fibroblasts were transplanted into CD4+ T cells from elderly humans. Elderly CD4^+^ T cells with or without mito-transfer were activated via surface cross-linking (CD3/CD28), T cells either received a single (single) or continuous (cont.) stimulation. The percentages of **A)** CD28- CD27, **B)** β-gal+, **C)** p16+, **D)** Ki67+, and **E**) ɣH2A.X+ human CD4+ cells at 7 days after intial activation with CD3/CD28/CD2. The percentages of **F**) CD45RO+ CD62L+, **G**) CD45RO+ CD62L, **H**) CD45RO- CD62L-, **I**) CD45RO- CD62L-, **J**) CD69+, **K**) PD-1+, **L**) PD-1+ LAG3+, **M**) PD-1+ TIM3+, **N**) CD28+, **O**) CD27+, **P**) CD28- CD27, **Q**) CD57+, **R**) KLRG1+, and **S**) β-Gal+, **T**) p16+, **U**) β-gal+ p16+ human CD4+ T cells at 14 days after continuous stimulation with CD3/CD28/CD2. 10 biological replicates per group, with p ≤ 0.05 = *, p ≤ 0.01 = **, using paired Student’s t-test.

