## Supplemental Tables 1 - 12 for "Mitochondrial Transplantation promotes protective effector and memory CD4^+^ T cell response during *Mycobacterium tuberculosis* infection and diminishes exhaustion and senescence in elderly CD4^+^ T cells"

| Uniprot ID | Gene Symbol | t-test | FDR (adj. P-val.) | O1 | O2 | O3 | OM1 | OM2 | OM3 |
|---|---|---|---|---|---|---|---|---|---|
| P35564 | Canx | 0.000 | 0.009 | 0.244 | 0.000 | -0.086 | 1.700 | 1.720 | 1.860 |
| P56959 | Fus | 0.739 | 1.000 | -0.231 | 0.140 | 0.000 | -0.070 | -0.107 | 0.255 |
| Q01147 | Creb1 | 0.091 | 0.232 | -0.101 | 0.000 | 0.267 | -0.384 | -0.333 | -0.078 |
| B2RQC6 | Cad | 0.675 | 0.952 | 0.244 | 0.000 | -0.142 | 0.662 | -0.451 | -0.695 |
| O09117 | Sypl1 | 0.366 | 0.613 | 0.244 | 0.000 | -0.323 | 0.744 | -0.281 | 0.538 |
| Q9JIK9 | Mrps34 | 0.049 | 0.154 | 0.000 | 0.757 | -0.497 | 1.210 | 1.140 | 1.010 |
| Q8K1Z0 | Coq9 | 0.032 | 0.117 | 0.000 | 0.186 | -0.023 | 0.862 | 0.388 | 0.472 |
| P12265 | Gusb | 0.016 | 0.079 | 0.311 | 0.000 | -0.297 | 0.822 | 0.616 | 0.837 |
| P0C0S6 | H2afz | 0.384 | 0.633 | -0.205 | 0.000 | 0.096 | -0.364 | -0.361 | 0.097 |
| Q3THW5 | H2afv | 0.384 | 0.633 | -0.205 | 0.000 | 0.096 | -0.364 | -0.361 | 0.097 |
| Q8BM55 | Tmem214 | 0.001 | 0.020 | 0.000 | 0.450 | -0.101 | 1.560 | 1.490 | 1.600 |
| P70227 | Itpr3 | 0.573 | 0.843 | -0.451 | 0.154 | 0.000 | 0.017 | -0.052 | 0.078 |
| O08579 | Emd | 0.292 | 0.518 | -0.552 | 0.000 | 0.097 | 0.094 | -0.100 | 0.655 |
| Q9CQ60 | Pgls | 0.384 | 0.633 | 0.000 | 1.190 | -0.787 | -0.304 | -2.870 | 0.284 |
| Q8K2K6 | Agfg1 | 0.896 | 1.000 | -0.030 | 0.000 | 1.700 | 0.753 | 0.090 | 1.090 |
| P97450 | Atp5j | 0.033 | 0.121 | 0.000 | 0.125 | -0.630 | 1.010 | 0.420 | 1.140 |
| O55022 | Pgrmc1 | 0.000 | 0.010 | 0.000 | -0.086 | 0.009 | 1.480 | 1.240 | 1.540 |
| Q9D0K2 | Oxct1 | 0.000 | 0.012 | 0.000 | 0.092 | -0.243 | 1.760 | 1.440 | 1.670 |
| Q3UZ39 | Lrrfip1 | 0.003 | 0.030 | 0.000 | -0.058 | 0.148 | -0.914 | -0.930 | -0.588 |
| P52825 | Cpt2 | 0.312 | 0.545 | 0.000 | 0.208 | -0.370 | 1.210 | 0.043 | 0.051 |
| P70460 | Vasp | 0.139 | 0.305 | 0.006 | -0.256 | 0.000 | -0.109 | -0.539 | -0.494 |
| Q62422 | Ostf1 | 0.025 | 0.102 | 0.488 | -0.040 | 0.000 | -0.345 | -0.835 | -0.846 |
| Q8BMK4 | Ckap4 | 0.000 | 0.010 | 0.093 | 0.000 | -0.598 | 3.600 | 3.310 | 3.710 |
| Q8BFY9 | Tnpo1 | 0.030 | 0.112 | -0.077 | 0.000 | 0.000 | -0.433 | -1.300 | -0.849 |
| Q9D6J6 | Ndufv2 | 0.049 | 0.154 | 0.000 | 0.342 | -0.183 | 0.718 | 0.363 | 0.856 |
| P51125 | Cast | 0.847 | 1.000 | -0.076 | 0.000 | 0.118 | 0.232 | -0.209 | -0.068 |
| P06800 | Ptprc | 0.199 | 0.394 | -0.090 | 0.130 | 0.000 | -0.302 | -0.348 | 0.046 |
| Q9Z277 | Baz1b | 0.955 | 1.000 | -0.231 | 0.238 | 0.000 | 0.030 | -0.197 | 0.206 |
| Q9CWL8 | Ctnnbl1 | 0.003 | 0.033 | 0.000 | 0.296 | -0.044 | -0.669 | -1.050 | -0.968 |
| Q7TPR4 | Actn1 | 0.680 | 0.956 | 0.455 | 0.000 | -0.439 | 0.662 | -0.251 | 0.099 |
| P35979 | Rpl12 | 0.491 | 0.755 | 0.266 | -0.151 | 0.000 | 0.271 | -0.094 | 0.359 |
| O09167 | Rpl21 | 0.336 | 0.573 | 0.282 | -1.900 | 0.000 | 0.415 | -0.668 | 2.110 |
| Q8R081 | Hnrnpl | 0.025 | 0.101 | -0.126 | 0.000 | 0.040 | -0.348 | -0.636 | -0.308 |
| P70315 | Was | 0.034 | 0.122 | 0.000 | -0.169 | 0.494 | -0.488 | -1.180 | -0.715 |
| Q91YH5 | Atl3 | 0.038 | 0.130 | -0.218 | 0.589 | 0.000 | 0.827 | 0.935 | 0.843 |
| P14824 | Anxa6 | 0.623 | 0.898 | -0.261 | 0.274 | 0.000 | -0.273 | -0.143 | 0.121 |
| Q99L45 | Eif2s2 | 0.540 | 0.807 | 0.342 | -0.259 | 0.000 | 0.266 | -0.631 | -0.179 |
| Q9Z2D6 | Mecp2 | 0.063 | 0.181 | -0.130 | 0.000 | 0.020 | -0.393 | -0.654 | -0.177 |
| Q9D1A2 | Cndp2 | 0.006 | 0.043 | 0.000 | 0.087 | -0.305 | -0.761 | -1.170 | -1.080 |
| Q6IRU2 | Tpm4 | 0.699 | 0.975 | 0.000 | 0.258 | -0.242 | 0.139 | -0.352 | -0.025 |
| Q9JKF1 | Iqgap1 | 0.611 | 0.885 | 0.080 | 0.000 | -0.164 | 0.186 | -0.420 | -0.164 |
| O08734 | Bak1 | 0.010 | 0.060 | 0.000 | 0.028 | -0.547 | 0.687 | 0.764 | 0.655 |
| Q9DBJ1 | Pgam1 | 0.020 | 0.088 | 0.582 | -0.030 | 0.000 | -0.473 | -1.080 | -1.040 |
| P24270 | Cat | 0.046 | 0.148 | 0.177 | 0.000 | -0.253 | 1.040 | 0.403 | 0.494 |
| P47757 | Capzb | 0.016 | 0.077 | 0.000 | 0.178 | -0.196 | -0.410 | -0.725 | -0.604 |
| Q8BK67 | Rcc2 | 0.253 | 0.466 | 0.487 | -0.092 | 0.000 | 0.079 | -0.473 | -0.172 |
| Q8C0G2 | Traf3ip3 | 0.025 | 0.100 | -0.492 | 0.050 | 0.000 | -0.704 | -0.767 | -0.938 |
| Q9CYN2 | Spcs2 | 0.069 | 0.192 | 0.000 | 0.270 | -0.026 | 0.525 | 0.226 | 0.467 |
| Q9D0F9 | Pgm1 | 0.000 | 0.014 | -0.069 | 0.000 | 0.150 | -1.850 | -2.270 | -1.690 |
| Q5SUA5 | Myo1g | 0.101 | 0.247 | 0.000 | -0.027 | 0.074 | -0.171 | -0.701 | -0.196 |
| Q8BJZ4 | Mrps35 | 0.007 | 0.051 | 0.062 | 0.000 | -0.636 | 1.340 | 1.050 | 0.901 |
| Q920B9 | Supt16h | 0.019 | 0.087 | 0.022 | 0.000 | -0.102 | -0.331 | -0.845 | -0.661 |
| D3Z7P3 | Gls | 0.500 | 0.765 | 0.000 | 0.006 | -0.037 | 0.732 | -0.191 | 0.043 |
| Q8BRG8 | Tmem209 | 0.181 | 0.368 | 0.000 | -1.610 | 0.980 | 0.868 | 1.100 | 1.090 |
| Q9DBE8 | Alg2 | 0.000 | 0.012 | 0.004 | 0.000 | -0.115 | 0.889 | 0.810 | 1.040 |
| Q9JIX8 | Acin1 | 0.008 | 0.051 | -0.012 | 0.228 | 0.000 | -0.410 | -0.622 | -0.382 |
| Q9JI39 | Abcb10 | 0.138 | 0.303 | 0.860 | -0.266 | 0.000 | 1.480 | 0.456 | 1.190 |

| | | | | | | | | | |
|---|---|---|---|---|---|---|---|---|---|
| O55201 | Supt5h | 0.014 | 0.073 | 0.000 | -0.043 | 0.190 | -0.500 | -1.070 | -0.621 |
| P47754 | Capza2 | 0.597 | 0.870 | 0.242 | -0.135 | 0.000 | -0.200 | -0.355 | 0.281 |
| Q9R190 | Mta2 | 0.043 | 0.142 | -0.073 | 0.000 | 0.242 | -0.313 | -0.546 | -0.196 |
| Q99PL5 | Rrbp1 | 0.476 | 0.736 | 0.000 | 0.054 | -0.281 | 0.493 | -0.283 | 0.146 |
| Q6DFW4 | Nop58 | 0.101 | 0.246 | 0.101 | -0.080 | 0.000 | -0.065 | -0.661 | -0.403 |
| P26041 | Msn | 0.350 | 0.592 | 0.047 | 0.000 | -0.054 | 0.016 | -0.303 | -0.044 |
| Q9DCT5 | Sdf2 | 0.001 | 0.020 | -0.108 | 0.000 | 0.394 | 2.260 | 1.720 | 1.880 |
| Q9CQJ8 | Ndufb9 | 0.003 | 0.033 | -0.196 | 0.220 | 0.000 | 0.740 | 0.744 | 0.811 |
| P97742 | Cpt1a | 0.015 | 0.076 | 0.000 | 0.296 | -0.452 | 0.925 | 0.756 | 0.903 |
| Q9JKR6 | Hyou1 | 0.008 | 0.053 | 0.225 | 0.000 | -0.131 | 1.020 | 0.600 | 0.825 |
| P13020 | Gsn | 0.025 | 0.102 | 0.242 | 0.000 | -0.363 | 0.994 | 0.554 | 0.643 |
| P11087 | Col1a1 | 0.000 | 0.002 | 0.000 | 0.029 | -0.359 | 7.740 | 7.340 | 7.640 |
| P62264 | Rps14 | 0.453 | 0.711 | 0.151 | -0.252 | 0.000 | 0.120 | -0.562 | -0.229 |
| P70441 | Slc9a3r1 | 0.003 | 0.033 | 0.000 | -0.076 | 0.282 | -1.040 | -1.680 | -1.200 |
| Q9D4J7 | Phf6 | 0.051 | 0.158 | 0.066 | -0.328 | 0.000 | -0.462 | -1.120 | -0.611 |
| P47738 | Aldh2 | 0.815 | 1.000 | 0.398 | 0.000 | -0.923 | 0.773 | -0.269 | -0.602 |
| Q00PI9 | Hnrnpul2 | 0.026 | 0.104 | -0.040 | 0.228 | 0.000 | -0.233 | -0.569 | -0.346 |
| Q8VE37 | Rcc1 | 0.021 | 0.092 | -0.073 | 0.000 | 0.088 | -0.349 | -0.451 | -0.184 |
| Q62351 | Tfrc | 0.848 | 1.000 | 1.330 | 0.000 | -0.491 | 1.130 | -0.397 | -0.350 |
| Q9Z2I8 | Suclg2 | 0.566 | 0.835 | 0.026 | 0.000 | -0.159 | 0.510 | -0.235 | 0.016 |
| Q6P5E4 | Uggt1 | 0.004 | 0.038 | 0.000 | 0.111 | -0.056 | 0.592 | 0.431 | 0.718 |
| P22892 | Ap1g1 | 0.026 | 0.103 | 0.208 | -0.088 | 0.000 | -0.441 | -1.260 | -1.380 |
| P68404 | Prkcb | 0.078 | 0.209 | 0.000 | 0.147 | -0.418 | -0.334 | -0.753 | -0.690 |
| Q9D554 | Sf3a3 | 0.380 | 0.629 | -0.549 | 0.464 | 0.000 | -1.380 | -0.697 | 0.305 |
| Q9DCX2 | Atp5h | 0.025 | 0.101 | 0.000 | 0.466 | -0.336 | 1.070 | 1.120 | 0.719 |
| P20152 | Vim | 0.215 | 0.412 | -0.323 | 0.000 | 0.034 | 0.296 | -0.083 | 0.213 |
| Q3THK7 | Gmps | 0.013 | 0.069 | 0.144 | 0.000 | -0.093 | -0.611 | -1.310 | -0.837 |
| P35550 | Fbl | 0.028 | 0.108 | 0.000 | 0.208 | -0.044 | -0.216 | -0.596 | -0.384 |
| Q8VE22 | Mrps23 | 0.006 | 0.044 | -0.054 | 0.132 | 0.000 | 0.889 | 0.501 | 0.810 |
| P60229 | Eif3e | 0.509 | 0.773 | 0.118 | 0.000 | -0.169 | 0.147 | -0.427 | -0.174 |
| Q569Z6 | Thrap3 | 0.018 | 0.083 | 0.000 | -0.095 | 0.105 | -0.329 | -0.780 | -0.538 |
| P50396 | Gdi1 | 0.000 | 0.008 | -0.087 | 0.000 | 0.016 | -1.240 | -1.480 | -1.310 |
| Q9DAR7 | Dcps | 0.002 | 0.027 | 0.000 | -0.191 | 0.360 | -1.520 | -1.940 | -2.320 |
| Q02053 | Uba1 | 0.001 | 0.019 | -0.136 | 0.000 | 0.019 | -0.926 | -1.240 | -1.310 |
| Q99L43 | Cds2 | 0.069 | 0.191 | 0.096 | 0.000 | -0.041 | 0.400 | 0.094 | 0.473 |
| P68373 | Tuba1c | 0.757 | 1.000 | -0.157 | 0.382 | 0.000 | 0.181 | -0.134 | -0.005 |
| Q8R550 | Sh3kbp1 | 0.065 | 0.185 | -0.456 | 0.012 | 0.000 | -0.738 | -0.809 | -0.400 |
| Q64511 | Top2b | 0.552 | 0.820 | -0.077 | 0.020 | 0.000 | -0.131 | -0.265 | 0.115 |
| Q62418 | Dbnl | 0.002 | 0.028 | -0.131 | 0.000 | 0.056 | -1.040 | -1.690 | -1.410 |
| Q9D404 | Oxsm | 0.917 | 1.000 | 0.000 | 0.611 | -0.220 | 0.234 | 0.026 | 0.046 |
| Q8BPU7 | Elmo1 | 0.003 | 0.033 | -0.217 | 0.000 | 0.110 | -0.699 | -0.889 | -0.683 |
| Q99JB2 | Stoml2 | 0.007 | 0.048 | 0.000 | 0.206 | -0.196 | 0.991 | 0.626 | 0.828 |
| P11499 | Hsp90ab1 | 0.018 | 0.084 | 0.026 | 0.000 | -0.037 | -0.346 | -0.932 | -0.757 |
| P21619 | Lmnb2 | 0.066 | 0.186 | -0.179 | 0.000 | 0.032 | -0.265 | -0.402 | -0.177 |
| Q6NZC7 | Sec23ip | 0.136 | 0.301 | 0.293 | 0.000 | -0.031 | -0.215 | -1.310 | -0.279 |
| Q922R8 | Pdia6 | 0.018 | 0.083 | 0.000 | 0.091 | -0.433 | 1.380 | 1.200 | 0.553 |
| P49710 | Hcls1 | 0.004 | 0.034 | -0.319 | 0.000 | 0.235 | -1.270 | -1.490 | -1.060 |
| Q9CY58 | Serbp1 | 0.592 | 0.864 | -0.242 | 0.000 | 0.112 | 0.215 | -0.198 | 0.139 |
| O89090 | Sp1 | 0.189 | 0.379 | 0.000 | 1.060 | -0.104 | -0.224 | -0.256 | -0.338 |
| P59017 | Bcl2l13 | 0.118 | 0.274 | 0.000 | -0.077 | 0.194 | -0.067 | -0.784 | -0.362 |
| Q9JI13 | Utp3 | 0.366 | 0.612 | 1.970 | -0.999 | 0.000 | 1.660 | 0.754 | 1.350 |
| P27659 | Rpl3 | 0.218 | 0.417 | 0.230 | -0.104 | 0.000 | 0.727 | 0.075 | 0.273 |
| P97807 | Fh | 0.048 | 0.152 | 0.000 | 0.319 | -0.305 | 0.689 | 0.384 | 0.857 |
| P11983 | Tcp1 | 0.019 | 0.087 | 0.010 | -0.033 | 0.000 | -0.369 | -0.934 | -0.590 |
| Q9QY81 | Nup210 | 0.079 | 0.209 | 0.000 | 0.154 | -0.034 | -0.133 | -0.253 | -0.062 |
| Q9EQP2 | Ehd4 | 0.227 | 0.429 | 0.000 | 0.001 | -0.845 | 0.451 | 0.115 | -0.049 |
| Q8R010 | Aimp2 | 0.173 | 0.357 | 0.013 | 0.000 | -0.531 | -0.551 | -1.030 | -0.312 |
| Q8CBY8 | Dctn4 | 0.037 | 0.129 | 0.000 | -0.092 | 0.128 | -0.321 | -0.625 | -0.243 |
| Q3TBT3 | Tmem173 | 0.649 | 0.925 | -0.395 | 0.164 | 0.000 | -0.291 | -0.323 | 0.074 |

| | | | | | | | | | |
|---|---|---|---|---|---|---|---|---|---|
| P14733 | Lmnb1 | 0.060 | 0.175 | -0.138 | 0.000 | 0.107 | -0.348 | -0.608 | -0.184 |
| P17225 | Ptbp1 | 0.985 | 1.000 | -0.053 | 0.090 | 0.000 | -0.272 | -0.102 | 0.336 |
| P61979 | Hnrnpk | 0.004 | 0.034 | -0.236 | 0.011 | 0.000 | -0.622 | -0.888 | -0.763 |
| P46935 | Nedd4 | 0.009 | 0.057 | 1.270 | 0.000 | -0.301 | 3.520 | 2.530 | 2.910 |
| Q99KV1 | Dnajb11 | 0.002 | 0.025 | 0.038 | 0.000 | -0.045 | 0.752 | 0.934 | 1.200 |
| Q9JJA4 | Wdr12 | 0.164 | 0.343 | 0.164 | 0.000 | -0.215 | -0.313 | -0.653 | -0.091 |
| Q9JHU4 | Dync1h1 | 0.241 | 0.450 | 0.000 | 0.014 | -0.111 | 0.549 | -0.014 | 0.098 |
| Q5PSV9 | Mdc1 | 0.020 | 0.088 | 0.000 | -0.100 | 0.030 | -0.538 | -0.554 | -0.234 |
| P07356 | Anxa2 | 0.002 | 0.028 | -0.294 | 0.146 | 0.000 | 0.920 | 0.824 | 1.030 |
| Q6PDM2 | Srsf1 | 0.015 | 0.076 | 0.288 | 0.000 | -0.003 | -0.389 | -0.976 | -0.902 |
| Q9Z1Q9 | Vars | 0.018 | 0.083 | 0.137 | -0.048 | 0.000 | -0.459 | -1.210 | -0.842 |
| P22646 | Cd3e | 0.501 | 0.766 | -0.349 | 0.000 | 0.371 | -0.475 | -0.277 | 0.156 |
| Q923D5 | Wbp11 | 0.022 | 0.095 | -0.054 | 0.000 | 0.115 | -0.351 | -0.733 | -0.346 |
| O08583 | Alyref | 0.509 | 0.773 | -0.288 | 0.193 | 0.000 | -0.401 | 0.188 | 1.140 |
| Q8BGH2 | Samm50 | 0.000 | 0.013 | 0.000 | 0.083 | -0.194 | 0.973 | 0.871 | 0.934 |
| Q99KK7 | Dpp3 | 0.000 | 0.013 | -0.119 | 0.102 | 0.000 | -1.230 | -1.360 | -1.600 |
| P16546 | Sptan1 | 0.055 | 0.165 | -0.057 | 0.095 | 0.000 | 0.323 | 0.097 | 0.320 |
| Q8VIJ6 | Sfpq | 0.012 | 0.066 | -0.229 | 0.123 | 0.000 | -0.574 | -0.596 | -0.442 |
| Q9CZR8 | Tsfm | 0.707 | 0.983 | 0.245 | 0.000 | -0.207 | 0.426 | -0.475 | -0.282 |
| Q9D1J3 | Sarnp | 0.287 | 0.512 | -0.551 | 0.000 | 0.172 | -0.814 | -0.430 | -0.184 |
| P23116 | Eif3a | 0.339 | 0.578 | 0.109 | 0.000 | -0.242 | 0.055 | -0.402 | -0.373 |
| Q8K4Z5 | Sf3a1 | 0.029 | 0.110 | -0.065 | 0.185 | 0.000 | -0.670 | -0.424 | -0.241 |
| Q9WVA3 | Bub3 | 0.226 | 0.429 | -0.180 | 0.000 | 0.110 | -0.149 | -0.518 | -0.085 |
| Q8BFW7 | Lpp | 0.000 | 0.009 | 0.000 | 0.546 | -0.074 | 4.720 | 4.080 | 4.430 |
| Q3TZZ7 | Esyt2 | 0.004 | 0.037 | -0.192 | 0.000 | 0.048 | 0.458 | 0.367 | 0.491 |
| P80314 | Cct2 | 0.030 | 0.113 | -0.001 | 0.000 | 0.055 | -0.220 | -0.792 | -0.640 |
| Q6PB66 | Lrpprc | 0.035 | 0.125 | 0.000 | 0.121 | -0.292 | 0.941 | 0.428 | 0.430 |
| E9PVX6 | Mki67 | 0.500 | 0.765 | 0.712 | -0.541 | 0.000 | 0.430 | -0.788 | -0.647 |
| Q7TSV4 | Pgm2 | 0.000 | 0.013 | 0.000 | -0.203 | 0.179 | -1.780 | -2.280 | -2.070 |
| Q9R0P3 | Esd | 0.086 | 0.222 | 0.000 | 0.047 | -0.224 | -0.212 | -0.757 | -0.429 |
| Q9D824 | Fip1l1 | 0.562 | 0.831 | -0.526 | 0.000 | 0.347 | -0.360 | -0.548 | 0.118 |
| Q8CI51 | Pdlim5 | 0.002 | 0.025 | 0.018 | 0.000 | -0.246 | 2.180 | 1.340 | 1.770 |
| Q99MR6 | Srrt | 0.012 | 0.069 | 0.089 | -0.063 | 0.000 | -0.600 | -1.420 | -1.250 |
| P55194 | Sh3bp1 | 0.018 | 0.083 | -0.199 | 0.000 | 0.014 | -0.550 | -1.070 | -0.614 |
| Q922P9 | Glyr1 | 0.200 | 0.394 | -0.174 | 0.000 | 0.112 | -0.349 | -0.510 | 0.002 |
| Q9D819 | Ppa1 | 0.012 | 0.069 | 0.055 | -0.145 | 0.000 | -1.050 | -1.690 | -0.821 |
| Q9QUM0 | Itga2b | 0.351 | 0.592 | 0.000 | 0.515 | -0.869 | -0.489 | -0.209 | -1.310 |
| Q80U93 | Nup214 | 0.291 | 0.518 | -0.291 | 0.118 | 0.000 | -0.193 | -0.509 | -0.103 |
| Q8BVY0 | Rsl1d1 | 0.050 | 0.155 | 0.000 | -0.234 | 0.038 | -0.355 | -1.110 | -0.679 |
| O09106 | Hdac1 | 0.745 | 1.000 | -0.195 | 0.107 | 0.000 | -0.161 | -0.227 | 0.147 |
| O09111 | Ndufb11 | 0.001 | 0.018 | 0.019 | -0.197 | 0.000 | 0.578 | 0.575 | 0.648 |
| Q9D6R2 | Idh3a | 0.546 | 0.814 | 0.000 | 0.280 | -0.166 | 0.748 | -0.097 | 0.040 |
| Q80WJ7 | Mtdh | 0.233 | 0.439 | 0.000 | -0.069 | 0.038 | 0.134 | -0.029 | 0.382 |
| Q60611 | Satb1 | 0.855 | 1.000 | 0.000 | -0.019 | 0.104 | 0.646 | -1.180 | 0.293 |
| P40124 | Cap1 | 0.001 | 0.015 | 0.000 | 0.284 | -0.079 | -1.070 | -1.130 | -1.270 |
| Q9ERU9 | Ranbp2 | 0.147 | 0.315 | 0.000 | -0.026 | 0.041 | 0.012 | -0.328 | -0.218 |
| P08113 | Hsp90b1 | 0.004 | 0.035 | 0.176 | 0.000 | -0.151 | 1.090 | 0.696 | 0.949 |
| P63024 | Vamp3 | 0.003 | 0.033 | 0.045 | 0.000 | -0.153 | 0.674 | 0.557 | 0.918 |
| P54823 | Ddx6 | 0.098 | 0.242 | 0.066 | 0.000 | -0.011 | -0.081 | -0.692 | -0.327 |
| P13864 | Dnmt1 | 0.113 | 0.267 | 0.336 | 0.000 | -0.055 | -0.098 | -0.924 | -0.353 |
| O08784 | Tcof1 | 0.113 | 0.266 | -0.009 | 0.000 | 0.150 | -0.099 | -0.536 | -0.133 |
| Q9D1G1 | Rab1b | 0.303 | 0.533 | 0.158 | -0.095 | 0.000 | -0.172 | -0.134 | 0.029 |
| Q99JY9 | Actr3 | 0.188 | 0.378 | 0.048 | -0.092 | 0.000 | 0.345 | 0.013 | 0.107 |
| P60710 | Actb | 0.156 | 0.330 | -0.073 | 0.121 | 0.000 | -0.014 | -0.514 | -0.237 |
| P63260 | Actg1 | 0.156 | 0.330 | -0.073 | 0.121 | 0.000 | -0.014 | -0.514 | -0.237 |
| Q80V86 | Ints8 | 0.570 | 0.839 | -0.504 | 3.710 | 0.000 | 0.053 | 1.310 | -0.890 |
| Q61655 | Ddx19a | 0.390 | 0.639 | 0.118 | -0.022 | 0.000 | 0.132 | -0.526 | -0.084 |
| P62830 | Rpl23 | 0.643 | 0.919 | 0.106 | -0.204 | 0.000 | 0.399 | -0.679 | -0.312 |
| Q9CQ92 | Fis1 | 0.078 | 0.209 | 0.240 | -0.067 | 0.000 | -0.226 | -0.500 | -0.127 |

| | | | | | | | | | |
|---|---|---|---|---|---|---|---|---|---|
| P54276 | Msh6 | 0.033 | 0.120 | 0.135 | 0.000 | -0.179 | -0.428 | -1.340 | -0.957 |
| P06240 | Lck | 0.294 | 0.521 | -0.202 | 0.459 | 0.000 | -0.260 | -0.257 | -0.001 |
| Q07235 | Serpine2 | 0.277 | 0.498 | 0.428 | 0.000 | -0.162 | 0.144 | -0.483 | -0.384 |
| P30681 | Hmgb2 | 0.095 | 0.238 | -0.015 | 0.000 | 0.040 | -0.144 | -0.615 | -0.195 |
| Q60953 | Pml | 0.062 | 0.178 | 0.071 | 0.000 | -0.108 | -0.221 | -1.170 | -0.818 |
| Q9ESX5 | Dkc1 | 0.025 | 0.102 | 0.012 | 0.000 | -0.193 | -0.277 | -0.447 | -0.330 |
| Q62189 | Snrpa | 0.029 | 0.111 | 0.000 | -0.283 | 0.089 | -0.543 | -0.927 | -0.493 |
| Q8VDD5 | Myh9 | 0.002 | 0.027 | 0.000 | 0.052 | -0.141 | 0.796 | 0.520 | 0.751 |
| Q922U1 | Prpf3 | 0.013 | 0.071 | -0.006 | 0.000 | 0.443 | -0.588 | -0.960 | -0.545 |
| Q78PY7 | Snd1 | 0.431 | 0.687 | 0.121 | 0.000 | -0.224 | 0.151 | -0.543 | -0.309 |
| Q80UG5 | Sept 9 | 0.237 | 0.444 | -0.272 | 0.086 | 0.000 | -0.200 | -0.429 | -0.138 |
| P24452 | Capg | 0.026 | 0.104 | -0.297 | 0.189 | 0.000 | -0.617 | -0.622 | -0.450 |
| P70372 | Elavl1 | 0.445 | 0.702 | -0.426 | 0.124 | 0.000 | -0.293 | -0.336 | -0.126 |
| Q9D0W5 | Ppil1 | 0.009 | 0.056 | -0.094 | 0.000 | 0.110 | -0.708 | -1.100 | -0.574 |
| P17182 | Eno1 | 0.002 | 0.028 | 0.055 | -0.141 | 0.000 | -0.948 | -1.530 | -1.280 |
| Q61599 | Arhgdib | 0.001 | 0.015 | 0.372 | -0.115 | 0.000 | -1.450 | -1.770 | -1.570 |
| P29758 | Oat | 0.001 | 0.018 | 0.000 | 0.170 | -0.473 | 1.810 | 1.920 | 2.200 |
| Q01320 | Top2a | 0.345 | 0.585 | 0.420 | 0.000 | -0.276 | 0.181 | -0.632 | -0.423 |
| Q9CYA0 | Creld2 | 0.202 | 0.398 | 0.461 | 0.000 | -0.153 | 0.057 | -0.768 | -0.358 |
| Q99KP6 | Prpf19 | 0.018 | 0.083 | 0.000 | -0.070 | 0.105 | -0.384 | -0.924 | -0.583 |
| Q9WV60 | Gsk3b | 0.103 | 0.249 | 0.000 | -0.045 | 0.044 | -0.511 | -1.120 | -0.157 |
| Q80VD1 | Fam98b | 0.136 | 0.301 | 0.000 | -0.052 | 0.036 | -0.205 | -0.328 | -0.014 |
| Q9Z2G6 | Sel1l | 0.015 | 0.077 | 0.000 | 0.785 | -0.235 | 1.410 | 1.440 | 1.460 |
| Q8QZY1 | Eif3l | 0.322 | 0.557 | 0.286 | 0.000 | -0.080 | 0.235 | -0.651 | -0.334 |
| Q9DBC7 | Prkar1a | 0.005 | 0.042 | -0.209 | 0.003 | 0.000 | -0.670 | -0.996 | -0.671 |
| Q9QZE5 | Copg1 | 0.155 | 0.329 | -0.425 | 0.159 | 0.000 | -0.405 | -0.615 | -0.289 |
| O08749 | Dld | 0.005 | 0.039 | 0.000 | 0.188 | -0.179 | 0.904 | 0.625 | 0.763 |
| Q9QXS1 | Plec | 0.017 | 0.081 | -0.164 | 0.000 | 0.083 | 0.797 | 0.326 | 0.702 |
| P38647 | Hspa9 | 0.072 | 0.196 | 0.035 | 0.000 | -0.356 | 1.510 | 0.407 | 0.498 |
| P63038 | Hspd1 | 0.006 | 0.044 | 0.000 | 0.119 | -0.240 | 1.090 | 0.650 | 0.822 |
| Q99KJ8 | Dctn2 | 0.536 | 0.803 | -0.313 | 0.028 | 0.000 | -0.031 | -0.389 | -0.172 |
| Q9DBL1 | Acadsb | 0.001 | 0.022 | 0.000 | 0.020 | -0.201 | 1.980 | 1.390 | 1.380 |
| Q61335 | Bcap31 | 0.080 | 0.212 | 0.000 | -0.052 | 0.226 | 1.140 | 0.266 | 0.636 |
| Q9Z2I0 | Letm1 | 0.002 | 0.025 | 0.000 | 0.057 | -0.198 | 0.565 | 0.495 | 0.591 |
| Q62318 | Trim28 | 0.170 | 0.352 | -0.106 | 0.039 | 0.000 | -0.141 | -0.502 | -0.101 |
| P84104 | Srsf3 | 0.070 | 0.193 | 0.568 | -0.014 | 0.000 | -0.104 | -0.988 | -0.770 |
| P62196 | Psmc5 | 0.040 | 0.136 | 0.000 | 0.308 | -0.008 | -0.201 | -0.818 | -0.531 |
| Q9D8E6 | Rpl4 | 0.054 | 0.163 | 0.321 | -0.021 | 0.000 | 0.782 | 0.316 | 0.637 |
| Q9CXR1 | Dhrs7 | 0.113 | 0.266 | 0.279 | 0.000 | -0.073 | 0.747 | 0.268 | 0.322 |
| Q9CWZ3 | Rbm8a | 0.096 | 0.238 | 0.000 | 0.642 | -0.100 | -0.984 | -1.000 | -0.037 |
| Q07813 | Bax | 0.323 | 0.559 | -0.110 | 0.000 | 0.085 | 0.671 | -0.114 | 0.210 |
| P68368 | Tuba4a | 0.020 | 0.087 | 0.146 | 0.000 | -0.226 | -0.480 | -1.030 | -1.100 |
| Q99JY0 | Hadhb | 0.462 | 0.721 | -0.070 | 0.350 | 0.000 | 0.625 | -0.024 | 0.237 |
| Q8BX90 | Fndc3a | 0.002 | 0.025 | 0.055 | 0.000 | -0.428 | 1.370 | 1.110 | 1.570 |
| Q3UPF5 | Zc3hav1 | 0.070 | 0.193 | -0.342 | 0.000 | 0.012 | -0.376 | -0.579 | -0.367 |
| Q91X20 | Ash2l | 0.063 | 0.180 | -0.240 | 0.077 | 0.000 | -0.381 | -0.538 | -0.236 |
| Q8BTM8 | Flna | 0.293 | 0.520 | 0.000 | 0.121 | -0.418 | 0.299 | 0.000 | 0.080 |
| P60122 | Ruvbl1 | 0.055 | 0.165 | 0.000 | -0.042 | 0.052 | -0.122 | -0.461 | -0.238 |
| Q61823 | Pdcd4 | 0.011 | 0.065 | -0.098 | 0.000 | 0.177 | -1.090 | -2.030 | -1.100 |
| P60335 | Pcbp1 | 0.101 | 0.246 | 0.188 | -0.140 | 0.000 | -0.087 | -0.939 | -0.627 |
| O88569 | Hnrnpa2b1 | 0.148 | 0.318 | -0.304 | 0.073 | 0.000 | -0.472 | -0.454 | -0.145 |
| Q9DBG6 | Rpn2 | 0.000 | 0.012 | 0.000 | 0.145 | -0.216 | 1.580 | 1.360 | 1.520 |
| P17426 | Ap2a1 | 0.687 | 0.963 | 0.021 | 0.000 | -0.170 | 0.374 | -0.291 | 0.030 |
| Q9D903 | Ebna1bp2 | 0.166 | 0.347 | -0.081 | 0.000 | 0.201 | -0.212 | -0.623 | -0.032 |
| Q9Z110 | Aldh18a1 | 0.109 | 0.261 | 0.000 | 0.052 | -0.331 | 1.350 | 0.197 | 0.449 |
| Q8VDK1 | Nit1 | 0.269 | 0.488 | 0.029 | 0.000 | -0.100 | -0.074 | -0.053 | -0.391 |
| Q9CPY7 | Lap3 | 0.806 | 1.000 | 0.158 | 0.000 | -0.383 | 0.568 | -0.516 | 0.000 |
| P14211 | Calr | 0.017 | 0.080 | 0.000 | 0.269 | -0.256 | 0.717 | 0.570 | 0.599 |
| Q91VI7 | Rnh1 | 0.157 | 0.333 | 0.147 | 0.000 | -0.013 | 0.060 | -0.933 | -0.519 |

| | | | | | | | | | |
|---|---|---|---|---|---|---|---|---|---|
| Q9JHJ0 | Tmod3 | 0.961 | 1.000 | 0.000 | 0.019 | -0.389 | 0.016 | -0.217 | -0.146 |
| P10107 | Anxa1 | 0.215 | 0.412 | 1.740 | 0.000 | -0.057 | 2.140 | 1.090 | 1.410 |
| O70145 | Ncf2 | 0.290 | 0.517 | 1.390 | 0.000 | -0.200 | 0.382 | -0.806 | -0.670 |
| P70429 | Evl | 0.271 | 0.490 | 0.032 | -0.049 | 0.000 | -0.053 | -1.080 | -0.145 |
| Q3U9G9 | Lbr | 0.634 | 0.910 | -0.053 | 0.000 | 0.182 | 0.005 | -0.311 | 0.185 |
| P50516 | Atp6v1a | 0.725 | 1.000 | 0.551 | 0.000 | -0.094 | 0.623 | -0.032 | 0.181 |
| P50247 | Ahcy | 0.001 | 0.020 | -0.059 | 0.000 | 0.042 | -1.130 | -1.720 | -1.480 |
| P40142 | Tkt | 0.039 | 0.132 | 0.533 | -0.106 | 0.000 | -0.340 | -1.110 | -0.864 |
| Q8BH59 | Slc25a12 | 0.037 | 0.127 | -0.084 | 0.153 | 0.000 | 0.368 | 0.204 | 0.480 |
| Q9R0X4 | Acot9 | 0.058 | 0.171 | 0.000 | 0.694 | -0.021 | 1.030 | 0.770 | 0.829 |
| O08914 | Faah | 0.027 | 0.106 | 0.000 | 0.057 | -0.048 | -0.181 | -0.299 | -0.124 |
| Q61550 | Rad21 | 0.309 | 0.542 | 0.000 | -0.027 | 0.146 | -0.235 | -0.456 | 0.157 |
| P52875 | Tmem165 | 0.029 | 0.110 | 0.372 | 0.000 | -0.179 | 1.140 | 0.576 | 1.390 |
| Q3TIX9 | Usp39 | 0.018 | 0.083 | 0.270 | -0.160 | 0.000 | -0.496 | -0.734 | -0.449 |
| P55302 | Lrpap1 | 0.001 | 0.015 | 0.000 | 0.001 | -0.525 | 1.780 | 1.710 | 2.060 |
| Q3U0V2 | Tradd | 0.076 | 0.204 | 0.000 | 0.234 | -0.079 | -0.090 | -0.626 | -0.805 |
| P21958 | Tap1 | 0.423 | 0.677 | -0.148 | 0.237 | 0.000 | -0.065 | -0.190 | 0.007 |
| Q3TXS7 | Psmd1 | 0.130 | 0.293 | 0.328 | -0.138 | 0.000 | -0.098 | -0.383 | -0.247 |
| Q99PV0 | Prpf8 | 0.020 | 0.087 | -0.023 | 0.225 | 0.000 | -0.196 | -0.284 | -0.261 |
| Q9Z0P5 | Twf2 | 0.004 | 0.034 | 0.062 | -0.102 | 0.000 | -1.230 | -1.830 | -1.120 |
| Q62261 | Sptbn1 | 0.014 | 0.072 | 0.000 | 0.243 | -0.084 | 0.494 | 0.512 | 0.429 |
| F6ZDS4 | Tpr | 0.023 | 0.096 | -0.076 | 0.000 | 0.080 | -0.279 | -0.547 | -0.266 |
| P11835 | Itgb2 | 0.068 | 0.188 | 0.255 | -0.053 | 0.000 | -0.095 | -0.485 | -0.320 |
| Q9CR51 | Atp6v1g1 | 0.816 | 1.000 | 0.620 | 0.000 | -0.365 | 0.327 | -0.391 | 0.055 |
| P32921 | Wars | 0.102 | 0.248 | 0.000 | -0.106 | 0.007 | -0.096 | -0.377 | -0.196 |
| Q9JIK5 | Ddx21 | 0.023 | 0.096 | 0.038 | 0.000 | -0.075 | -0.306 | -0.743 | -0.429 |
| P43404 | Zap70 | 0.038 | 0.131 | -0.465 | 0.041 | 0.000 | -0.669 | -0.946 | -0.589 |
| Q5FWK3 | Arhgap1 | 0.296 | 0.524 | 0.076 | -0.093 | 0.000 | 0.053 | -0.363 | -0.173 |
| Q8BH61 | F13a1 | 0.387 | 0.635 | 0.693 | 0.000 | -0.022 | 0.346 | -0.483 | -0.175 |
| Q8BWT1 | Acaa2 | 0.089 | 0.228 | 0.000 | 0.088 | -0.212 | 0.592 | 0.136 | 0.245 |
| P19973 | Lsp1 | 0.002 | 0.026 | 0.000 | -0.001 | 0.158 | -1.090 | -1.130 | -0.706 |
| P48678 | Lmna | 0.002 | 0.028 | 0.000 | 0.046 | -0.348 | 0.967 | 0.745 | 0.972 |
| G5E870 | Trip12 | 0.026 | 0.105 | -0.220 | 0.190 | 0.000 | -0.368 | -0.610 | -0.474 |
| Q9JHS9 | Cwc15 | 0.035 | 0.124 | 0.000 | -0.060 | 0.174 | -0.357 | -0.766 | -1.240 |
| O54824 | Il16 | 0.400 | 0.650 | -0.327 | 0.000 | 0.082 | -0.272 | -0.328 | -0.065 |
| Q91YI4 | Arrb2 | 0.043 | 0.141 | 0.421 | -0.090 | 0.000 | -0.306 | -1.050 | -0.648 |
| Q99JY3 | Gimap4 | 0.000 | 0.014 | -0.308 | 0.000 | 0.029 | -1.380 | -1.590 | -1.690 |
| P08003 | Pdia4 | 0.111 | 0.264 | 0.208 | 0.000 | -0.016 | 0.512 | 0.143 | 0.322 |
| O35295 | Purb | 0.019 | 0.085 | -0.254 | 0.142 | 0.000 | -0.477 | -0.948 | -0.748 |
| P62821 | Rab1A | 0.807 | 1.000 | 0.232 | -0.032 | 0.000 | 0.737 | -0.085 | -0.211 |
| Q08943 | Ssrp1 | 0.047 | 0.149 | 0.000 | 0.059 | -0.129 | -0.264 | -0.982 | -0.659 |
| Q9CW46 | Raver1 | 0.212 | 0.409 | -0.429 | 0.000 | 0.009 | -0.636 | -0.311 | -0.288 |
| Q8BL97 | Srsf7 | 0.044 | 0.142 | 0.415 | 0.000 | -0.051 | -0.237 | -0.986 | -0.723 |
| Q8C7E9 | Cstf2t | 0.231 | 0.435 | 0.000 | -0.128 | 0.732 | -4.370 | -0.082 | -0.715 |
| P49722 | Psma2 | 0.000 | 0.014 | -0.245 | 0.000 | 0.011 | -1.230 | -1.480 | -1.240 |
| Q9D8W5 | Psmd12 | 0.045 | 0.146 | 0.000 | 0.153 | -0.090 | -0.282 | -1.060 | -0.637 |
| Q9WUB3 | Pygm | 0.033 | 0.121 | -0.282 | 0.000 | 0.033 | -1.330 | -0.756 | -0.562 |
| Q99N84 | Mrps18b | 0.001 | 0.019 | -0.100 | 0.262 | 0.000 | 1.290 | 1.020 | 1.220 |
| Q99LX0 | Park7 | 0.001 | 0.015 | 0.016 | -0.257 | 0.000 | -1.140 | -1.450 | -1.350 |
| Q3UHJ0 | Aak1 | 0.314 | 0.547 | -0.257 | 0.000 | 0.048 | -0.279 | -0.575 | -0.009 |
| Q9DAU1 | Cnpy3 | 0.036 | 0.125 | 0.021 | 0.000 | -0.450 | 0.436 | 0.377 | 0.266 |
| Q60710 | Samhd1 | 0.000 | 0.013 | -0.120 | 0.000 | 0.000 | -0.783 | -1.010 | -0.907 |
| Q91ZW3 | Smarca5 | 0.092 | 0.233 | 0.000 | 0.038 | -0.092 | -0.173 | -0.514 | -0.168 |
| Q9CQF9 | Pcyox1 | 0.003 | 0.031 | 0.261 | 0.000 | -0.610 | 1.640 | 1.550 | 1.550 |
| Q3UN02 | Lclat1 | 0.023 | 0.096 | -0.171 | 0.817 | 0.000 | 1.370 | 1.200 | 1.660 |
| Q80YR5 | Safb2 | 0.033 | 0.119 | -0.279 | 0.036 | 0.000 | -0.582 | -0.899 | -0.420 |
| P26039 | Tln1 | 0.041 | 0.137 | 0.059 | 0.000 | -0.282 | -0.358 | -1.020 | -1.000 |
| Q76MZ3 | Ppp2r1a | 0.013 | 0.069 | 0.000 | 0.029 | -0.018 | -0.282 | -0.657 | -0.641 |
| P97371 | Psme1 | 0.001 | 0.022 | -0.041 | 0.476 | 0.000 | -1.210 | -1.170 | -1.120 |

| | | | | | | | | | |
|---|---|---|---|---|---|---|---|---|---|
| Q9QUJ7 | Acsl4 | 0.326 | 0.563 | 0.186 | 0.000 | -0.010 | 0.509 | -0.057 | 0.321 |
| Q9CQ06 | Mrpl24 | 0.031 | 0.115 | 0.158 | 0.000 | -0.175 | 0.302 | 0.820 | 0.865 |
| Q80X90 | Flnb | 0.001 | 0.014 | 0.069 | 0.000 | -0.289 | 2.350 | 1.740 | 2.050 |
| Q9R0E1 | Plod3 | 0.000 | 0.009 | 0.000 | 0.480 | -0.111 | 3.740 | 3.370 | 3.760 |
| Q9WU78 | Pdcd6ip | 0.102 | 0.247 | -0.082 | 0.153 | 0.000 | -0.146 | -0.697 | -0.246 |
| Q8BIQ5 | Cstf2 | 0.421 | 0.675 | -0.437 | 0.000 | 0.414 | -1.200 | -0.354 | 0.232 |
| P67984 | Rpl22 | 0.254 | 0.467 | 0.247 | 0.000 | -0.018 | 0.449 | 0.053 | 0.300 |
| P47740 | Aldh3a2 | 0.057 | 0.169 | 0.000 | 0.097 | -0.121 | 0.310 | 0.102 | 0.328 |
| Q9CPV4 | Glod4 | 0.023 | 0.096 | 0.000 | 0.150 | -0.049 | -0.326 | -0.824 | -0.470 |
| P26040 | Ezr | 0.241 | 0.451 | 0.000 | 0.009 | -0.085 | 0.234 | -0.093 | 0.286 |
| P42227 | Stat3 | 0.103 | 0.250 | 0.000 | -0.155 | 0.131 | -0.089 | -0.977 | -0.688 |
| Q9Z1N5 | Ddx39b | 0.760 | 1.000 | -0.016 | 0.782 | 0.000 | 0.109 | 0.138 | 0.256 |
| P01831 | Thy1 | 0.024 | 0.098 | 0.000 | -0.028 | 0.249 | 0.606 | 0.378 | 0.771 |
| Q8K0C4 | Cyp51a1 | 0.000 | 0.008 | 0.000 | 0.282 | -0.189 | 3.810 | 3.390 | 3.830 |
| Q9D0L7 | Armc10 | 0.027 | 0.107 | -0.131 | 0.052 | 0.000 | -0.301 | -0.431 | -0.208 |
| Q8BMP6 | Acbd3 | 0.380 | 0.629 | 0.334 | -0.249 | 0.000 | 0.447 | -1.140 | -0.709 |
| O35892 | Sp100 | 0.119 | 0.276 | 0.000 | -0.023 | 0.093 | -0.048 | -0.538 | -0.220 |
| P36371 | Tap2 | 0.787 | 1.000 | -0.225 | 0.273 | 0.000 | -0.153 | -0.078 | 0.133 |
| Q9CZ44 | Nsfl1c | 0.000 | 0.011 | -0.089 | 0.000 | 0.161 | -1.340 | -1.640 | -1.510 |
| Q9CR61 | Ndufb7 | 0.020 | 0.088 | 0.000 | -0.043 | 0.113 | 0.717 | 0.435 | 1.080 |
| P23492 | Pnp | 0.010 | 0.060 | 0.405 | 0.000 | -0.271 | -0.763 | -1.050 | -1.040 |
| O88685 | Psmc3 | 0.008 | 0.054 | 0.000 | 0.126 | -0.124 | -0.422 | -0.569 | -0.370 |
| P62960 | Ybx1 | 0.498 | 0.764 | 0.017 | 0.000 | -0.011 | 0.279 | -0.611 | -0.238 |
| Q8C0I1 | Agps | 0.043 | 0.142 | 0.639 | 0.000 | -0.154 | 1.310 | 0.764 | 0.956 |
| Q8R326 | Pspc1 | 0.029 | 0.110 | 0.000 | 0.345 | -0.071 | -0.607 | -0.409 | -0.294 |
| Q9DC69 | Ndufa9 | 0.004 | 0.037 | -0.072 | 0.102 | 0.000 | 0.741 | 0.442 | 0.635 |
| Q80YV3 | Trrap | 0.070 | 0.193 | -0.087 | 0.000 | 0.024 | -0.210 | -0.361 | -0.101 |
| P47941 | Crkl | 0.001 | 0.017 | 0.039 | -0.088 | 0.000 | -1.070 | -1.520 | -1.270 |
| P11276 | Fn1 | 0.000 | 0.013 | 0.759 | 0.000 | -0.029 | 4.210 | 3.600 | 3.790 |
| Q9D1M7 | Fkbp11 | 0.001 | 0.014 | 0.000 | 0.157 | -0.617 | 2.940 | 2.470 | 2.970 |
| P56399 | Usp5 | 0.041 | 0.136 | -0.064 | 0.055 | 0.000 | -0.344 | -1.040 | -1.390 |
| Q9CQN1 | Trap1 | 0.459 | 0.719 | 0.170 | 0.000 | -0.139 | 0.175 | -0.335 | -0.254 |
| O89023 | Tpp1 | 0.463 | 0.722 | 0.691 | 0.000 | -0.365 | 0.480 | -0.634 | -0.724 |
| Q8BHD7 | Ptbp3 | 0.069 | 0.192 | 0.000 | 0.316 | -0.167 | -0.178 | -0.748 | -0.545 |
| P11881 | Itpr1 | 0.010 | 0.059 | 0.000 | 0.236 | -0.244 | 0.838 | 0.632 | 0.646 |
| Q99KI3 | Emc3 | 0.001 | 0.015 | 0.000 | 0.228 | -0.156 | 1.460 | 1.280 | 1.590 |
| Q9WTI7 | Myo1c | 0.000 | 0.012 | 0.000 | 0.063 | -0.194 | 2.750 | 2.170 | 2.720 |
| Q61210 | Arhgef1 | 0.005 | 0.041 | 0.000 | 0.001 | -0.005 | -0.416 | -0.797 | -0.644 |
| P48036 | Anxa5 | 0.005 | 0.042 | 0.000 | 0.177 | -0.377 | 0.835 | 0.814 | 0.988 |
| B2RY56 | Rbm25 | 0.013 | 0.071 | 0.000 | -0.016 | 0.053 | -0.439 | -1.080 | -0.945 |
| P15702 | Spn | 0.027 | 0.105 | -0.280 | 0.000 | 0.175 | -0.428 | -0.534 | -0.621 |
| P12815 | Pdcd6 | 0.518 | 0.782 | -0.178 | 0.236 | 0.000 | -0.267 | 0.110 | -0.131 |
| Q9D6Z1 | Nop56 | 0.058 | 0.171 | -0.091 | 0.006 | 0.000 | -0.160 | -0.587 | -0.343 |
| Q9EPB4 | Pycard | 0.002 | 0.023 | 0.029 | 0.000 | -0.084 | -1.520 | -1.930 | -1.240 |
| D0QMC3 | Mndal | 0.015 | 0.076 | -0.179 | 0.041 | 0.000 | -0.432 | -0.756 | -0.459 |
| P80313 | Cct7 | 0.026 | 0.104 | 0.099 | 0.000 | -0.035 | -0.245 | -0.699 | -0.421 |
| Q9Z2Z6 | Slc25a20 | 0.006 | 0.043 | 0.161 | 0.000 | -0.081 | 0.953 | 0.633 | 0.619 |
| Q8VHE0 | Sec63 | 0.001 | 0.014 | 0.000 | 0.208 | -0.401 | 2.140 | 1.810 | 1.980 |
| Q64674 | Srm | 0.120 | 0.277 | 0.297 | 0.000 | -0.271 | -0.147 | -0.522 | -0.530 |
| P99027 | Rplp2 | 0.515 | 0.779 | 0.066 | -0.014 | 0.000 | -0.010 | -0.366 | 0.116 |
| Q921H8 | Acaa1a | 0.168 | 0.349 | 0.077 | 0.000 | -0.377 | 0.732 | -0.098 | 0.503 |
| P21550 | Eno3 | 0.001 | 0.022 | 0.000 | 0.285 | -0.011 | -0.823 | -1.150 | -1.090 |
| Q08024 | Cbfb | 0.018 | 0.083 | -0.165 | 0.091 | 0.000 | -0.345 | -0.749 | -0.625 |
| Q6P2L6 | Whsc1l1 | 0.352 | 0.593 | -0.300 | 0.000 | 0.252 | -0.124 | -1.230 | -0.019 |
| Q62523 | Zyx | 0.182 | 0.369 | -0.086 | 0.110 | 0.000 | 0.768 | -0.019 | 0.410 |
| Q9DB05 | Napa | 0.787 | 1.000 | 0.217 | 0.000 | -0.130 | 0.223 | -0.284 | -0.005 |
| Q99K48 | Nono | 0.023 | 0.096 | -0.200 | 0.000 | 0.002 | -0.434 | -0.635 | -0.333 |
| O09044 | Snap23 | 0.002 | 0.023 | 0.028 | 0.000 | -0.068 | 0.650 | 0.420 | 0.604 |
| Q61233 | Lcp1 | 0.000 | 0.008 | 0.000 | 0.104 | -0.046 | -1.290 | -1.480 | -1.380 |

| | | | | | | | | | |
|---|---|---|---|---|---|---|---|---|---|
| P24527 | Lta4h | 0.001 | 0.019 | 0.296 | 0.000 | -0.168 | -1.520 | -2.110 | -1.850 |
| P52480 | Pkm | 0.003 | 0.031 | 0.033 | 0.000 | -0.045 | -0.658 | -1.130 | -1.020 |
| Q60520 | Sin3a | 0.049 | 0.154 | -0.146 | 0.000 | 0.086 | -0.240 | -0.604 | -0.309 |
| Q68FL6 | Mars | 0.598 | 0.871 | 0.310 | 0.000 | -0.218 | 0.343 | -0.524 | -0.239 |
| Q8BV49 | Pyhin1 | 0.026 | 0.105 | 0.000 | -0.130 | 0.140 | -0.427 | -0.852 | -0.407 |
| Q01730 | Rsu1 | 0.373 | 0.622 | 0.000 | 0.204 | -0.850 | 0.353 | 0.275 | -0.182 |
| O08553 | Dpysl2 | 0.008 | 0.055 | -0.381 | 0.023 | 0.000 | -0.799 | -1.040 | -0.780 |
| Q3U7R1 | Esyt1 | 0.377 | 0.625 | -0.054 | 0.266 | 0.000 | 0.169 | 0.087 | 0.307 |
| Q9JJX6 | P2rx4 | 0.610 | 0.885 | 0.645 | -0.068 | 0.000 | 0.517 | -0.345 | -0.170 |
| Q91W90 | Txndc5 | 0.003 | 0.033 | 0.000 | 0.085 | -0.303 | 0.869 | 0.642 | 0.871 |
| P28740 | Kif2a | 0.128 | 0.291 | 0.126 | 0.000 | -0.368 | -0.159 | -0.728 | -0.764 |
| Q9WVA4 | Tagln2 | 0.004 | 0.038 | -0.036 | 0.121 | 0.000 | -0.400 | -0.593 | -0.378 |
| Q99P72 | Rtn4 | 0.000 | 0.006 | 0.000 | 0.066 | -0.069 | 1.910 | 1.670 | 1.840 |
| P09405 | Ncl | 0.005 | 0.042 | 0.000 | 0.013 | -0.184 | -0.731 | -1.300 | -1.140 |
| Q9ERS2 | Ndufa13 | 0.056 | 0.168 | 0.038 | -0.179 | 0.000 | 0.652 | 0.143 | 0.876 |
| P29391 | Ftl1 | 0.048 | 0.152 | 0.334 | -0.304 | 0.000 | -0.448 | -1.460 | -0.987 |
| P70288 | Hdac2 | 0.731 | 1.000 | 0.000 | 0.251 | -0.259 | -0.156 | -0.163 | 0.118 |
| Q8VE70 | Pdcd10 | 0.005 | 0.039 | 0.000 | -0.139 | 0.036 | -0.455 | -0.496 | -0.690 |
| P19783 | Cox4i1 | 0.004 | 0.038 | 0.100 | -0.024 | 0.000 | 0.711 | 0.422 | 0.696 |
| Q9CZ42 | Naxd | 0.001 | 0.018 | 0.148 | 0.000 | -0.050 | 1.030 | 0.765 | 0.960 |
| P61967 | Ap1s1 | 0.008 | 0.054 | 0.000 | -0.021 | 0.223 | -0.424 | -0.864 | -0.773 |
| P42225 | Stat1 | 0.005 | 0.039 | -0.079 | 0.271 | 0.000 | -0.603 | -0.992 | -0.866 |
| Q9CVB6 | Arpc2 | 0.043 | 0.142 | 0.217 | -0.103 | 0.000 | 0.408 | 0.250 | 0.503 |
| Q99LC3 | Ndufa10 | 0.006 | 0.047 | 0.000 | 0.010 | -0.249 | 0.718 | 0.460 | 0.860 |
| Q6PFD9 | Nup98 | 0.115 | 0.270 | -0.095 | 0.000 | 0.190 | -0.155 | -0.472 | -0.120 |
| Q8R2Y8 | Ptrh2 | 0.015 | 0.077 | -0.265 | 0.190 | 0.000 | 1.240 | 1.150 | 2.290 |
| P54728 | Rad23b | 0.023 | 0.096 | -0.121 | 0.083 | 0.000 | -0.526 | -1.300 | -0.766 |
| Q9WTM5 | Ruvbl2 | 0.218 | 0.417 | 0.000 | 0.106 | -0.153 | -0.146 | -0.350 | -0.056 |
| Q9R1P4 | Psma1 | 0.012 | 0.067 | -0.187 | 0.086 | 0.000 | -0.997 | -2.060 | -1.320 |
| Q8R4N0 | Clybl | 0.061 | 0.177 | 0.182 | 0.000 | -0.138 | 0.704 | 0.176 | 0.694 |
| Q91VC3 | Eif4a3 | 0.006 | 0.046 | 0.000 | -0.016 | 0.085 | -0.409 | -0.644 | -0.359 |
| O88696 | Clpp | 0.016 | 0.078 | 0.000 | 0.043 | -0.364 | 0.863 | 0.401 | 0.665 |
| P70699 | Gaa | 0.001 | 0.022 | 0.000 | -0.360 | 0.035 | 1.170 | 0.910 | 1.070 |
| Q99NB9 | Sf3b1 | 0.056 | 0.166 | -0.164 | 0.007 | 0.000 | -0.508 | -0.725 | -0.208 |
| Q8BHL5 | Elmo2 | 0.073 | 0.198 | -0.116 | 0.032 | 0.000 | -0.361 | -0.697 | -1.500 |
| Q9D1Q6 | Erp44 | 0.003 | 0.033 | 0.018 | 0.000 | -0.372 | 0.878 | 0.686 | 0.710 |
| D3Z6Q9 | Bin2 | 0.005 | 0.039 | -0.352 | 0.269 | 0.000 | -1.050 | -1.470 | -1.300 |
| Q80Y81 | Elac2 | 0.624 | 0.899 | 0.000 | -0.208 | 0.155 | 0.131 | -0.831 | 0.116 |
| P24668 | M6pr | 0.297 | 0.524 | 0.000 | -1.460 | 0.258 | 0.120 | 0.295 | 0.320 |
| Q8BH43 | Wasf2 | 0.373 | 0.621 | -0.073 | 0.000 | 0.024 | 0.173 | -0.125 | 0.272 |
| P61358 | Rpl27 | 0.051 | 0.157 | 0.000 | 0.389 | -0.168 | 0.470 | 0.530 | 0.698 |
| Q8BP47 | Nars | 0.300 | 0.529 | 0.063 | 0.000 | -0.627 | 0.134 | -0.049 | 0.174 |
| P42208 | Sept 2 | 0.001 | 0.022 | 0.181 | 0.000 | -0.114 | 0.895 | 0.778 | 1.010 |
| Q9CRD0 | Ociad1 | 0.184 | 0.372 | -0.379 | 0.247 | 0.000 | -0.636 | -0.622 | -0.097 |
| Q6PDG5 | Smarcc2 | 0.084 | 0.220 | 0.000 | -0.027 | 0.113 | -0.028 | -0.406 | -0.317 |
| Q64105 | Spr | 0.003 | 0.033 | -0.370 | 0.096 | 0.000 | -1.650 | -1.710 | -1.160 |
| P25206 | Mcm3 | 0.066 | 0.185 | 0.280 | 0.000 | -0.196 | -0.205 | -1.110 | -0.943 |
| P08752 | Gnai2 | 0.756 | 1.000 | 0.074 | 0.000 | -0.107 | 0.169 | -0.194 | -0.132 |
| O35382 | Exoc4 | 0.173 | 0.357 | 0.171 | 0.000 | -0.131 | 0.080 | -0.688 | -0.729 |
| O08788 | Dctn1 | 0.369 | 0.616 | 0.018 | -0.153 | 0.000 | 0.258 | -0.787 | -0.596 |
| P24161 | Cd247 | 0.811 | 1.000 | -0.617 | 0.000 | 0.247 | -0.627 | 0.147 | 0.420 |
| Q9CW03 | Smc3 | 0.037 | 0.128 | 0.000 | -0.055 | 0.001 | -0.243 | -0.484 | -0.187 |
| Q8CIE6 | Copa | 0.118 | 0.274 | 0.168 | 0.000 | -0.012 | -0.029 | -0.703 | -0.326 |
| Q9ESP1 | Sdf2l1 | 0.816 | 1.000 | 0.000 | 0.023 | -0.647 | -0.293 | -0.576 | 0.035 |
| P09528 | Fth1 | 0.085 | 0.220 | 0.442 | -0.233 | 0.000 | -0.225 | -1.190 | -0.715 |
| Q9DB77 | Uqcrc2 | 0.140 | 0.306 | 0.000 | 0.125 | -0.097 | 0.556 | 0.046 | 0.312 |
| O08810 | Eftud2 | 0.022 | 0.095 | 0.000 | 0.151 | -0.093 | -0.237 | -0.287 | -0.220 |
| Q8C4Q6 | Aida | 0.026 | 0.105 | -0.212 | 0.550 | 0.000 | 1.430 | 1.020 | 0.842 |
| Q05186 | Rcn1 | 0.000 | 0.003 | -0.133 | 0.156 | 0.000 | 3.830 | 3.540 | 3.760 |

| | | | | | | | | | |
|---|---|---|---|---|---|---|---|---|---|
| Q8K183 | Pdxk | 0.017 | 0.081 | 0.000 | -0.383 | 0.018 | -0.842 | -1.590 | -1.010 |
| P17710 | Hk1 | 0.085 | 0.221 | -0.211 | 0.114 | 0.000 | -0.278 | -0.528 | -0.214 |
| Q61425 | Hadh | 0.375 | 0.623 | -0.733 | 0.214 | 0.000 | 0.578 | 0.284 | -0.255 |
| Q8C570 | Rae1 | 0.961 | 1.000 | -0.124 | 0.114 | 0.000 | -0.099 | 0.017 | 0.085 |
| O54941 | Smarce1 | 0.092 | 0.232 | -0.025 | 0.000 | 0.203 | -0.414 | -0.671 | -0.041 |
| Q9Z1Q2 | Abhd16a | 0.066 | 0.186 | 0.213 | -0.145 | 0.000 | 0.518 | 0.389 | 1.190 |
| P10518 | Alad | 0.007 | 0.050 | 0.086 | -0.152 | 0.000 | -1.240 | -2.500 | -1.940 |
| Q9Z1Z2 | Strap | 0.011 | 0.064 | 0.000 | -0.050 | 0.159 | -0.265 | -0.534 | -0.473 |
| Q99MN9 | Pccb | 0.003 | 0.032 | 0.000 | 0.238 | -0.023 | 0.894 | 0.712 | 1.040 |
| Q9DBG3 | Ap2b1 | 0.155 | 0.329 | 0.000 | 0.158 | -0.099 | 0.695 | 0.126 | 0.236 |
| Q8VDN2 | Atp1a1 | 0.012 | 0.069 | -0.147 | 0.017 | 0.000 | 0.628 | 0.290 | 0.682 |
| P14901 | Hmox1 | 0.394 | 0.643 | 0.207 | 0.000 | -0.907 | -0.150 | -0.921 | -0.834 |
| O35083 | Agpat1 | 0.035 | 0.123 | 0.411 | 0.000 | -0.054 | 1.020 | 0.828 | 0.500 |
| Q3U1F9 | Pag1 | 0.317 | 0.550 | 0.105 | -0.106 | 0.000 | -0.473 | -0.720 | 0.213 |
| Q8BZN6 | Dock10 | 0.210 | 0.406 | -0.532 | 0.000 | 0.228 | -0.481 | -1.600 | -0.303 |
| P24063 | Itgal | 0.054 | 0.163 | -0.246 | 0.010 | 0.000 | -0.641 | -0.514 | -0.247 |
| Q8BMS1 | Hadha | 0.932 | 1.000 | 0.027 | 0.000 | -0.269 | 0.492 | -0.475 | -0.178 |
| Q9R1T4 | Sept 6 | 0.016 | 0.078 | -0.316 | 0.001 | 0.000 | -0.583 | -0.779 | -0.525 |
| P38060 | Hmgcl | 0.061 | 0.177 | 0.000 | 0.148 | -0.178 | 0.592 | 0.202 | 0.328 |
| Q61191 | Hcfc1 | 0.920 | 1.000 | -0.021 | 0.000 | 0.544 | 0.085 | -0.064 | 0.590 |
| P06151 | Ldha | 0.001 | 0.018 | -0.165 | 0.115 | 0.000 | -0.980 | -1.190 | -0.934 |
| Q9CZ13 | Uqcrc1 | 0.015 | 0.076 | 0.081 | 0.000 | -0.051 | 0.660 | 0.292 | 0.464 |
| Q3TDQ1 | Stt3b | 0.003 | 0.031 | 0.054 | 0.000 | -0.451 | 1.190 | 0.890 | 1.070 |
| Q922W5 | Pycr1 | 0.000 | 0.014 | 0.208 | 0.000 | -0.219 | 2.510 | 1.920 | 2.420 |
| Q91W39 | Ncoa5 | 0.060 | 0.175 | 0.026 | 0.000 | -0.041 | -0.220 | -0.959 | -0.518 |
| P63168 | Dynll1 | 0.922 | 1.000 | 0.502 | -0.203 | 0.000 | 0.013 | -0.054 | 0.267 |
| Q9EQH3 | Vps35 | 0.223 | 0.424 | 0.352 | -0.082 | 0.000 | 0.112 | -0.639 | -0.304 |
| P97287 | Mcl1 | 0.225 | 0.427 | 0.197 | 0.000 | -0.777 | 0.487 | -0.153 | 1.300 |
| P50544 | Acadvl | 0.245 | 0.456 | 0.000 | 0.061 | -0.285 | 0.866 | -0.022 | 0.132 |
| Q9DC51 | Gnai3 | 0.950 | 1.000 | 0.028 | -0.131 | 0.000 | 0.178 | -0.388 | 0.145 |
| Q8BK64 | Ahsa1 | 0.072 | 0.197 | 0.000 | 0.218 | -0.021 | -0.131 | -1.130 | -0.729 |
| Q9WTX5 | Skp1 | 0.009 | 0.057 | 0.000 | 0.142 | -0.147 | -0.508 | -0.496 | -0.360 |
| P42209 | Sept 1 | 0.040 | 0.135 | -0.383 | 0.163 | 0.000 | -0.531 | -0.641 | -0.536 |
| P32233 | Drg1 | 0.735 | 1.000 | 0.167 | -0.118 | 0.000 | 0.094 | -0.600 | 0.256 |
| P26043 | Rdx | 0.042 | 0.138 | 0.000 | 0.671 | -0.017 | 0.763 | 1.170 | 1.520 |
| P63158 | Hmgb1 | 0.474 | 0.734 | 0.000 | -0.236 | 0.219 | 0.196 | -0.585 | -0.247 |
| Q9R1C7 | Prpf40a | 0.057 | 0.169 | 0.000 | 0.036 | -0.175 | -0.196 | -0.679 | -0.769 |
| P62192 | Psmc1 | 0.016 | 0.078 | 0.000 | 0.007 | -0.059 | -0.317 | -0.671 | -0.390 |
| Q61656 | Ddx5 | 0.021 | 0.090 | 0.145 | 0.000 | -0.259 | -0.449 | -0.909 | -0.754 |
| Q8CAQ8 | Immt | 0.003 | 0.032 | 0.000 | 0.109 | -0.145 | 0.915 | 0.583 | 0.795 |
| P56480 | Atp5b | 0.009 | 0.057 | 0.000 | 0.258 | -0.162 | 0.723 | 0.568 | 0.808 |
| P60867 | Rps20 | 0.015 | 0.077 | 0.165 | 0.000 | -0.084 | -0.470 | -0.687 | -0.324 |
| Q9QXZ0 | Macf1 | 0.523 | 0.789 | 0.215 | -0.056 | 0.000 | 0.505 | -0.100 | 0.160 |
| P97351 | Rps3a | 0.907 | 1.000 | 0.091 | -0.290 | 0.000 | 0.303 | -0.279 | -0.309 |
| Q8CEC0 | Nup88 | 0.199 | 0.394 | -0.116 | 0.129 | 0.000 | -0.168 | -0.370 | -0.019 |
| Q8R5J9 | Arl6ip5 | 0.031 | 0.115 | -0.357 | 0.100 | 0.000 | 0.322 | 0.545 | 0.805 |
| Q64514 | Tpp2 | 0.416 | 0.669 | 0.008 | 0.000 | -1.190 | -0.674 | -0.816 | -0.777 |
| Q6GQT9 | Nomo1 | 0.001 | 0.018 | 0.066 | 0.000 | -0.516 | 2.070 | 1.620 | 1.860 |
| Q9CYG7 | Tomm34 | 0.013 | 0.071 | -0.013 | 0.210 | 0.000 | -0.338 | -0.800 | -0.588 |
| Q9Z329 | Itpr2 | 0.713 | 0.989 | -0.251 | 0.000 | 0.041 | -0.190 | -0.184 | 0.027 |
| Q8CCJ3 | Ufl1 | 0.399 | 0.650 | 0.000 | -0.401 | 0.074 | -0.084 | -0.609 | -0.240 |
| P46460 | Nsf | 0.134 | 0.298 | 0.377 | 0.000 | -0.134 | 0.037 | -0.914 | -0.723 |
| P61164 | Actr1a | 0.105 | 0.252 | 0.017 | -0.031 | 0.000 | -0.230 | -0.245 | -0.015 |
| O35857 | Timm44 | 0.016 | 0.078 | 0.231 | 0.000 | -0.162 | 0.827 | 0.450 | 0.765 |
| Q3THE2 | Myl12b | 0.009 | 0.056 | -0.141 | 0.510 | 0.000 | 0.987 | 1.260 | 1.210 |
| Q60972 | Rbbp4 | 0.188 | 0.378 | -0.072 | 0.000 | 0.231 | -0.232 | -0.800 | -0.009 |
| P09411 | Pgk1 | 0.021 | 0.091 | 0.132 | -0.121 | 0.000 | -0.542 | -1.100 | -0.544 |
| Q9WUQ2 | Preb | 0.007 | 0.048 | -0.081 | 0.122 | 0.000 | 0.725 | 0.406 | 0.640 |
| Q8CC88 | Vwa8 | 0.942 | 1.000 | 0.000 | 0.045 | -0.313 | 0.847 | -0.752 | -0.481 |

| | | | | | | | | | |
|---|---|---|---|---|---|---|---|---|---|
| Q9D7N9 | Apmap | 0.049 | 0.154 | 0.018 | 0.000 | -0.061 | 0.406 | 0.078 | 0.386 |
| O54782 | Man2b2 | 0.003 | 0.033 | 0.000 | 0.230 | -0.073 | -1.030 | -1.770 | -1.620 |
| P80316 | Cct5 | 0.042 | 0.140 | 0.066 | -0.104 | 0.000 | -0.220 | -0.838 | -0.649 |
| Q8BMJ2 | Lars | 0.881 | 1.000 | 0.252 | 0.000 | -0.100 | 0.278 | -0.242 | 0.028 |
| Q9DBZ1 | Ikbip | 0.000 | 0.002 | 0.000 | -0.098 | 0.101 | 6.270 | 5.940 | 6.260 |
| P43406 | Itgav | 0.000 | 0.013 | 0.172 | -0.250 | 0.000 | 2.300 | 1.820 | 2.020 |
| P10711 | Tcea1 | 0.098 | 0.242 | 0.028 | -0.179 | 0.000 | -0.348 | -0.662 | -0.165 |
| Q9D2V8 | Mfsd10 | 0.007 | 0.051 | 0.000 | 0.022 | -0.232 | 0.826 | 0.424 | 0.713 |
| Q91VD9 | Ndufs1 | 0.002 | 0.025 | 0.000 | 0.197 | -0.065 | 0.847 | 0.648 | 0.786 |
| Q9D2V7 | Coro7 | 0.000 | 0.008 | 0.000 | -0.014 | 0.005 | -1.400 | -1.690 | -1.670 |
| Q8BFR5 | Tufm | 0.440 | 0.697 | 0.165 | 0.000 | -0.339 | 0.256 | -0.698 | -0.598 |
| P47911 | Rpl6 | 0.036 | 0.126 | 0.095 | 0.000 | -0.087 | 0.610 | 0.185 | 0.488 |
| Q14C51 | Ptcd3 | 0.001 | 0.019 | 0.000 | 0.011 | -0.187 | 0.717 | 0.554 | 0.741 |
| P62814 | Atp6v1b2 | 0.820 | 1.000 | 0.689 | 0.000 | -0.027 | 0.606 | -0.188 | -0.001 |
| Q9WV55 | Vapa | 0.035 | 0.124 | 0.000 | 0.399 | -0.125 | 0.518 | 0.621 | 0.719 |
| P49718 | Mcm5 | 0.432 | 0.688 | 0.140 | 0.000 | -0.032 | 0.135 | -0.802 | -0.001 |
| P97760 | Polr2c | 0.231 | 0.436 | -0.024 | 0.099 | 0.000 | -0.330 | -1.380 | 0.008 |
| P10605 | Ctsb | 0.113 | 0.266 | 0.436 | 0.000 | -0.385 | -0.190 | -1.040 | -0.842 |
| Q9CU62 | Smc1a | 0.086 | 0.222 | 0.000 | -0.003 | 0.011 | -0.155 | -0.506 | -0.143 |
| P97855 | G3bp1 | 0.321 | 0.557 | 0.068 | -0.159 | 0.000 | -0.136 | -1.010 | -0.025 |
| Q6PDQ2 | Chd4 | 0.367 | 0.615 | 0.000 | -0.102 | 0.117 | -0.201 | -0.551 | 0.134 |
| Q3TBD2 | Hmha1 | 0.000 | 0.014 | 0.000 | -0.131 | 0.051 | -1.160 | -1.530 | -1.440 |
| P26450 | Pik3r1 | 0.468 | 0.727 | 0.000 | 0.958 | -0.100 | -0.021 | 0.030 | 0.038 |
| Q62087 | Pon3 | 0.014 | 0.073 | -0.127 | 0.000 | 0.034 | 0.275 | 0.248 | 0.476 |
| Q7TMK9 | Syncrip | 0.677 | 0.953 | 0.000 | 0.301 | -0.343 | 0.126 | -0.176 | 0.326 |
| Q9CZX9 | Emc4 | 0.015 | 0.076 | -0.448 | 0.367 | 0.000 | 0.940 | 1.180 | 0.890 |
| Q91VK1 | Bzw2 | 0.115 | 0.269 | 0.350 | 0.000 | -0.019 | -0.027 | -0.652 | -0.298 |
| Q3TKT4 | Smarca4 | 0.343 | 0.583 | 0.057 | -0.076 | 0.000 | 0.065 | -0.478 | -0.132 |
| Q9DB85 | Rrp8 | 0.469 | 0.728 | -0.371 | 0.000 | 0.165 | -0.132 | -0.454 | -0.090 |
| P17427 | Ap2a2 | 0.383 | 0.632 | 0.415 | 0.000 | -0.703 | 0.757 | -0.016 | 0.153 |
| Q99L47 | St13 | 0.014 | 0.072 | -0.081 | 0.036 | 0.000 | -0.526 | -1.130 | -0.692 |
| Q99P58 | Rab27b | 0.322 | 0.557 | 0.000 | 0.955 | -1.050 | -0.404 | -0.436 | -1.680 |
| P50580 | Pa2g4 | 0.244 | 0.454 | 0.328 | -0.040 | 0.000 | 0.166 | -0.734 | -0.310 |
| Q8R0G9 | Nup133 | 0.563 | 0.832 | 0.000 | -0.023 | 0.096 | 0.037 | -0.449 | 0.137 |
| Q9QX60 | Dguok | 0.708 | 0.984 | -0.077 | 0.150 | 0.000 | 0.045 | -0.050 | 0.196 |
| Q9JJL8 | Sars2 | 0.446 | 0.703 | 0.281 | 0.000 | -0.050 | 0.559 | -0.022 | 0.196 |
| Q8CGC7 | Eprs | 0.795 | 1.000 | 0.120 | 0.000 | -0.150 | 0.337 | -0.462 | -0.109 |
| P57780 | Actn4 | 0.021 | 0.091 | -0.172 | 0.037 | 0.000 | -0.274 | -0.547 | -0.455 |
| P14152 | Mdh1 | 0.000 | 0.009 | -0.168 | 0.000 | 0.080 | -1.560 | -1.840 | -1.800 |
| Q6P4T2 | Snrnp200 | 0.013 | 0.069 | -0.020 | 0.137 | 0.000 | -0.250 | -0.524 | -0.332 |
| Q99JR1 | Sfxn1 | 0.018 | 0.083 | 0.000 | 0.180 | -0.179 | 0.403 | 0.408 | 0.393 |
| Q9DCD0 | Pgd | 0.071 | 0.195 | 0.917 | 0.000 | -0.095 | -0.280 | -1.050 | -0.718 |
| Q9Z0H3 | Smarcb1 | 0.030 | 0.113 | 0.000 | -0.066 | 0.179 | -0.294 | -0.516 | -0.219 |
| O89086 | Rbm3 | 0.009 | 0.056 | -0.209 | 0.057 | 0.000 | -0.430 | -0.622 | -0.681 |
| Q922V4 | Plrg1 | 0.064 | 0.181 | 0.000 | 0.843 | -0.233 | -0.721 | -0.595 | -0.591 |
| Q6A026 | Pds5a | 0.449 | 0.707 | -0.685 | 0.000 | 0.044 | -0.448 | -0.612 | -0.234 |
| Q80Y14 | Glrx5 | 0.112 | 0.265 | 0.079 | 0.000 | -0.218 | 0.910 | 0.185 | 0.266 |
| P50136 | Bckdha | 0.099 | 0.243 | 0.031 | 0.000 | -0.341 | 0.421 | 0.014 | 0.389 |
| Q31125 | Slc39a7 | 0.027 | 0.105 | 0.000 | -0.045 | 0.406 | 0.591 | 0.957 | 1.230 |
| P54071 | Idh2 | 0.004 | 0.036 | 0.054 | 0.000 | -0.276 | 1.440 | 0.889 | 0.986 |
| F7BJB9 | Morc3 | 0.118 | 0.274 | 0.164 | -0.166 | 0.000 | -0.243 | -0.973 | -0.297 |
| Q61584 | Fxr1 | 0.129 | 0.291 | -0.026 | 0.120 | 0.000 | -0.203 | -0.299 | 0.011 |
| Q921T2 | Tor1aip1 | 0.690 | 0.965 | -0.053 | 0.000 | 0.307 | 0.445 | -0.208 | 0.311 |
| Q7TT37 | Ikbkap | 0.264 | 0.481 | 0.088 | 0.000 | -0.089 | 0.136 | -0.527 | -0.447 |
| Q9EQ61 | Pes1 | 0.132 | 0.295 | 0.201 | 0.000 | -0.198 | -0.092 | -0.992 | -0.526 |
| Q8C1A5 | Thop1 | 0.006 | 0.046 | 0.158 | -0.220 | 0.000 | -0.983 | -1.770 | -1.750 |
| O55222 | Ilk | 0.020 | 0.087 | 0.022 | 0.000 | -0.417 | 0.976 | 0.527 | 0.493 |
| Q91VV4 | Dennd2d | 0.016 | 0.079 | -0.287 | 0.000 | 0.220 | -0.634 | -0.864 | -0.591 |
| Q80XI4 | Pip4k2b | 0.801 | 1.000 | 0.042 | -0.073 | 0.000 | 0.261 | -0.324 | -0.109 |

| | | | | | | | | | |
|---|---|---|---|---|---|---|---|---|---|
| Q91V41 | Rab14 | 0.967 | 1.000 | 0.000 | 0.178 | -0.083 | 0.345 | -0.065 | -0.209 |
| P56960 | Exosc10 | 0.282 | 0.506 | 0.835 | 0.000 | -0.252 | 0.633 | 0.441 | 0.788 |
| Q8BXA1 | Golim4 | 0.007 | 0.050 | 0.892 | 0.000 | -0.477 | 2.510 | 2.100 | 2.200 |
| Q9WTP7 | Ak3 | 0.037 | 0.129 | -0.146 | 0.101 | 0.000 | 0.809 | 0.233 | 0.598 |
| Q3U0V1 | Khsrp | 0.421 | 0.676 | -0.157 | 0.000 | 0.015 | -0.296 | -0.296 | 0.081 |
| O54692 | Zw10 | 0.471 | 0.730 | 0.324 | -0.222 | 0.000 | 0.295 | 0.022 | 0.210 |
| Q9DBR7 | Ppp1r12a | 0.013 | 0.071 | -0.181 | 0.000 | 0.044 | -0.554 | -1.010 | -0.600 |
| Q61881 | Mcm7 | 0.037 | 0.127 | 0.253 | 0.000 | -0.194 | -0.352 | -1.070 | -0.804 |
| Q9EP69 | Sacm1l | 0.008 | 0.051 | -0.137 | 0.052 | 0.000 | 0.613 | 0.321 | 0.520 |
| P63330 | Ppp2ca | 0.032 | 0.118 | 0.000 | -0.177 | 0.241 | -0.502 | -0.750 | -0.337 |
| Q9CQE5 | Rgs10 | 0.001 | 0.018 | -0.183 | 0.000 | 0.053 | -1.900 | -1.880 | -1.380 |
| Q6NVE9 | Pptc7 | 0.557 | 0.825 | 0.000 | -0.054 | 0.016 | 0.205 | -0.490 | -0.140 |
| Q7TMF3 | Ndufa12 | 0.011 | 0.064 | -0.161 | 0.000 | 0.141 | 0.792 | 0.547 | 1.110 |
| O35874 | Slc1a4 | 0.008 | 0.051 | -2.820 | 0.000 | 0.039 | 3.660 | 3.800 | 3.980 |
| P11032 | Gzma | 0.448 | 0.706 | -1.520 | 0.000 | 0.204 | -1.490 | -1.050 | -0.367 |
| Q80VW7 | Akna | 0.096 | 0.238 | -0.012 | 0.069 | 0.000 | -0.965 | -1.600 | -0.137 |
| Q921M7 | Fam49b | 0.000 | 0.009 | 0.127 | 0.000 | -0.104 | -1.080 | -1.160 | -1.120 |
| O35465 | Fkbp8 | 0.521 | 0.786 | 0.000 | 0.167 | -0.029 | 0.317 | -0.227 | -0.452 |
| P12970 | Rpl7a | 0.049 | 0.154 | 0.141 | -0.105 | 0.000 | 0.570 | 0.175 | 0.428 |
| O35643 | Ap1b1 | 0.185 | 0.374 | 0.145 | 0.000 | -0.102 | -0.013 | -0.683 | -0.260 |
| P61982 | Ywhag | 0.016 | 0.078 | 0.076 | 0.000 | -0.065 | -0.399 | -0.977 | -0.678 |
| Q9CXY6 | Ilf2 | 0.540 | 0.807 | 0.000 | 0.631 | -0.035 | -0.337 | 0.060 | 0.300 |
| O70503 | Hsd17b12 | 0.014 | 0.072 | 0.001 | 0.000 | -0.328 | 1.430 | 0.851 | 0.653 |
| Q62376 | Snrnp70 | 0.125 | 0.285 | -0.253 | 0.000 | 1.590 | -0.713 | -0.893 | -0.482 |
| P62702 | Rps4x | 0.826 | 1.000 | 0.036 | 0.000 | -0.324 | 0.307 | -0.757 | -0.072 |
| Q61029 | Tmpo | 0.756 | 1.000 | -0.325 | 0.000 | 0.312 | -0.177 | -0.353 | 0.258 |
| Q8K3G5 | Vrk3 | 0.070 | 0.193 | -0.071 | 0.000 | 0.328 | -0.503 | -0.539 | -0.092 |
| Q8R016 | Blmh | 0.021 | 0.091 | -0.097 | 0.256 | 0.000 | -1.320 | -1.100 | -0.472 |
| A2AF47 | Dock11 | 0.001 | 0.018 | 0.000 | 0.134 | -0.011 | -0.552 | -0.771 | -0.648 |
| Q9EQQ9 | Mgea5 | 0.066 | 0.186 | -0.118 | 0.185 | 0.000 | -0.295 | -0.900 | -0.343 |
| Q8VCW4 | Unc93b1 | 0.699 | 0.975 | 0.532 | -0.017 | 0.000 | 0.616 | 0.022 | 0.192 |
| Q60932 | Vdac1 | 0.006 | 0.046 | 0.000 | 0.313 | -0.052 | 1.240 | 1.010 | 1.700 |
| P68040 | Rack1 | 0.095 | 0.237 | 0.132 | 0.000 | -0.220 | -0.151 | -0.721 | -0.490 |
| Q6ZQI3 | Mlec | 0.323 | 0.558 | 0.076 | 0.000 | -0.122 | 1.290 | -0.531 | 1.210 |
| Q60597 | Ogdh | 0.007 | 0.048 | 0.000 | 0.045 | -0.239 | 0.747 | 0.432 | 0.587 |
| Q61316 | Hspa4 | 0.002 | 0.027 | 0.000 | -0.007 | 0.001 | -0.700 | -1.140 | -0.857 |
| P26369 | U2af2 | 0.063 | 0.181 | 0.204 | 0.000 | -0.149 | -0.162 | -0.840 | -0.740 |
| Q9ESZ8 | Gtf2i | 0.049 | 0.154 | -0.230 | 0.072 | 0.000 | -0.284 | -0.490 | -0.318 |
| O08900 | Ikzf3 | 0.585 | 0.857 | -0.536 | 0.000 | 0.072 | -0.517 | -0.398 | 0.008 |
| Q99P88 | Nup155 | 0.894 | 1.000 | -0.030 | 0.199 | 0.000 | 0.139 | -0.001 | -0.006 |
| Q99K85 | Psat1 | 0.006 | 0.044 | 0.000 | 0.027 | -0.015 | -0.509 | -0.996 | -0.778 |
| Q69Z99 | Znf512 | 0.404 | 0.655 | -0.111 | 0.000 | 0.395 | -0.071 | -0.745 | 0.203 |
| Q78IK4 | Apool | 0.010 | 0.060 | -0.062 | 0.205 | 0.000 | 0.712 | 0.459 | 0.823 |
| P35821 | Ptpn1 | 0.031 | 0.116 | -0.040 | 0.235 | 0.000 | -0.162 | -0.284 | -0.302 |
| Q91YT0 | Ndufv1 | 0.001 | 0.019 | 0.000 | 0.034 | -0.076 | 0.817 | 0.553 | 0.688 |
| Q91ZA3 | Pcca | 0.004 | 0.037 | 0.000 | 0.273 | -0.086 | 0.822 | 0.667 | 0.847 |
| P46471 | Psmc2 | 0.133 | 0.297 | 0.000 | 0.036 | -0.095 | -0.069 | -0.984 | -0.512 |
| Q65Z40 | Wapl | 0.122 | 0.280 | 0.000 | 0.100 | -0.214 | -0.278 | -0.555 | -0.160 |
| Q5SSI6 | Utp18 | 0.118 | 0.274 | 0.035 | 0.000 | -0.326 | -0.227 | -0.724 | -0.437 |
| Q61464 | Znf638 | 0.087 | 0.224 | -0.106 | 0.232 | 0.000 | -0.212 | -0.535 | -0.164 |
| Q8VDP4 | Ccar2 | 0.001 | 0.019 | -0.213 | 0.000 | 0.026 | -1.020 | -1.310 | -1.030 |
| P54923 | Adprh | 0.020 | 0.089 | -0.015 | 0.000 | 0.174 | -1.590 | -3.200 | -1.430 |
| Q9QZQ8 | H2afy | 0.640 | 0.916 | 0.308 | -0.121 | 0.000 | 0.027 | 0.161 | 0.210 |
| Q3UV70 | Pdp1 | 0.009 | 0.056 | -0.168 | 0.000 | 0.071 | 0.763 | 0.401 | 0.775 |
| Q922D8 | Mthfd1 | 0.080 | 0.212 | 0.193 | 0.000 | -0.406 | -0.292 | -0.898 | -0.884 |
| Q6ZQ38 | Cand1 | 0.009 | 0.057 | -0.021 | 0.028 | 0.000 | -0.389 | -0.813 | -0.557 |
| P02468 | Lamc1 | 0.000 | 0.011 | 0.000 | 0.405 | -0.653 | 4.850 | 4.390 | 4.910 |
| Q9WUP7 | Uchl5 | 0.001 | 0.022 | 0.106 | 0.000 | -0.173 | -0.684 | -0.856 | -0.786 |
| O70133 | Dhx9 | 0.008 | 0.054 | -0.147 | 0.112 | 0.000 | -0.357 | -0.417 | -0.383 |

| | | | | | | | | | |
|---|---|---|---|---|---|---|---|---|---|
| P57776 | Eef1d | 0.010 | 0.058 | 0.131 | -0.085 | 0.000 | -0.355 | -0.700 | -0.662 |
| Q9D1J1 | Necap2 | 0.072 | 0.197 | 0.000 | 0.059 | -0.305 | -0.584 | -0.378 | -1.140 |
| P51174 | Acadl | 0.039 | 0.132 | 0.000 | 0.343 | -0.012 | 0.526 | 0.424 | 0.475 |
| Q8VI75 | Ipo4 | 0.133 | 0.296 | 0.315 | -0.020 | 0.000 | 1.170 | 0.277 | 0.490 |
| O55131 | Sept 7 | 0.373 | 0.621 | 0.000 | 0.100 | -0.111 | 0.425 | -0.163 | 0.292 |
| Q9D0F3 | Lman1 | 0.008 | 0.054 | 0.000 | 0.082 | -0.462 | 2.150 | 1.680 | 1.050 |
| P06797 | Ctsl | 0.055 | 0.166 | 0.060 | -0.053 | 0.000 | -0.223 | -0.865 | -0.442 |
| Q8K4Z3 | Naxe | 0.013 | 0.071 | 0.012 | 0.000 | -0.125 | 0.324 | 0.340 | 0.156 |
| P26638 | Sars | 0.134 | 0.298 | 0.000 | 0.015 | -0.002 | -0.210 | -0.327 | 0.004 |
| Q9ERK4 | Cse1l | 0.041 | 0.138 | 0.073 | -0.029 | 0.000 | -0.350 | -1.430 | -0.960 |
| Q9R233 | Tapbp | 0.550 | 0.819 | -0.278 | 0.096 | 0.000 | -0.148 | -0.045 | -0.232 |
| Q78ZA7 | Nap1l4 | 0.009 | 0.056 | -0.293 | 0.000 | 0.003 | -0.919 | -1.530 | -0.967 |
| Q9D7S9 | Chmp5 | 0.646 | 0.923 | 0.226 | -2.780 | 0.000 | -0.323 | -0.670 | -0.105 |
| Q8CHY6 | Gatad2a | 0.072 | 0.196 | -0.214 | 0.000 | 0.239 | -0.313 | -1.050 | -0.493 |
| Q5SQX6 | Cyfip2 | 0.133 | 0.297 | 0.000 | -0.073 | 0.396 | -0.157 | -0.338 | -0.099 |
| Q61103 | Dpf2 | 0.005 | 0.039 | -0.147 | 0.000 | 0.059 | -0.516 | -0.628 | -0.419 |
| Q80TH2 | Erbin | 0.102 | 0.248 | 0.004 | 0.000 | -0.191 | 0.359 | -0.014 | 0.435 |
| Q9DB29 | Iah1 | 0.002 | 0.024 | -0.084 | 0.000 | 0.019 | -0.725 | -1.100 | -0.844 |
| P08775 | Polr2a | 0.010 | 0.059 | 0.000 | -0.066 | 0.050 | -0.270 | -0.476 | -0.290 |
| A2ADY9 | Ddi2 | 0.042 | 0.140 | 0.078 | -0.450 | 0.000 | -0.584 | -1.520 | -1.060 |
| Q9Z0M6 | Cd97 | 0.222 | 0.423 | 0.067 | -0.258 | 0.000 | -0.186 | -0.412 | -0.150 |
| Q99KI0 | Aco2 | 0.135 | 0.299 | 0.000 | 0.042 | -0.166 | 0.553 | 0.087 | 0.138 |
| P26443 | Glud1 | 0.007 | 0.048 | -0.208 | 0.207 | 0.000 | 0.756 | 0.579 | 0.813 |
| P01899 | H2-D1 | 0.854 | 1.000 | 0.000 | 0.322 | -0.021 | 0.070 | 0.088 | 0.212 |
| Q9QWR8 | Naga | 0.347 | 0.588 | 0.711 | -0.217 | 0.000 | 0.309 | -0.481 | -0.619 |
| P56395 | Cyb5a | 0.102 | 0.247 | 0.000 | 0.774 | -0.122 | 0.724 | 0.805 | 0.954 |
| Q9QYC0 | Add1 | 0.099 | 0.243 | 0.258 | -0.225 | 0.000 | -0.105 | -0.792 | -0.814 |
| Q9DBS1 | Tmem43 | 0.001 | 0.019 | 0.127 | -0.398 | 0.000 | 1.340 | 1.310 | 1.460 |
| Q8CCF0 | Prpf31 | 0.340 | 0.579 | 0.000 | -0.040 | 0.902 | 0.248 | -0.333 | -0.206 |
| Q8BMF4 | Dlat | 0.002 | 0.025 | 0.000 | 0.245 | -0.071 | 1.060 | 0.823 | 1.080 |
| P70398 | Usp9x | 0.156 | 0.330 | 0.000 | 0.167 | -0.051 | 0.059 | -0.847 | -0.528 |
| P09103 | P4hb | 0.001 | 0.019 | 0.181 | 0.000 | -0.258 | 1.550 | 1.180 | 1.470 |
| P46062 | Sipa1 | 0.122 | 0.280 | 0.000 | -0.687 | 0.146 | -1.190 | -0.856 | -0.453 |
| Q8BRH0 | Tmtc3 | 0.001 | 0.019 | -0.049 | 0.564 | 0.000 | 2.350 | 1.990 | 2.490 |
| Q9Z2I9 | Sucla2 | 0.144 | 0.311 | 0.045 | 0.000 | -0.226 | 0.968 | 0.063 | 0.320 |
| Q9DCS9 | Ndufb10 | 0.006 | 0.043 | 0.000 | 0.250 | -0.106 | 0.628 | 0.918 | 0.805 |
| P17751 | Tpi1 | 0.001 | 0.019 | -0.153 | 0.052 | 0.000 | -1.010 | -1.410 | -1.170 |
| Q99MN1 | Kars | 0.095 | 0.237 | 0.000 | 0.246 | -0.487 | -0.474 | -0.558 | -0.660 |
| Q8C3X8 | Lmf2 | 0.001 | 0.022 | -0.449 | 0.291 | 0.000 | 1.800 | 2.180 | 2.370 |
| Q8R123 | Flad1 | 0.186 | 0.376 | 0.107 | 0.000 | -0.001 | -0.107 | -0.914 | -0.144 |
| Q8VCT3 | Rnpep | 0.006 | 0.047 | 0.307 | 0.000 | -0.067 | -0.773 | -1.480 | -1.230 |
| Q8BGW0 | Themis | 0.127 | 0.288 | -1.320 | 0.000 | 0.443 | -1.890 | -1.260 | -1.100 |
| Q8CI94 | Pygb | 0.009 | 0.058 | -0.215 | 0.120 | 0.000 | -0.661 | -1.230 | -0.895 |
| P70295 | Aup1 | 0.223 | 0.424 | -0.237 | 0.153 | 0.000 | 0.075 | 0.109 | 0.499 |
| Q8BSY0 | Asph | 0.000 | 0.014 | 0.141 | 0.000 | -0.607 | 3.050 | 2.490 | 2.970 |
| Q91WJ8 | Fubp1 | 0.460 | 0.719 | -0.102 | 0.177 | 0.000 | -0.201 | -0.216 | 0.143 |
| Q8VBT0 | Tmx1 | 0.047 | 0.149 | 0.000 | 0.316 | -0.123 | 0.858 | 0.556 | 0.392 |
| P10852 | Slc3a2 | 0.031 | 0.115 | 0.358 | -0.051 | 0.000 | 1.070 | 0.533 | 0.706 |
| Q6NV83 | U2surp | 0.018 | 0.083 | 0.000 | 0.005 | -0.147 | -0.422 | -0.985 | -0.714 |
| Q61937 | Npm1 | 0.054 | 0.164 | 0.268 | -0.137 | 0.000 | -0.286 | -1.550 | -1.340 |
| Q9JMH6 | Txnrd1 | 0.030 | 0.114 | 0.000 | -0.115 | 0.093 | -0.408 | -1.270 | -1.320 |
| Q8VEK3 | Hnrnpu | 0.023 | 0.096 | 0.000 | 0.135 | -0.052 | -0.278 | -0.465 | -0.202 |
| P62843 | Rps15 | 0.638 | 0.914 | 0.152 | -0.159 | 0.000 | 0.269 | -0.156 | 0.113 |
| P08207 | S100a10 | 0.150 | 0.321 | -1.010 | 0.000 | 0.570 | 0.678 | 0.482 | 0.974 |
| Q9CSH3 | Dis3 | 0.019 | 0.087 | -0.143 | 0.044 | 0.000 | -0.478 | -0.855 | -0.426 |
| Q3UZA1 | Rcsd1 | 0.010 | 0.060 | 0.278 | -0.314 | 0.000 | -0.995 | -1.930 | -1.520 |
| Q8BP92 | Rcn2 | 0.000 | 0.011 | -0.104 | 0.495 | 0.000 | 3.300 | 2.950 | 3.320 |
| P51150 | Rab7a | 0.539 | 0.807 | 0.095 | 0.000 | -0.023 | 0.282 | -0.085 | 0.100 |
| O35226 | Psmd4 | 0.124 | 0.283 | 0.193 | 0.000 | -0.170 | -0.493 | -1.620 | -0.319 |

| | | | | | | | | | |
|---|---|---|---|---|---|---|---|---|---|
| P42669 | Pura | 0.003 | 0.033 | -0.125 | 0.129 | 0.000 | -0.642 | -0.748 | -0.504 |
| Q9D2G2 | Dlst | 0.003 | 0.033 | 0.000 | 0.150 | -0.204 | 0.862 | 0.626 | 0.867 |
| Q9CRB2 | Nhp2 | 0.145 | 0.314 | 0.135 | -0.347 | 0.000 | -0.114 | -0.877 | -0.684 |
| Q9WV32 | Arpc1b | 0.004 | 0.034 | 0.000 | -0.036 | 0.060 | 0.284 | 0.204 | 0.307 |
| Q8CDM1 | Atad2 | 0.200 | 0.394 | -0.108 | 0.046 | 0.000 | -0.369 | -0.528 | 0.035 |
| Q9D154 | Serpinb1a | 0.050 | 0.155 | 1.640 | 0.000 | -0.016 | -0.530 | -1.950 | -1.950 |
| Q8BVE3 | Atp6v1h | 0.761 | 1.000 | 0.447 | 0.000 | -0.035 | 0.476 | -0.415 | 0.057 |
| O09110 | Map2k3 | 0.051 | 0.158 | 0.000 | 0.028 | -0.378 | 0.174 | 0.286 | 0.414 |
| P14206 | Rpsa | 0.136 | 0.301 | 0.104 | -0.167 | 0.000 | -0.043 | -0.645 | -0.462 |
| P62849 | Rps24 | 0.381 | 0.630 | 0.000 | 0.389 | -0.243 | 0.128 | 0.548 | 0.148 |
| Q8K224 | Nat10 | 0.136 | 0.300 | 0.000 | 0.122 | -0.081 | 0.017 | -0.643 | -0.503 |
| P63101 | Ywhaz | 0.001 | 0.015 | 0.000 | -0.021 | 0.113 | -1.140 | -1.610 | -1.370 |
| Q80ZK0 | Mrps10 | 0.027 | 0.107 | 0.000 | 0.562 | -0.484 | 1.350 | 0.904 | 1.690 |
| O08547 | Sec22b | 0.089 | 0.229 | -0.213 | 0.184 | 0.000 | 0.660 | 0.106 | 0.678 |
| Q9WTQ5 | Akap12 | 0.001 | 0.019 | 0.602 | 0.000 | -1.060 | 4.110 | 3.850 | 4.090 |
| P37040 | Por | 0.001 | 0.018 | 0.000 | 0.126 | -0.066 | 0.875 | 0.704 | 0.940 |
| P05064 | Aldoa | 0.039 | 0.132 | 0.022 | -0.139 | 0.000 | -0.265 | -0.709 | -0.415 |
| Q8CGY8 | Ogt | 0.046 | 0.149 | 0.000 | -0.266 | 0.000 | -0.418 | -0.946 | -0.498 |
| P00920 | Ca2 | 0.051 | 0.158 | 0.887 | 0.000 | -0.365 | -0.645 | -2.020 | -2.740 |
| Q9CRB9 | Chchd3 | 0.005 | 0.039 | -0.052 | 0.233 | 0.000 | 0.868 | 0.646 | 1.000 |
| P63017 | Hspa8 | 0.008 | 0.054 | 0.046 | 0.000 | -0.090 | -0.386 | -0.759 | -0.718 |
| Q9DAM5 | Slc25a19 | 0.512 | 0.776 | 0.000 | 0.235 | -0.215 | 0.146 | 0.162 | -1.530 |
| Q8BG32 | Psmd11 | 0.097 | 0.240 | 0.000 | 0.008 | -0.152 | -0.205 | -0.649 | -0.258 |
| Q9Z2U0 | Psma7 | 0.001 | 0.015 | -0.029 | 0.109 | 0.000 | -1.070 | -1.380 | -1.020 |
| Q7M6Y3 | Picalm | 0.391 | 0.640 | -0.430 | 1.660 | 0.000 | 1.050 | 1.440 | 0.683 |
| Q8BVK9 | Sp110 | 0.142 | 0.309 | -0.296 | 0.000 | 0.048 | -0.447 | -0.417 | -0.158 |
| Q99J39 | Mlycd | 0.122 | 0.280 | 0.000 | 0.662 | -0.127 | 0.733 | 0.678 | 0.583 |
| A2AN08 | Ubr4 | 0.208 | 0.404 | 0.203 | 0.000 | -0.345 | 0.064 | -0.983 | -0.892 |
| Q9EPK7 | Xpo7 | 0.014 | 0.072 | 0.000 | -0.014 | 0.137 | -0.561 | -1.410 | -1.080 |
| Q9DAW9 | Cnn3 | 0.001 | 0.019 | 1.350 | 0.000 | -0.597 | 6.050 | 5.170 | 5.740 |
| Q9CQ65 | Mtap | 0.004 | 0.036 | 0.000 | -0.105 | 0.108 | -1.190 | -1.490 | -0.845 |
| Q99LI7 | Cstf3 | 0.246 | 0.457 | -0.352 | 0.110 | 0.000 | -0.471 | -0.629 | -0.048 |
| Q9D0I9 | Rars | 0.267 | 0.485 | 0.171 | 0.000 | -0.071 | 0.100 | -0.634 | -0.231 |
| P08228 | Sod1 | 0.000 | 0.013 | -0.224 | 0.010 | 0.000 | -1.080 | -1.290 | -1.170 |
| Q99LC5 | Etfa | 0.310 | 0.542 | 0.000 | 0.213 | -0.198 | 0.705 | -0.164 | 0.467 |
| Q6DIC0 | Smarca2 | 0.021 | 0.090 | 0.000 | 0.161 | -0.064 | -0.474 | -0.802 | -0.336 |
| Q8CFE3 | Rcor1 | 0.012 | 0.068 | 0.000 | -0.039 | 0.014 | -0.534 | -1.040 | -0.568 |
| P61965 | Wdr5 | 0.036 | 0.125 | 0.000 | -0.113 | 0.084 | -0.312 | -0.817 | -0.427 |
| Q9CQQ7 | Atp5f1 | 0.000 | 0.014 | 0.081 | 0.000 | -0.116 | 1.040 | 0.810 | 0.987 |
| Q8R1Q8 | Dync1li1 | 0.384 | 0.633 | 0.000 | -0.009 | 0.173 | 0.263 | -0.011 | 0.215 |
| Q9CQF3 | Nudt21 | 0.017 | 0.080 | 0.226 | 0.000 | -0.079 | -0.412 | -0.346 | -0.277 |
| Q60864 | Stip1 | 0.004 | 0.035 | -0.047 | 0.027 | 0.000 | -0.677 | -1.220 | -1.030 |
| Q8CFI7 | Polr2b | 0.032 | 0.118 | -0.057 | 0.000 | 0.025 | -0.407 | -0.600 | -0.191 |
| Q8K124 | Plekho2 | 0.967 | 1.000 | 0.080 | -0.182 | 0.000 | 0.317 | -0.430 | -0.020 |
| Q8BH04 | Pck2 | 0.009 | 0.056 | 0.068 | 0.000 | -0.323 | 1.320 | 0.741 | 0.823 |
| Q99PT1 | Arhgdia | 0.005 | 0.039 | -0.050 | 0.000 | 0.040 | -1.120 | -1.610 | -0.907 |
| Q9JL26 | Fmnl1 | 0.003 | 0.033 | -0.034 | 0.000 | 0.049 | -0.746 | -1.290 | -0.945 |
| Q9Z0V7 | Timm17b | 0.705 | 0.982 | 0.000 | -0.059 | 0.096 | 0.339 | -0.298 | 0.243 |
| Q9WTQ8 | Timm23 | 0.001 | 0.017 | 0.000 | 0.198 | -0.057 | 0.799 | 0.774 | 0.865 |
| P28352 | Apex1 | 0.028 | 0.107 | 0.000 | -0.288 | 0.047 | -0.452 | -0.881 | -0.582 |
| O88874 | Ccnk | 0.240 | 0.449 | 0.303 | -0.212 | 0.000 | -1.210 | -0.367 | 0.032 |
| Q8R2E9 | Ero1b | 0.121 | 0.278 | 0.069 | 0.000 | -0.488 | -0.320 | -0.915 | -0.631 |
| Q80XN0 | Bdh1 | 0.208 | 0.404 | 0.000 | 0.147 | -0.057 | 0.047 | -0.615 | -0.246 |
| Q9CWJ9 | Atic | 0.007 | 0.048 | 0.044 | -0.094 | 0.000 | -0.791 | -1.580 | -1.370 |
| Q3TEA8 | Hp1bp3 | 0.107 | 0.256 | -0.152 | 0.000 | 0.110 | -0.294 | -0.484 | -0.100 |
| Q61081 | Cdc37 | 0.002 | 0.027 | 0.019 | 0.000 | -0.009 | -0.837 | -1.370 | -1.050 |
| Q61543 | Glg1 | 0.076 | 0.203 | -0.003 | 1.010 | 0.000 | 1.350 | 1.020 | 1.130 |
| P19426 | Nelfe | 0.017 | 0.081 | -0.074 | 0.000 | 0.154 | -0.345 | -0.656 | -0.356 |
| P27612 | Plaa | 0.023 | 0.096 | -0.191 | 0.007 | 0.000 | -0.710 | -1.900 | -1.320 |

| | | | | | | | | | |
|---|---|---|---|---|---|---|---|---|---|
| Q9CXI5 | Manf | 0.067 | 0.186 | 0.000 | 0.366 | -0.328 | 0.585 | 0.395 | 0.788 |
| Q3TCN2 | Plbd2 | 0.916 | 1.000 | 0.612 | -0.005 | 0.000 | 0.484 | -0.125 | 0.341 |
| P27546 | Map4 | 0.159 | 0.336 | -0.314 | 0.000 | 0.103 | 0.247 | 0.014 | 0.453 |
| A2AR02 | Ppig | 0.007 | 0.048 | 0.137 | 0.000 | -0.311 | -0.770 | -1.200 | -1.030 |
| Q9DCH4 | Eif3f | 0.200 | 0.395 | 0.000 | 0.017 | -0.327 | -0.119 | -0.590 | -0.416 |
| Q61598 | Gdi2 | 0.000 | 0.013 | -0.038 | 0.000 | 0.024 | -0.676 | -0.913 | -0.798 |
| Q8BVI4 | Qdpr | 0.149 | 0.319 | 0.000 | -0.293 | 0.016 | -0.220 | -0.562 | -0.276 |
| Q9CQ62 | Decr1 | 0.431 | 0.687 | -0.053 | 0.110 | 0.000 | 0.728 | -0.038 | 0.024 |
| P58681 | Tlr7 | 0.683 | 0.960 | 0.829 | -0.061 | 0.000 | 0.779 | 0.114 | 0.332 |
| Q6GV12 | Kdsr | 0.056 | 0.168 | -1.720 | 0.162 | 0.000 | 0.995 | 0.983 | 1.380 |
| Q8VEK0 | Tmem30a | 0.141 | 0.308 | 0.264 | 0.000 | -0.064 | 0.477 | 0.148 | 0.337 |
| Q8CDG3 | Vcpip1 | 0.072 | 0.197 | -0.095 | 0.062 | 0.000 | -0.188 | -1.310 | -0.985 |
| P62754 | Rps6 | 0.169 | 0.350 | 0.062 | -0.038 | 0.000 | 0.478 | -0.019 | 0.316 |
| Q8VBZ3 | Clptm1 | 0.007 | 0.050 | 0.000 | 0.091 | -0.179 | 0.984 | 0.517 | 0.882 |
| Q8VDM6 | Hnrnpul1 | 0.168 | 0.350 | 0.000 | 0.612 | -0.432 | -0.761 | -0.381 | -0.344 |
| Q569Z5 | Ddx46 | 0.021 | 0.092 | 0.180 | -0.247 | 0.000 | -0.684 | -1.790 | -1.560 |
| O35286 | Dhx15 | 0.013 | 0.071 | -0.082 | 0.000 | 0.065 | -0.392 | -0.774 | -0.454 |
| Q9JM14 | Nt5c | 0.002 | 0.023 | -0.329 | 0.393 | 0.000 | -1.550 | -1.680 | -1.570 |
| O08539 | Bin1 | 0.015 | 0.076 | -0.300 | 0.232 | 0.000 | -0.726 | -1.520 | -1.220 |
| Q9WV85 | Nme3 | 0.093 | 0.234 | 0.001 | -0.129 | 0.000 | -0.343 | -0.578 | -0.122 |
| Q91YW3 | Dnajc3 | 0.001 | 0.022 | 0.000 | 0.021 | -0.207 | 1.010 | 0.731 | 0.763 |
| Q9CT10 | Ranbp3 | 0.004 | 0.038 | -0.211 | 0.000 | 0.146 | -1.020 | -1.690 | -1.230 |
| Q8K1B8 | Fermt3 | 0.253 | 0.466 | 0.000 | 0.165 | -0.366 | -0.172 | -0.300 | -0.423 |
| P47962 | Rpl5 | 0.228 | 0.432 | 0.125 | -0.082 | 0.000 | 0.097 | -0.370 | -0.689 |
| Q9Z1G4 | Atp6v0a1 | 0.225 | 0.427 | 0.713 | -0.329 | 0.000 | 2.070 | -0.094 | 1.480 |
| Q9EQK5 | Mvp | 0.055 | 0.166 | -0.217 | 0.000 | 0.037 | 0.473 | 0.098 | 0.340 |
| Q99J99 | Mpst | 0.334 | 0.571 | 0.163 | 0.000 | -0.160 | 0.378 | -0.932 | -0.891 |
| Q9D967 | Mdp1 | 0.009 | 0.057 | -0.283 | 0.000 | 0.020 | -2.080 | -3.970 | -2.490 |
| O88487 | Dync1i2 | 0.412 | 0.665 | 0.000 | 0.172 | -0.041 | 0.321 | -0.021 | 0.156 |
| Q8C129 | Lnpep | 0.689 | 0.965 | 0.030 | 0.000 | -0.056 | 0.124 | -0.155 | 0.131 |
| Q3UQ84 | Tars2 | 0.008 | 0.053 | 0.000 | 0.010 | -0.095 | 0.543 | 0.329 | 0.293 |
| Q8CD10 | Micu2 | 0.000 | 0.014 | -0.023 | 0.080 | 0.000 | 1.030 | 0.770 | 0.952 |
| P61290 | Psme3 | 0.001 | 0.021 | 0.000 | 0.420 | -0.049 | -1.120 | -1.150 | -1.340 |
| Q8JZN5 | Acad9 | 0.215 | 0.412 | 0.000 | 0.153 | -0.210 | 0.571 | 0.123 | 0.083 |
| Q8CJF7 | Ahctf1 | 0.969 | 1.000 | -0.733 | 0.000 | 0.004 | -0.494 | -0.354 | 0.158 |
| P35293 | Rab18 | 0.022 | 0.093 | 0.000 | 0.058 | -0.235 | 0.574 | 0.244 | 0.439 |
| Q9JLI8 | Sart3 | 0.010 | 0.058 | 0.000 | 0.228 | -0.111 | -0.934 | -1.600 | -0.875 |
| Q08288 | Lyar | 0.211 | 0.408 | 0.005 | 0.000 | -0.310 | -0.096 | -0.931 | -0.453 |
| P53994 | Rab2a | 0.195 | 0.387 | 0.000 | 0.207 | -0.108 | 0.262 | 0.082 | 0.656 |
| P62774 | Mtpn | 0.005 | 0.041 | 0.000 | -0.056 | 0.340 | -0.915 | -1.670 | -1.290 |
| Q7TQI3 | Otub1 | 0.007 | 0.048 | 0.000 | -0.084 | 0.037 | -0.752 | -1.500 | -1.260 |
| P58252 | Eef2 | 0.192 | 0.384 | 0.070 | 0.000 | -0.273 | -0.046 | -0.716 | -0.492 |
| Q91V61 | Sfxn3 | 0.006 | 0.043 | -0.166 | 0.192 | 0.000 | 0.721 | 0.546 | 0.762 |
| Q68FD5 | Cltc | 0.377 | 0.625 | 0.000 | 0.035 | -0.230 | 0.063 | -0.437 | -0.339 |
| P68510 | Ywhah | 0.001 | 0.015 | -0.038 | 0.020 | 0.000 | -0.755 | -1.060 | -0.878 |
| Q91WD5 | Ndufs2 | 0.006 | 0.045 | 0.000 | 0.406 | -0.118 | 0.946 | 0.914 | 1.040 |
| P27773 | Pdia3 | 0.006 | 0.047 | 0.000 | 0.162 | -0.091 | 0.836 | 0.494 | 0.787 |
| Q9R0E2 | Plod1 | 0.000 | 0.010 | 0.686 | -0.258 | 0.000 | 4.510 | 4.250 | 4.540 |
| Q9JI11 | Stk4 | 0.009 | 0.057 | 0.000 | -0.122 | 0.047 | -0.744 | -1.550 | -1.450 |
| A2BH40 | Arid1a | 0.092 | 0.233 | -0.206 | 0.044 | 0.000 | -0.488 | -0.528 | -0.129 |
| Q9ERR7 | Sep 15 | 0.054 | 0.164 | 0.000 | 0.000 | 0.112 | 0.582 | 0.658 | 0.160 |
| Q8BKZ9 | Pdhx | 0.019 | 0.086 | 0.000 | 0.072 | -0.571 | 0.880 | 0.548 | 0.654 |
| P00405 | Mtco2 | 0.007 | 0.048 | 0.000 | 0.129 | -0.089 | 0.646 | 0.391 | 0.643 |
| Q3UVK0 | Ermp1 | 0.187 | 0.376 | 0.000 | 0.130 | -0.198 | 0.330 | -0.022 | 0.382 |
| Q8BL66 | Eea1 | 0.108 | 0.259 | -0.525 | 0.502 | 0.000 | -0.939 | -0.986 | -0.339 |
| P40336 | Vps26a | 0.557 | 0.825 | 0.032 | 0.000 | -0.325 | 0.061 | -0.478 | -0.248 |
| O08795 | Prkcsh | 0.006 | 0.047 | 0.000 | 0.339 | -0.092 | 0.821 | 0.722 | 0.867 |
| Q9WVK4 | Ehd1 | 0.325 | 0.561 | 0.320 | 0.000 | -0.219 | 0.523 | 0.037 | 0.249 |
| Q8VCF0 | Mavs | 0.900 | 1.000 | -0.276 | 0.000 | 0.068 | -0.034 | -0.283 | 0.051 |

| | | | | | | | | | |
|---|---|---|---|---|---|---|---|---|---|
| P62301 | Rps13 | 0.889 | 1.000 | 0.000 | 0.263 | -0.044 | -0.105 | 0.013 | 0.390 |
| O08917 | Flot1 | 0.054 | 0.164 | 0.190 | -0.046 | 0.000 | 0.567 | 0.218 | 0.752 |
| Q9D898 | Arpc5l | 0.171 | 0.353 | -0.177 | 0.054 | 0.000 | 0.015 | 0.105 | 0.281 |
| A2AJI0 | Map7d1 | 0.249 | 0.462 | -0.128 | 0.647 | 0.000 | 1.090 | 0.041 | 1.120 |
| Q9JMA1 | Usp14 | 0.074 | 0.201 | 0.000 | 0.381 | -0.373 | -0.303 | -0.956 | -1.110 |
| Q60930 | Vdac2 | 0.002 | 0.023 | 0.000 | 0.161 | -0.063 | 1.040 | 0.946 | 1.360 |
| P62900 | Rpl31 | 0.502 | 0.767 | 0.126 | -0.033 | 0.000 | 0.385 | -0.004 | 0.012 |
| F8VPU2 | Farp1 | 0.001 | 0.014 | -0.677 | 0.000 | 0.355 | 4.280 | 3.500 | 3.700 |
| Q9JMG7 | Hdgfrp3 | 0.136 | 0.301 | -0.347 | 0.000 | 0.147 | -0.879 | -0.986 | -0.099 |
| Q922L6 | Nelfcd | 0.012 | 0.066 | 0.000 | -0.045 | 0.091 | -0.415 | -0.543 | -0.247 |
| Q52KI8 | Srrm1 | 0.120 | 0.276 | 0.238 | -0.450 | 0.000 | -0.234 | -2.830 | -1.770 |
| Q8BG81 | Poldip3 | 0.098 | 0.242 | -0.525 | 0.000 | 0.195 | -0.771 | -0.787 | -0.391 |
| Q921Z5 | Tnfaip8 | 0.082 | 0.214 | 0.388 | 0.000 | -0.292 | -0.278 | -0.792 | -0.547 |
| Q3UJU9 | Rmdn3 | 0.332 | 0.570 | 0.215 | 0.000 | -0.020 | -0.077 | -0.868 | 0.115 |
| Q8CB44 | Gramd4 | 0.430 | 0.686 | 0.000 | 0.225 | -0.025 | 0.064 | 0.109 | 0.327 |
| Q922K7 | Nop2 | 0.086 | 0.223 | 0.000 | 0.003 | -0.303 | -0.235 | -0.800 | -0.577 |
| P49717 | Mcm4 | 0.023 | 0.097 | 0.346 | 0.000 | -0.046 | -0.328 | -0.924 | -0.787 |
| Q3UHX2 | Pdap1 | 0.179 | 0.365 | 0.000 | 0.188 | -0.220 | -0.300 | -0.993 | -0.140 |
| Q9R1P1 | Psmb3 | 0.004 | 0.035 | 0.033 | -0.340 | 0.000 | -1.020 | -1.120 | -0.826 |
| Q61941 | Nnt | 0.000 | 0.013 | 1.030 | -0.638 | 0.000 | 5.910 | 5.440 | 5.700 |
| P52293 | Kpna2 | 0.813 | 1.000 | 0.707 | 0.000 | -0.101 | 0.887 | -0.132 | 0.151 |
| Q05512 | Mark2 | 0.392 | 0.641 | 0.223 | -0.177 | 0.000 | 0.293 | -0.698 | -0.474 |
| Q5XJY5 | Arcn1 | 0.085 | 0.221 | 0.106 | -0.155 | 0.000 | -0.126 | -0.943 | -0.727 |
| Q99JR8 | Smarcd2 | 0.103 | 0.250 | 0.000 | -0.058 | 0.257 | -0.113 | -0.328 | -0.116 |
| Q9D0T1 | Snu13 | 0.001 | 0.016 | 0.000 | 0.041 | -0.075 | -0.503 | -0.523 | -0.409 |
| Q8BFZ9 | Erlin2 | 0.006 | 0.047 | 0.000 | 0.127 | -0.748 | 1.380 | 1.200 | 1.190 |
| Q9D0S9 | Hint2 | 0.816 | 1.000 | 0.000 | 1.130 | -1.100 | -0.268 | 1.220 | -0.316 |
| P21107 | Tpm3 | 0.005 | 0.039 | 0.000 | 0.179 | -0.001 | -0.958 | -1.660 | -1.080 |
| Q921S7 | Mrpl37 | 0.249 | 0.462 | 0.174 | 0.000 | -0.144 | 0.783 | -0.030 | 0.302 |
| Q3V3R1 | Mthfd1l | 0.570 | 0.839 | 0.201 | 0.000 | -0.215 | 0.497 | -0.751 | -0.501 |
| Q91VW3 | Sh3bgrl3 | 0.003 | 0.033 | -0.165 | 0.000 | 0.130 | -1.470 | -2.560 | -2.250 |
| E9Q555 | Rnf213 | 0.959 | 1.000 | 0.447 | 0.000 | -0.205 | 0.553 | -0.424 | 0.057 |
| Q8K411 | Pitrm1 | 0.003 | 0.030 | 0.219 | 0.000 | -0.224 | 1.320 | 0.923 | 1.190 |
| Q9D1L0 | Chchd2 | 0.596 | 0.869 | -0.345 | 0.000 | 0.075 | 0.112 | -1.140 | 0.029 |
| Q9QYR9 | Acot2 | 0.001 | 0.018 | -0.148 | 0.041 | 0.000 | 0.854 | 0.676 | 0.934 |
| P16858 | Gapdh | 0.043 | 0.142 | -0.051 | 0.026 | 0.000 | -0.153 | -0.557 | -0.357 |
| P61222 | Abce1 | 0.300 | 0.529 | 0.517 | -0.034 | 0.000 | 0.313 | -0.505 | -0.467 |
| P61161 | Actr2 | 0.282 | 0.505 | 0.071 | -0.134 | 0.000 | 0.212 | -0.043 | 0.126 |
| Q8VCH8 | Ubxn4 | 0.013 | 0.069 | 0.099 | 0.000 | -0.026 | 0.633 | 0.446 | 0.951 |
| O55023 | Impa1 | 0.000 | 0.013 | -0.248 | 0.000 | 0.009 | -1.540 | -1.950 | -1.710 |
| P70248 | Myo1f | 0.137 | 0.302 | 0.249 | -0.171 | 0.000 | -0.054 | -0.811 | -0.449 |
| Q9JHW2 | Nit2 | 0.309 | 0.542 | -0.141 | 0.431 | 0.000 | 0.296 | 0.245 | 0.362 |
| Q9JHW4 | Eefsec | 0.087 | 0.224 | 0.249 | -0.072 | 0.000 | -0.272 | -1.360 | -0.515 |
| O88736 | Hsd17b7 | 0.000 | 0.004 | 0.000 | 0.367 | -0.023 | 4.050 | 3.850 | 4.130 |
| P57759 | Erp29 | 0.022 | 0.094 | 0.000 | 0.074 | -0.443 | 0.673 | 0.369 | 0.667 |
| O35654 | Pold2 | 0.012 | 0.068 | -0.024 | 0.000 | 0.141 | -0.399 | -0.668 | -0.328 |
| Q62419 | Sh3gl1 | 0.450 | 0.708 | 0.000 | -0.110 | 0.019 | 0.287 | -0.209 | 0.241 |
| P61082 | Ube2m | 0.204 | 0.400 | 0.058 | -0.064 | 0.000 | 0.007 | -0.882 | -0.322 |
| Q8K1E0 | Stx5 | 0.141 | 0.307 | 0.053 | -0.343 | 0.000 | 0.073 | 0.123 | 0.426 |
| Q60715 | P4ha1 | 0.000 | 0.008 | 0.000 | 0.805 | -0.211 | 6.380 | 6.030 | 6.340 |
| Q9D8Y0 | Efhd2 | 0.003 | 0.033 | 0.045 | 0.000 | -0.014 | -0.957 | -1.570 | -1.060 |
| Q9R0P5 | Dstn | 0.888 | 1.000 | 0.713 | 0.000 | -1.130 | 0.571 | -0.381 | -0.332 |
| Q9QYC7 | Ggcx | 0.000 | 0.003 | -0.235 | 0.000 | 0.166 | 5.000 | 4.810 | 5.100 |
| Q6ZWX6 | Eif2s1 | 0.438 | 0.695 | 0.263 | 0.000 | -0.164 | 0.248 | -0.476 | -0.325 |
| Q9DB20 | Atp5o | 0.001 | 0.019 | 0.000 | 0.129 | -0.067 | 0.904 | 0.673 | 0.901 |
| Q8BR65 | Suds3 | 0.139 | 0.304 | 0.057 | 0.000 | -0.089 | -0.026 | -0.482 | -0.292 |
| Q9QZ88 | Vps29 | 0.132 | 0.296 | 0.190 | -0.256 | 0.000 | -0.377 | -1.590 | -0.445 |
| O08709 | Prdx6 | 0.003 | 0.031 | 0.085 | 0.000 | -0.156 | -1.090 | -1.810 | -1.670 |
| Q9CQ22 | Lamtor1 | 0.492 | 0.756 | 0.000 | -0.355 | 0.051 | 0.095 | -0.158 | 0.109 |

| | | | | | | | | | |
|---|---|---|---|---|---|---|---|---|---|
| Q8BIJ6 | Iars2 | 0.008 | 0.053 | 0.000 | 0.073 | -0.104 | 0.819 | 0.408 | 0.655 |
| Q8CIN4 | Pak2 | 0.001 | 0.022 | -0.049 | 0.000 | 0.006 | -1.000 | -1.520 | -1.160 |
| Q9CZW4 | Acsl3 | 0.001 | 0.022 | 0.554 | -0.639 | 0.000 | 3.110 | 2.640 | 3.120 |
| Q61739 | Itga6 | 0.274 | 0.495 | 0.000 | 0.607 | -0.319 | -0.246 | -0.143 | -0.388 |
| Q8K2Z4 | Ncapd2 | 0.437 | 0.694 | 0.059 | -0.356 | 0.000 | 0.217 | -0.607 | -0.761 |
| Q91YQ5 | Rpn1 | 0.001 | 0.018 | 0.000 | 0.016 | -0.293 | 1.400 | 1.040 | 1.180 |
| Q9DC61 | Pmpca | 0.012 | 0.068 | 0.000 | 0.351 | -0.158 | 1.180 | 0.802 | 0.788 |
| Q3UJB9 | Edc4 | 0.867 | 1.000 | 0.128 | -0.076 | 0.000 | 0.309 | -0.341 | -0.021 |
| P18242 | Ctsd | 0.694 | 0.969 | 0.717 | 0.000 | -0.835 | 0.208 | -0.516 | -0.451 |
| Q99L13 | Hibadh | 0.000 | 0.014 | 0.000 | 0.215 | -0.163 | 1.320 | 1.240 | 1.450 |
| Q03265 | Atp5a1 | 0.002 | 0.025 | 0.000 | 0.050 | -0.113 | 0.804 | 0.540 | 0.809 |
| Q9QXX4 | Slc25a13 | 0.004 | 0.038 | -0.014 | 0.047 | 0.000 | 0.518 | 0.393 | 0.699 |
| O54879 | Hmgb3 | 0.121 | 0.278 | 0.188 | -0.126 | 0.000 | -0.298 | -0.889 | -0.166 |
| Q9QY06 | Myo9b | 0.117 | 0.273 | 0.000 | -0.054 | 0.168 | 0.018 | -0.602 | -0.651 |
| Q9Z1F9 | Uba2 | 0.005 | 0.042 | 0.000 | -0.064 | 0.102 | -1.020 | -1.890 | -1.320 |
| P62855 | Rps26 | 0.563 | 0.832 | 0.024 | -0.672 | 0.000 | 0.355 | -0.968 | -0.977 |
| P63005 | Pafah1b1 | 0.015 | 0.075 | 0.000 | 0.142 | -0.104 | -0.432 | -0.897 | -0.558 |
| Q6PGG2 | Gmip | 0.008 | 0.055 | 0.062 | 0.000 | -0.212 | -1.290 | -1.810 | -0.933 |
| O35855 | Bcat2 | 0.331 | 0.569 | 0.049 | 0.000 | -0.484 | 0.516 | -0.134 | 0.039 |
| P29351 | Ptpn6 | 0.095 | 0.238 | 0.228 | -0.143 | 0.000 | -0.147 | -0.756 | -0.370 |
| Q8BPG6 | Sumf2 | 0.000 | 0.013 | -0.094 | 0.054 | 0.000 | 0.610 | 0.632 | 0.744 |
| Q9CQC7 | Ndufb4 | 0.005 | 0.039 | 0.000 | 0.250 | -0.034 | 0.719 | 0.760 | 0.569 |
| Q99KH8 | Stk24 | 0.035 | 0.123 | -0.121 | 0.133 | 0.000 | -0.235 | -0.505 | -0.703 |
| Q9Z2N8 | Actl6a | 0.006 | 0.047 | -0.016 | 0.057 | 0.000 | -0.380 | -0.667 | -0.402 |
| Q91ZV0 | Mia2 | 0.257 | 0.472 | -0.554 | 0.166 | 0.000 | 0.561 | -0.135 | 0.378 |
| Q8CGK3 | Lonp1 | 0.159 | 0.335 | 0.242 | 0.000 | -0.158 | 1.280 | 0.204 | 0.415 |
| Q8C4J7 | Tbl3 | 0.049 | 0.154 | 0.189 | 0.000 | -0.100 | -0.174 | -0.628 | -0.759 |
| Q8VEM8 | Slc25a3 | 0.001 | 0.019 | 0.000 | 0.026 | -0.130 | 1.170 | 0.803 | 0.950 |
| P60762 | Morf4l1 | 0.009 | 0.057 | -0.318 | 0.087 | 0.000 | -0.962 | -0.761 | -0.658 |
| Q9D8U8 | Snx5 | 0.176 | 0.360 | 0.255 | 0.000 | -0.005 | 0.060 | -0.518 | -0.216 |
| Q9Z2U1 | Psma5 | 0.001 | 0.018 | -0.148 | 0.000 | 0.027 | -0.997 | -1.330 | -1.400 |
| P27641 | Xrcc5 | 0.003 | 0.030 | -0.102 | 0.000 | 0.052 | -1.050 | -1.770 | -1.530 |
| Q80UU9 | Pgrmc2 | 0.036 | 0.125 | 0.101 | -1.420 | 0.000 | 1.290 | 0.890 | 1.240 |
| Q9D051 | Pdhb | 0.332 | 0.570 | 0.000 | 0.004 | -0.255 | 0.792 | -0.134 | 0.064 |
| P47758 | Srprb | 0.009 | 0.057 | -0.191 | 0.223 | 0.000 | 0.773 | 0.554 | 0.890 |
| Q9CYR0 | Ssbp1 | 0.351 | 0.592 | 0.000 | 0.078 | 0.000 | 0.385 | -0.069 | 0.185 |
| Q91ZX7 | Lrp1 | 0.002 | 0.025 | 0.234 | 0.000 | -1.060 | 2.740 | 2.500 | 2.710 |
| Q9D1P4 | Chordc1 | 0.031 | 0.116 | 0.000 | -0.375 | 0.231 | -0.843 | -1.750 | -0.898 |
| Q8VCB1 | Ndc1 | 0.083 | 0.216 | -0.054 | 0.125 | 0.000 | 0.620 | 0.106 | 0.452 |
| P62889 | Rpl30 | 0.194 | 0.386 | 0.236 | -0.137 | 0.000 | 0.653 | 0.049 | 0.359 |
| Q64521 | Gpd2 | 0.010 | 0.061 | 0.000 | 0.125 | -0.395 | 0.903 | 0.600 | 0.693 |
| O70152 | Dpm1 | 0.009 | 0.058 | -0.055 | 0.148 | 0.000 | 0.621 | 0.542 | 0.972 |
| Q64727 | Vcl | 0.889 | 1.000 | 0.423 | 0.000 | -0.698 | 0.350 | -0.352 | -0.100 |
| Q61792 | Lasp1 | 0.591 | 0.864 | -0.323 | 0.046 | 0.000 | 0.162 | -0.267 | 0.147 |
| O88532 | Zfr | 0.756 | 1.000 | -0.374 | 0.000 | 0.242 | -0.033 | -0.343 | 0.030 |
| Q64012 | Raly | 0.019 | 0.086 | 0.000 | 0.086 | -0.035 | -0.193 | -0.356 | -0.175 |
| Q9EPE9 | Atp13a1 | 0.000 | 0.013 | 0.082 | 0.000 | -0.043 | 0.982 | 1.020 | 1.240 |
| Q08093 | Cnn2 | 0.711 | 0.987 | -0.069 | 0.221 | 0.000 | 0.526 | -0.210 | 0.112 |
| Q9Z0X1 | Aifm1 | 0.917 | 1.000 | 0.000 | 0.036 | -0.025 | 0.160 | -0.134 | 0.015 |
| Q8K297 | Colgalt1 | 0.000 | 0.013 | 0.020 | 0.000 | -0.798 | 3.140 | 2.810 | 2.900 |
| P27601 | Gna13 | 0.966 | 1.000 | -0.229 | 0.139 | 0.000 | 0.035 | -0.168 | 0.062 |
| P68372 | Tubb4b | 0.036 | 0.125 | 0.000 | 0.073 | -0.162 | -0.272 | -0.840 | -0.694 |
| Q01853 | Vcp | 0.002 | 0.025 | 0.000 | 0.056 | -0.134 | -0.465 | -0.616 | -0.590 |
| Q8K1R3 | Pnpt1 | 0.333 | 0.571 | 0.128 | 0.000 | -0.456 | 1.050 | 0.609 | -0.428 |
| Q6NS46 | Pdcd11 | 0.115 | 0.270 | 0.288 | 0.000 | -0.052 | 0.029 | -0.613 | -0.698 |
| Q91V92 | Acly | 0.836 | 1.000 | 0.521 | 0.000 | -0.217 | 0.528 | -0.126 | 0.096 |
| Q61687 | Atrx | 0.020 | 0.089 | -0.063 | 0.030 | 0.000 | -0.511 | -1.080 | -0.541 |
| O35887 | Calu | 0.000 | 0.010 | 0.000 | 0.381 | -0.028 | 2.430 | 2.320 | 2.650 |
| Q8VBV7 | Cops8 | 0.011 | 0.063 | -0.099 | 0.297 | 0.000 | -0.536 | -0.993 | -1.100 |

| | | | | | | | | | |
|---|---|---|---|---|---|---|---|---|---|
| Q9D892 | Itpa | 0.015 | 0.075 | 0.000 | 0.571 | -0.294 | -0.901 | -1.130 | -1.500 |
| P32067 | Ssb | 0.024 | 0.099 | 0.085 | 0.000 | -0.109 | -0.360 | -1.020 | -0.784 |
| Q8BI72 | Cdkn2aip | 0.009 | 0.058 | -0.178 | 0.025 | 0.000 | -0.523 | -0.712 | -0.421 |
| P97372 | Psme2 | 0.004 | 0.035 | -0.129 | 0.542 | 0.000 | -1.040 | -1.290 | -1.300 |
| Q9WUM5 | Suclg1 | 0.579 | 0.850 | 0.279 | 0.000 | -0.064 | 1.160 | -0.031 | -0.140 |
| P97452 | Bop1 | 0.146 | 0.315 | 0.031 | 0.000 | -0.439 | -0.274 | -0.489 | -0.693 |
| Q6WVG3 | Kctd12 | 0.656 | 0.932 | 0.315 | 0.000 | -0.034 | 0.280 | -0.225 | -0.040 |
| Q9CXW3 | Cacybp | 0.796 | 1.000 | 0.000 | 0.321 | -0.259 | 0.140 | -0.448 | 0.154 |
| Q8C7V8 | Ccdc134 | 0.086 | 0.222 | 0.343 | -0.195 | 0.000 | 1.030 | 0.218 | 1.050 |
| Q9EPU0 | Upf1 | 0.254 | 0.467 | 0.070 | 0.000 | -0.442 | -0.174 | -0.567 | -0.419 |
| Q8K2B3 | Sdha | 0.017 | 0.081 | 0.000 | 0.002 | -0.204 | 0.722 | 0.297 | 0.454 |
| P67778 | Phb | 0.003 | 0.033 | 0.000 | 0.017 | -0.309 | 0.843 | 0.579 | 0.735 |
| Q6NZJ6 | Eif4g1 | 0.192 | 0.384 | 0.001 | -0.411 | 0.000 | 0.581 | -0.101 | 0.237 |
| P35235 | Ptpn11 | 0.002 | 0.029 | -0.286 | 0.010 | 0.000 | -0.906 | -1.190 | -0.916 |
| Q922Q4 | Pycr2 | 0.016 | 0.078 | 0.102 | 0.000 | -0.106 | 1.190 | 0.520 | 0.764 |
| P47753 | Capza1 | 0.230 | 0.435 | -0.227 | 0.804 | 0.000 | -0.433 | -0.018 | -0.420 |
| Q61102 | Abcb7 | 0.585 | 0.857 | -0.410 | 0.169 | 0.000 | 0.145 | -0.268 | 0.316 |
| P47802 | Mtx1 | 0.295 | 0.522 | 0.000 | 0.014 | -0.282 | 0.283 | -0.076 | 0.042 |
| P03958 | Ada | 0.014 | 0.072 | 0.754 | -0.343 | 0.000 | -1.270 | -2.090 | -1.410 |
| P63321 | Rala | 0.001 | 0.019 | 0.000 | 0.267 | -0.010 | 1.050 | 1.270 | 1.390 |
| P27808 | Mgat1 | 0.011 | 0.063 | 0.341 | -0.109 | 0.000 | 1.110 | 0.673 | 1.070 |
| Q91V08 | Clec2d | 0.595 | 0.868 | 0.000 | 4.180 | -0.694 | -1.230 | 2.370 | -0.961 |
| P21279 | Gnaq | 0.001 | 0.019 | 0.000 | 0.266 | -0.175 | 1.290 | 1.150 | 1.370 |
| Q3TWW8 | Srsf6 | 0.078 | 0.209 | 0.100 | 0.000 | -0.133 | -0.141 | -1.130 | -1.060 |
| Q80WQ2 | Vac14 | 0.093 | 0.234 | 0.126 | 0.000 | -0.005 | -0.024 | -0.634 | -0.440 |
| P19157 | Gstp1 | 0.004 | 0.036 | -0.138 | 0.000 | 0.106 | -1.050 | -1.380 | -0.809 |
| Q99JF8 | Psip1 | 0.090 | 0.230 | -0.095 | 0.000 | 0.152 | -0.474 | -0.546 | -0.054 |
| Q9EPU4 | Cpsf1 | 0.146 | 0.315 | 0.000 | 0.097 | -0.063 | -0.601 | -0.441 | 0.028 |
| P61028 | Rab8b | 0.484 | 0.747 | 0.000 | 0.106 | -1.460 | -0.015 | -0.228 | 0.075 |
| Q9EQH2 | Erap1 | 0.628 | 0.904 | -0.026 | 0.272 | 0.000 | 0.049 | -0.104 | 0.119 |
| P63276 | Rps17 | 0.251 | 0.463 | 0.554 | -0.143 | 0.000 | 0.393 | 0.274 | 1.260 |
| P08249 | Mdh2 | 0.023 | 0.096 | -0.067 | 0.250 | 0.000 | 0.700 | 0.373 | 0.737 |
| Q8C0E2 | Vps26b | 0.098 | 0.242 | 0.000 | -0.173 | 0.155 | -0.331 | -1.450 | -0.538 |
| P06537 | Nr3c1 | 0.043 | 0.141 | -0.291 | 0.000 | 0.306 | -0.491 | -0.996 | -0.556 |
| P84096 | Rhog | 0.210 | 0.406 | 0.099 | -0.086 | 0.000 | -0.027 | -0.680 | -0.190 |
| Q8K273 | Mmgt1 | 0.040 | 0.135 | 0.123 | -0.229 | 0.000 | 1.120 | 0.385 | 1.530 |
| Q6P5D8 | Smchd1 | 0.071 | 0.194 | -0.012 | 0.080 | 0.000 | -0.246 | -0.640 | -0.156 |
| Q99KC8 | Vwa5a | 0.041 | 0.136 | -0.167 | 0.112 | 0.000 | 0.212 | 0.207 | 0.290 |
| Q6A0A9 | FAM120A | 0.722 | 1.000 | -0.005 | 0.168 | 0.000 | 0.747 | -0.800 | -0.310 |
| P26231 | Ctnna1 | 0.001 | 0.018 | 0.052 | -0.640 | 0.000 | 3.370 | 2.460 | 2.860 |
| Q6ZPE2 | Sbf1 | 0.006 | 0.044 | 0.052 | -0.183 | 0.000 | -0.773 | -1.370 | -1.020 |
| Q9Z1K5 | Arih1 | 0.021 | 0.092 | 0.126 | -0.138 | 0.000 | -0.818 | -2.130 | -1.350 |
| Q3UUQ7 | Pgap1 | 0.013 | 0.072 | 0.463 | -0.528 | 0.000 | 1.170 | 1.170 | 1.220 |
| P42125 | Eci1 | 0.192 | 0.384 | 0.000 | 0.429 | -0.075 | 0.361 | 0.416 | 0.326 |
| Q8C9B9 | Dido1 | 0.033 | 0.120 | -0.193 | 0.041 | 0.000 | 0.473 | 0.154 | 0.515 |
| Q80U72 | Scrib | 0.649 | 0.926 | -0.386 | 0.062 | 0.000 | 1.500 | -0.647 | -0.191 |
| O35143 | Atpif1 | 0.013 | 0.069 | 0.259 | 0.000 | -0.321 | 1.590 | 1.150 | 0.813 |
| Q8K310 | Matr3 | 0.033 | 0.119 | -0.160 | 0.000 | 0.057 | -0.247 | -0.430 | -0.265 |
| Q8BXZ1 | Tmx3 | 0.000 | 0.008 | -0.015 | 0.000 | 0.010 | 0.683 | 0.570 | 0.683 |
| Q9JJT0 | Rcl1 | 0.006 | 0.044 | -0.094 | 0.000 | 0.033 | -0.925 | -1.080 | -1.640 |
| P35278 | Rab5c | 0.471 | 0.730 | -0.070 | 0.126 | 0.000 | 0.187 | -0.097 | 0.261 |
| Q91VR5 | Ddx1 | 0.525 | 0.791 | 0.054 | 0.000 | -0.008 | 0.345 | -0.164 | 0.182 |
| P84084 | Arf5 | 0.092 | 0.233 | 0.000 | 0.100 | -0.067 | -0.116 | -1.250 | -0.806 |
| P14685 | Psmd3 | 0.048 | 0.152 | 0.000 | -0.102 | 0.036 | -0.240 | -0.752 | -0.399 |
| P80315 | Cct4 | 0.030 | 0.114 | -0.089 | 0.028 | 0.000 | -0.246 | -0.738 | -0.523 |
| Q7TPD0 | Ints3 | 0.037 | 0.128 | 0.000 | -0.118 | 0.448 | -0.365 | -1.040 | -0.669 |
| O88531 | Ppt1 | 0.102 | 0.248 | 0.288 | 0.000 | -0.559 | -0.350 | -1.070 | -1.500 |
| Q921M3 | Sf3b3 | 0.003 | 0.031 | -0.017 | 0.117 | 0.000 | -0.436 | -0.637 | -0.422 |
| Q8VH51 | Rbm39 | 0.044 | 0.143 | 0.079 | 0.000 | -0.126 | -0.211 | -0.764 | -0.586 |

| | | | | | | | | | |
|---|---|---|---|---|---|---|---|---|---|
| Q8BHN3 | Ganab | 0.006 | 0.047 | 0.180 | 0.000 | -0.140 | 0.719 | 0.487 | 0.700 |
| Q9CZW5 | Tomm70 | 0.062 | 0.179 | 0.000 | 0.008 | -0.151 | 0.239 | 0.035 | 0.226 |
| O35972 | Mrpl23 | 0.003 | 0.033 | 0.000 | 0.107 | -0.240 | 0.886 | 0.665 | 0.669 |
| P62315 | Snrpd1 | 0.645 | 0.922 | 0.000 | -1.410 | 0.030 | -0.557 | -0.619 | 0.778 |
| O35350 | Capn1 | 0.018 | 0.084 | -0.014 | 0.000 | 0.021 | -0.236 | -0.579 | -0.351 |
| Q60737 | Csnk2a1 | 0.841 | 1.000 | 0.000 | 0.031 | -0.134 | -0.077 | -0.483 | 0.308 |
| P97376 | Frg1 | 0.010 | 0.059 | 0.020 | -0.044 | 0.000 | -0.541 | -0.851 | -0.422 |
| P68037 | Ube2l3 | 0.133 | 0.297 | 0.000 | 0.339 | -0.029 | -0.282 | -1.350 | -0.224 |
| P54103 | Dnajc2 | 0.094 | 0.236 | 0.144 | 0.000 | 0.000 | -0.125 | -0.980 | -0.423 |
| O08915 | Aip | 0.025 | 0.102 | 0.016 | -0.067 | 0.000 | -0.672 | -1.950 | -1.290 |
| O35841 | Api5 | 0.032 | 0.116 | -0.044 | 0.000 | 0.040 | -0.768 | -0.726 | -0.225 |
| Q8CHP6 | Phc3 | 0.947 | 1.000 | -0.400 | 0.000 | 0.610 | 0.040 | 0.081 | 0.027 |
| Q8K284 | Gtf3c1 | 0.105 | 0.252 | 0.000 | 0.163 | -0.024 | -0.119 | -1.060 | -0.453 |
| Q09014 | Ncf1 | 0.407 | 0.658 | 1.740 | -0.022 | 0.000 | 1.230 | -1.690 | -0.706 |
| P81269 | Atf1 | 0.311 | 0.544 | 0.000 | -2.120 | 0.502 | 0.421 | -0.113 | 1.130 |
| O35343 | Kpna4 | 0.217 | 0.415 | 0.000 | 0.365 | -0.431 | -1.080 | -0.695 | -0.005 |
| Q9CSU0 | Rprd1b | 0.000 | 0.009 | -0.009 | 0.058 | 0.000 | -0.946 | -1.160 | -1.000 |
| Q3UQ44 | Iqgap2 | 0.285 | 0.509 | 0.213 | 0.000 | -0.073 | 0.235 | -0.465 | -0.837 |
| Q6P5F9 | Xpo1 | 0.204 | 0.400 | 0.004 | 0.000 | -0.804 | -0.562 | -0.972 | -0.620 |
| Q9D0D3 | Mtpap | 0.039 | 0.134 | -0.042 | 0.000 | 0.411 | 0.553 | 0.544 | 0.831 |
| P51859 | Hdgf | 0.004 | 0.034 | -0.010 | 0.000 | 0.062 | -0.903 | -1.530 | -1.040 |
| P11942 | Cd3g | 0.167 | 0.349 | -0.367 | 0.000 | 0.062 | -0.670 | -0.495 | -0.158 |
| Q91VE6 | Nifk | 0.244 | 0.454 | 0.009 | 0.000 | -0.055 | 0.059 | -0.834 | -0.332 |
| O88543 | Cops3 | 0.037 | 0.128 | -0.261 | 0.000 | 0.061 | -0.532 | -1.550 | -0.983 |
| Q9D6S7 | Mrrf | 0.086 | 0.223 | 0.000 | 0.433 | -0.651 | 0.785 | 0.917 | 0.434 |
| Q9CXT8 | Pmpcb | 0.017 | 0.082 | 0.000 | 0.156 | -0.441 | 1.050 | 0.643 | 0.643 |
| O09172 | Gclm | 0.514 | 0.778 | 0.497 | 0.000 | -0.914 | 0.147 | -0.892 | -0.812 |
| O70370 | Ctss | 0.016 | 0.078 | 0.174 | 0.000 | -0.123 | -0.289 | -0.491 | -0.485 |
| Q9ES52 | Inpp5d | 0.069 | 0.191 | -0.054 | 0.183 | 0.000 | -0.248 | -0.967 | -0.380 |
| Q8CHT0 | Aldh4a1 | 0.875 | 1.000 | 0.335 | -0.067 | 0.000 | 1.090 | -0.715 | -0.397 |
| O35326 | Srsf5 | 0.051 | 0.157 | 0.057 | 0.000 | -0.537 | -0.598 | -1.720 | -1.290 |
| P61255 | Rpl26 | 0.969 | 1.000 | 0.146 | 0.000 | -0.044 | 0.302 | -0.198 | -0.021 |
| Q9WV54 | Asah1 | 0.436 | 0.693 | 0.241 | 0.000 | -0.799 | -0.033 | -0.720 | -0.851 |
| Q5SWD9 | Tsr1 | 0.007 | 0.048 | 0.044 | -0.144 | 0.000 | -0.590 | -0.766 | -0.441 |
| Q9Z0N1 | Eif2s3x | 0.715 | 0.992 | 0.118 | 0.000 | -0.435 | 0.405 | -0.210 | -0.202 |
| Q9JIG8 | Praf2 | 0.001 | 0.022 | -0.152 | 0.388 | 0.000 | 1.550 | 1.310 | 1.510 |
| P17742 | Ppia | 0.001 | 0.020 | -0.069 | 0.072 | 0.000 | -0.744 | -1.100 | -0.888 |
| Q9D1N9 | Mrpl21 | 0.001 | 0.019 | 0.000 | 0.089 | -0.218 | 0.892 | 0.811 | 1.010 |
| P21460 | Cst3 | 0.344 | 0.583 | 0.555 | 0.000 | -0.002 | 0.303 | -0.365 | -0.291 |
| Q8QZT1 | Acat1 | 0.050 | 0.156 | 0.000 | 0.181 | -0.006 | 0.765 | 0.228 | 0.592 |
| P60843 | Eif4a1 | 0.768 | 1.000 | 0.365 | 0.000 | -0.115 | 0.496 | -0.360 | -0.167 |
| Q9DCS3 | Mecr | 0.121 | 0.279 | 0.000 | 0.207 | -0.206 | 1.210 | 0.269 | 0.392 |
| Q9QUI0 | Rhoa | 0.092 | 0.233 | 0.000 | -0.019 | 0.008 | -0.050 | -0.555 | -0.397 |
| Q7TPH6 | Mycbp2 | 0.256 | 0.470 | 0.044 | -0.016 | 0.000 | 0.048 | -0.906 | -0.239 |
| Q8C0C0 | Zhx2 | 0.138 | 0.303 | -0.142 | 0.000 | 0.004 | -0.455 | -0.390 | -0.046 |
| B2RXR6 | Ankrd44 | 0.004 | 0.036 | 0.067 | -0.178 | 0.000 | -1.230 | -2.070 | -1.460 |
| O89053 | Coro1a | 0.001 | 0.021 | -0.088 | 0.000 | 0.009 | -0.623 | -0.917 | -0.740 |
| Q9Z0E6 | Gbp2 | 0.096 | 0.238 | -0.595 | 0.652 | 0.000 | -0.971 | -1.400 | -0.481 |
| Q8VCM8 | Ncln | 0.000 | 0.013 | 0.163 | 0.000 | -0.176 | 2.230 | 1.750 | 2.080 |
| P39054 | Dnm2 | 0.326 | 0.562 | 0.075 | -0.001 | 0.000 | 0.448 | -0.098 | 0.271 |
| O55106 | Strn | 0.053 | 0.162 | -0.090 | 0.271 | 0.000 | -0.484 | -0.533 | -0.141 |
| Q9QX47 | Son | 0.342 | 0.582 | -0.273 | 0.117 | 0.000 | -0.128 | -0.274 | -0.155 |
| Q9D883 | U2af1 | 0.235 | 0.442 | 0.393 | -0.112 | 0.000 | 0.274 | -0.934 | -0.764 |
| Q7TMR0 | Prcp | 0.537 | 0.805 | 0.534 | 0.000 | -0.186 | 0.295 | -0.429 | -0.127 |
| Q8JZQ9 | Eif3b | 0.260 | 0.476 | 0.275 | -0.073 | 0.000 | 0.153 | -0.697 | -0.305 |
| Q3V0C5 | Usp48 | 0.002 | 0.023 | -0.188 | 0.240 | 0.000 | -1.040 | -1.390 | -1.230 |
| Q922B2 | Dars | 0.310 | 0.543 | 0.005 | 0.000 | -0.176 | -0.007 | -0.316 | -0.229 |
| Q3THK3 | Gtf2f1 | 0.002 | 0.024 | 0.000 | 0.200 | -0.068 | -1.120 | -1.740 | -1.450 |
| Q8CFX1 | H6pd | 0.003 | 0.031 | 0.000 | 0.021 | -0.537 | 1.370 | 1.020 | 1.180 |

| | | | | | | | | | |
|---|---|---|---|---|---|---|---|---|---|
| Q8K1I7 | Wipf1 | 0.146 | 0.315 | -0.001 | 0.000 | 0.402 | -0.311 | -0.419 | 0.062 |
| Q4VA53 | Pds5b | 0.031 | 0.116 | -0.082 | 0.000 | 0.035 | -0.347 | -0.548 | -0.198 |
| Q7TPV4 | Mybbp1a | 0.078 | 0.208 | 0.135 | 0.000 | -0.117 | -0.129 | -0.809 | -0.532 |
| Q3ULD5 | Mccc2 | 0.004 | 0.036 | 0.000 | 0.315 | -0.197 | 1.080 | 0.893 | 0.981 |
| Q8BJS4 | Sun2 | 0.725 | 1.000 | 0.000 | 0.079 | -0.089 | 0.028 | -0.229 | 0.073 |
| Q64324 | Stxbp2 | 0.030 | 0.112 | 0.000 | 0.170 | -0.102 | -0.258 | -0.789 | -0.715 |
| Q9CWK8 | Snx2 | 0.007 | 0.050 | 0.074 | 0.000 | -0.016 | -0.329 | -0.673 | -0.635 |
| Q9DCC8 | Tomm20 | 0.641 | 0.917 | -0.091 | 0.350 | 0.000 | 0.068 | 0.058 | 0.398 |
| O88342 | Wdr1 | 0.001 | 0.019 | -0.112 | 0.000 | 0.084 | -0.978 | -1.390 | -1.190 |
| P55096 | Abcd3 | 0.092 | 0.232 | 0.042 | 0.000 | -0.929 | 1.360 | 1.060 | 0.053 |
| Q80W54 | Zmpste24 | 0.080 | 0.211 | 0.027 | 0.000 | -0.220 | 0.715 | 0.060 | 0.482 |
| P32020 | Scp2 | 0.103 | 0.248 | 0.172 | 0.000 | -0.126 | 0.611 | 0.091 | 0.465 |
| Q8BUH8 | Senp7 | 0.036 | 0.125 | -0.092 | 0.000 | 0.105 | -0.578 | -0.970 | -0.310 |
| Q07797 | Lgals3bp | 0.056 | 0.168 | 0.660 | 0.000 | -0.240 | 1.040 | 0.809 | 0.805 |
| Q9QYB1 | Clic4 | 0.007 | 0.051 | 0.074 | 0.000 | -0.173 | 1.240 | 0.654 | 0.833 |
| Q9CR16 | Ppid | 0.050 | 0.155 | 0.063 | 0.000 | -0.400 | -0.464 | -0.946 | -0.621 |
| P19096 | Fasn | 0.558 | 0.826 | 0.157 | 0.000 | -0.066 | 0.282 | -0.464 | -0.161 |
| Q8BTS4 | Nup54 | 0.174 | 0.357 | -0.078 | 0.000 | 0.036 | -0.224 | -0.263 | 0.002 |
| P61750 | Arf4 | 0.057 | 0.168 | 0.156 | -0.200 | 0.000 | 1.220 | 0.315 | 0.670 |
| P43247 | Msh2 | 0.040 | 0.135 | 0.112 | 0.000 | -0.005 | -0.303 | -0.900 | -0.393 |
| Q9CXF4 | Tbc1d15 | 0.442 | 0.698 | 0.404 | 0.000 | -0.414 | 0.186 | -0.566 | -0.489 |
| P47856 | Gfpt1 | 0.260 | 0.476 | 0.041 | -0.007 | 0.000 | 0.080 | -0.490 | -0.204 |
| Q640M1 | Utp14a | 0.054 | 0.163 | 0.000 | -0.037 | 0.384 | -0.252 | -0.903 | -0.422 |
| Q9QZM0 | Ubqln2 | 0.002 | 0.025 | 0.000 | -0.073 | 0.286 | -0.713 | -0.834 | -0.761 |
| Q60739 | Bag1 | 0.001 | 0.015 | -0.089 | 0.000 | 0.076 | -1.060 | -1.260 | -0.917 |
| Q8BH95 | Echs1 | 0.035 | 0.124 | 0.000 | 0.240 | -0.097 | 0.915 | 0.414 | 0.524 |
| Q9JLV6 | Pnkp | 0.033 | 0.121 | 0.186 | 0.000 | -0.012 | -0.137 | -0.517 | -0.435 |
| P56391 | Cox6b1 | 0.015 | 0.076 | 0.030 | 0.000 | -0.083 | 0.406 | 0.174 | 0.409 |
| P48758 | Cbr1 | 0.045 | 0.146 | 0.000 | -0.362 | 0.229 | -0.750 | -2.180 | -1.160 |
| Q60668 | Hnrnpd | 0.029 | 0.111 | -0.167 | 0.000 | 0.014 | -0.668 | -0.810 | -0.301 |
| Q99LF4 | Rtcb | 0.551 | 0.820 | 0.000 | 0.036 | -0.061 | 0.119 | -0.108 | 0.122 |
| Q91V64 | Isoc1 | 0.027 | 0.106 | 0.263 | 0.000 | -0.176 | -0.967 | -1.210 | -2.380 |
| Q9D3E6 | Stag1 | 0.097 | 0.241 | -0.110 | 0.000 | 0.040 | -0.192 | -0.391 | -0.105 |
| Q9DBE9 | Ftsj3 | 0.149 | 0.319 | 0.213 | 0.000 | -0.414 | -0.104 | -0.908 | -1.010 |
| Q9JLV5 | Cul3 | 0.171 | 0.353 | 0.033 | -0.575 | 0.000 | -0.300 | -1.190 | -0.676 |
| Q99N96 | Mrpl1 | 0.091 | 0.231 | 0.000 | -1.980 | 0.201 | 0.820 | 0.852 | 1.270 |
| Q8K2T8 | Paf1 | 0.000 | 0.013 | 0.000 | -0.060 | 0.046 | -0.832 | -0.873 | -0.672 |
| O09159 | Man2b1 | 0.658 | 0.934 | 0.557 | -0.062 | 0.000 | 0.520 | 0.127 | 0.181 |
| Q9Z1X4 | Ilf3 | 0.531 | 0.798 | -0.320 | 0.120 | 0.000 | -0.211 | -0.422 | 0.045 |
| P42567 | Eps15 | 0.012 | 0.067 | 0.268 | -0.005 | 0.000 | -0.508 | -1.210 | -0.973 |
| Q9CPR7 | Sike1 | 0.285 | 0.509 | -0.334 | 0.039 | 0.000 | -0.127 | -0.901 | -0.255 |
| Q9D0R2 | Tars | 0.045 | 0.146 | -0.112 | 0.000 | 0.099 | -0.225 | -0.415 | -0.184 |
| Q07076 | Anxa7 | 0.833 | 1.000 | 0.000 | 0.052 | -0.034 | 0.275 | -0.212 | 0.053 |
| Q9QYE6 | Golga5 | 0.161 | 0.339 | 0.042 | -0.319 | 0.000 | 0.172 | 0.003 | 0.235 |
| Q61768 | Kif5b | 0.532 | 0.799 | 0.107 | 0.000 | -0.003 | 0.348 | -0.450 | -0.300 |
| Q99KE1 | Me2 | 0.952 | 1.000 | 0.000 | 0.140 | -0.205 | 0.190 | -0.212 | -0.072 |
| Q60931 | Vdac3 | 0.007 | 0.050 | 0.000 | 0.252 | -0.058 | 1.080 | 0.902 | 1.570 |
| Q63850 | Nup62 | 0.333 | 0.571 | 0.264 | -3.320 | 0.000 | 0.165 | -0.153 | 0.862 |
| O55098 | Stk10 | 0.435 | 0.692 | -0.223 | 0.161 | 0.000 | -0.488 | -0.706 | 0.292 |
| Q8BZH4 | Pogz | 0.153 | 0.326 | -0.120 | 0.000 | 0.033 | -0.410 | -0.449 | -0.010 |
| Q9QYJ0 | Dnaja2 | 0.688 | 0.965 | 0.097 | 0.000 | -0.099 | 0.154 | -0.236 | -0.084 |
| Q8CDN6 | Txnl1 | 0.008 | 0.051 | 0.000 | 0.040 | -0.283 | -0.969 | -1.380 | -0.821 |
| Q8R5C5 | Actr1b | 0.301 | 0.529 | -0.159 | 0.417 | 0.000 | 0.472 | 0.374 | 0.126 |
| P09602 | Hmgn2 | 0.380 | 0.629 | -1.030 | 0.000 | 0.113 | -0.438 | -0.284 | -3.170 |
| P09055 | Itgb1 | 0.001 | 0.019 | 0.000 | 0.434 | 0.000 | 1.880 | 1.520 | 1.680 |
| Q924W5 | Smc6 | 0.171 | 0.353 | 0.165 | 0.000 | -0.119 | -0.122 | -0.954 | -0.241 |
| O08663 | Metap2 | 0.390 | 0.640 | 0.466 | -3.020 | 0.000 | -1.320 | -1.100 | -4.830 |
| P09925 | Surf1 | 0.000 | 0.013 | 0.069 | 0.000 | -0.073 | 0.682 | 0.590 | 0.722 |
| O89100 | Grap2 | 0.049 | 0.154 | -0.257 | 0.110 | 0.000 | -0.559 | -2.360 | -1.760 |

| | | | | | | | | | |
|---|---|---|---|---|---|---|---|---|---|
| P06745 | Gpi | 0.005 | 0.039 | 0.533 | -0.070 | 0.000 | -1.010 | -1.620 | -1.360 |
| Q9EQU5 | Set | 0.009 | 0.055 | 0.000 | 0.489 | -0.112 | -0.929 | -0.885 | -0.687 |
| P58742 | Aaas | 0.006 | 0.044 | 0.000 | 0.068 | -0.024 | 0.523 | 0.402 | 0.731 |
| Q8BMD8 | Slc25a24 | 0.023 | 0.096 | -0.447 | 0.176 | 0.000 | 0.526 | 0.574 | 0.757 |
| O70172 | Pip4k2a | 0.136 | 0.301 | 0.051 | 0.000 | -0.088 | -0.007 | -0.946 | -0.646 |
| P62880 | Gnb2 | 0.106 | 0.254 | 0.181 | -0.405 | 0.000 | 0.353 | 0.177 | 0.720 |
| P11103 | Parp1 | 0.506 | 0.770 | -0.125 | 0.040 | 0.000 | -0.190 | -0.321 | 0.120 |
| Q99LC2 | Cstf1 | 0.099 | 0.242 | -0.452 | 0.024 | 0.000 | -0.702 | -0.620 | -0.334 |
| P11352 | Gpx1 | 0.131 | 0.294 | 0.457 | 0.000 | -0.235 | -0.028 | -0.804 | -0.877 |
| Q04750 | Top1 | 0.149 | 0.319 | 0.118 | -0.057 | 0.000 | -0.028 | -0.407 | -0.160 |
| Q8VE47 | Uba5 | 0.001 | 0.020 | 0.048 | -0.035 | 0.000 | -0.621 | -0.940 | -0.867 |
| Q8K4G5 | Ablim1 | 0.436 | 0.693 | -0.562 | 0.000 | 0.335 | -0.503 | -0.450 | -0.049 |
| P06332 | Cd4 | 0.187 | 0.377 | -0.346 | 0.000 | 0.028 | -0.404 | -0.507 | -0.161 |
| Q8BRF7 | Scfd1 | 0.745 | 1.000 | 0.000 | -0.031 | 0.146 | 0.060 | -0.193 | 0.131 |
| P04627 | Araf | 0.149 | 0.319 | 0.195 | -0.430 | 0.000 | -2.140 | -1.090 | -7.290 |
| Q61545 | Ewsr1 | 0.344 | 0.584 | -0.147 | 0.000 | 0.141 | -0.110 | -0.516 | 0.030 |
| Q9CY64 | Blvra | 0.001 | 0.017 | 0.070 | 0.000 | -0.027 | -1.010 | -1.480 | -1.270 |
| P54116 | Stom | 0.556 | 0.825 | 0.677 | 0.000 | -0.243 | 0.446 | -0.539 | -0.245 |
| Q0P678 | Zc3h18 | 0.031 | 0.114 | 0.000 | -0.085 | 0.167 | -0.421 | -1.190 | -0.663 |
| P59235 | Nup43 | 0.560 | 0.829 | -0.120 | 0.383 | 0.000 | -0.494 | 0.070 | 0.192 |
| Q3UW53 | Fam129a | 0.316 | 0.550 | -0.006 | 0.203 | 0.000 | 0.205 | -0.653 | -0.238 |
| Q02819 | Nucb1 | 0.009 | 0.056 | 0.039 | 0.000 | -0.189 | 0.852 | 0.418 | 0.648 |
| P62334 | Psmc6 | 0.204 | 0.400 | -0.038 | 0.043 | 0.000 | 0.062 | -0.326 | -0.538 |
| P99029 | Prdx5 | 0.996 | 1.000 | 0.234 | 0.000 | -0.266 | 0.051 | -0.201 | 0.121 |
| P56183 | Rrp1 | 0.829 | 1.000 | 1.090 | 0.000 | -2.470 | 0.744 | -3.520 | 0.226 |
| Q9DCR2 | Ap3s1 | 0.247 | 0.458 | 0.237 | -0.394 | 0.000 | 0.805 | 0.129 | 0.105 |
| Q9JLZ8 | Sigirr | 0.827 | 1.000 | -1.170 | 0.459 | 0.000 | -0.526 | -0.206 | -0.321 |
| Q9QUR6 | Prep | 0.027 | 0.106 | 0.102 | -0.262 | 0.000 | -0.737 | -1.200 | -0.530 |
| Q9DBP5 | Cmpk1 | 0.018 | 0.084 | 0.000 | 0.122 | -0.056 | -0.724 | -1.950 | -1.410 |
| Q5SFM8 | Rbm27 | 0.299 | 0.528 | -0.487 | 0.001 | 0.000 | 0.226 | -0.114 | 0.081 |
| Q7JJ13 | Brd2 | 0.032 | 0.117 | -0.064 | 0.006 | 0.000 | -0.586 | -1.290 | -0.544 |
| Q6PHZ2 | Camk2d | 0.078 | 0.209 | -0.861 | 0.391 | 0.000 | -1.070 | -2.170 | -0.983 |
| P29416 | Hexa | 0.886 | 1.000 | 0.576 | -0.195 | 0.000 | 0.462 | -0.293 | 0.066 |
| O70252 | Hmox2 | 0.188 | 0.378 | 0.000 | 0.189 | -0.038 | 0.252 | 0.067 | 0.428 |
| Q9DB73 | Cyb5r1 | 0.000 | 0.008 | 0.000 | 0.094 | -0.277 | 2.440 | 2.150 | 2.440 |
| Q9DBH5 | Lman2 | 0.002 | 0.025 | 0.000 | 0.055 | -0.225 | 0.807 | 0.612 | 0.822 |
| P62242 | Rps8 | 0.850 | 1.000 | 0.000 | 0.181 | -0.355 | 0.070 | -0.089 | -0.056 |
| Q6P069 | Sri | 0.434 | 0.691 | -0.202 | 0.027 | 0.000 | -0.084 | -0.301 | -0.063 |
| Q9CZX8 | Rps19 | 0.508 | 0.772 | 0.243 | -0.007 | 0.000 | 0.331 | -0.433 | -0.184 |
| Q07417 | Acads | 0.730 | 1.000 | 0.089 | 0.000 | -0.125 | 0.435 | -0.537 | -0.263 |
| Q8R1B4 | Eif3c | 0.348 | 0.589 | 0.218 | 0.000 | -0.044 | 0.214 | -0.467 | -0.267 |
| O88379 | Baz1a | 0.138 | 0.303 | 0.035 | 0.000 | -0.139 | -0.281 | -0.452 | -0.062 |
| P14429 | H2-Q7 | 0.206 | 0.403 | -0.383 | 0.114 | 0.000 | -0.472 | -0.369 | -0.198 |
| P70452 | Stx4 | 0.379 | 0.628 | 0.000 | 0.361 | -1.230 | 0.582 | 0.088 | -0.010 |
| P12787 | Cox5a | 0.002 | 0.028 | 0.000 | 0.139 | -0.028 | 0.629 | 0.454 | 0.629 |
| Q8BKE6 | Cyp20a1 | 0.000 | 0.014 | 0.000 | 0.088 | -0.320 | 1.730 | 1.440 | 1.780 |
| P04104 | Krt1 | 0.277 | 0.498 | -0.073 | 0.000 | 0.585 | -0.156 | 1.390 | 1.350 |
| P17439 | Gba | 0.007 | 0.049 | 0.708 | 0.000 | -0.006 | 1.550 | 1.370 | 1.490 |
| Q9D0M3 | Cyc1 | 0.192 | 0.384 | 0.243 | 0.000 | -0.038 | 0.016 | 0.587 | 0.597 |
| Q7TSI3 | Ppp6r1 | 0.005 | 0.040 | 0.000 | -0.040 | 0.250 | -0.738 | -1.240 | -0.832 |
| Q80X85 | Mrps7 | 0.051 | 0.158 | 0.000 | 0.066 | -0.627 | 0.765 | 0.358 | 0.418 |
| Q99ME9 | Gtpbp4 | 0.155 | 0.329 | 0.013 | -0.133 | 0.000 | -0.088 | -0.821 | -0.361 |
| Q00417 | Tcf7 | 0.842 | 1.000 | -1.160 | 0.000 | 0.065 | -0.570 | -0.372 | 0.138 |
| Q922Q8 | Lrrc59 | 0.002 | 0.028 | 0.000 | 0.070 | -0.353 | 0.875 | 0.825 | 0.768 |
| P24547 | Impdh2 | 0.232 | 0.437 | 0.041 | 0.000 | -0.289 | -0.073 | -0.748 | -0.361 |
| P70670 | Naca | 0.788 | 1.000 | 0.000 | 0.099 | -0.468 | -0.113 | -0.429 | -0.013 |
| Q61152 | Ptpn18 | 0.001 | 0.020 | -0.173 | 0.000 | 0.344 | -1.430 | -1.960 | -1.780 |
| Q8BGQ7 | Aars | 0.540 | 0.807 | 0.245 | 0.000 | -0.060 | 0.323 | -0.318 | -0.276 |
| P35486 | Pdha1 | 0.264 | 0.482 | 0.044 | 0.000 | -0.235 | 0.792 | 0.073 | -0.010 |

| | | | | | | | | | |
|---|---|---|---|---|---|---|---|---|---|
| Q9CYZ2 | Tpd52l2 | 0.037 | 0.129 | 0.129 | -0.440 | 0.000 | -0.566 | -0.943 | -1.290 |
| P48024 | Eif1 | 0.101 | 0.247 | 0.057 | 0.000 | -0.056 | -0.341 | -0.676 | -0.087 |
| Q9CXU9 | Eif1b | 0.101 | 0.247 | 0.057 | 0.000 | -0.056 | -0.341 | -0.676 | -0.087 |
| P70168 | Kpnb1 | 0.204 | 0.400 | -0.040 | 0.088 | 0.000 | -0.081 | -0.545 | -0.064 |
| P80318 | Cct3 | 0.041 | 0.138 | 0.007 | 0.000 | -0.029 | -0.227 | -0.751 | -0.412 |
| O88665 | Brd7 | 0.381 | 0.630 | 0.000 | -0.139 | 0.144 | -0.197 | -0.413 | 0.108 |
| Q1HFZ0 | Nsun2 | 0.102 | 0.247 | 0.289 | 0.000 | -0.086 | -0.029 | -1.110 | -0.849 |
| P70333 | Hnrnph2 | 0.656 | 0.932 | -0.681 | 0.027 | 0.000 | -0.074 | -0.027 | -0.211 |
| Q80ZW2 | Them6 | 0.880 | 1.000 | 0.000 | 2.290 | -4.050 | -2.070 | 0.845 | 0.459 |
| Q9D1D4 | Tmed10 | 0.017 | 0.081 | 0.000 | 0.326 | -0.089 | 1.230 | 0.979 | 2.020 |
| Q61166 | Mapre1 | 0.016 | 0.079 | 0.211 | -0.152 | 0.000 | -0.500 | -0.907 | -1.160 |
| Q6P542 | Abcf1 | 0.342 | 0.582 | 0.361 | -0.057 | 0.000 | 0.327 | -0.666 | -0.406 |
| Q9DCE5 | Pak1ip1 | 0.033 | 0.120 | 0.064 | -0.123 | 0.000 | -0.401 | -0.996 | -0.500 |
| Q9JMH9 | Myo18a | 0.465 | 0.723 | -0.001 | 0.000 | 0.173 | 0.109 | -0.451 | 0.058 |
| O88456 | Capns1 | 0.707 | 0.983 | -0.351 | 0.263 | 0.000 | -0.211 | -0.070 | -0.033 |
| Q8BJ71 | Nup93 | 0.605 | 0.878 | 0.000 | 0.099 | -0.117 | 0.056 | -0.115 | 0.244 |
| O88520 | Shoc2 | 0.850 | 1.000 | 0.000 | -0.539 | 0.000 | -0.051 | -0.911 | 0.193 |
| P53810 | Pitpna | 0.002 | 0.023 | 0.000 | 0.117 | -0.135 | -1.010 | -1.530 | -1.350 |
| P61804 | Dad1 | 0.002 | 0.025 | 0.054 | 0.000 | -0.232 | 1.770 | 1.190 | 1.270 |
| P39749 | Fen1 | 0.169 | 0.351 | 0.002 | -0.215 | 0.000 | -0.243 | -0.778 | -0.197 |
| Q9WU81 | Slc37a2 | 0.276 | 0.497 | 0.431 | 0.000 | -0.628 | 0.442 | 0.392 | 0.173 |
| P14148 | Rpl7 | 0.006 | 0.044 | 0.207 | -0.004 | 0.000 | 0.767 | 0.494 | 0.739 |
| P61620 | Sec61a1 | 0.001 | 0.019 | 0.000 | 0.072 | -0.131 | 1.560 | 1.060 | 1.370 |
| Q61171 | Prdx2 | 0.040 | 0.134 | 0.089 | 0.000 | -0.208 | -0.353 | -0.755 | -0.402 |
| Q9R1P3 | Psmb2 | 0.006 | 0.044 | 0.222 | 0.000 | -0.028 | -0.810 | -1.210 | -1.570 |
| Q8C7X2 | Emc1 | 0.001 | 0.015 | 0.000 | 0.211 | -0.285 | 1.450 | 1.340 | 1.440 |
| Q9CQB5 | Cisd2 | 0.179 | 0.365 | 0.000 | 0.161 | -0.086 | 0.224 | 0.101 | 0.145 |
| Q8CHP5 | Pym1 | 0.018 | 0.083 | -0.344 | 0.000 | 0.040 | -0.801 | -1.330 | -0.753 |
| Q8K3A0 | Hscb | 0.378 | 0.627 | 0.020 | 0.000 | -7.410 | -0.071 | 0.082 | -0.054 |
| Q3V1T4 | P3h1 | 0.001 | 0.022 | 0.735 | -0.994 | 0.000 | 4.580 | 3.830 | 4.310 |
| Q8R3N6 | Thoc1 | 0.334 | 0.571 | 0.080 | 0.000 | -0.098 | -0.259 | -0.213 | 0.073 |
| Q64737 | Gart | 0.258 | 0.473 | 0.691 | 0.000 | -0.051 | 0.448 | -0.834 | -1.050 |
| Q920Q6 | Msi2 | 0.276 | 0.497 | 0.000 | 0.706 | -0.324 | 1.510 | 1.050 | -0.068 |
| Q9Z1Q5 | Clic1 | 0.005 | 0.039 | -0.013 | 0.084 | 0.000 | -0.653 | -1.240 | -1.010 |
| Q9D6K5 | Synj2bp | 0.056 | 0.167 | 0.000 | 0.262 | -0.061 | 0.573 | 0.279 | 0.805 |
| O54786 | Dffa | 0.079 | 0.210 | 0.000 | -0.630 | 0.300 | -0.522 | -1.400 | -1.040 |
| Q9EPL9 | Acox3 | 0.674 | 0.951 | 0.021 | 0.000 | -0.636 | 0.263 | -0.494 | -0.944 |
| P97384 | Anxa11 | 0.008 | 0.053 | 0.041 | 0.000 | -0.141 | -0.485 | -0.611 | -0.358 |
| O35381 | Anp32a | 0.000 | 0.012 | -0.159 | 0.084 | 0.000 | -1.290 | -1.590 | -1.380 |
| P07901 | Hsp90aa1 | 0.021 | 0.090 | 0.000 | 0.010 | -0.335 | -0.557 | -1.180 | -1.010 |
| Q64518 | Atp2a3 | 0.176 | 0.360 | 0.078 | 0.000 | -0.382 | -0.145 | -0.536 | -0.666 |
| Q9QZD8 | Slc25a10 | 0.003 | 0.030 | 0.000 | 0.214 | -0.415 | 1.190 | 1.260 | 1.510 |
| P52633 | Stat6 | 0.079 | 0.210 | 0.000 | -0.088 | 0.044 | -0.100 | -0.740 | -0.588 |
| Q02111 | Prkcq | 0.722 | 1.000 | 0.000 | 1.430 | -0.213 | -0.229 | 0.752 | 0.016 |
| P62305 | Snrpe | 0.123 | 0.283 | 0.000 | 0.203 | -0.302 | -0.339 | -0.394 | -1.200 |
| Q80YV2 | Zc3hc1 | 0.030 | 0.113 | -0.583 | 0.055 | 0.000 | -0.937 | -0.866 | -1.450 |
| Q5SUR0 | Pfas | 0.006 | 0.045 | -0.054 | 0.678 | 0.000 | -1.340 | -1.880 | -1.200 |
| Q80X50 | Ubap2l | 0.003 | 0.032 | 0.000 | -0.103 | 0.353 | 1.140 | 0.971 | 1.270 |
| Q9D8C4 | Ifi35 | 0.019 | 0.085 | -0.500 | 0.000 | 0.050 | -1.010 | -1.030 | -1.700 |
| Q7TMY8 | Huwe1 | 0.222 | 0.423 | 0.263 | -0.109 | 0.000 | 0.159 | -0.989 | -0.543 |
| Q9JII6 | Akr1a1 | 0.500 | 0.765 | 0.262 | -0.011 | 0.000 | 0.150 | -0.272 | 0.031 |
| P25425 | Pou2f1 | 0.006 | 0.047 | -0.147 | 0.120 | 0.000 | -0.422 | -0.622 | -0.496 |
| O89017 | Lgmn | 0.545 | 0.813 | 1.600 | -0.051 | 0.000 | 0.284 | -0.932 | 0.738 |
| Q9CWW6 | Pin4 | 0.376 | 0.624 | 0.000 | 0.180 | -0.252 | -0.110 | -0.344 | -0.071 |
| Q61733 | Mrps31 | 0.132 | 0.296 | 0.000 | 0.114 | -0.108 | 0.841 | -0.005 | 0.687 |
| Q8BHZ0 | Fam49a | 0.018 | 0.083 | 0.100 | 0.000 | -0.048 | -0.759 | -2.060 | -1.560 |
| Q61074 | Ppm1g | 0.015 | 0.076 | 0.076 | 0.000 | 0.000 | -0.823 | -1.880 | -1.110 |
| Q8R2U0 | Seh1l | 0.165 | 0.347 | -0.177 | 0.102 | 0.000 | -0.189 | -0.511 | -0.117 |
| Q64378 | Fkbp5 | 0.009 | 0.056 | -0.004 | 0.047 | 0.000 | -0.637 | -1.400 | -1.220 |

| | | | | | | | | | |
|---|---|---|---|---|---|---|---|---|---|
| Q91YE7 | Rbm5 | 0.008 | 0.052 | 0.000 | -0.307 | 0.016 | -0.952 | -1.520 | -0.983 |
| Q8BQ30 | Ppp1r18 | 0.409 | 0.660 | -0.391 | 0.000 | 0.142 | -0.536 | -0.383 | 0.028 |
| Q8CBW3 | Abi1 | 0.213 | 0.409 | -0.378 | 0.171 | 0.000 | 0.201 | 0.069 | 0.305 |
| P25976 | Ubtf | 0.014 | 0.072 | -0.184 | 0.000 | 0.021 | -0.544 | -0.926 | -0.527 |
| Q9CQH3 | Ndufb5 | 0.003 | 0.032 | 0.000 | 0.202 | -0.151 | 0.682 | 0.684 | 0.812 |
| Q8BH74 | Nup107 | 0.749 | 1.000 | -0.109 | 0.284 | 0.000 | -0.001 | -0.230 | 0.225 |
| P97429 | Anxa4 | 0.589 | 0.861 | 0.000 | 0.197 | -0.094 | 0.014 | -0.212 | 0.083 |
| P03921 | Mtnd5 | 0.051 | 0.156 | 0.000 | 0.416 | -0.059 | 0.695 | 0.639 | 1.400 |
| Q9CY50 | Ssr1 | 0.002 | 0.025 | 0.000 | 0.307 | -0.320 | 1.390 | 1.330 | 1.580 |
| P20060 | Hexb | 0.512 | 0.776 | 0.530 | -0.073 | 0.000 | 0.432 | -0.428 | -0.245 |
| Q922Y1 | Ubxn1 | 0.019 | 0.087 | 0.000 | -0.150 | 0.077 | -0.496 | -0.940 | -0.493 |
| P26350 | Ptma | 0.061 | 0.176 | 0.136 | 0.000 | -0.615 | -0.753 | -2.920 | -2.010 |
| Q8C3J5 | Dock2 | 0.006 | 0.047 | 0.009 | 0.000 | -0.011 | -0.628 | -1.100 | -0.665 |
| O35593 | Psmd14 | 0.036 | 0.125 | 0.000 | -0.104 | 0.054 | -0.306 | -0.715 | -0.334 |
| Q64213 | Sf1 | 0.008 | 0.054 | 0.000 | 0.089 | -0.065 | -0.205 | -0.298 | -0.238 |
| Q8C9V1 | Tbc1d10c | 0.042 | 0.139 | 0.000 | -0.523 | 0.236 | -0.798 | -1.680 | -0.940 |
| Q8K2I4 | Manba | 0.513 | 0.777 | 0.000 | 0.090 | -0.405 | 0.151 | -0.227 | 0.198 |
| Q7TNV0 | Dek | 0.006 | 0.047 | 0.137 | -0.007 | 0.000 | -0.513 | -1.050 | -0.888 |
| P62748 | Hpcal1 | 0.005 | 0.041 | 0.000 | 0.247 | -0.109 | -0.560 | -0.589 | -0.515 |
| P62918 | Rpl8 | 0.638 | 0.914 | 0.175 | -0.165 | 0.000 | 0.451 | -0.129 | -0.005 |
| Q9Z1J3 | Nfs1 | 0.866 | 1.000 | 0.239 | 0.000 | -0.042 | 0.878 | -0.349 | -0.124 |
| P17095 | Hmga1 | 0.277 | 0.498 | 0.000 | -0.453 | 0.203 | -0.194 | -1.160 | -0.274 |
| Q8VHR5 | Gatad2b | 0.198 | 0.392 | -0.373 | 0.000 | 0.093 | -0.487 | -0.543 | -0.135 |
| Q8BHG1 | Nrdc | 0.000 | 0.008 | -0.090 | 0.116 | 0.000 | -1.730 | -1.490 | -1.520 |
| P83741 | Wnk1 | 0.046 | 0.149 | -0.144 | 0.000 | 0.437 | -0.714 | -1.090 | -0.318 |
| O08582 | Gtpbp1 | 0.039 | 0.133 | 0.342 | -0.062 | 0.000 | -0.272 | -1.110 | -0.866 |
| Q9D0E1 | Hnrnpm | 0.116 | 0.271 | -0.089 | 0.068 | 0.000 | -0.065 | -0.477 | -0.243 |
| Q60631 | Grb2 | 0.005 | 0.041 | -0.091 | 0.403 | 0.000 | -1.080 | -1.510 | -0.944 |
| Q3UDW8 | Hgsnat | 0.535 | 0.803 | 0.177 | 0.000 | -0.116 | 0.499 | -0.142 | 0.120 |
| P58059 | Mrps21 | 0.054 | 0.163 | 0.000 | 0.089 | -0.066 | 0.647 | 0.140 | 0.589 |
| P14094 | Atp1b1 | 0.004 | 0.034 | -0.400 | 0.000 | 0.313 | 2.120 | 1.460 | 2.270 |
| P53395 | Dbt | 0.046 | 0.149 | 0.000 | 0.286 | -0.049 | 0.968 | 0.326 | 0.859 |
| Q9ESW4 | Agk | 0.183 | 0.372 | 0.000 | -0.572 | 0.077 | 0.527 | -0.141 | 0.703 |
| Q8BKS9 | Pum3 | 0.196 | 0.389 | 0.136 | 0.000 | -0.234 | -0.103 | -1.360 | -0.447 |
| O89110 | Casp8 | 0.012 | 0.067 | 0.166 | -0.032 | 0.000 | -1.050 | -1.120 | -2.060 |
| P42230 | Stat5a | 0.015 | 0.076 | -0.331 | 0.072 | 0.000 | -0.731 | -1.550 | -1.370 |
| Q9Z2L7 | Crlf3 | 0.015 | 0.076 | 0.078 | -0.288 | 0.000 | -0.879 | -2.030 | -1.700 |
| Q9CZ83 | Mrpl55 | 0.003 | 0.031 | 0.173 | 0.000 | -0.267 | 0.903 | 0.758 | 0.928 |
| O70492 | Snx3 | 0.413 | 0.665 | 0.054 | 0.000 | -0.003 | 0.688 | -0.177 | 0.226 |
| Q8BXA5 | Clptm1l | 0.009 | 0.056 | 0.000 | 0.069 | -0.288 | 0.729 | 0.507 | 0.942 |
| Q8BG48 | Stk17b | 0.005 | 0.039 | 0.000 | -0.182 | 0.023 | -1.390 | -1.950 | -1.130 |
| P46664 | Adss | 0.051 | 0.157 | 0.264 | -0.674 | 0.000 | -0.768 | -1.940 | -1.330 |
| Q9DCT2 | Ndufs3 | 0.021 | 0.090 | -0.343 | 0.080 | 0.000 | 0.597 | 0.506 | 1.070 |
| P62082 | Rps7 | 0.231 | 0.435 | 0.000 | 0.021 | -0.201 | 0.193 | -0.095 | 0.370 |
| Q3UM45 | Ppp1r7 | 0.011 | 0.061 | -0.459 | 0.000 | 0.147 | -1.720 | -1.920 | -1.040 |
| Q62465 | Vat1 | 0.029 | 0.110 | 0.420 | 0.000 | -0.104 | 1.170 | 0.561 | 1.040 |
| Q61107 | Gbp4 | 0.002 | 0.026 | 0.000 | -0.009 | 0.293 | -1.020 | -1.590 | -1.530 |
| P43275 | Hist1h1a | 0.473 | 0.733 | 0.000 | -0.398 | 0.015 | -0.138 | -0.623 | -0.127 |
| Q91VH2 | Snx9 | 0.007 | 0.051 | 0.000 | 0.205 | -0.383 | 1.650 | 1.140 | 0.994 |
| Q9CRD2 | Emc2 | 0.000 | 0.012 | 0.000 | 0.161 | -0.136 | 1.450 | 1.330 | 1.590 |
| Q99JX7 | Nxf1 | 0.309 | 0.542 | 0.393 | -0.510 | 0.000 | -0.281 | -0.727 | -0.190 |
| P63094 | Gnas | 0.019 | 0.087 | -0.209 | 0.111 | 0.000 | 0.694 | 0.307 | 0.605 |
| Q11011 | Npepps | 0.059 | 0.174 | 0.306 | 0.000 | -0.087 | -0.142 | -0.600 | -0.447 |
| P02535 | Krt10 | 0.910 | 1.000 | -0.437 | 0.000 | 0.988 | -0.544 | 0.928 | 0.386 |
| P97314 | Csrp2 | 0.001 | 0.019 | -1.840 | 0.000 | 0.161 | 5.250 | 4.780 | 5.010 |
| P18760 | Cfl1 | 0.003 | 0.033 | -0.271 | 0.034 | 0.000 | -0.898 | -1.220 | -0.868 |
| P62962 | Pfn1 | 0.000 | 0.012 | 0.000 | 0.175 | -0.052 | -1.240 | -1.510 | -1.260 |
| P49312 | Hnrnpa1 | 0.225 | 0.427 | -0.046 | 0.209 | 0.000 | -0.160 | -0.235 | 0.055 |
| P97333 | Nrp1 | 0.223 | 0.425 | -0.500 | 0.000 | 0.295 | -0.584 | -0.652 | -0.167 |

| | | | | | | | | | |
|---|---|---|---|---|---|---|---|---|---|
| Q8VDP6 | Cdipt | 0.044 | 0.142 | -0.068 | 0.082 | 0.000 | 0.492 | 0.133 | 0.466 |
| P51660 | Hsd17b4 | 0.034 | 0.122 | 0.027 | 0.000 | -0.287 | 1.110 | 0.293 | 0.795 |
| Q9D8N0 | Eef1g | 0.431 | 0.687 | 0.665 | -0.080 | 0.000 | 0.863 | -1.110 | -0.938 |
| Q05920 | Pc | 0.015 | 0.076 | 0.211 | 0.000 | -0.913 | 2.850 | 1.420 | 1.760 |
| P10649 | Gstm1 | 0.260 | 0.476 | 1.250 | 0.000 | -0.306 | 0.019 | -0.763 | -0.382 |
| P61226 | Rap2b | 0.010 | 0.059 | 0.419 | 0.000 | -0.170 | 1.050 | 0.866 | 0.897 |
| Q3TDN2 | Faf2 | 0.182 | 0.369 | -0.615 | 0.705 | 0.000 | 0.072 | 1.380 | 2.500 |
| Q8K358 | Pigu | 0.174 | 0.357 | 0.343 | 0.000 | -1.340 | 0.317 | 1.360 | 5.340 |
| P49442 | Inpp1 | 0.013 | 0.070 | -0.686 | 0.009 | 0.000 | -2.310 | -3.380 | -1.760 |
| P61957 | Sumo2 | 0.255 | 0.469 | 0.000 | -0.147 | 0.071 | -0.215 | -3.690 | -0.576 |
| Q8VCW8 | Acsf2 | 0.374 | 0.623 | 0.347 | -0.054 | 0.000 | 0.597 | -1.130 | -0.804 |
| Q8K1J6 | Trnt1 | 0.089 | 0.227 | -0.056 | 0.000 | 0.076 | -0.250 | -0.666 | -0.153 |
| Q02248 | Ctnnb1 | 0.001 | 0.018 | 0.000 | 1.470 | -0.278 | 6.150 | 5.320 | 5.830 |
| Q80VP0 | Tecpr1 | 0.211 | 0.407 | -0.458 | 0.000 | 0.797 | -0.779 | -0.878 | -0.031 |
| Q9WUN2 | Tbk1 | 0.159 | 0.336 | 0.000 | -0.120 | 0.216 | -0.002 | -1.180 | -0.558 |
| Q9CPR5 | Mrpl15 | 0.032 | 0.119 | 0.144 | -1.190 | 0.000 | 1.030 | 0.900 | 1.190 |
| Q8CH25 | Sltm | 0.673 | 0.950 | -0.415 | 0.000 | 0.073 | -0.058 | -0.117 | 0.051 |
| Q8CBA2 | Slfn5 | 0.102 | 0.247 | 0.000 | -0.213 | 0.161 | -0.133 | -0.594 | -0.422 |
| Q9DCW4 | Etfb | 0.322 | 0.558 | 0.252 | 0.000 | -0.128 | 0.867 | -0.052 | 0.291 |
| Q9DBR0 | Akap8 | 0.021 | 0.092 | 0.000 | 0.298 | -0.040 | -0.520 | -0.856 | -0.359 |
| Q3UQU0 | Brd9 | 0.145 | 0.314 | -0.026 | 0.000 | 0.187 | -0.021 | -1.600 | -0.720 |
| Q8VD26 | Tmem143 | 0.001 | 0.022 | -0.252 | 0.062 | 0.000 | 0.775 | 0.891 | 1.040 |
| O35638 | Stag2 | 0.144 | 0.311 | 0.000 | 0.186 | -0.178 | -0.177 | -0.493 | -1.410 |
| Q8C1D8 | Iws1 | 0.115 | 0.270 | 0.000 | -0.155 | 0.078 | -0.329 | -0.906 | -0.206 |
| Q9Z315 | Sart1 | 0.004 | 0.038 | -0.166 | 0.000 | 0.034 | -0.707 | -1.120 | -0.776 |
| Q9CQ54 | Ndufc2 | 0.001 | 0.022 | 0.044 | 0.000 | -0.180 | 0.889 | 0.645 | 0.923 |
| P17918 | Pcna | 0.142 | 0.309 | 0.516 | 0.000 | -0.203 | -0.004 | -0.794 | -0.676 |
| Q9CX86 | Hnrnpa0 | 0.237 | 0.445 | -0.227 | 0.000 | 0.302 | 0.083 | -0.505 | -0.595 |
| Q8VCY6 | Utp6 | 0.039 | 0.133 | 0.158 | -0.187 | 0.000 | -0.292 | -0.675 | -0.422 |
| P99024 | Tubb5 | 0.970 | 1.000 | 0.000 | -0.069 | 0.025 | 0.542 | -0.296 | -0.257 |
| P62317 | Snrpd2 | 0.037 | 0.129 | -0.005 | 0.066 | 0.000 | -0.224 | -0.407 | -0.116 |
| Q9CQ75 | Ndufa2 | 0.016 | 0.078 | -0.063 | 0.189 | 0.000 | 0.742 | 0.367 | 0.640 |
| P45376 | Akr1b1 | 0.002 | 0.023 | -0.133 | 0.414 | 0.000 | -1.310 | -1.620 | -1.300 |
| Q00899 | Yy1 | 0.078 | 0.208 | 0.000 | -0.096 | 0.192 | -0.240 | -1.020 | -0.413 |
| P23780 | Glb1 | 0.709 | 0.985 | 0.541 | 0.000 | -0.380 | 0.396 | 0.028 | 0.086 |
| Q8BKJ9 | Sirt7 | 0.048 | 0.152 | 0.000 | -0.263 | 0.089 | -0.314 | -0.839 | -0.582 |
| Q3UUV5 | Skap1 | 0.002 | 0.023 | -0.314 | 0.000 | 0.236 | -1.930 | -2.300 | -1.630 |
| Q91VM5 | Rbmxl1 | 0.172 | 0.354 | -0.370 | 0.000 | 0.091 | -0.513 | -0.512 | -0.165 |
| P09671 | Sod2 | 0.023 | 0.096 | -0.061 | 0.348 | 0.000 | 0.715 | 0.476 | 0.682 |
| Q9D958 | Spcs1 | 0.952 | 1.000 | 0.410 | -1.850 | 0.000 | 0.580 | -2.270 | 0.466 |
| O88844 | Idh1 | 0.617 | 0.891 | 0.610 | 0.000 | -0.573 | 0.268 | 0.140 | 0.187 |
| Q9DB42 | Znf593 | 0.300 | 0.529 | -0.360 | 0.000 | 0.352 | -0.645 | -0.477 | 0.060 |
| P04441 | Cd74 | 0.860 | 1.000 | 0.000 | 0.083 | -0.265 | -0.059 | -0.135 | -0.049 |
| O55029 | Copb2 | 0.051 | 0.158 | 0.023 | 0.000 | -0.067 | -0.281 | -1.100 | -0.638 |
| P52431 | Pold1 | 0.005 | 0.043 | -0.206 | 0.235 | 0.000 | -0.797 | -1.220 | -0.903 |
| Q9Z1D1 | Eif3g | 0.484 | 0.747 | 0.391 | 0.000 | -0.762 | 0.185 | -1.290 | -0.524 |
| P27048 | Snrpb | 0.016 | 0.079 | 0.000 | 0.086 | -0.061 | -0.366 | -0.857 | -0.543 |
| Q8K4L3 | Svil | 0.677 | 0.953 | 0.060 | -0.124 | 0.000 | 0.329 | -0.177 | -0.004 |
| Q99LP6 | Grpel1 | 0.003 | 0.030 | 0.000 | 0.185 | -0.177 | 1.010 | 0.736 | 0.977 |
| Q9CR62 | Slc25a11 | 0.026 | 0.105 | 0.013 | 0.000 | -0.081 | 0.658 | 0.233 | 0.369 |
| O35309 | Nmi | 0.000 | 0.008 | 0.000 | -0.038 | 0.026 | -0.966 | -1.090 | -0.910 |
| Q8BKC5 | Ipo5 | 0.805 | 1.000 | 0.128 | 0.000 | -0.716 | 0.189 | -0.657 | -0.408 |
| Q8BH97 | Rcn3 | 0.000 | 0.010 | 0.000 | 0.300 | -1.020 | 5.700 | 5.470 | 5.630 |
| P42932 | Cct8 | 0.038 | 0.129 | 0.019 | 0.000 | -0.027 | -0.248 | -0.835 | -0.494 |
| Q8BHJ5 | Tbl1xr1 | 0.162 | 0.341 | -0.117 | 0.000 | 0.120 | -0.368 | -2.330 | -0.539 |
| Q9WUK4 | Rfc2 | 0.057 | 0.169 | 0.113 | 0.000 | -0.228 | -0.245 | -0.935 | -0.770 |
| Q8BH24 | Tm9sf4 | 0.003 | 0.032 | 0.024 | 0.000 | -0.422 | 0.875 | 0.752 | 0.962 |
| Q921G8 | Tubgcp2 | 0.065 | 0.184 | 0.000 | -0.034 | 0.085 | -0.377 | -0.541 | -0.083 |
| Q3UPL0 | Sec31a | 0.955 | 1.000 | 0.030 | -0.312 | 0.000 | 0.370 | -0.502 | -0.200 |

| | | | | | | | | | |
|---|---|---|---|---|---|---|---|---|---|
| P45952 | Acadm | 0.441 | 0.697 | 0.020 | 0.000 | -0.423 | 0.852 | -0.186 | -0.129 |
| Q9Z2E1 | Mbd2 | 0.061 | 0.176 | -0.257 | 0.000 | 0.089 | -0.475 | -0.467 | -0.235 |
| Q6ZQF7 | Jade2 | 0.007 | 0.048 | -0.162 | 0.087 | 0.000 | -0.756 | -0.903 | -0.512 |
| Q9ERN0 | Scamp2 | 0.659 | 0.935 | 0.258 | 0.000 | -0.346 | 0.217 | -0.347 | -0.321 |
| O35900 | Lsm2 | 0.000 | 0.011 | -0.100 | 0.000 | 0.134 | -1.180 | -1.020 | -1.150 |
| Q9CX30 | Yif1b | 0.001 | 0.018 | -0.142 | 0.000 | 0.081 | 1.740 | 1.320 | 1.910 |
| P51912 | Slc1a5 | 0.002 | 0.025 | 0.161 | 0.000 | -0.159 | 0.905 | 0.762 | 1.060 |
| Q80ZS3 | Mrps26 | 0.050 | 0.155 | -0.354 | 0.562 | 0.000 | 0.950 | 0.666 | 0.952 |
| Q99LH2 | Ptdss1 | 0.004 | 0.038 | 0.073 | 0.000 | -0.255 | 0.781 | 1.180 | 0.770 |
| Q01965 | Ly9 | 0.539 | 0.807 | 0.248 | -0.106 | 0.000 | -0.213 | -0.306 | 0.254 |
| P23198 | Cbx3 | 0.053 | 0.162 | -0.100 | 0.000 | 0.464 | -0.392 | -0.638 | -0.271 |
| O09061 | Psmb1 | 0.002 | 0.022 | 0.000 | 0.060 | -0.048 | -0.871 | -1.280 | -0.911 |
| P63028 | Tpt1 | 0.111 | 0.264 | -0.589 | 1.150 | 0.000 | -2.170 | -1.060 | -0.523 |
| Q9Z2Q5 | Mrpl40 | 0.003 | 0.031 | 0.154 | -0.056 | 0.000 | 0.713 | 0.525 | 0.546 |
| Q9ES46 | Parvb | 0.600 | 0.873 | 0.000 | 0.158 | -1.990 | -0.638 | -0.108 | -3.010 |
| E9PVA8 | Gcn1 | 0.557 | 0.825 | 0.132 | 0.000 | -0.329 | 0.500 | -0.383 | 0.255 |
| O88291 | Znf326 | 0.531 | 0.798 | -0.617 | 0.000 | 0.031 | -0.141 | -0.078 | 0.089 |
| E9Q394 | Akap13 | 0.012 | 0.068 | -0.244 | 0.000 | 0.099 | -0.813 | -1.320 | -0.739 |
| Q8C2K5 | Rasal3 | 0.018 | 0.083 | -0.082 | 0.141 | 0.000 | -0.512 | -1.320 | -0.924 |
| P05555 | Itgam | 0.903 | 1.000 | 1.640 | -0.097 | 0.000 | 1.180 | 0.078 | 0.022 |
| P23475 | Xrcc6 | 0.009 | 0.056 | 0.040 | 0.000 | -0.446 | -0.978 | -1.630 | -1.300 |
| Q04447 | Ckb | 0.070 | 0.192 | 0.000 | 0.157 | -0.135 | -0.164 | -0.636 | -0.365 |
| Q924T2 | Mrps2 | 0.014 | 0.073 | 0.000 | 0.348 | -0.330 | 0.736 | 1.060 | 0.937 |
| Q9Z1M8 | Ik | 0.021 | 0.092 | 0.000 | 0.281 | -0.201 | -0.702 | -0.501 | -0.461 |
| D3YXK2 | Safb | 0.010 | 0.060 | -0.321 | 0.077 | 0.000 | -0.723 | -0.601 | -0.827 |
| Q9CQA3 | Sdhb | 0.005 | 0.042 | 0.000 | 0.126 | -0.143 | 0.648 | 0.433 | 0.552 |
| O70251 | Eef1b | 0.042 | 0.140 | 0.083 | -0.155 | 0.000 | -0.239 | -0.809 | -0.633 |
| Q6Q899 | Ddx58 | 0.144 | 0.311 | 0.000 | 0.060 | -0.766 | -0.397 | -1.090 | -1.250 |
| Q9CQ71 | Rpa3 | 0.181 | 0.368 | -0.082 | 0.000 | 0.202 | -0.347 | -0.889 | -3.700 |
| P21447 | Abcb1a | 0.208 | 0.404 | -0.389 | 0.059 | 0.000 | -0.616 | -0.615 | -0.099 |
| Q99JI4 | Psmd6 | 0.028 | 0.107 | 0.000 | 0.155 | -0.067 | -0.324 | -0.253 | -0.600 |
| Q9DAS9 | Gng12 | 0.017 | 0.080 | 1.250 | -0.499 | 0.000 | 2.910 | 2.090 | 2.560 |
| Q8CH02 | Sugp1 | 0.095 | 0.237 | -0.273 | 0.000 | 0.142 | -0.752 | -0.679 | -0.159 |
| O55128 | Sap18 | 0.043 | 0.142 | -0.163 | 0.000 | 0.168 | -0.507 | -2.010 | -1.380 |
| Q8CG76 | Akr7a2 | 0.383 | 0.632 | -0.121 | 0.000 | 0.029 | 0.397 | -0.088 | 0.045 |
| Q9QY24 | Zbp1 | 0.179 | 0.364 | -0.179 | 0.000 | 0.456 | -0.213 | -0.556 | -0.097 |
| Q8BJW6 | Eif2a | 0.743 | 1.000 | 0.280 | -0.084 | 0.000 | 0.391 | -0.452 | -0.025 |
| Q8C2Q3 | Rbm14 | 0.597 | 0.870 | -0.370 | 0.000 | 0.306 | -0.338 | -0.331 | 0.164 |
| Q9D3P8 | Plgrkt | 0.012 | 0.069 | -0.004 | 0.232 | 0.000 | 0.479 | 0.379 | 0.528 |
| Q3UJD6 | Usp19 | 0.246 | 0.457 | 0.704 | -0.028 | 0.000 | 0.999 | 0.247 | 0.759 |
| Q9ER69 | Wtap | 0.018 | 0.084 | 0.000 | -0.142 | 0.111 | -0.903 | -1.710 | -0.814 |
| Q9D8B3 | Chmp4b | 0.033 | 0.119 | 0.000 | -0.068 | 0.070 | -0.181 | -0.595 | -0.445 |
| Q9WVJ2 | Psmd13 | 0.040 | 0.134 | 0.000 | -0.037 | 0.010 | -0.179 | -0.699 | -0.638 |
| Q9D358 | Acp1 | 0.010 | 0.060 | 0.000 | 0.034 | -0.377 | -0.763 | -1.300 | -1.250 |
| P11031 | Sub1 | 0.128 | 0.291 | 0.485 | 0.000 | -0.229 | -0.017 | -1.030 | -0.866 |
| Q6ZWU9 | Rps27 | 0.484 | 0.747 | 0.000 | 0.208 | -0.051 | 0.074 | -0.521 | 0.101 |
| Q64282 | Ifit1 | 0.387 | 0.636 | 0.484 | -0.260 | 0.000 | 0.741 | -0.075 | 0.507 |
| Q9JK48 | Sh3glb1 | 0.086 | 0.222 | 0.000 | 0.081 | -0.382 | -0.354 | -0.952 | -0.542 |
| Q8BY89 | Slc44a2 | 0.203 | 0.400 | 0.143 | -0.178 | 0.000 | 0.730 | -0.130 | 0.759 |
| P98083 | Shc1 | 0.721 | 1.000 | 0.042 | -0.011 | 0.000 | 1.180 | -0.727 | 0.209 |
| Q9EQ28 | Pold3 | 0.000 | 0.013 | 0.036 | 0.000 | -0.085 | -0.691 | -0.831 | -0.668 |
| Q8BTI8 | Srrm2 | 0.272 | 0.492 | -0.683 | 0.000 | 0.080 | -0.469 | -0.866 | -0.362 |
| Q9CPW4 | Arpc5 | 0.450 | 0.708 | 0.000 | -0.098 | 0.034 | 0.163 | -0.086 | 0.066 |
| Q9WUU7 | Ctsz | 0.118 | 0.274 | 0.656 | -0.102 | 0.000 | 0.028 | -1.080 | -0.862 |
| Q922Q1 | Mosc 2 | 0.217 | 0.416 | 0.000 | 0.380 | -0.531 | 0.398 | 0.188 | 0.486 |
| Q3THS6 | Mat2a | 0.000 | 0.013 | 0.099 | -0.109 | 0.000 | -0.881 | -1.060 | -0.919 |
| Q6NVF9 | Cpsf6 | 0.003 | 0.032 | 0.000 | -0.136 | 0.019 | -0.650 | -0.911 | -0.604 |
| P05202 | Got2 | 0.002 | 0.028 | 0.000 | 0.188 | -0.073 | 0.799 | 0.617 | 0.665 |
| P62874 | Gnb1 | 0.000 | 0.013 | 0.000 | 0.012 | -0.274 | 1.780 | 1.420 | 1.780 |

| | | | | | | | | | |
|---|---|---|---|---|---|---|---|---|---|
| O08756 | Hsd17b10 | 0.087 | 0.224 | -0.012 | 0.447 | 0.000 | 0.862 | 0.393 | 0.563 |
| Q9JL35 | Hmgn5 | 0.058 | 0.171 | -0.153 | 0.000 | 0.090 | -0.490 | -0.558 | -0.153 |
| Q05816 | Fabp5 | 0.681 | 0.957 | 0.000 | 0.842 | -2.280 | 0.637 | -5.690 | 0.568 |
| Q9QYB5 | Add3 | 0.122 | 0.281 | 0.000 | -0.422 | 0.087 | -0.199 | -1.070 | -0.858 |
| Q64133 | Maoa | 0.000 | 0.011 | 0.151 | 0.000 | -0.761 | 3.640 | 3.580 | 3.770 |
| P22315 | Fech | 0.179 | 0.364 | 0.378 | -1.300 | 0.000 | 1.010 | 0.491 | 0.277 |
| Q80V26 | Impad1 | 0.000 | 0.014 | 0.210 | -0.225 | 0.000 | 1.400 | 1.310 | 1.370 |
| Q9DAA6 | Exosc1 | 0.980 | 1.000 | 0.000 | 1.410 | -0.356 | -0.065 | 0.682 | 0.391 |
| Q921K9 | Bcl7b | 0.026 | 0.105 | -0.081 | 0.372 | 0.000 | -0.672 | -1.020 | -0.387 |
| Q99PV8 | Bcl11b | 0.314 | 0.547 | -0.287 | 0.000 | 0.223 | -0.356 | -0.661 | 0.065 |
| Q9CPQ8 | Atp5l | 0.001 | 0.022 | -0.234 | 0.085 | 0.000 | 0.683 | 0.763 | 0.798 |
| P24369 | Ppib | 0.003 | 0.033 | 0.000 | 0.096 | -0.188 | 0.909 | 0.584 | 0.780 |
| Q99LE6 | Abcf2 | 0.419 | 0.673 | 0.556 | -0.007 | 0.000 | 0.704 | 0.144 | 0.369 |
| Q8BHB4 | Wdr3 | 0.325 | 0.561 | -0.953 | 0.000 | 0.292 | -0.509 | -0.492 | -1.080 |
| P21956 | Mfge8 | 0.010 | 0.061 | 0.000 | -0.252 | 0.038 | 1.480 | 0.660 | 1.380 |
| O55126 | Gbas | 0.027 | 0.106 | 1.180 | -0.355 | 0.000 | 1.730 | 1.880 | 2.050 |
| O35368 | Ifi203 | 0.019 | 0.087 | -0.210 | 0.033 | 0.000 | -0.572 | -1.050 | -1.390 |
| Q8CH18 | Ccar1 | 0.158 | 0.335 | -0.956 | 0.094 | 0.000 | -0.824 | -1.620 | -0.710 |
| Q3UMR5 | Mcu | 0.004 | 0.038 | 0.000 | 0.207 | -0.231 | 0.923 | 0.701 | 0.918 |
| P84099 | Rpl19 | 0.926 | 1.000 | 0.090 | 0.000 | -0.109 | 0.249 | -0.392 | 0.186 |
| Q9R0H0 | Acox1 | 0.000 | 0.012 | 0.521 | 0.000 | -0.037 | 2.920 | 2.570 | 2.770 |
| Q9D024 | Ccdc47 | 0.003 | 0.031 | 0.000 | 0.288 | -0.108 | 1.170 | 0.992 | 1.430 |
| Q9WUR2 | Eci2 | 0.392 | 0.641 | -0.002 | 0.400 | 0.000 | 0.637 | 0.088 | 0.274 |
| Q60766 | Irgm1 | 0.018 | 0.083 | 0.051 | 0.000 | -0.187 | 0.495 | 0.228 | 0.373 |
| P62983 | Rps27a | 0.185 | 0.374 | 0.000 | -0.150 | 0.017 | -0.164 | -0.623 | -0.142 |
| Q6ZWV3 | Rpl10 | 0.061 | 0.176 | 0.051 | -0.075 | 0.000 | 0.587 | 0.110 | 0.392 |
| B9EJ86 | Osbpl8 | 0.137 | 0.302 | 0.203 | 0.000 | -0.375 | 0.463 | 0.106 | 0.376 |
| Q9Z2X1 | Hnrnpf | 0.015 | 0.076 | -0.128 | 0.274 | 0.000 | -0.417 | -0.823 | -0.664 |
| P35700 | Prdx1 | 0.130 | 0.293 | 0.028 | 0.000 | -0.030 | 0.421 | -0.006 | 0.325 |
| P16330 | Cnp | 0.135 | 0.298 | 0.000 | 0.133 | -0.102 | -0.008 | -0.671 | -0.448 |
| Q61735 | Cd47 | 0.827 | 1.000 | 0.608 | 0.000 | -0.357 | -0.209 | -0.255 | 1.080 |
| Q05CL8 | Larp7 | 0.002 | 0.029 | 0.038 | -0.083 | 0.000 | -0.704 | -1.150 | -0.895 |
| Q61578 | Fdxr | 0.099 | 0.243 | 0.000 | 0.172 | -0.061 | -0.043 | -0.801 | -0.540 |
| Q61136 | Prpf4b | 0.129 | 0.291 | 0.000 | 0.121 | -0.928 | -0.820 | -0.932 | -0.971 |
| O88559 | Men1 | 0.002 | 0.025 | -0.161 | 0.000 | 0.096 | -0.644 | -0.763 | -0.597 |
| Q8K363 | Ddx18 | 0.197 | 0.391 | 0.093 | -0.221 | 0.000 | 0.055 | -0.960 | -0.698 |
| Q8JZQ2 | Afg3l2 | 0.015 | 0.076 | 0.000 | 0.090 | -0.109 | 0.828 | 0.358 | 0.590 |
| P62313 | Lsm6 | 0.017 | 0.081 | 0.000 | -0.578 | 0.234 | -1.060 | -1.350 | -1.030 |
| P62259 | Ywhae | 0.027 | 0.105 | 0.000 | 0.068 | -0.183 | -0.420 | -1.160 | -0.884 |
| Q61753 | Phgdh | 0.305 | 0.536 | 0.093 | 0.000 | -0.202 | 0.150 | -0.694 | -0.523 |
| Q505F5 | Lrrc47 | 0.193 | 0.385 | 0.492 | 0.000 | -0.011 | 0.172 | -0.814 | -0.432 |
| P07742 | Rrm1 | 0.156 | 0.330 | 1.310 | -0.668 | 0.000 | -0.320 | -1.880 | -1.010 |
| Q9CSN1 | Snw1 | 0.101 | 0.247 | 0.000 | -0.182 | 0.300 | -0.364 | -0.567 | -0.141 |
| Q9D880 | Timm50 | 0.034 | 0.123 | 0.000 | 0.165 | -0.056 | 0.558 | 0.221 | 0.486 |
| Q6ZQ88 | Kdm1a | 0.170 | 0.353 | 0.000 | 0.467 | -0.476 | -0.467 | -0.683 | -0.320 |
| Q08642 | Padi2 | 0.002 | 0.027 | 0.000 | -0.067 | 0.169 | -1.410 | -1.390 | -0.906 |
| Q7TSG2 | Ctdp1 | 0.053 | 0.162 | -0.333 | 0.000 | 0.669 | -0.845 | -1.500 | -0.585 |
| P97496 | Smarcc1 | 0.887 | 1.000 | -0.284 | 1.510 | 0.000 | 0.430 | 1.030 | -0.553 |
| Q9CPR4 | Rpl17 | 0.058 | 0.171 | 0.038 | 0.000 | -0.028 | 0.436 | 0.083 | 0.437 |
| Q6A068 | Cdc5l | 0.072 | 0.196 | -0.551 | 0.000 | 0.012 | -0.825 | -0.927 | -0.473 |
| P53986 | Slc16a1 | 0.170 | 0.352 | 0.555 | 0.000 | -0.036 | 1.090 | 0.354 | 0.539 |
| Q60902 | Eps15l1 | 0.605 | 0.878 | 0.310 | -0.100 | 0.000 | -0.085 | -0.310 | 0.258 |
| Q9QXT0 | Cnpy2 | 0.001 | 0.019 | 0.000 | 0.051 | -0.133 | 1.060 | 0.742 | 0.879 |
| Q6A4J8 | Usp7 | 0.012 | 0.067 | 0.000 | -0.009 | 0.033 | -0.659 | -1.400 | -0.850 |
| Q9CYL5 | Glipr2 | 0.076 | 0.205 | 0.000 | -0.110 | 0.013 | 0.537 | 0.043 | 0.419 |
| Q9CQW1 | Ykt6 | 0.330 | 0.568 | -0.082 | 0.999 | 0.000 | 0.328 | 0.902 | 1.060 |
| Q61595 | Ktn1 | 0.022 | 0.095 | 0.000 | -0.106 | 0.163 | 0.570 | 0.650 | 1.220 |
| Q6PF93 | Pik3c3 | 0.004 | 0.034 | 0.000 | 0.082 | -0.066 | -0.656 | -0.946 | -1.160 |
| P70296 | Pebp1 | 0.000 | 0.012 | -0.033 | 0.000 | 0.005 | -1.230 | -1.610 | -1.490 |

| | | | | | | | | | |
|---|---|---|---|---|---|---|---|---|---|
| P28271 | Aco1 | 0.002 | 0.024 | 0.000 | -0.026 | 0.042 | -0.794 | -1.280 | -1.150 |
| Q9QXB9 | Drg2 | 0.746 | 1.000 | 0.545 | -0.088 | 0.000 | 0.489 | -0.337 | -0.021 |
| P29341 | Pabpc1 | 0.216 | 0.413 | 0.000 | 0.523 | -0.083 | -0.141 | -0.164 | -0.096 |
| Q64735 | Cr1l | 0.871 | 1.000 | 0.107 | 0.000 | -0.145 | 0.063 | -0.267 | 0.095 |
| Q8BGU5 | Ccny | 0.233 | 0.438 | -0.007 | 0.027 | 0.000 | 0.181 | -0.036 | 0.360 |
| Q8K2C9 | Hacd3 | 0.081 | 0.213 | -0.211 | 0.067 | 0.000 | 0.124 | 0.145 | 0.335 |
| O70439 | Stx7 | 0.141 | 0.308 | 0.241 | 0.000 | -0.122 | -0.013 | -0.452 | -0.348 |
| Q8K2C7 | Os9 | 0.015 | 0.075 | -0.244 | 0.661 | 0.000 | 1.410 | 1.240 | 1.200 |
| P62482 | Kcnab2 | 0.556 | 0.825 | 0.000 | 0.764 | -1.840 | -0.436 | -2.640 | -0.126 |
| Q501J6 | Ddx17 | 0.007 | 0.048 | -0.046 | 0.116 | 0.000 | -0.532 | -0.750 | -0.397 |
| P20065 | Tmsb4x | 0.010 | 0.058 | 0.187 | 0.000 | -0.161 | -1.300 | -2.470 | -1.480 |
| P23708 | Nfya | 0.195 | 0.387 | 0.098 | 0.000 | -0.157 | -0.432 | -0.614 | -3.200 |
| Q922Q9 | Chid1 | 0.852 | 1.000 | 0.030 | 0.000 | -0.341 | -0.419 | 0.214 | -0.239 |
| P70399 | Tp53bp1 | 0.062 | 0.178 | 0.267 | 0.000 | -0.013 | -0.418 | -1.440 | -0.510 |
| Q3TIR3 | Ric8a | 0.860 | 1.000 | 0.000 | -0.572 | 0.168 | 0.033 | -1.140 | 0.413 |
| Q5SSL4 | Abr | 0.067 | 0.188 | 0.000 | -0.140 | 0.026 | -0.313 | -1.340 | -0.724 |
| Q9CZM2 | Rpl15 | 0.017 | 0.080 | 0.110 | 0.000 | -0.154 | 0.555 | 0.302 | 0.373 |
| Q80UW8 | Polr2e | 0.036 | 0.126 | 0.000 | -0.030 | 0.211 | -0.203 | -0.628 | -0.347 |
| Q9Z2A7 | Dgat1 | 0.041 | 0.136 | 0.000 | 0.214 | -0.404 | 0.617 | 0.361 | 0.650 |
| Q99LJ6 | Gpx7 | 0.000 | 0.012 | 0.150 | -1.130 | 0.000 | 5.360 | 4.910 | 5.340 |
| Q9WTX6 | Cul1 | 0.660 | 0.936 | 0.000 | -2.550 | 0.489 | 0.572 | -1.490 | -3.190 |
| Q924Z4 | Cers2 | 0.033 | 0.119 | 0.000 | 0.616 | -0.229 | 0.912 | 0.981 | 0.929 |
| Q9Z103 | Adnp | 0.796 | 1.000 | -0.620 | 0.000 | 0.145 | -0.140 | -0.320 | 0.219 |
| Q07113 | Igf2r | 0.000 | 0.008 | 0.000 | -0.092 | 0.121 | 1.850 | 1.700 | 1.910 |
| Q5XKN4 | Jagn1 | 0.013 | 0.069 | -0.346 | 0.043 | 0.000 | 0.703 | 0.682 | 1.270 |
| Q9D706 | Rpap3 | 0.076 | 0.205 | 0.123 | -0.216 | 0.000 | -0.201 | -0.665 | -0.416 |
| Q9JHS3 | Lamtor2 | 0.211 | 0.407 | 0.426 | -0.037 | 0.000 | 0.206 | -0.733 | -0.512 |
| A2AGT5 | Ckap5 | 0.082 | 0.216 | 0.000 | -0.236 | 0.124 | -0.162 | -0.706 | -0.630 |
| Q60865 | Caprin1 | 0.278 | 0.499 | 0.134 | 0.000 | -0.385 | -0.068 | -0.729 | -0.380 |
| Q8K019 | Bclaf1 | 0.022 | 0.095 | -0.039 | 0.072 | 0.000 | -0.301 | -0.823 | -0.523 |
| Q8BWF2 | Gimap5 | 0.015 | 0.075 | -0.145 | 0.000 | 0.167 | -0.476 | -0.802 | -0.463 |
| Q99MI6 | Gimap3 | 0.015 | 0.075 | -0.145 | 0.000 | 0.167 | -0.476 | -0.802 | -0.463 |
| Q8R323 | Rfc3 | 0.159 | 0.336 | 0.374 | 0.000 | -0.078 | 0.115 | -1.040 | -0.693 |
| P97315 | Csrp1 | 0.051 | 0.158 | -2.340 | 0.000 | 1.880 | 2.360 | 4.050 | 4.750 |
| Q8C522 | Endod1 | 0.757 | 1.000 | 0.000 | 0.145 | -0.042 | 0.232 | -0.229 | -0.045 |
| Q9CQV8 | Ywhab | 0.002 | 0.028 | -0.049 | 0.104 | 0.000 | -0.851 | -1.410 | -1.150 |
| Q921F2 | Tardbp | 0.080 | 0.211 | 0.004 | 0.000 | -0.021 | -0.142 | -0.728 | -0.349 |
| P63280 | Ube2i | 0.764 | 1.000 | 0.000 | 0.245 | -0.255 | -0.134 | -0.038 | 0.016 |
| P68254 | Ywhaq | 0.006 | 0.046 | -0.139 | 0.000 | 0.050 | -0.858 | -1.630 | -1.230 |
| Q8VDL4 | Adpgk | 0.170 | 0.352 | 0.785 | 0.000 | -0.022 | 1.230 | 0.505 | 0.737 |
| Q5XJE5 | Leo1 | 0.033 | 0.119 | -0.326 | 0.153 | 0.000 | -0.839 | -0.635 | -1.470 |
| Q4FK66 | Prpf38a | 0.020 | 0.088 | 0.219 | -0.245 | 0.000 | -0.513 | -0.993 | -0.699 |
| Q9D855 | Uqcrb | 0.013 | 0.069 | 0.000 | 0.173 | -0.233 | 0.669 | 0.448 | 0.554 |
| Q5M8N4 | Sdr39u1 | 0.975 | 1.000 | -0.090 | 1.850 | 0.000 | -0.753 | 1.040 | 1.390 |
| Q8BRN9 | Cc2d1b | 0.026 | 0.105 | 0.000 | 0.668 | -0.145 | -1.050 | -1.940 | -0.829 |
| Q8BH69 | Sephs1 | 0.009 | 0.056 | -0.042 | 0.000 | 0.303 | -1.280 | -1.270 | -0.629 |
| P70302 | Stim1 | 0.048 | 0.152 | 0.074 | 0.000 | -0.139 | 0.247 | 0.124 | 0.379 |
| P19536 | Cox5b | 0.287 | 0.512 | -0.201 | 0.104 | 0.000 | 0.204 | -0.054 | 0.216 |
| Q62093 | Srsf2 | 0.063 | 0.181 | 0.470 | -0.085 | 0.000 | -0.167 | -1.240 | -1.120 |
| Q8R050 | Gspt1 | 0.272 | 0.492 | 0.345 | -0.104 | 0.000 | 0.231 | -0.805 | -0.456 |
| Q791V5 | Mtch2 | 0.007 | 0.048 | 0.000 | 0.565 | -0.017 | 1.160 | 1.140 | 1.190 |
| P35279 | Rab6a | 0.697 | 0.973 | 0.155 | 0.000 | -0.059 | 0.176 | -0.128 | 0.205 |
| P57784 | Snrpa1 | 0.013 | 0.070 | -0.141 | 0.155 | 0.000 | -0.593 | -0.649 | -0.350 |
| P14869 | Rplp0 | 0.093 | 0.235 | 0.077 | 0.000 | -0.078 | 0.564 | 0.057 | 0.406 |
| P61022 | Chp1 | 0.001 | 0.019 | 0.138 | -0.075 | 0.000 | 1.490 | 1.050 | 1.450 |
| Q8BLF1 | Nceh1 | 0.001 | 0.019 | 0.000 | 0.056 | -0.064 | 0.628 | 0.446 | 0.552 |
| O54890 | Itgb3 | 0.385 | 0.633 | 0.000 | 0.662 | -0.700 | -0.246 | -0.257 | -0.808 |
| Q9JJ28 | Flii | 0.231 | 0.435 | 0.000 | 0.161 | -0.371 | -0.054 | -0.816 | -0.487 |
| O70310 | Nmt1 | 0.544 | 0.811 | 0.098 | -0.428 | 0.000 | 0.144 | -1.000 | -0.217 |

| | | | | | | | | | |
|---|---|---|---|---|---|---|---|---|---|
| Q3U1J4 | Ddb1 | 0.021 | 0.091 | -0.034 | 0.216 | 0.000 | -0.354 | -1.040 | -0.931 |
| Q9Z130 | Hnrnpdl | 0.499 | 0.764 | 0.000 | 0.040 | -2.550 | -0.201 | -0.411 | 0.032 |
| P62257 | Ube2h | 0.503 | 0.767 | 0.000 | -0.443 | 0.167 | -0.211 | -0.606 | -0.014 |
| P62915 | Gtf2b | 0.008 | 0.054 | 0.170 | 0.000 | -0.046 | -0.454 | -0.911 | -0.672 |
| O35250 | Exoc7 | 0.017 | 0.080 | -0.249 | 0.000 | 0.056 | -0.422 | -0.781 | -0.742 |
| P50172 | Hsd11b1 | 0.261 | 0.477 | 0.205 | 0.000 | -0.348 | -0.044 | -0.433 | -0.522 |
| Q6NZB0 | Dnajc8 | 0.099 | 0.243 | -0.028 | 0.000 | 0.279 | -0.162 | -0.578 | -0.129 |
| Q9CZY3 | Ube2v1 | 0.009 | 0.057 | 0.000 | -0.037 | 0.271 | -0.626 | -1.190 | -0.739 |
| Q9CY57 | Chtop | 0.143 | 0.309 | -0.418 | 0.117 | 0.000 | -0.544 | -0.832 | -0.228 |
| Q9ET26 | Rnf114 | 0.004 | 0.033 | -0.147 | 0.271 | 0.000 | -1.180 | -1.710 | -2.010 |
| P35276 | Rab3d | 0.330 | 0.567 | 0.916 | -0.442 | 0.000 | 0.434 | -2.060 | -0.646 |
| Q91WN1 | Dnajc9 | 0.034 | 0.122 | -0.034 | 0.000 | 0.155 | -0.381 | -0.634 | -0.195 |
| Q7TNG5 | Eml2 | 0.001 | 0.016 | 0.205 | -0.013 | 0.000 | -1.330 | -1.840 | -1.470 |
| Q924K8 | Mta3 | 0.030 | 0.114 | -0.251 | 0.006 | 0.000 | -0.565 | -0.629 | -0.314 |
| Q9QXY6 | Ehd3 | 0.016 | 0.079 | -0.307 | 0.000 | 0.187 | -0.555 | -0.968 | -0.899 |
| Q62167 | Ddx3x | 0.437 | 0.694 | 0.572 | -0.079 | 0.000 | 0.360 | -0.179 | -0.557 |
| Q60996 | Ppp2r5c | 0.122 | 0.280 | 0.000 | -0.686 | 0.064 | -0.519 | -1.030 | -0.730 |
| Q8BJY1 | Psmd5 | 0.005 | 0.039 | 0.000 | -0.013 | 0.093 | -0.877 | -1.520 | -0.988 |
| Q61699 | Hsph1 | 0.043 | 0.141 | 0.078 | -0.155 | 0.000 | -0.310 | -1.080 | -0.742 |
| Q99K41 | Emilin1 | 0.416 | 0.670 | 0.000 | 0.316 | -1.210 | 0.517 | 0.065 | -0.112 |
| P62911 | Rpl32 | 0.562 | 0.831 | 0.000 | 0.444 | -0.453 | 0.516 | -0.354 | 0.583 |
| Q9CWX9 | Ddx47 | 0.062 | 0.178 | 0.000 | -0.105 | 0.012 | -0.508 | -1.220 | -0.386 |
| Q04207 | Rela | 0.003 | 0.030 | 0.000 | -0.006 | 0.032 | -0.840 | -1.440 | -1.170 |
| B2RUP2 | Unc13d | 0.001 | 0.019 | -0.112 | 0.000 | 0.027 | -0.618 | -0.760 | -0.569 |
| Q7TQK1 | Ints7 | 0.079 | 0.211 | -0.115 | 0.068 | 0.000 | -0.097 | -0.432 | -0.299 |
| P21278 | Gna11 | 0.000 | 0.008 | -0.109 | 0.262 | 0.000 | 3.110 | 2.800 | 3.100 |
| Q60605 | Myl6 | 0.043 | 0.141 | -0.374 | 0.009 | 0.000 | 0.591 | 0.152 | 0.570 |
| Q80T69 | Rsbn1 | 0.064 | 0.183 | 0.000 | -0.349 | 0.070 | -0.907 | -0.869 | -0.298 |
| P09450 | Junb | 0.437 | 0.694 | -0.450 | 0.000 | 0.615 | -0.225 | 0.461 | 1.360 |
| Q8VDD9 | Phip | 0.465 | 0.724 | -0.464 | 0.000 | 0.167 | -0.384 | -0.642 | 0.057 |
| Q811D0 | Dlg1 | 0.287 | 0.512 | 0.849 | -0.648 | 0.000 | 0.548 | -2.010 | -1.710 |
| Q8K4B0 | Mta1 | 0.791 | 1.000 | 0.000 | -0.152 | 0.426 | 0.202 | -0.311 | 0.179 |
| P70280 | Vamp7 | 0.346 | 0.587 | -0.078 | 0.033 | 0.000 | 0.104 | -0.400 | -0.234 |
| Q9QZE7 | Tsnax | 0.001 | 0.022 | 0.000 | 0.385 | -0.165 | -1.410 | -1.900 | -1.600 |
| Q8R3N1 | Nop14 | 0.332 | 0.570 | -0.160 | 0.000 | 0.304 | 0.053 | -0.677 | -0.095 |
| Q8BFP9 | Pdk1 | 0.386 | 0.634 | 0.043 | 0.000 | -0.101 | 0.101 | -0.625 | -0.165 |
| Q99P81 | Abcg3 | 0.243 | 0.453 | 0.251 | 0.000 | -0.027 | 0.093 | -0.358 | -0.158 |
| Q9CZ30 | Ola1 | 0.090 | 0.230 | 0.164 | 0.000 | -0.069 | -0.178 | -0.868 | -0.330 |
| Q6A0D4 | Rftn1 | 0.286 | 0.511 | 0.030 | 0.000 | -0.148 | -0.005 | -0.496 | -0.184 |
| Q99PP7 | Trim33 | 0.047 | 0.150 | -0.262 | 0.000 | 0.007 | -0.335 | -0.889 | -0.588 |
| Q99LM2 | Cdk5rap3 | 0.442 | 0.699 | 0.000 | -0.004 | 0.018 | 0.014 | -0.017 | 0.147 |
| Q6P1Y8 | Inpp4b | 0.035 | 0.125 | -0.467 | 0.000 | 0.045 | -1.560 | -3.990 | -2.020 |
| O35604 | Npc1 | 0.052 | 0.160 | 0.406 | -0.105 | 0.000 | 0.918 | 0.427 | 0.686 |
| P25444 | Rps2 | 0.856 | 1.000 | 0.223 | -0.069 | 0.000 | 0.402 | -0.165 | 0.026 |
| Q8R2M2 | Dnttip2 | 0.473 | 0.733 | -0.541 | 0.000 | 0.436 | -0.515 | -0.348 | -0.001 |
| Q3UMU9 | Hdgfrp2 | 0.090 | 0.230 | 0.082 | -0.189 | 0.000 | -0.315 | -2.380 | -1.430 |
| Q9JM76 | Arpc3 | 0.056 | 0.166 | 0.000 | -0.035 | 0.058 | 0.405 | 0.091 | 0.299 |
| Q9EQ20 | Aldh6a1 | 0.368 | 0.615 | 0.300 | 0.000 | -0.150 | 1.460 | -0.061 | 0.226 |
| O55135 | Eif6 | 0.023 | 0.096 | 0.057 | -0.012 | 0.000 | -0.371 | -0.879 | -1.130 |
| Q9Z0S1 | Bpnt1 | 0.290 | 0.515 | -1.300 | 0.000 | 0.199 | -3.920 | -2.080 | 0.204 |
| Q3UYV9 | Ncbp1 | 0.003 | 0.030 | -0.103 | 0.026 | 0.000 | -0.664 | -0.913 | -0.589 |
| Q8R0X7 | Sgpl1 | 0.003 | 0.033 | 0.000 | 0.081 | -0.094 | 1.270 | 0.727 | 1.060 |
| Q8K2M0 | Mrpl38 | 0.002 | 0.024 | 0.114 | 0.000 | -0.089 | 0.995 | 0.671 | 0.855 |
| P16125 | Ldhb | 0.000 | 0.013 | -0.212 | 0.132 | 0.000 | -1.330 | -1.570 | -1.550 |
| Q9DC50 | Crot | 0.119 | 0.276 | 0.000 | -0.124 | 0.034 | -0.050 | -0.367 | -0.340 |
| P62071 | Rras2 | 0.017 | 0.081 | 0.120 | 0.000 | -0.083 | 2.130 | 0.816 | 1.860 |
| Q80XU3 | Nucks1 | 0.047 | 0.149 | 0.000 | -0.362 | 0.201 | -0.682 | -2.200 | -1.300 |
| P23591 | Tsta3 | 0.050 | 0.155 | 0.000 | -0.683 | 0.362 | -0.877 | -1.760 | -2.630 |
| P97300 | Nptn | 0.044 | 0.143 | -0.405 | 0.611 | 0.000 | 0.707 | 1.630 | 1.810 |

| | | | | | | | | | |
|---|---|---|---|---|---|---|---|---|---|
| Q9Z127 | Slc7a5 | 0.005 | 0.043 | 0.399 | 0.000 | -0.183 | 1.980 | 1.200 | 1.790 |
| G5E829 | Atp2b1 | 0.001 | 0.020 | -0.137 | 0.083 | 0.000 | 1.270 | 0.902 | 1.290 |
| O35435 | Dhodh | 0.072 | 0.196 | 0.000 | 0.359 | -1.330 | 0.638 | 1.240 | 1.160 |
| Q9D7X8 | Ggct | 0.001 | 0.019 | -0.269 | 0.000 | 0.357 | -2.090 | -1.980 | -2.600 |
| O35691 | Pnn | 0.019 | 0.085 | -0.156 | 0.000 | 0.067 | -0.334 | -0.570 | -0.348 |
| Q91YU8 | Ppan | 0.916 | 1.000 | -0.220 | 0.149 | 0.000 | 0.107 | -0.314 | 0.077 |
| Q9QXG4 | Acss2 | 0.016 | 0.078 | -0.138 | 0.694 | 0.000 | -1.240 | -2.430 | -1.330 |
| P40237 | Cd82 | 0.269 | 0.488 | -0.180 | 0.212 | 0.000 | -0.365 | -0.262 | 0.030 |
| Q922J3 | Clip1 | 0.034 | 0.122 | 0.415 | -0.534 | 0.000 | -1.450 | -1.440 | -0.712 |
| Q6P5B0 | Rrp12 | 0.301 | 0.529 | 0.369 | -0.321 | 0.000 | 0.174 | -0.805 | -0.596 |
| P21995 | Emb | 0.102 | 0.247 | -0.018 | 0.022 | 0.000 | 0.401 | 0.018 | 0.398 |
| Q6P9J9 | Ano6 | 0.002 | 0.023 | 0.138 | 0.000 | -0.273 | 1.220 | 0.975 | 1.320 |
| Q8BUM3 | Ptpn7 | 0.004 | 0.038 | 0.125 | -0.871 | 0.000 | -2.120 | -2.240 | -1.970 |
| P50518 | Atp6v1e1 | 0.837 | 1.000 | 0.431 | 0.000 | -0.221 | 0.368 | -0.169 | 0.175 |
| Q8BYW1 | Arhgap25 | 0.003 | 0.033 | -0.351 | 0.267 | 0.000 | -1.140 | -1.230 | -1.400 |
| O88441 | Mtx2 | 0.276 | 0.496 | 0.000 | 0.113 | -0.108 | 0.703 | -0.135 | 0.393 |
| Q6P9Q4 | Fhod1 | 0.095 | 0.238 | 0.000 | 0.192 | -0.114 | -0.066 | -0.781 | -0.583 |
| Q9R0I7 | Ylpm1 | 0.058 | 0.171 | 0.000 | 0.437 | -0.223 | -0.318 | -0.631 | -0.534 |
| Q921N6 | Ddx27 | 0.244 | 0.454 | 0.486 | 0.000 | -0.037 | 0.216 | -0.895 | -0.357 |
| Q9Z1Z0 | Uso1 | 0.335 | 0.573 | 0.000 | 0.009 | -0.654 | 0.355 | -2.660 | -1.290 |
| Q8R127 | Sccpdh | 0.064 | 0.183 | -0.667 | 0.216 | 0.000 | 0.428 | 0.527 | 0.710 |
| B1AZI6 | Thoc2 | 0.998 | 1.000 | 0.453 | -0.343 | 0.000 | -0.050 | -0.330 | 0.488 |
| Q18PI6 | Cd84 | 0.101 | 0.247 | -0.282 | 0.224 | 0.000 | -0.350 | -1.080 | -0.397 |
| Q8BY87 | Usp47 | 0.774 | 1.000 | -1.190 | 0.000 | 0.428 | -0.271 | -1.430 | 0.297 |
| Q8BJ03 | Cox15 | 0.015 | 0.075 | 0.000 | -0.638 | 0.346 | 1.240 | 1.100 | 1.810 |
| Q8R164 | Bphl | 0.031 | 0.114 | 0.197 | 0.000 | -0.133 | 0.714 | 0.305 | 0.778 |
| Q3THG9 | Aarsd1 | 0.001 | 0.019 | -0.043 | 0.000 | 0.016 | -0.899 | -1.240 | -1.330 |
| Q920E5 | Fdps | 0.104 | 0.252 | 1.020 | 0.000 | -0.336 | 1.880 | 0.878 | 1.120 |
| Q9JII5 | Dazap1 | 0.028 | 0.107 | 0.000 | 0.377 | -0.062 | -0.451 | -0.655 | -0.308 |
| Q8K003 | Tma7 | 0.203 | 0.399 | 0.000 | -0.024 | 0.550 | 0.396 | 0.384 | 0.845 |
| P28656 | Nap1l1 | 0.113 | 0.267 | 0.177 | -0.254 | 0.000 | -0.193 | -0.671 | -0.353 |
| Q9DAW6 | Prpf4 | 0.005 | 0.039 | 0.000 | 0.312 | -0.251 | -0.873 | -0.940 | -1.050 |
| Q3TW96 | Uap1l1 | 0.784 | 1.000 | -0.947 | 0.000 | 0.254 | 0.223 | -0.282 | -0.282 |
| Q62383 | Supt6h | 0.838 | 1.000 | 0.314 | -0.950 | 0.000 | 0.804 | 0.583 | -1.490 |
| O09130 | Nfatc2ip | 0.031 | 0.115 | 0.000 | -0.134 | 0.049 | -0.815 | -2.260 | -1.240 |
| Q8VDM4 | Psmd2 | 0.239 | 0.447 | 0.042 | -0.316 | 0.000 | -0.103 | -0.642 | -0.329 |
| Q9DBF1 | Aldh7a1 | 0.000 | 0.009 | 0.000 | 0.318 | -0.191 | 2.700 | 2.460 | 2.520 |
| Q99MR8 | Mccc1 | 0.382 | 0.631 | 0.014 | 0.000 | -0.317 | 0.811 | -0.251 | 0.108 |
| Q6ZQB6 | Ppip5k2 | 0.003 | 0.033 | 0.000 | 0.042 | -0.056 | -0.461 | -0.578 | -0.345 |
| P28063 | Psmb8 | 0.001 | 0.014 | -0.160 | 0.000 | 0.024 | -1.350 | -1.790 | -1.440 |
| P11438 | Lamp1 | 0.050 | 0.156 | 0.166 | -0.304 | 0.000 | 0.879 | 0.236 | 0.715 |
| Q60790 | Rasa3 | 0.846 | 1.000 | 0.000 | 0.108 | -0.282 | 0.257 | -0.193 | -0.123 |
| Q9CY97 | Ssu72 | 0.032 | 0.118 | -0.131 | 0.000 | 0.023 | -0.703 | -1.630 | -0.740 |
| Q925I1 | Atad3 | 0.245 | 0.456 | 0.000 | 0.064 | -0.526 | 0.689 | -0.048 | 0.094 |
| Q8R149 | Bud13 | 0.404 | 0.655 | 0.211 | -0.128 | 0.000 | 0.438 | -0.129 | -5.790 |
| Q8BJ05 | Zc3h14 | 0.250 | 0.462 | 0.000 | -0.321 | 0.346 | -1.400 | -0.546 | 0.077 |
| Q8BG05 | Hnrnpa3 | 0.572 | 0.842 | 0.000 | 0.531 | -0.203 | -0.478 | -0.064 | 0.294 |
| Q9WVL0 | Gstz1 | 0.112 | 0.265 | 0.000 | 0.163 | -0.024 | 1.190 | 0.487 | 0.239 |
| Q9DB25 | Alg5 | 0.009 | 0.058 | 0.000 | -0.448 | 0.027 | 0.870 | 0.529 | 0.843 |
| O55143 | Atp2a2 | 0.001 | 0.018 | 0.000 | 0.045 | -0.311 | 1.460 | 1.100 | 1.270 |
| Q6QD59 | Bnip1 | 0.831 | 1.000 | 0.000 | -0.712 | 0.039 | -0.271 | -0.640 | 0.026 |
| P56376 | Acyp1 | 0.087 | 0.224 | 0.000 | -0.217 | 0.506 | -0.816 | -1.530 | -0.252 |
| P58771 | Tpm1 | 0.375 | 0.623 | 0.000 | 0.208 | -0.022 | 0.058 | -0.868 | 0.051 |
| Q7TNC4 | Luc7l2 | 0.063 | 0.180 | 0.454 | -0.011 | 0.000 | -0.152 | -1.330 | -1.050 |
| Q922H2 | Pdk3 | 0.193 | 0.385 | 0.065 | 0.000 | -0.219 | 0.781 | -0.001 | 0.229 |
| Q9CQJ6 | Denr | 0.330 | 0.568 | 0.249 | -0.263 | 0.000 | 0.475 | -0.043 | 0.254 |
| Q00612 | G6pdx | 0.184 | 0.373 | 1.060 | -0.237 | 0.000 | 0.116 | -0.895 | -0.924 |
| P29452 | Casp1 | 0.002 | 0.027 | -0.321 | 0.027 | 0.000 | -1.120 | -1.610 | -1.380 |
| O54957 | Lat | 0.507 | 0.771 | -0.255 | 0.567 | 0.000 | 0.081 | 0.134 | 0.862 |

| | | | | | | | | | |
|---|---|---|---|---|---|---|---|---|---|
| Q05117 | Acp5 | 0.229 | 0.433 | 0.477 | 0.000 | -0.821 | -0.165 | -0.959 | -1.720 |
| Q3TZM9 | Alg11 | 0.001 | 0.021 | 0.265 | 0.000 | -0.147 | 1.390 | 1.080 | 1.260 |
| Q6ZQ03 | Fnbp4 | 0.001 | 0.017 | 0.000 | -0.040 | 0.153 | -0.833 | -1.160 | -1.090 |
| Q99J62 | Rfc4 | 0.246 | 0.457 | 0.532 | -0.054 | 0.000 | 0.414 | -0.865 | -1.300 |
| Q9CQU3 | Rer1 | 0.001 | 0.015 | 0.049 | 0.000 | -0.033 | 0.920 | 0.802 | 1.120 |
| P56812 | Pdcd5 | 0.007 | 0.049 | -0.173 | 0.137 | 0.000 | -1.160 | -1.590 | -0.851 |
| Q5SVQ0 | Kat7 | 0.002 | 0.025 | -0.004 | 0.000 | 0.180 | -0.725 | -1.080 | -0.767 |
| Q8BGT7 | Smndc1 | 0.168 | 0.349 | -0.203 | 0.000 | 0.691 | -1.140 | -1.620 | 0.190 |
| Q08481 | Pecam1 | 0.631 | 0.907 | 0.000 | 1.060 | -0.637 | -0.841 | 0.355 | -0.038 |
| Q4VAA2 | Cdv3 | 0.723 | 1.000 | 0.000 | -0.140 | 0.131 | 0.438 | -0.344 | 0.172 |
| P61087 | Ube2k | 0.014 | 0.072 | 0.193 | 0.000 | -0.032 | -0.552 | -1.380 | -1.130 |
| P59114 | Pcif1 | 0.284 | 0.508 | 0.000 | -0.966 | 0.255 | -0.704 | -2.550 | -0.344 |
| Q6PAM1 | Txlna | 0.425 | 0.679 | 0.512 | -0.234 | 0.000 | 1.380 | 0.111 | 0.072 |
| Q9EPL8 | Ipo7 | 0.513 | 0.776 | 0.430 | -0.700 | 0.000 | 0.125 | -1.260 | -0.270 |
| Q9JMA2 | Qtrt1 | 0.001 | 0.019 | 0.000 | 0.059 | -0.158 | -0.700 | -0.758 | -0.891 |
| Q640N3 | Arhgap30 | 0.048 | 0.151 | -0.074 | 0.000 | 0.135 | -0.461 | -1.900 | -1.130 |
| P41241 | Csk | 0.102 | 0.248 | 0.002 | 0.000 | -0.016 | -0.072 | -0.837 | -0.511 |
| Q3UBX0 | Tmem109 | 0.186 | 0.376 | 0.000 | -4.730 | 0.023 | 0.839 | 0.983 | 1.030 |
| Q8R146 | Apeh | 0.003 | 0.031 | -0.048 | 0.000 | 0.071 | -1.000 | -1.700 | -1.640 |
| Q8R0L9 | Tada3 | 0.199 | 0.394 | -0.089 | 0.639 | 0.000 | -0.084 | -0.212 | -0.232 |
| Q6P9N1 | Fam126a | 0.432 | 0.687 | -0.621 | 2.050 | 0.000 | 0.332 | 1.680 | 1.890 |
| Q8VCX5 | Micu1 | 0.567 | 0.836 | -0.165 | 0.177 | 0.000 | 1.300 | -0.698 | 0.513 |
| Q99JW4 | Lims1 | 0.478 | 0.740 | 0.000 | 0.120 | -0.598 | 0.349 | -0.126 | -0.073 |
| Q6ZWR6 | Syne1 | 0.727 | 1.000 | 0.859 | -1.990 | 0.000 | 0.386 | 0.188 | -0.688 |
| Q99M87 | Dnaja3 | 0.029 | 0.112 | 0.003 | 0.000 | -0.329 | 0.713 | 0.216 | 0.625 |
| Q80SU7 | Gvin1 | 0.117 | 0.273 | 0.231 | 0.000 | -0.019 | 0.045 | -0.881 | -0.733 |
| B2RY04 | Dock5 | 0.009 | 0.057 | -0.206 | 0.037 | 0.000 | -0.766 | -1.150 | -0.647 |
| Q5SUF2 | Luc7l3 | 0.021 | 0.091 | 0.264 | 0.000 | -0.087 | -0.609 | -1.770 | -1.470 |
| Q09200 | B4galnt1 | 0.004 | 0.036 | 0.052 | -0.019 | 0.000 | -0.568 | -0.695 | -0.380 |
| Q06138 | Cab39 | 0.004 | 0.036 | 0.105 | -0.146 | 0.000 | -0.761 | -1.280 | -1.010 |
| Q9D379 | Ephx1 | 0.000 | 0.012 | 0.114 | -0.040 | 0.000 | 1.700 | 1.320 | 1.580 |
| Q9JIF7 | Copb1 | 0.018 | 0.084 | 0.000 | 0.412 | -0.073 | -0.488 | -1.040 | -1.230 |
| Q9ESE1 | Lrba | 0.000 | 0.013 | -0.292 | 0.080 | 0.000 | -1.440 | -1.500 | -1.340 |
| Q9CQ40 | Mrpl49 | 0.092 | 0.233 | 0.339 | 0.000 | -0.837 | 0.637 | 0.883 | 0.445 |
| O08573 | Lgals9 | 0.391 | 0.640 | 0.475 | 0.000 | -0.125 | 0.251 | -0.413 | -0.271 |
| Q6PR54 | Rif1 | 0.046 | 0.148 | 0.000 | -0.091 | 0.018 | -0.254 | -0.829 | -0.460 |
| Q9ESV0 | Ddx24 | 0.175 | 0.360 | 0.000 | 0.189 | -0.128 | 0.133 | -1.170 | -0.925 |
| Q8CIM8 | Ints4 | 0.007 | 0.048 | -0.070 | 0.095 | 0.000 | -0.300 | -0.542 | -0.502 |
| Q6P6J9 | Txndc15 | 0.896 | 1.000 | 0.417 | 0.000 | -1.790 | -0.136 | -1.240 | 0.342 |
| Q8VBW6 | Nae1 | 0.028 | 0.108 | -0.019 | 0.000 | 0.206 | -0.776 | -2.140 | -1.120 |
| Q3TGF2 | Fam107b | 0.166 | 0.347 | 0.000 | -0.310 | 0.205 | -0.458 | -0.714 | -0.106 |
| P50171 | Hsd17b8 | 0.804 | 1.000 | -0.055 | 0.000 | 0.133 | 0.345 | -0.437 | -0.015 |
| Q68FL4 | Ahcyl2 | 0.099 | 0.244 | 0.091 | -0.030 | 0.000 | -0.016 | -0.637 | -0.577 |
| Q60960 | Kpna1 | 0.132 | 0.296 | 0.000 | 0.990 | -0.334 | -1.180 | -0.338 | -0.508 |
| Q6PD26 | Pigs | 0.001 | 0.020 | 0.000 | 0.161 | -0.055 | 0.792 | 0.665 | 0.633 |
| P62267 | Rps23 | 0.689 | 0.965 | 0.013 | -0.028 | 0.000 | 0.271 | -0.356 | 0.363 |
| P16675 | Ctsa | 0.839 | 1.000 | 0.308 | 0.000 | -0.004 | 1.220 | -0.818 | -0.515 |
| Q8R3F5 | Mcat | 0.181 | 0.368 | 0.020 | 0.000 | -0.178 | 0.816 | -0.052 | 0.333 |
| Q9DCN2 | Cyb5r3 | 0.000 | 0.005 | 0.000 | 0.161 | -0.051 | 2.080 | 1.950 | 2.120 |
| O08550 | Kmt2b | 0.024 | 0.100 | 0.000 | -0.032 | 0.324 | -0.352 | -0.570 | -0.287 |
| Q9DB15 | Mrpl12 | 0.006 | 0.044 | 0.000 | 0.110 | -0.240 | 0.803 | 0.551 | 0.609 |
| Q921L3 | Tmco1 | 0.013 | 0.070 | 0.210 | 0.000 | -0.647 | 1.070 | 1.350 | 0.878 |
| Q99NB1 | Acss1 | 0.101 | 0.247 | 0.000 | 0.064 | -0.261 | -0.149 | -0.573 | -0.593 |
| Q9EQI8 | Mrpl46 | 0.017 | 0.081 | 0.194 | 0.000 | -0.083 | 1.000 | 0.471 | 0.691 |
| Q8R574 | Prpsap2 | 0.015 | 0.076 | 0.000 | -0.221 | 0.127 | -0.692 | -1.420 | -0.898 |
| O35639 | Anxa3 | 0.011 | 0.063 | 0.503 | 0.000 | -0.324 | 1.340 | 1.040 | 1.270 |
| Q9WVJ3 | Cpq | 0.146 | 0.315 | 0.375 | 0.000 | -0.250 | -0.094 | -0.745 | -0.448 |
| Q68FH4 | Galk2 | 0.009 | 0.056 | 0.000 | 0.100 | -0.032 | -0.657 | -1.280 | -0.781 |
| Q8C163 | Exog | 0.071 | 0.194 | -0.225 | 0.654 | 0.000 | 0.703 | 0.765 | 1.070 |

| | | | | | | | | | |
|---|---|---|---|---|---|---|---|---|---|
| O35609 | Scamp3 | 0.623 | 0.898 | -0.255 | 0.000 | 0.047 | -0.117 | -0.199 | -0.056 |
| P03930 | Mtatp8 | 0.027 | 0.105 | -0.070 | 0.024 | 0.000 | 0.738 | 0.252 | 0.830 |
| P62270 | Rps18 | 0.436 | 0.693 | 0.154 | -0.073 | 0.000 | 0.130 | -0.385 | -0.087 |
| Q80UM7 | Mogs | 0.021 | 0.090 | 0.000 | 0.122 | -0.001 | 0.478 | 0.248 | 0.578 |
| Q8C1B7 | Sept 11 | 0.137 | 0.302 | -0.140 | 0.191 | 0.000 | 0.444 | 0.081 | 0.326 |
| Q80X82 | Sympk | 0.294 | 0.522 | 0.000 | 1.070 | -0.036 | -0.491 | -0.519 | 0.363 |
| Q5U458 | Dnajc11 | 0.066 | 0.185 | 0.771 | -0.042 | 0.000 | 1.780 | 1.580 | 0.657 |
| P97311 | Mcm6 | 0.032 | 0.116 | 0.272 | -0.017 | 0.000 | -0.280 | -1.070 | -0.927 |
| Q8VDJ3 | Hdlbp | 0.079 | 0.209 | 0.211 | 0.000 | -0.388 | 1.260 | 0.312 | 0.588 |
| Q61292 | Lamb2 | 0.013 | 0.070 | 0.471 | -3.200 | 0.000 | 4.150 | 3.830 | 4.190 |
| O08600 | Endog | 0.058 | 0.171 | -0.172 | 0.406 | 0.000 | -0.439 | -0.313 | -0.402 |
| Q9ER00 | Stx12 | 0.134 | 0.298 | 0.205 | 0.000 | -1.040 | 1.720 | -0.121 | 1.470 |
| Q61263 | Soat1 | 0.202 | 0.397 | -1.560 | 0.617 | 0.000 | 0.381 | 0.756 | 1.000 |
| P08030 | Aprt | 0.013 | 0.071 | 0.398 | -0.078 | 0.000 | -0.482 | -0.881 | -0.702 |
| O35459 | Ech1 | 0.045 | 0.146 | 0.000 | 0.120 | -0.262 | 0.291 | 0.279 | 0.264 |
| P47968 | Rpia | 0.160 | 0.337 | 0.718 | -0.577 | 0.000 | -0.741 | -2.220 | -0.389 |
| Q91XD7 | Creld1 | 0.012 | 0.067 | 0.000 | 0.730 | -0.301 | 1.590 | 1.500 | 2.140 |
| Q6ZPQ6 | Pitpnm2 | 0.007 | 0.048 | -0.296 | 0.000 | 0.192 | -0.918 | -1.170 | -0.791 |
| Q924C1 | Xpo5 | 0.198 | 0.392 | 0.232 | 0.000 | -0.052 | 0.150 | -1.420 | -0.684 |
| Q8VDC0 | Lars2 | 0.390 | 0.640 | 0.288 | 0.000 | -4.090 | 0.945 | -0.292 | -0.213 |
| Q8CEC6 | Ppwd1 | 0.002 | 0.025 | 0.000 | 0.104 | -0.053 | -0.865 | -0.835 | -0.565 |
| Q8K021 | Scamp1 | 0.737 | 1.000 | 0.000 | 0.045 | -0.281 | -0.544 | -0.201 | 1.040 |
| P62908 | Rps3 | 0.186 | 0.376 | 0.099 | 0.000 | -0.178 | -0.074 | -0.632 | -0.252 |
| Q497V5 | Srbd1 | 0.629 | 0.904 | 0.000 | -0.755 | 0.097 | -0.082 | -0.855 | -0.278 |
| P34022 | Ranbp1 | 0.071 | 0.195 | 0.406 | -0.097 | 0.000 | -0.243 | -1.450 | -0.795 |
| Q9CRA8 | Exosc5 | 0.006 | 0.046 | -0.019 | 0.000 | 0.245 | -0.631 | -0.827 | -0.461 |
| Q8K2X3 | Obfc1 | 0.025 | 0.101 | -0.192 | 0.301 | 0.000 | -0.567 | -1.010 | -0.532 |
| Q3UHB1 | Nt5dc3 | 0.001 | 0.014 | 0.000 | 0.448 | -0.077 | 2.160 | 2.160 | 2.560 |
| Q9ESU6 | Brd4 | 0.610 | 0.884 | 0.000 | 0.102 | -0.420 | -0.365 | -0.366 | 0.059 |
| Q60775 | Elf1 | 0.189 | 0.380 | -0.298 | 0.000 | 0.179 | -0.917 | -2.490 | -0.101 |
| Q9JJK2 | Lancl2 | 0.097 | 0.240 | 0.477 | -0.052 | 0.000 | -0.074 | -0.884 | -0.483 |
| Q80YW0 | Cyth4 | 0.013 | 0.070 | 0.000 | -0.147 | 0.123 | -0.448 | -0.948 | -0.724 |
| Q9CWW7 | Cxxc1 | 0.381 | 0.630 | -0.019 | 0.834 | 0.000 | -0.213 | 0.075 | 0.075 |
| Q9DCJ5 | Ndufa8 | 0.075 | 0.202 | 0.000 | 1.300 | -0.142 | 1.220 | 1.620 | 2.010 |
| O08529 | Capn2 | 0.746 | 1.000 | -0.207 | 0.000 | 0.072 | 0.305 | -0.569 | -0.147 |
| Q9D020 | Nt5c3a | 0.008 | 0.053 | 0.000 | 0.443 | -0.151 | -0.917 | -1.350 | -0.880 |
| Q62393 | Tpd52 | 0.002 | 0.027 | 0.298 | -0.017 | 0.000 | -0.642 | -0.624 | -0.600 |
| Q9Z0H4 | Celf2 | 0.075 | 0.202 | -0.319 | 0.000 | 0.094 | -0.466 | -0.487 | -0.283 |
| Q8VDP3 | Mical1 | 0.105 | 0.252 | -0.304 | 0.256 | 0.000 | -0.186 | -1.030 | -1.600 |
| Q921G7 | Etfdh | 0.009 | 0.055 | 0.011 | 0.000 | -0.217 | 0.785 | 0.394 | 0.573 |
| Q9JHS4 | Clpx | 0.557 | 0.825 | 0.335 | 0.000 | -0.328 | 1.540 | -0.635 | 0.363 |
| Q8BK72 | Mrps27 | 0.012 | 0.068 | 0.000 | 0.528 | -0.256 | 1.200 | 1.050 | 1.510 |
| O70475 | Ugdh | 0.023 | 0.096 | 0.529 | 0.000 | -0.191 | 2.030 | 0.941 | 1.470 |
| Q93092 | Taldo1 | 0.006 | 0.044 | 0.013 | 0.000 | -0.200 | -1.010 | -1.870 | -1.490 |
| O70293 | Grk6 | 0.047 | 0.150 | 0.170 | -0.136 | 0.000 | -0.401 | -1.250 | -0.613 |
| Q99LH1 | Gnl2 | 0.400 | 0.651 | -0.012 | 0.347 | 0.000 | 0.181 | 0.019 | -0.682 |
| Q03347 | Runx1 | 0.085 | 0.221 | -0.073 | 0.000 | 0.462 | -0.343 | -0.360 | -0.146 |
| Q9DBR1 | Xrn2 | 0.619 | 0.893 | 0.182 | -3.030 | 0.000 | -0.079 | -0.512 | -0.558 |
| Q8BVQ5 | Ppme1 | 0.046 | 0.149 | 0.000 | -0.304 | 0.094 | -0.600 | -2.320 | -1.730 |
| P58389 | Ppp2r4 | 0.004 | 0.035 | 0.000 | 0.159 | -0.258 | -0.987 | -1.580 | -1.380 |
| Q9D7S7 | Rpl22l1 | 0.056 | 0.168 | 0.639 | -0.126 | 0.000 | 1.190 | 0.683 | 0.870 |
| Q9CQI3 | Gmfb | 0.027 | 0.105 | -0.614 | 0.344 | 0.000 | -1.170 | -1.820 | -1.060 |
| Q6P8X1 | Snx6 | 0.080 | 0.211 | 0.179 | -0.169 | 0.000 | -0.355 | -1.340 | -0.535 |
| Q922S4 | Pde2a | 0.073 | 0.199 | 0.280 | -0.058 | 0.000 | -0.909 | -2.760 | -0.790 |
| Q9Z2W0 | Dnpep | 0.888 | 1.000 | -0.137 | 0.062 | 0.000 | 0.083 | -0.199 | 0.092 |
| Q6ZPR5 | Smpd4 | 0.671 | 0.948 | -0.120 | 0.000 | 0.029 | 0.054 | -0.374 | 0.028 |
| Q8R0F5 | Rbmx2 | 0.033 | 0.119 | 0.125 | 0.000 | -0.033 | -0.510 | -1.190 | -0.479 |
| Q62186 | Ssr4 | 0.004 | 0.036 | 0.000 | 0.264 | -0.140 | 1.460 | 1.390 | 2.160 |
| Q8BFZ3 | Actbl2 | 0.527 | 0.794 | 0.000 | -1.890 | 1.230 | -0.488 | 0.989 | 0.984 |

| | | | | | | | | | |
|---|---|---|---|---|---|---|---|---|---|
| P48962 | Slc25a4 | 0.000 | 0.013 | 0.000 | 0.359 | -0.136 | 2.120 | 1.860 | 2.070 |
| P14234 | Fgr | 0.214 | 0.412 | 0.781 | 0.000 | -0.781 | -0.032 | -1.290 | -1.570 |
| Q61805 | Lbp | 0.929 | 1.000 | 1.500 | 0.000 | -0.715 | 0.961 | -0.240 | -0.151 |
| Q8K3H0 | Appl1 | 0.122 | 0.280 | -0.019 | 0.000 | 0.087 | 0.005 | -0.899 | -0.625 |
| Q9CQU0 | Txndc12 | 0.033 | 0.119 | 0.153 | -0.268 | 0.000 | 0.484 | 0.520 | 1.080 |
| Q9D2H6 | Sp2 | 0.814 | 1.000 | -0.748 | 0.000 | 0.314 | -0.304 | -0.704 | 0.256 |
| Q9Z183 | Padi4 | 0.348 | 0.589 | 0.465 | -0.149 | 0.000 | 0.230 | -0.725 | -0.248 |
| Q6ZPZ3 | Zc3h4 | 0.051 | 0.157 | 0.058 | -0.071 | 0.000 | -0.559 | -1.210 | -0.369 |
| Q9CQX2 | Cyb5b | 0.013 | 0.071 | -0.164 | 0.064 | 0.000 | 0.975 | 0.423 | 0.947 |
| P30412 | Ppic | 0.000 | 0.004 | 0.211 | 0.000 | -0.045 | 4.510 | 4.650 | 5.030 |
| Q8R1V4 | Tmed4 | 0.003 | 0.033 | 0.009 | 0.000 | -0.106 | 0.740 | 0.516 | 0.912 |
| Q02257 | Jup | 0.002 | 0.027 | -0.066 | 0.000 | 0.472 | 1.290 | 1.510 | 1.590 |
| Q8CG46 | Smc5 | 0.067 | 0.188 | 0.000 | -0.111 | 0.174 | -0.168 | -0.552 | -0.273 |
| P42232 | Stat5b | 0.007 | 0.048 | -0.226 | 0.000 | 0.150 | -0.984 | -1.490 | -0.914 |
| P55200 | Kmt2a | 0.043 | 0.142 | -0.334 | 0.008 | 0.000 | -0.582 | -1.070 | -0.506 |
| Q91W98 | Slc15a4 | 0.614 | 0.889 | 0.885 | -1.730 | 0.000 | 0.497 | -0.217 | 0.175 |
| Q9CPQ3 | Tomm22 | 0.010 | 0.061 | -0.029 | 0.000 | 0.186 | 1.040 | 0.523 | 0.946 |
| Q8C147 | Dock8 | 0.007 | 0.050 | -0.108 | 0.071 | 0.000 | -0.609 | -1.200 | -0.943 |
| Q9R1T2 | Sae1 | 0.001 | 0.022 | -0.100 | 0.000 | 0.086 | -1.310 | -1.900 | -1.410 |
| Q8BFQ4 | Wdr82 | 0.490 | 0.754 | -0.084 | 1.340 | 0.000 | -0.615 | -2.410 | 1.500 |
| P70349 | Hint1 | 0.003 | 0.030 | -0.064 | 0.000 | 0.148 | -1.010 | -1.670 | -1.260 |
| Q9D2M8 | Ube2v2 | 0.013 | 0.069 | -0.093 | 0.000 | 0.331 | -0.566 | -1.180 | -0.850 |
| O70318 | Epb41l2 | 0.014 | 0.073 | 0.087 | -0.042 | 0.000 | 1.630 | 0.720 | 1.050 |
| P70335 | Rock1 | 0.917 | 1.000 | -0.464 | 0.059 | 0.000 | 0.136 | -0.564 | 0.117 |
| Q8BWY3 | Etf1 | 0.036 | 0.125 | 0.100 | -0.068 | 0.000 | -0.716 | -1.340 | -0.441 |
| Q01405 | Sec23a | 0.212 | 0.409 | 0.000 | -0.038 | 0.008 | 0.653 | -0.087 | 0.364 |
| Q9D287 | Bcas2 | 0.044 | 0.145 | -0.673 | 0.000 | 0.145 | -1.230 | -1.000 | -0.769 |
| P70224 | Gimap1 | 0.037 | 0.127 | -0.253 | 0.025 | 0.000 | -0.508 | -0.642 | -0.305 |
| Q6ZQ08 | Cnot1 | 0.189 | 0.380 | 0.131 | 0.000 | -0.006 | 0.096 | -0.868 | -0.442 |
| Q9D1M0 | Sec13 | 0.774 | 1.000 | 0.000 | 0.076 | -0.324 | -0.229 | 0.354 | -0.169 |
| P18155 | Mthfd2 | 0.034 | 0.122 | 0.000 | 0.357 | -0.941 | 0.795 | 1.290 | 1.380 |
| O70546 | Kdm6a | 0.111 | 0.263 | 0.014 | -0.113 | 0.000 | -0.286 | -0.636 | -0.129 |
| P70697 | Urod | 0.023 | 0.096 | 0.306 | 0.000 | -0.652 | -1.230 | -2.010 | -1.240 |
| Q9QYJ3 | Dnajb1 | 0.023 | 0.096 | 0.060 | 0.000 | -0.117 | -0.733 | -1.600 | -0.788 |
| P26323 | Fli1 | 0.274 | 0.494 | 0.000 | -0.113 | 0.253 | 0.070 | -0.670 | -0.183 |
| Q9DCZ4 | Apoo | 0.004 | 0.037 | 0.000 | 0.124 | -0.197 | 0.733 | 0.566 | 0.860 |
| P15920 | Atp6v0a2 | 0.003 | 0.033 | 0.000 | 0.059 | -0.066 | 0.643 | 0.434 | 0.744 |
| Q61466 | Smarcd1 | 0.126 | 0.287 | -0.202 | 0.000 | 0.061 | -0.260 | -0.833 | -0.252 |
| Q8BSQ9 | Pbrm1 | 0.031 | 0.114 | 0.000 | 0.231 | -0.058 | -0.850 | -1.020 | -0.291 |
| Q9QVP9 | Ptk2b | 0.038 | 0.131 | -1.180 | 0.133 | 0.000 | -1.480 | -1.840 | -1.650 |
| P62996 | Tra2b | 0.066 | 0.186 | 0.283 | -0.028 | 0.000 | -0.167 | -0.937 | -1.350 |
| O88890 | Sh2d1a | 0.185 | 0.374 | -0.831 | 0.461 | 0.000 | -2.030 | -0.784 | -0.473 |
| Q8CHI8 | Ep400 | 0.615 | 0.889 | -2.080 | 0.000 | 0.300 | -0.174 | -0.489 | 0.144 |
| Q8K2H4 | Acap1 | 0.037 | 0.129 | -0.174 | 0.000 | 0.030 | -0.286 | -0.504 | -0.266 |
| Q8BWU5 | Osgep | 0.031 | 0.114 | 0.000 | 0.021 | -0.337 | -1.120 | -2.230 | -0.990 |
| Q9DCM0 | Ethe1 | 0.636 | 0.912 | 0.000 | 0.137 | -0.820 | 0.135 | -0.445 | 0.180 |
| Q99N93 | Mrpl16 | 0.172 | 0.355 | 0.000 | 0.020 | -0.274 | 0.737 | -0.068 | 0.326 |
| Q91WQ5 | Taf5l | 0.070 | 0.193 | -0.572 | 0.072 | 0.000 | -0.475 | -1.060 | -0.998 |
| Q78XF5 | Ostc | 0.000 | 0.012 | 0.000 | 0.306 | -0.226 | 2.330 | 2.190 | 2.510 |
| P62897 | Cycs | 0.205 | 0.401 | 0.000 | 0.081 | -0.533 | -0.300 | -0.469 | -0.708 |
| Q9CXE7 | Tmed5 | 0.012 | 0.068 | 0.000 | 0.150 | -0.065 | 0.610 | 0.334 | 0.624 |
| Q8BWG8 | Arrb1 | 0.003 | 0.032 | 0.094 | 0.000 | -0.186 | -0.771 | -1.060 | -0.747 |
| Q6NVG1 | Lpcat4 | 0.557 | 0.825 | 1.640 | -0.603 | 0.000 | 0.733 | -2.380 | 0.392 |
| P97352 | S100a13 | 0.076 | 0.205 | 0.121 | -0.269 | 0.000 | -0.471 | -1.150 | -0.410 |
| Q61990 | Pcbp2 | 0.167 | 0.348 | 0.037 | 0.000 | -0.006 | -0.016 | -0.517 | -0.184 |
| O35598 | Adam10 | 0.952 | 1.000 | 0.337 | 0.000 | -0.128 | 0.350 | -0.150 | 0.048 |
| P54729 | Nub1 | 0.979 | 1.000 | 0.840 | 0.000 | -0.638 | 1.160 | 0.303 | -1.330 |
| P84089 | Erh | 0.027 | 0.107 | -0.352 | 0.000 | 0.217 | -1.060 | -2.580 | -1.450 |
| Q8BQ47 | Cnpy4 | 0.431 | 0.687 | -0.487 | 0.000 | 0.258 | -1.090 | -0.890 | 0.397 |

| | | | | | | | | | |
|---|---|---|---|---|---|---|---|---|---|
| P13379 | Cd5 | 0.724 | 1.000 | 0.000 | 0.432 | -2.100 | -0.529 | -0.610 | 0.443 |
| P31786 | Dbi | 0.001 | 0.019 | 0.000 | 0.100 | -0.167 | -1.290 | -1.720 | -1.290 |
| P16045 | Lgals1 | 0.944 | 1.000 | 0.000 | 1.160 | -0.293 | 0.381 | 0.217 | 0.367 |
| Q9WUK2 | Eif4h | 0.613 | 0.888 | 0.000 | -0.268 | 0.057 | -0.012 | -0.659 | 0.052 |
| P97821 | Ctsc | 0.036 | 0.126 | 0.633 | -0.023 | 0.000 | -0.313 | -1.010 | -0.904 |
| P63325 | Rps10 | 0.296 | 0.524 | 0.877 | 0.000 | -0.932 | 1.130 | 0.005 | 1.120 |
| Q9R045 | Angptl2 | 0.066 | 0.186 | 0.000 | 1.160 | -3.870 | 3.180 | 2.590 | 3.020 |
| Q91WM3 | Rrp9 | 0.048 | 0.152 | 0.000 | -0.070 | 0.006 | -0.208 | -0.689 | -0.376 |
| P97370 | Atp1b3 | 0.009 | 0.056 | 0.000 | 0.096 | -0.048 | 0.414 | 0.419 | 0.703 |
| P62751 | Rpl23a | 0.082 | 0.216 | 0.090 | -0.198 | 0.000 | 0.425 | 0.101 | 0.242 |
| Q9D0B0 | Srsf9 | 0.011 | 0.066 | 0.020 | 0.000 | -0.159 | -0.652 | -1.270 | -0.798 |
| Q9R062 | Gyg1 | 0.243 | 0.454 | 0.654 | 0.000 | -0.020 | 0.224 | -0.695 | -0.316 |
| Q921I1 | Tf | 0.819 | 1.000 | 0.000 | -0.479 | 0.122 | 0.389 | -0.261 | -0.276 |
| Q9DAT2 | Mrgbp | 0.207 | 0.404 | -0.081 | 0.000 | 0.111 | -0.097 | -1.350 | -0.296 |
| Q9R1E0 | Foxo1 | 0.005 | 0.039 | -0.131 | 0.075 | 0.000 | -0.360 | -0.460 | -0.388 |
| Q9WTR1 | Trpv2 | 0.227 | 0.430 | 0.000 | 0.025 | -0.046 | 0.378 | -0.095 | 0.370 |
| Q80TG1 | Kansl1 | 0.986 | 1.000 | -1.300 | 0.000 | 0.119 | -0.573 | -0.707 | 0.071 |
| O70194 | Eif3d | 0.591 | 0.864 | 0.071 | 0.000 | -0.109 | 0.166 | -0.463 | -0.074 |
| Q99LR1 | Abhd12 | 0.002 | 0.026 | 0.137 | 0.000 | -0.385 | 1.300 | 1.020 | 1.320 |
| P97310 | Mcm2 | 0.021 | 0.090 | 0.214 | -0.065 | 0.000 | -0.388 | -1.080 | -0.810 |
| Q62095 | Ddx3y | 0.466 | 0.724 | 0.597 | -0.003 | 0.000 | 0.486 | -0.267 | -0.483 |
| Q9JLI6 | Scly | 0.000 | 0.013 | -0.263 | 0.000 | 0.067 | -1.830 | -1.890 | -1.550 |
| P26645 | Marcks | 0.049 | 0.154 | 0.063 | 0.000 | -0.390 | 2.040 | 0.474 | 1.140 |
| Q3UKJ7 | Smu1 | 0.022 | 0.095 | -0.083 | 0.468 | 0.000 | -0.675 | -0.709 | -0.392 |
| Q9DCU6 | Mrpl4 | 0.041 | 0.138 | 0.000 | 0.039 | -0.885 | 0.969 | 0.968 | 0.394 |
| Q9ERL7 | Gmfg | 0.000 | 0.011 | 0.000 | 0.112 | -0.049 | -1.510 | -1.900 | -1.610 |
| Q8R180 | Ero1a | 0.132 | 0.296 | 0.317 | 0.000 | -0.244 | 0.636 | 0.184 | 0.436 |
| Q8K190 | Saysd1 | 0.726 | 1.000 | -0.379 | 0.000 | 0.155 | -0.127 | -0.154 | 0.302 |
| E9Q1P8 | Irf2bp2 | 0.832 | 1.000 | 0.177 | 0.000 | -0.061 | -0.107 | -0.185 | 0.296 |
| Q99KU0 | Vmp1 | 0.001 | 0.018 | 0.000 | 0.098 | -0.354 | 1.510 | 1.260 | 1.280 |
| Q8R5K4 | Nol6 | 0.229 | 0.434 | -0.275 | 0.000 | 0.070 | -0.056 | -0.418 | -0.407 |
| P51863 | Atp6v0d1 | 0.117 | 0.273 | 0.335 | 0.000 | -0.216 | 0.754 | 0.302 | 0.348 |
| Q9CQW9 | Ifitm3 | 0.039 | 0.133 | 1.730 | 0.000 | -0.642 | 2.840 | 2.290 | 2.540 |
| P62281 | Rps11 | 0.867 | 1.000 | 0.187 | -0.095 | 0.000 | 0.460 | -0.436 | -0.080 |
| A2A432 | Cul4b | 0.030 | 0.112 | -0.179 | 0.020 | 0.000 | -0.399 | -1.010 | -0.636 |
| O35984 | Pbx2 | 0.209 | 0.405 | -0.193 | 0.047 | 0.000 | -0.257 | -0.992 | -0.140 |
| P16460 | Ass1 | 0.143 | 0.310 | -0.587 | 0.000 | 0.018 | 0.512 | 0.153 | 0.075 |
| Q9DCG9 | Trmt112 | 0.673 | 0.950 | 0.000 | -4.950 | 0.195 | -0.811 | -3.810 | -2.730 |
| P27870 | Vav1 | 0.049 | 0.154 | 0.325 | 0.000 | -0.114 | -0.272 | -1.240 | -1.350 |
| P01901 | H2-K1 | 0.918 | 1.000 | 0.000 | 1.100 | -0.068 | -0.245 | 0.777 | 0.664 |
| Q8VDG3 | Parn | 0.009 | 0.055 | 0.000 | 0.031 | -0.430 | -0.991 | -1.720 | -1.620 |
| Q6P8I4 | Pcnp | 0.120 | 0.277 | 0.000 | 1.530 | -0.170 | -0.543 | -0.732 | -0.579 |
| P63154 | Crnkl1 | 0.013 | 0.069 | -0.110 | 0.068 | 0.000 | -0.499 | -0.964 | -0.564 |
| Q9CR58 | Slc25a30 | 0.011 | 0.062 | 0.000 | 0.243 | -0.162 | 0.866 | 0.561 | 0.664 |
| P62717 | Rpl18a | 0.672 | 0.949 | 0.000 | -0.370 | 0.023 | 0.764 | -0.398 | -0.192 |
| Q791T5 | Mtch1 | 0.001 | 0.019 | 0.000 | 0.409 | -0.005 | 1.910 | 1.480 | 1.660 |
| P10630 | Eif4a2 | 0.000 | 0.013 | -0.146 | 0.000 | 0.209 | -1.250 | -1.450 | -1.260 |
| Q9D7G0 | Prps1 | 0.019 | 0.087 | 0.000 | -0.080 | 0.012 | -0.415 | -1.060 | -1.060 |
| Q9DB70 | Fundc1 | 0.256 | 0.470 | -0.517 | 1.650 | 0.000 | 2.470 | 0.712 | 1.270 |
| Q9QWT9 | Kifc1 | 0.255 | 0.469 | 0.245 | 0.000 | -0.236 | 0.102 | -1.330 | -0.505 |
| P18654 | Rps6ka3 | 0.028 | 0.108 | 0.000 | -0.035 | 0.109 | -0.413 | -1.370 | -1.310 |
| Q9JJI8 | Rpl38 | 0.723 | 1.000 | 0.346 | -0.062 | 0.000 | 0.418 | 0.040 | 0.031 |
| Q99KF1 | Tmed9 | 0.007 | 0.047 | 0.000 | 0.304 | -0.089 | 0.916 | 0.738 | 1.130 |
| P00375 | Dhfr | 0.067 | 0.187 | 0.141 | -0.471 | 0.000 | -0.723 | -1.620 | -0.675 |
| P37913 | Lig1 | 0.180 | 0.367 | 0.049 | 0.000 | -0.161 | 0.072 | -0.870 | -1.240 |
| P29533 | Vcam1 | 0.146 | 0.315 | 0.746 | 0.000 | -0.120 | 0.114 | -1.040 | -0.893 |
| P59325 | Eif5 | 0.094 | 0.236 | 0.365 | 0.000 | -0.125 | -0.059 | -1.180 | -0.979 |
| Q9DC16 | Ergic1 | 0.000 | 0.008 | 0.000 | 0.202 | 0.000 | 1.680 | 1.520 | 1.720 |
| P43276 | Hist1h1b | 0.047 | 0.151 | -0.012 | 0.000 | 0.126 | -0.123 | -0.379 | -0.159 |

| | | | | | | | | | |
|---|---|---|---|---|---|---|---|---|---|
| O88673 | Dgka | 0.006 | 0.046 | 0.000 | 0.135 | -0.029 | -0.968 | -1.940 | -1.570 |
| Q61712 | Dnajc1 | 0.294 | 0.522 | 0.000 | -0.008 | 0.013 | 0.295 | -0.127 | 0.666 |
| Q9CXZ1 | Ndufs4 | 0.012 | 0.067 | -0.105 | 0.323 | 0.000 | 0.881 | 0.660 | 1.120 |
| O70400 | Pdlim1 | 0.000 | 0.014 | -0.049 | 0.191 | 0.000 | -0.883 | -0.928 | -0.774 |
| P27046 | Man2a1 | 0.004 | 0.038 | 0.100 | 0.000 | -0.187 | 0.809 | 0.590 | 0.982 |
| Q3TVI8 | Pbxip1 | 0.001 | 0.017 | -0.130 | 0.192 | 0.000 | 2.250 | 1.830 | 1.640 |
| Q8BY71 | Hat1 | 0.097 | 0.239 | 0.491 | 0.000 | -0.180 | -0.101 | -1.180 | -0.855 |
| Q9CQD1 | Rab5a | 0.358 | 0.603 | -0.260 | 0.000 | 0.058 | 0.299 | -0.184 | 0.248 |
| Q06185 | Atp5i | 0.001 | 0.020 | 0.000 | -0.021 | 0.001 | 0.709 | 0.524 | 0.804 |
| Q80X71 | Tmem106b | 0.015 | 0.076 | -0.073 | 0.000 | 0.115 | 1.150 | 0.542 | 0.683 |
| Q6Q477 | Atp2b4 | 0.616 | 0.890 | -0.793 | 0.069 | 0.000 | -0.045 | -0.917 | -0.375 |
| Q9JK81 | Myg1 | 0.012 | 0.068 | -0.941 | 0.000 | 0.003 | -1.670 | -1.890 | -1.610 |
| Q62388 | Atm | 0.107 | 0.257 | -0.265 | 0.116 | 0.000 | -0.451 | -1.290 | -0.368 |
| Q6DVA0 | Lemd2 | 0.002 | 0.024 | -0.038 | 0.000 | 0.064 | 0.752 | 0.567 | 0.883 |
| Q8BMS9 | Rassf2 | 0.010 | 0.059 | 0.048 | -0.167 | 0.000 | -0.794 | -1.680 | -1.350 |
| P18653 | Rps6ka1 | 0.130 | 0.293 | 0.310 | -0.305 | 0.000 | -0.035 | -2.120 | -1.510 |
| Q9JMD0 | Znf207 | 0.085 | 0.221 | -0.017 | 0.438 | 0.000 | -0.369 | -0.827 | -0.122 |
| P62137 | Ppp1ca | 0.822 | 1.000 | 0.237 | -0.916 | 0.000 | -0.030 | -0.612 | -0.318 |
| Q8BQZ5 | Cpsf4 | 0.017 | 0.080 | 0.000 | 0.329 | -0.003 | -0.392 | -0.615 | -0.331 |
| A2BE28 | Las1l | 0.144 | 0.311 | 0.000 | 0.020 | -0.132 | -0.086 | -0.921 | -0.445 |
| Q8BFV2 | Pcid2 | 0.056 | 0.167 | 0.000 | -0.229 | 0.168 | -0.404 | -1.510 | -0.871 |
| Q9CQN7 | Mrpl41 | 0.001 | 0.019 | -0.057 | 0.242 | 0.000 | 1.080 | 0.872 | 1.000 |
| Q8BHS3 | Rbm22 | 0.033 | 0.120 | -0.099 | 0.000 | 0.141 | -0.234 | -0.540 | -0.298 |
| P08920 | Cd2 | 0.013 | 0.069 | -0.128 | 0.000 | 0.180 | -0.418 | -0.328 | -0.464 |
| O35114 | Scarb2 | 0.362 | 0.607 | 0.450 | -0.680 | 0.000 | 0.717 | 0.016 | 0.235 |
| O35129 | Phb2 | 0.003 | 0.030 | 0.000 | 0.062 | -0.164 | 0.922 | 0.591 | 0.718 |
| P61027 | Rab10 | 0.511 | 0.776 | 0.322 | 0.000 | -0.168 | -0.098 | 0.011 | -0.079 |
| Q9D8T2 | Gsdmdc1 | 0.014 | 0.073 | 0.032 | 0.000 | -0.234 | -0.817 | -1.860 | -1.440 |
| Q9JI44 | Dmap1 | 0.183 | 0.371 | 0.000 | -0.120 | 0.075 | -0.268 | -0.446 | -0.006 |
| Q80TY0 | Fnbp1 | 0.000 | 0.014 | -0.105 | 0.201 | 0.000 | -1.180 | -1.500 | -1.310 |
| O70404 | Vamp8 | 0.765 | 1.000 | 0.004 | -0.083 | 0.000 | 0.196 | -0.274 | -0.138 |
| Q8BYK4 | Rdh12 | 0.365 | 0.612 | 0.037 | 0.000 | -0.061 | 0.037 | -0.072 | -0.880 |
| Q3UMY5 | Eml4 | 0.063 | 0.180 | 0.000 | -0.076 | 0.798 | -1.170 | -0.342 | -0.609 |
| Q62426 | Cstb | 0.075 | 0.202 | 0.000 | 0.417 | -0.437 | -0.446 | -0.934 | -0.684 |
| P17047 | Lamp2 | 0.008 | 0.053 | 0.337 | 0.000 | -0.296 | 1.220 | 0.851 | 1.180 |
| P97823 | Lypla1 | 0.168 | 0.350 | 0.000 | -0.250 | 0.196 | 0.058 | -1.140 | -0.961 |
| P35585 | Ap1m1 | 0.027 | 0.107 | 0.008 | -0.381 | 0.000 | -0.626 | -1.250 | -0.796 |
| P54726 | Rad23a | 0.000 | 0.014 | -0.136 | 0.081 | 0.000 | -1.310 | -1.450 | -1.100 |
| P62858 | Rps28 | 0.085 | 0.221 | 0.213 | -0.107 | 0.000 | -0.184 | -0.653 | -0.247 |
| Q61503 | Nt5e | 0.368 | 0.615 | -0.517 | 0.000 | 0.009 | -0.654 | -0.461 | -0.109 |
| Q8BVF7 | Aph1a | 0.804 | 1.000 | 0.000 | -0.832 | 2.030 | 2.600 | -2.380 | 2.440 |
| Q9CXW2 | Mrps22 | 0.536 | 0.803 | 0.837 | -1.070 | 0.000 | 1.960 | 1.660 | -1.400 |
| Q91VT4 | Cbr4 | 0.890 | 1.000 | 0.000 | 0.188 | -0.136 | 0.418 | -0.152 | -0.122 |
| Q9JJZ4 | Ube2j1 | 0.339 | 0.578 | -0.229 | 0.072 | 0.000 | 0.103 | -0.107 | 0.459 |
| P61089 | Ube2n | 0.014 | 0.073 | -0.057 | 0.000 | 0.306 | -0.454 | -0.788 | -0.468 |
| Q9JKF7 | Mrpl39 | 0.494 | 0.759 | 0.000 | -3.790 | 0.010 | 1.100 | -0.144 | -1.440 |
| P20444 | Prkca | 0.030 | 0.113 | -0.268 | 0.109 | 0.000 | -0.390 | -0.460 | -0.434 |
| P32037 | Slc2a3 | 0.192 | 0.384 | 0.000 | 0.198 | -1.110 | -0.827 | -0.767 | -1.920 |
| P50543 | S100a11 | 0.359 | 0.603 | 0.112 | -0.422 | 0.000 | 0.231 | -1.160 | -0.762 |
| Q9JJZ6 | Klf13 | 0.026 | 0.103 | -0.094 | 0.000 | 0.094 | -0.359 | -0.751 | -1.050 |
| Q8BNW9 | Kbtbd11 | 0.007 | 0.048 | -0.061 | 0.000 | 0.260 | -1.280 | -2.450 | -1.660 |
| Q60787 | Lcp2 | 0.009 | 0.057 | -0.096 | 0.845 | 0.000 | -1.390 | -1.550 | -1.040 |
| Q02242 | Pdcd1 | 0.396 | 0.646 | -0.498 | 0.000 | 0.112 | -0.263 | 0.315 | 0.333 |
| Q9Z0L8 | Ggh | 0.394 | 0.643 | -0.367 | 0.486 | 0.000 | -0.505 | -0.272 | 0.051 |
| Q8VC03 | Eml3 | 0.027 | 0.105 | 0.000 | -0.105 | 0.575 | -0.465 | -0.766 | -0.650 |
| A2AQ19 | Rtf1 | 0.008 | 0.052 | 0.023 | -0.348 | 0.000 | -0.937 | -1.470 | -1.690 |
| Q8BG30 | Nelfa | 0.687 | 0.963 | -0.364 | 0.000 | 0.565 | -0.184 | 0.126 | -0.112 |
| Q91VR2 | Atp5c1 | 0.035 | 0.125 | 0.127 | 0.000 | -0.028 | 1.150 | 0.535 | 0.475 |
| Q62425 | Ndufa4 | 0.219 | 0.419 | -0.214 | 0.210 | 0.000 | 0.195 | 0.073 | 0.541 |

| | | | | | | | | | |
|---|---|---|---|---|---|---|---|---|---|
| P63242 | Eif5a | 0.208 | 0.404 | 0.466 | -0.135 | 0.000 | 0.222 | -1.650 | -0.813 |
| Q6ZQ58 | Larp1 | 0.589 | 0.861 | 0.238 | -0.291 | 0.000 | 0.329 | -0.257 | 0.307 |
| P46978 | Stt3a | 0.000 | 0.013 | 0.000 | 0.119 | -0.213 | 1.570 | 1.280 | 1.410 |
| Q9D5T0 | Atad1 | 0.387 | 0.636 | 0.005 | -0.010 | 0.000 | 0.504 | -0.072 | 0.068 |
| Q9JKX6 | Nudt5 | 0.000 | 0.014 | 0.033 | -0.029 | 0.000 | -1.110 | -1.520 | -1.260 |
| Q91YR7 | Prpf6 | 0.053 | 0.162 | 0.000 | 0.162 | -0.181 | -0.213 | -0.817 | -0.784 |
| Q6P1F6 | Ppp2r2a | 0.006 | 0.044 | -0.191 | 0.010 | 0.000 | -0.731 | -1.310 | -1.050 |
| Q8K212 | Pacs1 | 0.237 | 0.444 | -0.105 | 0.819 | 0.000 | 0.622 | 0.460 | 1.960 |
| P35980 | Rpl18 | 0.134 | 0.298 | 0.241 | -0.111 | 0.000 | 0.694 | 0.106 | 0.452 |
| Q9R099 | Tbl2 | 0.033 | 0.120 | 0.000 | 0.449 | -0.050 | 0.634 | 0.647 | 0.976 |
| Q64261 | Cdk6 | 0.020 | 0.089 | 0.186 | -0.124 | 0.000 | -0.539 | -1.450 | -1.390 |
| P63037 | Dnaja1 | 0.340 | 0.580 | 0.072 | 0.000 | -0.107 | 0.131 | -0.492 | -0.294 |
| Q3V009 | Tmed1 | 0.002 | 0.022 | 0.000 | 0.224 | -0.659 | 3.120 | 2.230 | 2.850 |
| Q8BGD9 | Eif4b | 0.065 | 0.184 | 0.178 | -0.283 | 0.000 | -0.447 | -1.800 | -1.000 |
| Q8QZS1 | Hibch | 0.404 | 0.655 | 0.000 | 0.318 | -0.092 | 0.156 | 0.135 | 0.318 |
| Q8BYB9 | Poglut1 | 0.033 | 0.121 | 0.293 | -0.547 | 0.000 | 1.410 | 0.557 | 1.140 |
| Q9EPJ9 | Arfgap1 | 0.189 | 0.379 | 0.000 | 0.443 | -0.044 | -0.772 | 0.231 | -1.060 |
| Q99J56 | Derl1 | 0.057 | 0.168 | 0.000 | 0.353 | -0.292 | 0.483 | 0.610 | 0.489 |
| Q8CBG9 | Rnf170 | 0.048 | 0.152 | -0.653 | 0.156 | 0.000 | 0.587 | 0.421 | 0.902 |
| P15379 | Cd44 | 0.076 | 0.205 | 0.008 | -0.076 | 0.000 | 0.622 | 0.100 | 0.309 |
| Q9CXT7 | Tmem192 | 0.444 | 0.701 | 0.035 | 0.000 | -0.127 | -0.080 | -0.516 | 0.051 |
| Q99L04 | Dhrs1 | 0.470 | 0.729 | 0.040 | 0.000 | -0.227 | 0.315 | -0.898 | -0.478 |
| Q4VBD2 | Tapt1 | 0.005 | 0.039 | 0.000 | 0.047 | -0.033 | 0.806 | 0.438 | 0.702 |
| P70158 | Smpdl3a | 0.093 | 0.234 | 0.547 | -0.004 | 0.000 | -0.204 | -1.990 | -0.899 |
| P05201 | Got1 | 0.000 | 0.013 | -0.145 | 0.019 | 0.000 | -1.300 | -1.680 | -1.440 |
| P83917 | Cbx1 | 0.031 | 0.115 | -0.155 | 0.623 | 0.000 | -1.040 | -0.915 | -0.466 |
| Q99KB8 | Hagh | 0.754 | 1.000 | 0.142 | 0.000 | -0.442 | -0.018 | -0.512 | -0.014 |
| Q8VD65 | Pik3r4 | 0.075 | 0.203 | 0.065 | 0.000 | -0.273 | -0.475 | -2.190 | -1.180 |
| P70388 | Rad50 | 0.176 | 0.360 | 0.000 | -0.036 | 0.010 | 0.053 | -0.597 | -0.457 |
| Q8BHA0 | Ino80c | 0.191 | 0.383 | -0.478 | 0.295 | 0.000 | -0.586 | -0.487 | -0.263 |
| Q9D1C9 | Rrp7a | 0.568 | 0.837 | 0.000 | 1.660 | -0.041 | -1.260 | 0.528 | 0.770 |
| Q05D44 | Eif5b | 0.751 | 1.000 | 0.544 | 0.000 | -0.060 | 1.210 | -0.708 | 0.591 |
| Q91ZR1 | Rab4b | 0.047 | 0.151 | 0.000 | -0.097 | 0.211 | -0.147 | -0.349 | -0.366 |
| Q91VX9 | Tmem168 | 0.078 | 0.208 | 0.000 | 0.153 | -0.166 | 0.492 | 0.337 | 0.135 |
| Q9CWS4 | Cpsf3l | 0.800 | 1.000 | -1.180 | 0.444 | 0.000 | -0.383 | -0.246 | 0.328 |
| Q0VGB7 | Ppp4r2 | 0.007 | 0.048 | 0.097 | -0.490 | 0.000 | -1.190 | -1.610 | -1.190 |
| Q8VCG3 | Wdr74 | 0.068 | 0.188 | 0.053 | 0.000 | -0.255 | -0.230 | -0.618 | -0.810 |
| P48377 | Rfx1 | 0.168 | 0.350 | 0.000 | 1.250 | -1.230 | -2.680 | -3.290 | -0.112 |
| P03888 | Mtnd1 | 0.570 | 0.839 | -0.830 | 0.541 | 0.000 | -0.768 | 0.658 | 1.100 |
| Q91XA2 | Golm1 | 0.144 | 0.311 | 0.162 | -0.030 | 0.000 | -0.479 | -0.405 | 0.053 |
| Q9JLJ2 | Aldh9a1 | 0.081 | 0.213 | 0.030 | -0.088 | 0.000 | -0.239 | -1.410 | -0.779 |
| Q9CR57 | Rpl14 | 0.016 | 0.078 | 0.142 | 0.000 | -0.271 | 0.753 | 0.423 | 0.557 |
| Q689Z5 | Sbno1 | 0.023 | 0.096 | 0.000 | 0.229 | -0.069 | -0.550 | -1.670 | -1.430 |
| P31266 | Rbpj | 0.029 | 0.110 | 0.068 | -0.003 | 0.000 | -0.262 | -0.886 | -0.617 |
| P62342 | Selt | 0.115 | 0.270 | 0.000 | 0.100 | -0.390 | 0.284 | 0.120 | 0.255 |
| P41245 | Mmp9 | 0.231 | 0.435 | 2.280 | 0.000 | -0.046 | 0.785 | -1.710 | -1.570 |
| Q03147 | Cdk7 | 0.029 | 0.110 | 0.000 | -0.008 | 0.017 | -0.257 | -0.564 | -0.237 |
| Q8K2Y9 | Ccm2 | 0.105 | 0.252 | 0.000 | -0.079 | 0.056 | -0.182 | -0.525 | -0.131 |
| P61205 | Arf3 | 0.041 | 0.138 | 0.110 | 0.000 | -0.115 | -0.215 | -0.803 | -0.684 |
| P84078 | Arf1 | 0.041 | 0.138 | 0.110 | 0.000 | -0.115 | -0.215 | -0.803 | -0.684 |
| Q8CG48 | Smc2 | 0.516 | 0.779 | 0.399 | -0.110 | 0.000 | 0.404 | -0.520 | -0.271 |
| Q6PFR5 | Tra2a | 0.863 | 1.000 | -0.783 | 1.460 | 0.000 | 0.388 | 0.513 | 0.140 |
| Q8C0M9 | Asrgl1 | 0.000 | 0.011 | 0.000 | -0.030 | 0.298 | -1.320 | -1.440 | -1.400 |
| Q8CI11 | Gnl3 | 0.503 | 0.767 | -0.054 | 0.404 | 0.000 | 0.261 | -0.582 | 0.031 |
| P59708 | Sf3b6 | 0.380 | 0.630 | 0.000 | 0.753 | -0.018 | -1.090 | 0.063 | 0.292 |
| Q9ERU3 | Znf22 | 0.139 | 0.304 | 0.026 | -0.097 | 0.000 | -0.267 | -2.210 | -0.809 |
| Q91VM9 | Ppa2 | 0.070 | 0.193 | -0.028 | 0.474 | 0.000 | 0.613 | 0.457 | 0.874 |
| P10126 | Eef1a1 | 0.463 | 0.721 | 0.222 | -0.135 | 0.000 | 0.343 | -0.593 | -0.404 |
| P13597 | Icam1 | 0.007 | 0.050 | 0.000 | 0.319 | -0.108 | 0.945 | 0.690 | 0.810 |

| | | | | | | | | | |
|---|---|---|---|---|---|---|---|---|---|
| Q8C0C7 | Farsa | 0.044 | 0.144 | 0.000 | 0.024 | -0.200 | -0.294 | -0.996 | -0.832 |
| P28658 | Atxn10 | 0.868 | 1.000 | 0.308 | 0.000 | -0.205 | 0.902 | -0.791 | -0.288 |
| P28474 | Adh5 | 0.003 | 0.032 | -0.238 | 0.142 | 0.000 | -1.150 | -1.810 | -1.400 |
| O70551 | Srpk1 | 0.012 | 0.067 | 0.029 | 0.000 | -0.348 | -0.612 | -0.911 | -0.979 |
| Q9D6T0 | Nosip | 0.272 | 0.491 | 0.000 | -0.462 | 0.101 | -0.324 | -0.810 | -0.199 |
| Q9ES97 | Rtn3 | 0.173 | 0.357 | 0.075 | 0.000 | -0.021 | 0.550 | 0.036 | 0.226 |
| P10639 | Txn | 0.016 | 0.079 | 0.196 | -0.365 | 0.000 | -0.914 | -2.060 | -1.810 |
| Q9D8P4 | Mrpl17 | 0.062 | 0.179 | -0.135 | 0.609 | 0.000 | 0.806 | 0.896 | 0.632 |
| Q922D4 | Ppp6r3 | 0.039 | 0.133 | 0.000 | 0.104 | -0.045 | -0.337 | -1.250 | -0.779 |
| Q61144 | Psen2 | 0.255 | 0.468 | -0.215 | 0.716 | 0.000 | 0.567 | 0.830 | 0.354 |
| P57716 | Ncstn | 0.012 | 0.067 | 0.058 | 0.000 | -0.094 | 1.230 | 0.526 | 1.070 |
| Q61187 | Tsg101 | 0.722 | 1.000 | 0.000 | 0.043 | -0.115 | 0.395 | -0.280 | 0.042 |
| Q9Z204 | Hnrnpc | 0.274 | 0.494 | -0.174 | 0.244 | 0.000 | -0.399 | -0.244 | 0.040 |
| Q923D2 | Blvrb | 0.072 | 0.196 | 0.052 | 0.000 | -0.959 | -0.794 | -2.040 | -1.690 |
| Q91WG4 | Elp2 | 0.084 | 0.220 | 0.228 | -0.048 | 0.000 | -0.173 | -1.420 | -0.763 |
| P47791 | Gsr | 0.018 | 0.084 | 0.341 | 0.000 | -0.118 | -0.466 | -1.080 | -1.080 |
| Q75N62 | Gimap8 | 0.012 | 0.069 | -0.070 | 0.000 | 0.009 | -0.447 | -0.926 | -0.575 |
| Q9JLT4 | Txnrd2 | 0.839 | 1.000 | 0.000 | 0.351 | -0.041 | 0.125 | 0.030 | 0.247 |
| Q9DCD2 | Xab2 | 0.010 | 0.058 | -0.080 | 0.000 | 0.077 | -0.302 | -0.602 | -0.563 |
| Q9CQ69 | Uqcrq | 0.004 | 0.038 | 0.000 | 0.165 | -0.022 | 0.638 | 0.470 | 0.464 |
| Q9CZU6 | Cs | 0.005 | 0.041 | 0.000 | 0.137 | -0.034 | 0.868 | 0.509 | 0.669 |
| Q9D662 | Sec23b | 0.820 | 1.000 | 0.129 | -2.060 | 0.000 | -0.568 | -1.170 | -0.727 |
| O88967 | Yme1l1 | 0.017 | 0.081 | 0.000 | -0.237 | 0.021 | 0.822 | 0.328 | 0.838 |
| Q8K1R7 | Nek9 | 0.096 | 0.238 | -0.055 | 0.220 | 0.000 | -0.017 | -0.634 | -0.594 |
| P68181 | Prkacb | 0.979 | 1.000 | -1.250 | 0.245 | 0.000 | -0.353 | -0.234 | -0.378 |
| Q9DBG5 | Plin3 | 0.127 | 0.288 | -0.512 | 0.000 | 0.399 | -0.702 | -1.290 | -0.333 |
| P48193 | Epb41 | 0.179 | 0.364 | 0.171 | 0.000 | -0.123 | 0.082 | -0.784 | -0.608 |
| Q5DU31 | Ipcef1 | 0.108 | 0.259 | -0.777 | 0.000 | 0.265 | -1.270 | -2.170 | -0.550 |
| Q60648 | Gm2a | 0.038 | 0.129 | 0.000 | -0.182 | 0.048 | -0.348 | -1.050 | -0.713 |
| P83940 | Tceb1 | 0.011 | 0.064 | -0.336 | 0.000 | 0.298 | -1.000 | -1.880 | -1.370 |
| Q9JHK5 | Plek | 0.404 | 0.655 | 0.273 | 0.000 | -0.425 | 0.193 | -0.584 | -0.757 |
| P35282 | Rab21 | 1.000 | 1.000 | -0.077 | 0.256 | 0.000 | 0.281 | -0.147 | 0.045 |
| Q6ZPY7 | Kdm3b | 0.027 | 0.107 | 0.000 | 0.495 | -0.124 | -0.530 | -1.600 | -1.240 |
| Q8BU14 | Sec62 | 0.007 | 0.049 | 0.047 | -0.056 | 0.000 | 0.844 | 0.419 | 0.756 |
| P46414 | Cdkn1b | 0.005 | 0.039 | -0.490 | 0.000 | 0.483 | -2.560 | -3.040 | -1.880 |
| Q99J72 | Apobec3 | 0.052 | 0.160 | 0.065 | -0.374 | 0.000 | -0.391 | -1.200 | -1.040 |
| Q811U4 | Mfn1 | 0.420 | 0.674 | 0.000 | 1.260 | -0.739 | 2.090 | 1.140 | -0.275 |
| P35329 | Cd22 | 0.083 | 0.218 | -0.729 | 0.548 | 0.000 | -3.200 | -2.990 | -0.495 |
| Q99J47 | Dhrs7b | 0.010 | 0.059 | 0.000 | 0.380 | -0.485 | 1.100 | 1.150 | 1.120 |
| Q91YI0 | Asl | 0.025 | 0.102 | -0.108 | 0.298 | 0.000 | -1.820 | -0.654 | -2.060 |
| P68433 | Hist1h3a | 0.422 | 0.676 | 0.323 | -0.184 | 0.000 | 0.348 | -0.110 | 0.569 |
| P84228 | Hist1h3b | 0.422 | 0.676 | 0.323 | -0.184 | 0.000 | 0.348 | -0.110 | 0.569 |
| P84244 | H3f3a | 0.422 | 0.676 | 0.323 | -0.184 | 0.000 | 0.348 | -0.110 | 0.569 |
| Q62433 | Ndrg1 | 0.116 | 0.271 | 0.000 | 0.021 | -0.415 | 0.448 | 0.000 | 0.329 |
| Q9D1R9 | Rpl34 | 0.210 | 0.407 | 0.167 | -0.106 | 0.000 | 0.638 | 0.073 | 0.196 |
| Q9JHI7 | Exosc9 | 0.019 | 0.087 | -0.189 | 0.046 | 0.000 | -0.782 | -0.947 | -0.406 |
| Q9CQM5 | Txndc17 | 0.008 | 0.055 | -0.143 | 0.362 | 0.000 | -1.500 | -1.760 | -2.830 |
| Q9D6J5 | Ndufb8 | 0.039 | 0.132 | -0.884 | 0.149 | 0.000 | 0.958 | 0.797 | 0.597 |
| Q9QZ06 | Tollip | 0.998 | 1.000 | 0.443 | 0.000 | -0.223 | 0.233 | -0.413 | 0.398 |
| Q8BJ64 | Chdh | 0.375 | 0.623 | 0.970 | -0.135 | 0.000 | 0.659 | -1.190 | -0.567 |
| Q9Z1R2 | Bag6 | 0.633 | 0.909 | 0.089 | 0.000 | -0.303 | 0.178 | -0.646 | -0.160 |
| Q6NZF1 | Zc3h11a | 0.003 | 0.032 | -0.026 | 0.242 | 0.000 | -0.859 | -0.855 | -0.564 |
| Q8R0F3 | Sumf1 | 0.002 | 0.024 | 0.000 | 0.232 | -0.568 | 1.860 | 1.650 | 1.660 |
| P48722 | Hspa4l | 0.006 | 0.047 | -0.299 | 0.139 | 0.000 | -1.170 | -0.804 | -1.320 |
| Q810A7 | Ddx42 | 0.810 | 1.000 | 0.110 | -5.730 | 0.000 | -0.901 | -2.390 | -0.792 |
| Q9CQN6 | Tmem14c | 0.066 | 0.186 | 0.146 | 0.000 | -0.210 | 0.760 | 0.282 | 0.302 |
| Q9CQL5 | Mrpl18 | 0.012 | 0.067 | -0.027 | 0.185 | 0.000 | 0.925 | 0.512 | 1.060 |
| Q8VEE4 | Rpa1 | 0.072 | 0.197 | 0.054 | -0.271 | 0.000 | -0.678 | -1.690 | -0.560 |
| Q03267 | Ikzf1 | 0.113 | 0.266 | -0.067 | 0.000 | 0.167 | -0.242 | -1.470 | -0.496 |

| | | | | | | | | | |
|---|---|---|---|---|---|---|---|---|---|
| Q8CI08 | Slain2 | 0.911 | 1.000 | -1.240 | 0.000 | 1.400 | -0.021 | -0.848 | 1.380 |
| Q9CQR2 | Rps21 | 0.095 | 0.237 | 0.311 | 0.000 | -0.025 | -0.024 | -0.691 | -0.461 |
| Q8R2Q4 | Gfm2 | 0.142 | 0.309 | 0.000 | 0.144 | -0.128 | 0.042 | -1.100 | -0.903 |
| P43274 | Hist1h1e | 0.210 | 0.406 | 0.191 | 0.000 | -0.177 | -0.111 | -0.417 | -0.115 |
| Q76KJ5 | Cd3eap | 0.295 | 0.522 | -0.247 | 0.366 | 0.000 | -0.128 | -0.249 | -0.160 |
| Q9D710 | Tmx2 | 0.003 | 0.030 | 0.122 | 0.000 | -0.249 | 0.828 | 0.670 | 0.867 |
| O35316 | Slc6a6 | 0.021 | 0.092 | 1.240 | 0.000 | -0.326 | 3.830 | 2.060 | 2.770 |
| Q9QYI3 | Dnajc7 | 0.064 | 0.183 | 0.049 | 0.000 | -0.092 | -0.189 | -0.999 | -0.668 |
| Q9JKB3 | Ybx3 | 0.504 | 0.768 | 0.000 | 0.000 | -0.501 | 0.146 | -0.242 | 0.042 |
| Q9CPT5 | Nop16 | 0.751 | 1.000 | 0.000 | 0.393 | -0.018 | 0.600 | -0.657 | 0.037 |
| P70403 | Cux1 | 0.336 | 0.573 | -2.080 | 0.000 | 1.250 | 0.628 | 0.354 | 1.600 |
| Q9D8V0 | Hm13 | 0.029 | 0.110 | -0.063 | 0.378 | 0.000 | 0.955 | 0.512 | 0.731 |
| Q9CPX6 | Atg3 | 0.178 | 0.364 | 0.037 | 0.000 | -0.063 | -0.014 | -0.809 | -0.341 |
| P97313 | Prkdc | 0.549 | 0.818 | 0.000 | -0.329 | 0.216 | -0.441 | -0.410 | 0.217 |
| Q9DBG7 | Srpra | 0.996 | 1.000 | 0.111 | 0.000 | -0.032 | 0.308 | -0.444 | 0.219 |
| P31230 | Aimp1 | 0.280 | 0.502 | 0.052 | 0.000 | -0.151 | 0.072 | -0.920 | -0.350 |
| Q99J95 | Cdk9 | 0.029 | 0.110 | 0.000 | -0.077 | 0.030 | -0.617 | -1.760 | -1.040 |
| Q8JZU2 | Slc25a1 | 0.000 | 0.012 | 0.000 | 0.212 | -0.083 | 1.600 | 1.430 | 1.730 |
| Q3TJD7 | Pdlim7 | 0.000 | 0.014 | -0.291 | 0.000 | 0.069 | 2.390 | 2.110 | 2.820 |
| Q6PD03 | Ppp2r5a | 0.406 | 0.658 | 0.305 | -0.812 | 0.000 | -0.080 | -1.290 | -0.489 |
| Q8CG72 | Adprhl2 | 0.003 | 0.033 | -0.365 | 0.161 | 0.000 | -1.260 | -1.820 | -1.390 |
| Q8BXV2 | Bri3bp | 0.054 | 0.163 | 0.300 | -0.665 | 0.000 | 1.270 | 0.551 | 0.721 |
| Q62311 | Taf6 | 0.460 | 0.719 | -0.606 | 0.000 | 0.112 | -0.200 | -0.935 | -0.177 |
| Q99K23 | Ufsp2 | 0.019 | 0.087 | 0.081 | 0.000 | -0.422 | 0.696 | 0.388 | 0.640 |
| P61924 | Copz1 | 0.013 | 0.070 | 0.000 | 0.191 | -0.211 | -0.551 | -0.896 | -0.603 |
| P31041 | Cd28 | 0.187 | 0.376 | -0.410 | 0.000 | 0.214 | -0.507 | -0.630 | -0.161 |
| Q60973 | Rbbp7 | 0.041 | 0.138 | -0.182 | 0.071 | 0.000 | -0.476 | -0.460 | -0.205 |
| Q9EP97 | Senp3 | 0.650 | 0.926 | -0.050 | 0.562 | 0.000 | 0.402 | -0.247 | -0.046 |
| Q8BU88 | Mrpl22 | 0.017 | 0.080 | 0.003 | -0.068 | 0.000 | 0.895 | 0.346 | 0.595 |
| P53702 | Hccs | 0.001 | 0.022 | -0.130 | 0.313 | 0.000 | 1.420 | 1.460 | 1.860 |
| Q6PGC1 | Dhx29 | 0.175 | 0.359 | -0.668 | 0.188 | 0.000 | -0.453 | -1.060 | -0.550 |
| Q9D787 | Ppil2 | 0.371 | 0.618 | -0.918 | 0.000 | 0.190 | -0.048 | -2.360 | -0.653 |
| Q80TJ7 | Phf8 | 0.268 | 0.487 | -0.121 | 1.600 | 0.000 | 0.400 | -1.350 | -0.462 |
| P04202 | Tgfb1 | 0.054 | 0.163 | 0.000 | 0.044 | -0.363 | -0.731 | -0.344 | -0.794 |
| Q9WUM4 | Coro1c | 0.649 | 0.925 | 0.000 | 0.820 | -0.303 | 0.531 | 0.356 | 0.152 |
| Q8R502 | Lrrc8c | 0.354 | 0.597 | -0.207 | 0.177 | 0.000 | -0.160 | -0.207 | -0.043 |
| O70572 | Smpd2 | 0.081 | 0.213 | 0.525 | -4.690 | 0.000 | 2.230 | 2.590 | 2.610 |
| Q61036 | Pak3 | 0.803 | 1.000 | -1.200 | 0.000 | 0.506 | -0.050 | -0.933 | -0.175 |
| P36552 | Cpox | 0.116 | 0.271 | 0.330 | 0.000 | -0.246 | -0.140 | -0.571 | -0.479 |
| Q9QUM9 | Psma6 | 0.010 | 0.060 | 0.000 | 0.763 | -0.042 | -1.220 | -1.070 | -0.858 |
| Q8R322 | Gle1 | 0.568 | 0.837 | 0.082 | 0.000 | -0.787 | -0.422 | -0.427 | -0.374 |
| P61021 | Rab5b | 0.370 | 0.617 | 0.000 | -1.020 | 0.280 | 0.548 | -2.430 | -1.880 |
| P99026 | Psmb4 | 0.001 | 0.021 | 0.000 | 0.003 | -0.115 | -1.070 | -1.620 | -1.350 |
| Q91V04 | Tram1 | 0.196 | 0.389 | 0.000 | 2.330 | -0.017 | 1.120 | 2.790 | 2.950 |
| Q9D0G0 | Mrps30 | 0.012 | 0.068 | 0.000 | 0.044 | -0.091 | 1.010 | 0.479 | 0.609 |
| Q9QYA2 | Tomm40 | 0.010 | 0.059 | 0.000 | 0.202 | -0.235 | 1.300 | 0.723 | 0.891 |
| P12023 | App | 0.032 | 0.117 | 4.640 | -1.820 | 0.000 | 7.180 | 7.130 | 7.230 |
| Q9Z0M5 | Lipa | 0.093 | 0.234 | 0.471 | 0.000 | -0.153 | -0.105 | -0.870 | -0.654 |
| Q62448 | Eif4g2 | 0.342 | 0.582 | 0.405 | -0.041 | 0.000 | 0.127 | -0.804 | -0.006 |
| Q9Z0W3 | Nup160 | 0.694 | 0.970 | 0.000 | 0.186 | -0.070 | -0.051 | -0.145 | 0.162 |
| P10922 | H1f0 | 0.130 | 0.293 | 0.000 | -0.145 | 0.190 | -0.040 | -1.060 | -0.629 |
| Q8CJG0 | Ago2 | 0.043 | 0.141 | -0.346 | 0.000 | 0.075 | -0.501 | -0.885 | -0.493 |
| O35144 | Terf2 | 0.061 | 0.176 | -0.147 | 0.408 | 0.000 | -0.504 | -0.507 | -0.217 |
| Q62446 | Fkbp3 | 0.063 | 0.181 | -0.025 | 0.000 | 0.005 | -0.303 | -0.475 | -0.092 |
| Q9JLQ0 | Cd2ap | 0.993 | 1.000 | 0.063 | -1.320 | 0.000 | -0.269 | -1.480 | 0.514 |
| Q3TLH4 | Prrc2c | 0.072 | 0.197 | -0.333 | 0.208 | 0.000 | -0.639 | -1.180 | -0.375 |
| Q8BYH7 | Tbc1d17 | 0.194 | 0.387 | 0.112 | -0.197 | 0.000 | 0.008 | -0.707 | -0.450 |
| P70290 | Mpp1 | 0.605 | 0.878 | 0.118 | 0.000 | -0.851 | -0.046 | -0.940 | -0.420 |
| P06342 | H2-Ab1 | 0.941 | 1.000 | 0.000 | 1.270 | -0.824 | -0.149 | 0.502 | 0.243 |

| | | | | | | | | | |
|---|---|---|---|---|---|---|---|---|---|
| Q8C208 | Ikzf4 | 0.461 | 0.719 | -0.864 | 0.212 | 0.000 | -0.745 | -0.405 | -0.361 |
| Q8VC28 | Akr1c13 | 0.001 | 0.019 | -0.340 | 0.000 | 0.003 | -1.700 | -2.330 | -1.900 |
| Q8BYM8 | Cars2 | 0.412 | 0.665 | 0.076 | 0.000 | -0.350 | 0.743 | -0.258 | 0.119 |
| Q64310 | Surf4 | 0.002 | 0.029 | 0.000 | 0.103 | -0.025 | 1.070 | 0.679 | 0.805 |
| Q91YK2 | Rrp1b | 0.028 | 0.107 | -0.039 | 0.046 | 0.000 | -0.265 | -0.819 | -0.549 |
| Q9CRA9 | Fgfr1op2 | 0.213 | 0.409 | -0.430 | 0.257 | 0.000 | -0.553 | -0.291 | -0.299 |
| Q6PGH1 | Bud31 | 0.005 | 0.039 | 0.183 | -0.025 | 0.000 | -0.445 | -0.774 | -0.762 |
| Q8BTX9 | Hsdl1 | 0.769 | 1.000 | 0.000 | 0.163 | -0.252 | 0.135 | -0.129 | 0.040 |
| Q61334 | Bcap29 | 0.670 | 0.948 | 0.026 | -0.020 | 0.000 | 0.419 | -0.310 | 0.194 |
| Q80US4 | Actr5 | 0.919 | 1.000 | -1.160 | 0.377 | 0.000 | 1.380 | -1.400 | -1.080 |
| P60766 | Cdc42 | 0.652 | 0.928 | 0.377 | -0.109 | 0.000 | 0.126 | -0.409 | 0.199 |
| Q61072 | Adam9 | 0.035 | 0.124 | 0.660 | -3.660 | 0.000 | 3.520 | 2.950 | 3.250 |
| P62331 | Arf6 | 0.347 | 0.587 | -0.116 | 0.120 | 0.000 | 0.221 | -0.074 | 0.255 |
| Q9CXW4 | Rpl11 | 0.252 | 0.466 | 0.375 | -0.080 | 0.000 | 0.127 | -0.615 | -0.243 |
| Q8CBE3 | Wdr37 | 0.204 | 0.400 | 0.000 | 0.548 | -0.350 | -0.276 | -0.350 | -0.371 |
| O88455 | Dhcr7 | 0.003 | 0.029 | -0.220 | 0.000 | 0.700 | 2.230 | 1.960 | 2.310 |
| Q63932 | Map2k2 | 0.693 | 0.969 | 0.000 | -0.792 | 0.019 | 0.453 | -1.150 | -0.779 |
| P14131 | Rps16 | 0.613 | 0.887 | 0.169 | 0.000 | -0.079 | 0.180 | -0.279 | -0.059 |
| P97434 | Mprip | 0.001 | 0.019 | 0.000 | -0.488 | 0.742 | 3.430 | 3.150 | 3.770 |
| Q8VE99 | Ccdc115 | 0.807 | 1.000 | -0.113 | 0.000 | 0.110 | -0.028 | -0.201 | 0.135 |
| Q60770 | Stxbp3 | 0.565 | 0.834 | 0.090 | 0.000 | -0.054 | 0.526 | -0.195 | 0.106 |
| Q80UM3 | Naa15 | 0.844 | 1.000 | 0.164 | -1.690 | 0.000 | 0.202 | -1.150 | -1.040 |
| Q9DBY8 | Nvl | 0.158 | 0.334 | 0.081 | -0.257 | 0.000 | -0.001 | -0.936 | -1.120 |
| Q920Q4 | Vps16 | 0.992 | 1.000 | 0.246 | 0.000 | -0.008 | 0.326 | -0.203 | 0.110 |
| Q9CZ28 | Snf8 | 0.469 | 0.728 | 0.006 | -0.006 | 0.000 | -0.028 | -0.403 | 0.079 |
| Q7TMM9 | Tubb2a | 0.111 | 0.264 | 0.000 | 0.407 | -0.148 | 1.530 | 0.414 | 0.633 |
| P04235 | Cd3d | 0.281 | 0.504 | -0.353 | 0.097 | 0.000 | -0.435 | -0.393 | -0.084 |
| Q9CX99 | Grap | 0.077 | 0.207 | -0.922 | 0.152 | 0.000 | -1.990 | -2.090 | -0.687 |
| Q99K70 | Rragc | 0.587 | 0.858 | 0.115 | 0.000 | -0.184 | 0.184 | -0.455 | -0.160 |
| Q8JZN7 | Rhot2 | 0.010 | 0.061 | 0.084 | 0.000 | -0.204 | 0.554 | 0.327 | 0.480 |
| P47963 | Rpl13 | 0.796 | 1.000 | -0.012 | 0.034 | 0.000 | 0.497 | -0.522 | 0.307 |
| Q571I9 | Aldh16a1 | 0.015 | 0.076 | 0.000 | 0.059 | -0.391 | -0.726 | -1.380 | -1.130 |
| B1AZA5 | Tmem245 | 0.094 | 0.236 | 0.000 | -4.310 | 0.381 | 1.750 | 1.800 | 2.530 |
| Q8R3C6 | Rbm19 | 0.085 | 0.220 | 0.110 | 0.000 | -0.230 | -0.172 | -1.180 | -0.999 |
| Q9Z0J0 | Npc2 | 0.678 | 0.954 | 0.038 | -0.325 | 0.000 | 0.294 | -0.329 | 0.035 |
| Q3UGR5 | Hdhd2 | 0.093 | 0.234 | 0.000 | 0.123 | -0.395 | -0.377 | -0.501 | -0.971 |
| P14115 | Rpl27a | 0.809 | 1.000 | 0.307 | 0.000 | -0.285 | 0.495 | -0.240 | -0.018 |
| Q9WUU9 | Mcm3ap | 0.519 | 0.784 | 0.121 | 0.000 | -0.223 | -0.095 | -0.723 | 0.133 |
| Q99KY4 | Gak | 0.193 | 0.385 | 0.471 | -0.202 | 0.000 | -0.838 | -3.880 | -0.314 |
| A2AL36 | Cntrl | 0.237 | 0.444 | 0.000 | -0.173 | 0.276 | -0.143 | -0.798 | -0.066 |
| Q80X41 | Vrk1 | 0.007 | 0.048 | 0.000 | -0.019 | 0.080 | -0.523 | -0.890 | -0.509 |
| O54734 | Ddost | 0.002 | 0.025 | 0.000 | 0.042 | -0.469 | 1.490 | 1.120 | 1.220 |
| Q9JIF0 | Prmt1 | 0.208 | 0.404 | -2.050 | 1.030 | 0.000 | -1.680 | -2.110 | -1.400 |
| Q149F5 | Tmem71 | 0.621 | 0.895 | -0.561 | 0.000 | 0.498 | -0.795 | -0.363 | 0.363 |
| Q9CXJ4 | Abcb8 | 0.014 | 0.074 | 0.000 | 0.642 | -0.065 | 1.350 | 1.020 | 1.280 |
| Q61142 | Spin1 | 0.161 | 0.339 | -0.044 | 0.000 | 0.623 | -0.259 | -0.903 | -0.017 |
| Q91WS0 | Cisd1 | 0.396 | 0.646 | -0.215 | 1.180 | 0.000 | 0.469 | 0.808 | 0.987 |
| O09174 | Amacr | 0.030 | 0.113 | 0.000 | 0.612 | -0.235 | 1.240 | 0.949 | 0.900 |
| P62852 | Rps25 | 0.696 | 0.972 | 0.125 | -0.126 | 0.000 | 0.233 | -0.473 | -0.035 |
| P60670 | Nploc4 | 0.740 | 1.000 | 1.190 | 0.000 | -0.041 | 0.721 | -0.556 | 0.388 |
| Q9JLN9 | Mtor | 0.116 | 0.271 | 0.095 | 0.000 | -0.092 | -0.022 | -0.391 | -0.444 |
| Q9D7N3 | Mrps9 | 0.122 | 0.280 | 0.267 | -0.306 | 0.000 | 1.060 | 0.047 | 1.140 |
| P22682 | Cbl | 0.000 | 0.013 | -0.024 | 0.000 | 0.154 | -1.190 | -1.550 | -1.380 |
| P35922 | Fmr1 | 0.469 | 0.729 | 0.174 | -0.169 | 0.000 | 0.691 | -1.830 | -0.615 |
| Q5SYD0 | Myo1d | 0.002 | 0.025 | 0.242 | 0.000 | -0.716 | 3.190 | 2.310 | 3.330 |
| Q6P9R1 | Ddx51 | 0.054 | 0.164 | 0.000 | 0.058 | -0.347 | -0.583 | -1.340 | -0.615 |
| Q8BWM0 | Ptges2 | 0.946 | 1.000 | -1.850 | 2.870 | 0.000 | 1.240 | 0.282 | -0.189 |
| Q8R5F7 | Ifih1 | 0.396 | 0.645 | 0.526 | 0.000 | -0.048 | 0.713 | -1.360 | -0.697 |
| P24788 | Cdk11b | 0.579 | 0.850 | 0.534 | -6.110 | 0.000 | 0.018 | -1.130 | -0.567 |

| | | | | | | | | | |
|---|---|---|---|---|---|---|---|---|---|
| Q64522 | Hist2h2ab | 0.685 | 0.962 | 0.000 | 0.243 | -0.083 | -0.414 | -0.437 | 0.562 |
| Q99PU8 | Dhx30 | 0.416 | 0.670 | 0.000 | 0.380 | -0.875 | 1.330 | -0.133 | -0.053 |
| Q9JL16 | Isg20 | 0.132 | 0.296 | 0.573 | 0.000 | -0.300 | -0.033 | -1.030 | -1.080 |
| Q8R035 | Ict1 | 0.016 | 0.077 | 0.201 | -0.053 | 0.000 | 1.010 | 0.503 | 0.676 |
| Q9CPT4 | Mydgf | 0.595 | 0.868 | -0.032 | 0.503 | 0.000 | 0.538 | 0.152 | -1.160 |
| Q8BK63 | Csnk1a1 | 0.143 | 0.310 | 0.250 | 0.000 | -0.233 | -0.046 | -0.568 | -0.775 |
| Q99020 | Hnrnpab | 0.020 | 0.088 | -0.016 | 0.000 | 0.002 | -0.493 | -0.788 | -0.309 |
| Q05144 | Rac2 | 0.029 | 0.112 | 0.254 | -0.036 | 0.000 | -0.369 | -1.240 | -0.844 |
| P97872 | Fmo5 | 0.485 | 0.748 | 0.755 | -0.101 | 0.000 | 0.639 | -0.667 | -0.436 |
| Q99J36 | Thumpd1 | 0.295 | 0.522 | -0.210 | 0.125 | 0.000 | -5.210 | -0.726 | -0.026 |
| Q7TMB8 | Cyfip1 | 0.108 | 0.259 | 0.000 | 0.124 | -1.260 | 0.703 | 0.315 | 0.700 |
| Q3URS9 | Ccdc51 | 0.007 | 0.048 | 0.194 | 0.000 | -0.221 | 0.635 | 0.568 | 0.670 |
| Q3KNM2 | Marchf5 | 0.008 | 0.055 | 0.127 | 0.000 | -0.352 | 1.370 | 0.744 | 1.400 |
| P62737 | Acta2 | 0.095 | 0.238 | 0.856 | 0.000 | -0.683 | 1.790 | 0.877 | 3.750 |
| P68033 | Actc1 | 0.095 | 0.238 | 0.856 | 0.000 | -0.683 | 1.790 | 0.877 | 3.750 |
| Q8C2K1 | Def6 | 0.035 | 0.125 | 0.000 | -0.029 | 0.144 | -0.322 | -0.929 | -0.442 |
| Q9CQW2 | Arl8b | 0.497 | 0.763 | 0.125 | 0.000 | -0.062 | 0.265 | -1.000 | -0.059 |
| P28798 | Grn | 0.144 | 0.311 | 0.111 | 0.000 | -1.300 | -0.647 | -1.980 | -2.100 |
| Q5DW34 | Ehmt1 | 0.432 | 0.687 | 0.000 | -0.729 | 0.184 | -0.148 | -1.100 | -0.351 |
| Q8BHZ4 | Znf592 | 0.469 | 0.728 | -0.184 | 0.000 | 0.011 | 0.150 | -1.000 | -0.161 |
| Q9ET30 | Tm9sf3 | 0.062 | 0.178 | 0.000 | 0.413 | -0.902 | 0.864 | 0.711 | 0.992 |
| Q8QZY9 | Sf3b4 | 0.531 | 0.798 | -2.010 | 0.478 | 0.000 | -3.370 | -0.510 | -0.237 |
| Q7TQH0 | Atxn2l | 0.095 | 0.238 | -0.117 | 0.000 | 0.068 | -0.604 | -0.867 | -0.096 |
| Q91W59 | Rbms1 | 0.033 | 0.121 | 0.806 | -0.066 | 0.000 | 2.090 | 0.979 | 1.920 |
| Q8VBV3 | Exosc2 | 0.039 | 0.133 | -0.349 | 0.471 | 0.000 | -0.627 | -0.957 | -0.651 |
| Q99LI2 | Clcc1 | 0.023 | 0.096 | -0.100 | 0.000 | 0.227 | 0.468 | 0.534 | 0.896 |
| Q9WUA3 | Pfkp | 0.033 | 0.119 | -0.074 | 0.000 | 0.023 | -0.325 | -1.040 | -0.682 |
| Q9DC70 | Ndufs7 | 0.005 | 0.040 | 0.000 | 0.110 | -0.184 | 0.684 | 0.466 | 0.575 |
| Q3UND0 | Skap2 | 0.206 | 0.403 | 0.518 | 0.000 | -0.624 | -0.258 | -0.599 | -1.130 |
| A6PWY4 | Wdr76 | 0.081 | 0.213 | 0.000 | -0.414 | 0.003 | -0.428 | -1.580 | -0.920 |
| O70494 | Sp3 | 0.078 | 0.208 | -0.010 | 0.000 | 0.263 | -0.254 | -1.040 | -0.332 |
| Q91X84 | Crtc3 | 0.778 | 1.000 | -1.050 | 0.000 | 0.635 | -0.708 | -0.435 | 0.214 |
| P84091 | Ap2m1 | 0.321 | 0.556 | 0.149 | 0.000 | -0.439 | 0.525 | 0.040 | -0.018 |
| P67871 | Csnk2b | 0.499 | 0.765 | 0.026 | -0.042 | 0.000 | 0.326 | -0.131 | 0.085 |
| P55258 | Rab8a | 0.478 | 0.740 | 0.000 | 0.081 | -1.520 | 0.007 | -0.248 | 0.041 |
| P48428 | Tbca | 0.000 | 0.003 | 0.052 | 0.000 | -0.010 | -1.980 | -2.160 | -2.200 |
| Q9CY27 | Tecr | 0.001 | 0.019 | 0.000 | 0.089 | -0.113 | 1.390 | 0.978 | 1.330 |
| Q9QZD9 | Eif3i | 0.115 | 0.269 | 0.000 | 0.168 | -0.055 | -0.088 | -1.120 | -0.527 |
| Q62158 | Trim27 | 0.431 | 0.687 | 0.111 | -0.657 | 0.000 | 0.009 | -1.310 | -0.441 |
| P26516 | Psmd7 | 0.010 | 0.059 | 0.000 | 0.058 | -0.009 | -0.723 | -0.864 | -0.382 |
| Q60634 | Flot2 | 0.111 | 0.264 | 0.405 | 0.000 | -0.394 | 1.280 | 0.318 | 0.643 |
| Q8CAS9 | Parp9 | 0.397 | 0.647 | 0.000 | -0.344 | 0.030 | 0.545 | -1.360 | -1.310 |
| Q91WQ3 | Yars | 0.766 | 1.000 | 0.202 | -0.103 | 0.000 | 0.408 | -0.347 | -0.198 |
| Q8CI32 | Bag5 | 0.037 | 0.129 | 0.971 | 0.000 | -0.099 | -0.561 | -1.150 | -0.939 |
| Q60749 | Khdrbs1 | 0.078 | 0.208 | -0.028 | 0.000 | 0.031 | -0.155 | -0.632 | -0.249 |
| Q99KG3 | Rbm10 | 0.006 | 0.046 | -0.268 | 0.000 | 0.234 | -0.923 | -1.030 | -0.744 |
| Q9EP89 | Lactb | 0.009 | 0.055 | 0.000 | 0.227 | -0.004 | 0.611 | 0.438 | 0.673 |
| Q9D3D9 | Atp5d | 0.126 | 0.287 | -0.224 | 0.095 | 0.000 | 0.573 | 0.056 | 0.267 |
| Q80W00 | Ppp1r10 | 0.269 | 0.487 | 0.000 | -0.002 | 0.060 | -0.042 | -1.260 | -0.145 |
| Q9DC48 | Cdc40 | 0.029 | 0.110 | 0.078 | 0.000 | -0.189 | -0.334 | -0.713 | -0.444 |
| Q9D0F6 | Rfc5 | 0.453 | 0.711 | 0.422 | 0.000 | -0.107 | 0.315 | -0.450 | -0.249 |
| E9Q6J5 | Bod1l | 0.006 | 0.047 | -0.187 | 0.000 | 0.048 | -0.937 | -1.340 | -0.761 |
| Q8BT60 | Cpne3 | 0.133 | 0.297 | 1.050 | -0.584 | 0.000 | -0.468 | -1.280 | -0.789 |
| Q9JL61 | Rfx5 | 0.105 | 0.252 | -0.505 | 0.086 | 0.000 | -1.050 | -0.889 | -0.307 |
| Q8C0L0 | Tmx4 | 0.533 | 0.800 | 0.070 | -0.116 | 0.000 | 0.113 | -0.118 | 0.169 |
| Q9CR67 | Tmem33 | 0.011 | 0.064 | 0.000 | 0.299 | -0.015 | 1.150 | 0.633 | 1.090 |
| O08912 | Galnt1 | 0.941 | 1.000 | 0.124 | -0.001 | 0.000 | 0.241 | -0.251 | 0.096 |
| O88384 | Vti1b | 0.769 | 1.000 | 0.080 | 0.000 | -0.234 | 0.387 | -0.713 | -0.140 |
| O88986 | Gcat | 0.640 | 0.917 | -0.398 | 0.000 | 0.308 | 0.250 | -0.442 | -0.349 |

| | | | | | | | | | |
|---|---|---|---|---|---|---|---|---|---|
| Q9WV80 | Snx1 | 0.053 | 0.162 | 0.124 | 0.000 | -0.011 | -0.162 | -0.776 | -0.438 |
| P70318 | Tial1 | 0.250 | 0.463 | -0.420 | 0.005 | 0.000 | -0.234 | -0.572 | -0.309 |
| Q9WU00 | Nrf1 | 0.250 | 0.462 | 0.000 | 0.342 | -0.076 | -0.339 | -0.325 | 0.125 |
| Q9JKX4 | Aatf | 0.350 | 0.592 | 0.000 | 0.709 | -1.030 | -0.247 | -0.828 | -1.010 |
| O54946 | Dnajb6 | 0.385 | 0.634 | -0.028 | 0.357 | 0.000 | -0.199 | -0.392 | 0.254 |
| Q6KCD5 | Nipbl | 0.198 | 0.392 | -0.044 | 0.000 | 0.205 | 0.043 | -0.828 | -0.282 |
| Q9EST5 | Anp32b | 0.003 | 0.033 | -0.081 | 0.000 | 0.074 | -0.928 | -1.530 | -1.580 |
| O35685 | Nudc | 0.046 | 0.149 | 0.279 | -0.199 | 0.000 | -0.351 | -1.340 | -0.968 |
| Q7TSC1 | Prrc2a | 0.109 | 0.260 | -0.311 | 0.000 | 0.082 | -0.383 | -0.282 | -0.324 |
| P59729 | Rin3 | 0.908 | 1.000 | 0.881 | -4.610 | 0.000 | -0.058 | -4.180 | -0.287 |
| O55234 | Psmb5 | 0.002 | 0.022 | 0.000 | -0.040 | 0.005 | -0.534 | -0.717 | -0.480 |
| Q9Z0P4 | Palm | 0.078 | 0.209 | 4.740 | 0.000 | -1.440 | 5.760 | 4.970 | 5.890 |
| Q9JHI5 | Ivd | 0.333 | 0.571 | 0.000 | 0.127 | -0.011 | 0.544 | -0.052 | 0.212 |
| Q922J9 | Far1 | 0.001 | 0.020 | 0.251 | 0.000 | -0.223 | 1.600 | 1.300 | 1.720 |
| Q8BLN5 | Lss | 0.000 | 0.011 | 0.000 | -0.076 | 0.465 | 3.020 | 2.690 | 3.120 |
| Q64685 | St6gal1 | 0.007 | 0.047 | -0.227 | 0.000 | 0.149 | -0.709 | -0.705 | -0.551 |
| O35864 | Cops5 | 0.075 | 0.202 | -0.163 | 0.000 | 0.098 | -1.120 | -1.010 | -3.290 |
| Q9CPP6 | Ndufa5 | 0.245 | 0.456 | -0.536 | 0.555 | 0.000 | 0.694 | 0.147 | 0.649 |
| P15532 | Nme1 | 0.004 | 0.038 | 0.000 | 0.373 | -0.032 | -0.827 | -1.420 | -1.230 |
| P70271 | Pdlim4 | 0.116 | 0.271 | 0.000 | 0.283 | -1.460 | 0.861 | 0.415 | 0.941 |
| Q9D0M1 | Prpsap1 | 0.155 | 0.330 | -0.116 | 0.000 | 0.125 | -0.059 | -2.380 | -1.090 |
| Q8CB77 | Tceb3 | 0.095 | 0.237 | -0.143 | 0.026 | 0.000 | -0.321 | -0.886 | -0.258 |
| Q99LD9 | Eif2b2 | 0.175 | 0.359 | 0.033 | -0.145 | 0.000 | -0.005 | -0.644 | -0.430 |
| P17879 | Hspa1b | 0.401 | 0.652 | 0.000 | -1.290 | 0.818 | 0.709 | 0.226 | 0.366 |
| Q61696 | Hspa1a | 0.401 | 0.652 | 0.000 | -1.290 | 0.818 | 0.709 | 0.226 | 0.366 |
| O88544 | Cops4 | 0.001 | 0.019 | -0.084 | 0.083 | 0.000 | -0.508 | -0.674 | -0.571 |
| O55125 | Nipsnap1 | 0.039 | 0.134 | -0.109 | 0.000 | 0.160 | 0.335 | 0.199 | 0.384 |
| Q8BZR9 | Ncbp3 | 0.042 | 0.139 | 0.000 | -0.357 | 0.139 | -0.557 | -1.920 | -1.740 |
| Q8BX10 | Pgam5 | 0.212 | 0.409 | -0.303 | 0.066 | 0.000 | 0.400 | -0.081 | 0.256 |
| Q8CG47 | Smc4 | 0.107 | 0.256 | 0.051 | 0.000 | -0.195 | -0.164 | -1.130 | -0.645 |
| Q9Z1T1 | Ap3b1 | 0.903 | 1.000 | 0.355 | 0.000 | -0.119 | 0.549 | -0.293 | 0.090 |
| P70245 | Ebp | 0.136 | 0.301 | -0.271 | 0.136 | 0.000 | 0.523 | 0.032 | 0.355 |
| P41105 | Rpl28 | 0.677 | 0.953 | 0.058 | -0.160 | 0.000 | 0.474 | -0.870 | -0.236 |
| Q8C5P7 | Tdrp | 0.289 | 0.514 | 0.000 | 0.540 | -3.900 | -0.646 | 0.680 | 3.180 |
| Q921G6 | Lrch4 | 0.128 | 0.290 | 0.000 | 0.144 | -0.354 | -0.491 | -0.524 | -1.780 |
| P48410 | Abcd1 | 0.804 | 1.000 | 0.401 | -0.082 | 0.000 | 0.518 | -0.244 | 0.258 |
| P07091 | S100a4 | 0.014 | 0.073 | -0.132 | 0.000 | 0.151 | -0.450 | -0.771 | -0.446 |
| Q9D1I2 | Card19 | 0.027 | 0.106 | -0.929 | 0.000 | 0.091 | 1.010 | 0.663 | 1.220 |
| Q99LI5 | Znf281 | 0.332 | 0.569 | -0.459 | 0.034 | 0.000 | -0.422 | -0.184 | -0.403 |
| Q99M28 | Rnps1 | 0.025 | 0.101 | 0.023 | 0.000 | -0.010 | -0.173 | -0.526 | -0.366 |
| P19253 | Rpl13a | 0.055 | 0.166 | 0.072 | -0.126 | 0.000 | 0.649 | 0.148 | 0.402 |
| Q9D0R4 | Ddx56 | 0.301 | 0.530 | -0.058 | 0.000 | 0.322 | 0.224 | -0.377 | -0.453 |
| Q9JJU8 | Sh3bgrl | 0.001 | 0.016 | -0.279 | 0.031 | 0.000 | -1.640 | -2.170 | -2.160 |
| Q9D753 | Exosc8 | 0.007 | 0.050 | -0.219 | 0.000 | 0.109 | -0.795 | -0.804 | -0.526 |
| Q8C8U0 | Ppfibp1 | 0.004 | 0.036 | -1.930 | 0.000 | 0.118 | 4.310 | 3.430 | 3.420 |
| P20491 | Fcer1g | 0.868 | 1.000 | 0.025 | 0.000 | -0.757 | 0.174 | -0.883 | -0.237 |
| P08226 | Apoe | 0.508 | 0.772 | 0.861 | -0.299 | 0.000 | 0.960 | 0.161 | 0.360 |
| Q62384 | Zpr1 | 0.001 | 0.019 | 0.000 | -0.018 | 0.224 | -1.120 | -1.420 | -1.640 |
| Q9DAV9 | Tmem38b | 0.014 | 0.073 | 0.128 | 0.000 | -0.142 | 0.489 | 0.347 | 0.671 |
| Q3UYC0 | Ppm1h | 0.126 | 0.287 | -0.049 | 0.465 | 0.000 | -0.111 | -0.156 | -0.485 |
| O35405 | Pld3 | 0.088 | 0.227 | 0.695 | -0.043 | 0.000 | -0.040 | -1.100 | -0.976 |
| P46061 | Rangap1 | 0.066 | 0.186 | -0.024 | 0.000 | 0.159 | -0.246 | -0.623 | -0.156 |
| Q6ZQH8 | Nup188 | 0.536 | 0.803 | 0.000 | 0.306 | -0.291 | -0.168 | -0.324 | 0.082 |
| Q9CQS8 | Sec61b | 0.002 | 0.025 | 0.034 | 0.000 | -0.122 | 1.150 | 0.720 | 0.971 |
| Q61686 | Cbx5 | 0.059 | 0.173 | 0.129 | -0.280 | 0.000 | -0.287 | -1.000 | -0.770 |
| P97480 | Eya3 | 0.133 | 0.296 | 0.000 | -0.166 | 0.589 | -0.090 | -2.020 | -0.889 |
| Q6A009 | Ltn1 | 0.985 | 1.000 | 0.052 | 0.000 | -0.178 | 0.434 | -0.683 | 0.102 |
| P12382 | Pfkl | 0.013 | 0.070 | -0.225 | 0.544 | 0.000 | -0.976 | -1.550 | -0.976 |
| Q8BGC0 | Htatsf1 | 0.081 | 0.213 | 0.000 | -0.444 | 0.487 | -1.040 | -4.100 | -1.620 |

| | | | | | | | | | |
|---|---|---|---|---|---|---|---|---|---|
| O35648 | Cetn3 | 0.061 | 0.176 | 0.000 | -0.110 | 0.216 | -0.144 | -0.637 | -0.458 |
| Q9R1K9 | Cetn2 | 0.007 | 0.048 | 0.000 | -0.087 | 0.218 | -0.638 | -1.140 | -0.782 |
| Q63844 | Mapk3 | 0.097 | 0.239 | 0.625 | -0.043 | 0.000 | -0.104 | -0.713 | -0.406 |
| Q8JZM7 | Cdc73 | 0.003 | 0.033 | 0.000 | 0.219 | -0.127 | -0.762 | -1.210 | -1.040 |
| P50429 | Arsb | 0.524 | 0.790 | 0.190 | -1.080 | 0.000 | -0.004 | -1.780 | -0.486 |
| Q3B7Z2 | Osbp | 0.301 | 0.529 | 0.000 | -0.645 | 0.139 | -0.287 | -1.600 | -0.361 |
| Q8BMC4 | Nop9 | 0.058 | 0.171 | -0.300 | 0.866 | 0.000 | -0.880 | -0.720 | -1.950 |
| P42228 | Stat4 | 0.004 | 0.034 | -0.425 | 0.000 | 0.128 | -1.440 | -1.590 | -1.150 |
| Q8VEH3 | Arl8a | 0.427 | 0.683 | 0.000 | 0.064 | -0.007 | 0.164 | -2.160 | 0.049 |
| Q8C7R4 | Uba6 | 0.049 | 0.154 | -0.131 | 0.000 | 0.079 | -0.551 | -1.370 | -0.519 |
| Q61216 | Mre11a | 0.103 | 0.249 | 0.000 | 0.005 | -0.020 | -0.225 | -0.399 | -0.043 |
| Q9CQT1 | Mri1 | 0.010 | 0.059 | -0.208 | 0.000 | 0.005 | -1.030 | -2.150 | -1.650 |
| Q99KK2 | Cmas | 0.118 | 0.274 | 0.000 | 0.039 | -0.028 | -0.189 | -0.483 | -0.065 |
| P58854 | Tubgcp3 | 0.950 | 1.000 | 0.322 | 0.000 | -0.040 | 0.316 | -0.178 | 0.107 |
| Q60854 | Serpinb6 | 0.023 | 0.096 | 0.000 | 0.176 | -0.003 | -0.265 | -0.736 | -0.439 |
| Q8K3K7 | Agpat2 | 0.136 | 0.300 | 0.673 | -0.697 | 0.000 | 0.782 | 0.680 | 0.732 |
| Q9D8S3 | Arfgap3 | 0.403 | 0.655 | 0.116 | -0.337 | 0.000 | 0.416 | -0.238 | 0.275 |
| O88942 | Nfatc1 | 0.007 | 0.049 | 0.000 | 0.025 | -0.018 | -0.426 | -0.651 | -0.342 |
| P30416 | Fkbp4 | 0.034 | 0.122 | 0.210 | -0.131 | 0.000 | -0.417 | -1.450 | -1.070 |
| Q9WTP6 | Ak2 | 0.009 | 0.057 | 0.187 | 0.000 | -0.011 | -0.394 | -0.833 | -0.616 |
| Q5I012 | Slc38a10 | 0.015 | 0.076 | 0.000 | 0.102 | -0.889 | 1.920 | 0.930 | 1.770 |
| Q8CIV2 | Tmem259 | 0.933 | 1.000 | 0.000 | -0.305 | 3.950 | 1.260 | 4.740 | -1.730 |
| Q8BG51 | Rhot1 | 0.015 | 0.076 | 0.000 | 0.146 | -0.220 | 0.857 | 0.556 | 0.462 |
| Q9ERI2 | Rab27a | 0.016 | 0.078 | -0.088 | 0.258 | 0.000 | -0.439 | -0.539 | -0.320 |
| Q8K327 | Champ1 | 0.072 | 0.197 | 0.000 | 0.682 | -0.760 | -1.110 | -0.962 | -1.050 |
| Q8K3Z9 | Pom121 | 0.636 | 0.912 | -0.467 | 0.020 | 0.000 | -0.303 | -0.455 | 1.110 |
| Q60759 | Gcdh | 0.450 | 0.708 | 0.000 | 0.355 | -0.371 | 0.773 | -0.053 | 0.092 |
| Q91X78 | Erlin1 | 0.006 | 0.047 | 0.506 | 0.000 | -0.214 | 1.620 | 1.150 | 1.490 |
| O08808 | Diaph1 | 0.008 | 0.053 | 0.000 | 0.331 | -0.056 | -0.629 | -1.230 | -1.130 |
| E9Q3L2 | Pi4ka | 0.023 | 0.096 | 0.000 | -0.187 | 0.098 | -0.443 | -1.100 | -0.794 |
| P47226 | Tes | 0.054 | 0.164 | -0.065 | 0.000 | 0.382 | -0.256 | -1.230 | -0.735 |
| P20108 | Prdx3 | 0.006 | 0.046 | 0.214 | -0.058 | 0.000 | 0.778 | 0.509 | 0.675 |
| P28076 | Psmb9 | 0.000 | 0.008 | 0.000 | 0.158 | -0.071 | -1.320 | -1.440 | -1.440 |
| Q922Q2 | Riok1 | 0.209 | 0.406 | 0.319 | -0.036 | 0.000 | 0.212 | -1.780 | -0.775 |
| Q3U2A8 | Vars2 | 0.089 | 0.227 | 0.000 | 0.110 | -0.376 | 0.144 | 0.368 | 0.307 |
| Q99NB8 | Ubqln4 | 0.918 | 1.000 | -0.973 | 0.205 | 0.000 | -0.248 | -0.431 | -0.210 |
| P61961 | Ufm1 | 0.109 | 0.260 | 0.161 | -0.247 | 0.000 | -0.168 | -0.732 | -0.429 |
| Q80YD1 | Supv3l1 | 0.046 | 0.149 | 0.000 | 0.195 | -0.039 | 0.533 | 0.318 | 0.254 |
| Q9WUM3 | Coro1b | 0.004 | 0.036 | 0.000 | 0.494 | -0.016 | -0.912 | -0.833 | -0.805 |
| P62245 | Rps15a | 0.129 | 0.291 | 0.000 | 0.085 | -0.316 | -0.152 | -0.630 | -0.945 |
| Q6ZWQ0 | Syne2 | 0.001 | 0.014 | -0.133 | 0.058 | 0.000 | -0.906 | -1.160 | -0.960 |
| Q9D162 | Ccdc167 | 0.095 | 0.237 | 0.528 | -0.276 | 0.000 | 1.370 | 0.371 | 2.040 |
| Q6NSR8 | Npepl1 | 0.008 | 0.053 | -0.164 | 0.000 | 0.177 | -0.998 | -1.940 | -1.350 |
| B1AY13 | Usp24 | 0.332 | 0.570 | 0.320 | -0.715 | 0.000 | 0.167 | -1.930 | -0.876 |
| Q6NZN0 | Rbm26 | 0.151 | 0.323 | 0.000 | -0.189 | 0.425 | -0.493 | -1.240 | -0.077 |
| O35737 | Hnrnph1 | 0.264 | 0.482 | -0.105 | 0.106 | 0.000 | 0.047 | -0.222 | -0.287 |
| Q01768 | Nme2 | 0.028 | 0.109 | 0.000 | 0.131 | -0.266 | -0.484 | -1.090 | -0.705 |
| Q9D8Y8 | Ing5 | 0.137 | 0.302 | -0.463 | 0.040 | 0.000 | -0.637 | -0.577 | -0.289 |
| Q921F4 | Hnrnpll | 0.081 | 0.213 | -0.224 | 0.000 | 0.360 | -0.213 | -0.501 | -0.736 |
| Q8K337 | Inpp5b | 0.001 | 0.019 | -0.129 | 0.000 | 0.150 | -1.020 | -1.310 | -1.430 |
| Q62077 | Plcg1 | 0.003 | 0.033 | -0.125 | 0.000 | 0.471 | -1.190 | -1.370 | -1.750 |
| Q9DBC0 | Selo | 0.169 | 0.350 | 0.118 | -0.027 | 0.000 | 0.290 | -2.830 | -2.140 |
| Q99KK9 | Hars2 | 0.327 | 0.564 | 0.000 | 0.815 | -0.658 | 0.593 | 0.065 | 1.500 |
| Q8BMQ2 | Gtf3c4 | 0.311 | 0.544 | -0.322 | 0.051 | 0.000 | -0.220 | -0.870 | -0.103 |
| Q99KP3 | Cryl1 | 0.002 | 0.027 | -0.510 | 0.000 | 0.271 | -1.670 | -2.060 | -2.120 |
| Q8BHG9 | Cggbp1 | 0.310 | 0.542 | -0.314 | 0.000 | 0.385 | -0.552 | -0.518 | 0.112 |
| Q3UDE2 | Ttll12 | 0.009 | 0.058 | -0.158 | 0.219 | 0.000 | -0.662 | -1.330 | -1.070 |
| P54822 | Adsl | 0.060 | 0.175 | -0.145 | 0.046 | 0.000 | -0.266 | -0.814 | -0.390 |
| P41242 | Matk | 0.064 | 0.183 | -0.128 | 0.000 | 0.032 | -0.458 | -1.140 | -0.374 |

| | | | | | | | | | |
|---|---|---|---|---|---|---|---|---|---|
| O54825 | Bysl | 0.040 | 0.134 | 0.000 | 0.199 | -0.082 | -0.345 | -0.495 | -0.172 |
| Q60591 | Nfatc2 | 0.930 | 1.000 | 0.183 | -1.590 | 0.000 | -0.316 | -0.458 | -0.478 |
| Q91V12 | Acot7 | 0.020 | 0.089 | -0.038 | 0.193 | 0.000 | 0.736 | 0.449 | 0.396 |
| Q8BWD8 | Cdk19 | 0.857 | 1.000 | -1.270 | 0.000 | 0.173 | -0.518 | -0.267 | -0.035 |
| Q8C5L3 | Cnot2 | 0.073 | 0.198 | -0.390 | 0.000 | 0.063 | -0.329 | -1.130 | -0.861 |
| Q6PEB6 | Mob4 | 0.779 | 1.000 | -0.331 | 0.004 | 0.000 | 0.295 | 0.251 | -0.595 |
| Q8BHF7 | Pgs1 | 0.017 | 0.080 | 0.000 | 1.120 | -0.175 | 2.030 | 1.790 | 2.010 |
| Q8K3J1 | Ndufs8 | 0.002 | 0.025 | 0.000 | 0.081 | -0.026 | 0.747 | 0.504 | 0.553 |
| Q64433 | Hspe1 | 0.001 | 0.018 | 0.000 | 0.100 | -0.080 | 1.090 | 0.779 | 0.991 |
| Q8C5N3 | Cwc22 | 0.023 | 0.096 | 0.000 | 0.191 | -0.464 | -0.905 | -2.000 | -1.370 |
| O70161 | Pip5k1c | 0.114 | 0.268 | -0.827 | 0.696 | 0.000 | 0.554 | 0.961 | 1.390 |
| Q80UJ7 | Rab3gap1 | 0.391 | 0.640 | -0.189 | 0.000 | 0.108 | 0.130 | -0.432 | -0.341 |
| Q80XP8 | Fam76b | 0.240 | 0.449 | 0.252 | -1.180 | 0.000 | 0.005 | -3.390 | -2.020 |
| Q3V1L4 | Nt5c2 | 0.047 | 0.149 | 0.000 | 0.109 | -0.392 | -0.430 | -1.090 | -0.869 |
| Q9CXG3 | Ppil4 | 0.025 | 0.103 | 0.000 | -0.294 | 0.128 | -0.723 | -1.650 | -0.961 |
| Q6ZWN5 | Rps9 | 0.784 | 1.000 | 0.205 | 0.000 | -0.149 | 0.324 | -0.286 | -0.170 |
| Q3ULJ0 | Gpd1l | 0.045 | 0.146 | 0.027 | 0.000 | -0.287 | -0.381 | -1.230 | -1.180 |
| O35215 | Ddt | 0.032 | 0.118 | -0.099 | 1.310 | 0.000 | -3.690 | -1.070 | -2.560 |
| Q9DB27 | Mcts1 | 0.959 | 1.000 | 0.152 | -0.168 | 0.000 | 0.247 | -0.337 | 0.042 |
| Q9WUP4 | Srd5a3 | 0.212 | 0.409 | 0.098 | -1.150 | 0.000 | 0.613 | -0.036 | 0.341 |
| Q9JKC8 | Ap3m1 | 0.127 | 0.288 | 0.404 | 0.000 | -0.143 | 0.882 | 0.358 | 0.385 |
| Q99KQ4 | Nampt | 0.129 | 0.291 | 0.000 | 0.160 | -0.599 | -1.710 | -1.790 | -0.193 |
| Q9WUA2 | Farsb | 0.086 | 0.223 | 0.000 | 0.165 | -0.086 | -0.105 | -0.380 | -0.183 |
| Q8K0D5 | Gfm1 | 0.253 | 0.467 | 0.000 | 0.253 | -0.248 | 1.020 | 0.078 | 0.217 |
| P62320 | Snrpd3 | 0.040 | 0.135 | 0.000 | 0.221 | -0.144 | -0.306 | -0.280 | -0.293 |
| P53026 | Rpl10a | 0.307 | 0.538 | 0.158 | -0.322 | 0.000 | 0.118 | -0.555 | -0.840 |
| P36993 | Ppm1b | 0.001 | 0.021 | 0.086 | -0.228 | 0.000 | -1.300 | -1.800 | -1.840 |
| Q61035 | Hars | 0.035 | 0.123 | -0.560 | 0.683 | 0.000 | -1.710 | -0.913 | -1.280 |
| Q8BP67 | Rpl24 | 0.126 | 0.288 | 0.000 | 0.147 | -0.328 | 0.301 | 0.143 | 0.228 |
| Q8BQZ4 | Ralgapb | 0.057 | 0.170 | -0.188 | 0.101 | 0.000 | -0.179 | -0.555 | -0.491 |
| Q99KN9 | Clint1 | 0.085 | 0.221 | 0.000 | -0.072 | 0.053 | -0.466 | -1.160 | -0.263 |
| Q6ZWV7 | Rpl35 | 0.412 | 0.665 | 0.100 | -0.032 | 0.000 | 0.532 | -0.166 | 0.271 |
| Q9QZL0 | Ripk3 | 0.408 | 0.660 | 0.000 | 0.733 | -0.238 | 0.654 | -1.470 | -0.577 |
| Q62219 | Tgfb1i1 | 0.001 | 0.022 | -0.852 | 0.592 | 0.000 | 5.540 | 3.950 | 5.150 |
| Q99M51 | Nck1 | 0.034 | 0.122 | 0.044 | 0.000 | -0.172 | -1.730 | -2.880 | -0.932 |
| Q8K157 | Galm | 0.001 | 0.018 | -0.472 | 0.286 | 0.000 | -2.590 | -2.270 | -2.830 |
| Q6R891 | Ppp1r9b | 0.007 | 0.048 | -0.300 | 0.179 | 0.000 | -1.040 | -1.590 | -1.050 |
| Q8VE97 | Srsf4 | 0.080 | 0.211 | 0.065 | -0.149 | 0.000 | -0.350 | -2.830 | -2.070 |
| P97868 | Rbbp6 | 0.093 | 0.234 | 0.000 | -0.157 | 0.004 | 0.164 | 0.016 | 0.278 |
| Q9R1J0 | Nsdhl | 0.000 | 0.010 | 0.212 | 0.000 | -0.154 | 1.860 | 1.730 | 1.980 |
| Q9JLQ2 | Git2 | 0.003 | 0.033 | -0.185 | 0.261 | 0.000 | -0.937 | -1.420 | -1.150 |
| Q99K51 | Pls3 | 0.002 | 0.027 | 0.000 | 0.943 | -0.084 | 3.190 | 2.810 | 3.720 |
| P54818 | Galc | 0.358 | 0.603 | 0.640 | -0.012 | 0.000 | 0.282 | -0.448 | -0.145 |
| P33609 | Pola1 | 0.060 | 0.176 | 0.058 | 0.000 | -0.834 | -1.570 | -1.450 | -0.752 |
| O70591 | Pfdn2 | 0.026 | 0.103 | 0.000 | 0.302 | -0.526 | -0.767 | -1.420 | -1.340 |
| Q60875 | Arhgef2 | 0.124 | 0.284 | 0.011 | -0.367 | 0.000 | -0.122 | -2.670 | -3.400 |
| Q66JX5 | Fgfr1op | 0.672 | 0.949 | 0.516 | 0.000 | -0.404 | -0.396 | -2.760 | 1.530 |
| P49117 | Nr2c2 | 0.046 | 0.147 | -0.415 | 0.000 | 0.288 | -1.000 | -1.560 | -0.578 |
| Q5EBH1 | Rassf5 | 0.008 | 0.052 | -0.317 | 0.000 | 0.413 | -1.520 | -2.820 | -2.000 |
| Q99N94 | Mrpl9 | 0.093 | 0.234 | 0.142 | 0.000 | -0.854 | 1.030 | 0.602 | 0.221 |
| Q8C3X2 | Ccdc90b | 0.005 | 0.040 | -0.005 | 0.303 | 0.000 | 1.660 | 1.020 | 1.250 |
| P81117 | Nucb2 | 0.002 | 0.027 | 0.376 | 0.000 | -0.305 | 1.870 | 1.440 | 1.640 |
| O54788 | Dffb | 0.056 | 0.166 | 0.000 | 1.160 | -0.052 | -1.010 | -0.828 | -0.474 |
| Q9QY76 | Vapb | 0.449 | 0.707 | 0.020 | 0.000 | -0.313 | 0.435 | 0.068 | -0.237 |
| P58404 | Strn4 | 0.482 | 0.745 | 0.000 | -0.039 | 0.280 | -0.045 | -0.554 | 0.248 |
| Q9CQI7 | Snrpb2 | 0.009 | 0.058 | 0.000 | -0.133 | 0.059 | -0.549 | -0.991 | -0.617 |
| Q9CQN4 | Sostdc1 | 0.262 | 0.478 | -0.797 | 0.000 | 0.018 | -1.030 | -0.651 | -0.382 |
| Q9JKV1 | Adrm1 | 0.232 | 0.436 | 0.000 | -0.776 | 0.108 | 0.423 | -4.760 | -2.820 |
| Q9JHC9 | Elf2 | 0.083 | 0.216 | -0.198 | 0.039 | 0.000 | -0.443 | -1.150 | -0.367 |

| | | | | | | | | | |
|---|---|---|---|---|---|---|---|---|---|
| P63087 | Ppp1cc | 0.072 | 0.197 | -0.142 | 0.000 | 0.013 | -0.169 | -0.780 | -0.518 |
| P70371 | Terf1 | 0.025 | 0.102 | 0.000 | 0.395 | -0.619 | -1.120 | -1.050 | -1.160 |
| P46467 | Vps4b | 0.061 | 0.177 | 0.037 | -0.210 | 0.000 | -0.384 | -1.500 | -0.859 |
| Q924A2 | Cic | 0.368 | 0.615 | 0.000 | -0.386 | 0.805 | -0.032 | -0.150 | -0.582 |
| P55012 | Slc12a2 | 0.024 | 0.099 | 0.000 | -2.210 | 1.390 | 3.800 | 3.040 | 3.760 |
| Q8R5A3 | Apbb1ip | 0.010 | 0.059 | 0.149 | 0.000 | -0.079 | -0.505 | -1.010 | -1.070 |
| Q9JM90 | Stap1 | 0.167 | 0.348 | 0.120 | -0.071 | 0.000 | -0.139 | -0.977 | -0.217 |
| Q99N87 | Mrps5 | 0.000 | 0.012 | 0.039 | 0.000 | -0.040 | 1.140 | 0.934 | 0.888 |
| Q8VDF2 | Uhrf1 | 0.486 | 0.749 | 0.987 | 0.000 | -0.817 | 0.598 | -0.873 | -1.370 |
| Q91W50 | Csde1 | 0.837 | 1.000 | 0.000 | 1.520 | -0.584 | 0.018 | -0.602 | 1.010 |
| Q6IRU5 | Cltb | 0.010 | 0.059 | 0.000 | -0.318 | 0.052 | -0.644 | -0.946 | -0.715 |
| Q8R3B7 | Brd8 | 0.126 | 0.286 | -0.614 | 0.116 | 0.000 | -0.398 | -2.410 | -1.310 |
| P25799 | Nfkb1 | 0.017 | 0.080 | 0.000 | -0.065 | 0.139 | -0.430 | -1.120 | -0.909 |
| Q9WTX8 | Mad1l1 | 0.041 | 0.136 | 0.000 | -0.010 | 0.350 | -0.176 | -0.603 | -0.413 |
| Q09XV5 | Chd8 | 0.586 | 0.857 | 0.000 | -1.090 | 0.094 | -0.128 | -0.357 | 0.228 |
| Q923G2 | Polr2h | 0.104 | 0.252 | -0.057 | 0.000 | 0.218 | -0.239 | -2.030 | -0.901 |
| Q61555 | Fbn2 | 0.000 | 0.013 | -0.574 | 0.551 | 0.000 | 3.680 | 3.550 | 3.750 |
| Q921Y2 | Imp3 | 0.195 | 0.387 | 0.000 | 0.390 | -0.130 | -0.035 | -0.345 | -0.201 |
| Q8C5W3 | Tbcel | 0.029 | 0.112 | 0.000 | 0.915 | -0.754 | -2.390 | -1.250 | -2.060 |
| Q8JZX4 | Rbm17 | 0.033 | 0.121 | 0.000 | 0.276 | -0.213 | -0.865 | -0.990 | -0.361 |
| Q9ER38 | Tor3a | 0.003 | 0.032 | 0.000 | 0.258 | -0.172 | 0.998 | 0.891 | 1.220 |
| P51881 | Slc25a5 | 0.203 | 0.400 | 0.000 | 1.090 | -0.121 | 0.477 | 1.260 | 1.440 |
| O88627 | Slc28a2 | 0.316 | 0.549 | 0.079 | -0.550 | 0.000 | -0.170 | -1.540 | -0.377 |
| Q8K4L0 | Ddx54 | 0.274 | 0.495 | 0.000 | -0.315 | 0.208 | 0.291 | -1.620 | -0.988 |
| Q8R1S0 | Coq6 | 0.178 | 0.364 | 0.000 | 0.621 | -0.148 | 0.936 | 0.515 | 0.414 |
| E2JF22 | Piezo1 | 0.024 | 0.098 | 0.000 | -2.270 | 2.080 | 4.710 | 3.970 | 4.720 |
| Q5SV85 | Synrg | 0.000 | 0.008 | 0.080 | -0.246 | 0.000 | 5.810 | 5.470 | 6.420 |
| Q9DBL7 | Coasy | 0.134 | 0.298 | 7.440 | -0.978 | 0.000 | 7.090 | 7.330 | 7.010 |
| P83870 | Phf5a | 0.029 | 0.110 | -0.244 | 0.000 | 0.161 | -0.640 | -1.280 | -0.627 |
| Q5HZI9 | Slc25a51 | 0.018 | 0.083 | -0.031 | 0.632 | 0.000 | 1.200 | 0.956 | 1.100 |
| Q91VC9 | Ghitm | 0.003 | 0.030 | 0.000 | 0.088 | -0.147 | 1.130 | 0.700 | 0.953 |
| Q8BGB5 | Limd2 | 0.130 | 0.293 | 0.000 | -0.039 | 0.115 | -0.066 | -0.503 | -0.158 |
| Q8CAK1 | Iba57 | 0.322 | 0.558 | 0.036 | 0.000 | -0.449 | 0.439 | -0.205 | 0.176 |
| Q4QQM4 | Trp53i11 | 0.420 | 0.674 | 0.000 | 0.090 | -0.068 | -0.104 | -0.396 | 0.108 |
| Q3UFY0 | Rrp36 | 0.228 | 0.431 | -0.058 | 0.250 | 0.000 | -0.563 | -0.442 | 0.160 |
| P59999 | Arpc4 | 0.531 | 0.798 | 0.000 | 0.139 | -0.184 | 0.125 | 0.054 | -0.015 |
| Q9D6K8 | Fundc2 | 0.013 | 0.071 | 0.000 | 0.474 | -0.255 | 1.690 | 1.410 | 0.958 |
| Q8BRT1 | Clasp2 | 0.206 | 0.402 | -0.569 | 0.000 | 0.003 | -0.061 | -1.620 | -1.150 |
| P62827 | Ran | 0.003 | 0.031 | -0.041 | 0.044 | 0.000 | -0.674 | -1.100 | -0.771 |
| Q9Z108 | Stau1 | 0.006 | 0.046 | 0.086 | 0.000 | -0.256 | 1.060 | 0.598 | 1.010 |
| P28867 | Prkcd | 0.874 | 1.000 | 0.163 | 0.000 | -1.170 | -0.357 | 0.299 | -1.260 |
| Q9ET54 | Palld | 0.003 | 0.031 | 0.370 | -0.667 | 0.000 | 2.810 | 2.010 | 2.300 |
| O08807 | Prdx4 | 0.256 | 0.470 | 0.038 | 0.000 | -0.002 | 0.439 | -0.070 | 0.262 |
| Q9QZ23 | Nfu1 | 0.022 | 0.094 | 0.000 | 0.235 | -0.727 | 1.080 | 0.755 | 1.350 |
| Q3UEB3 | Puf60 | 0.005 | 0.039 | 0.000 | -0.093 | 0.127 | -0.739 | -1.290 | -0.937 |
| Q8C4B4 | Unc119b | 0.448 | 0.706 | -0.956 | 0.211 | 0.000 | 0.115 | -0.456 | -2.910 |
| Q9JKB1 | Uchl3 | 0.006 | 0.045 | 0.000 | 0.255 | -0.128 | -1.230 | -2.330 | -1.720 |
| Q9D906 | Atg7 | 0.013 | 0.070 | 0.028 | -0.171 | 0.000 | -0.567 | -1.170 | -0.790 |
| P51175 | Ppox | 0.909 | 1.000 | 0.000 | 0.082 | -0.711 | 0.651 | -0.730 | -0.373 |
| P54775 | Psmc4 | 0.018 | 0.083 | -0.123 | 0.000 | 0.106 | -0.391 | -0.884 | -0.580 |
| Q9ES56 | Trappc4 | 0.865 | 1.000 | 0.000 | 0.560 | -0.421 | -0.618 | 0.358 | 0.174 |
| P51163 | Uros | 0.072 | 0.196 | 0.559 | -0.181 | 0.000 | -0.372 | -1.170 | -2.130 |
| P11440 | Cdk1 | 0.243 | 0.453 | 0.534 | 0.000 | -1.540 | 1.200 | -0.031 | 0.779 |
| Q810B6 | Ankfy1 | 0.392 | 0.641 | -0.006 | 0.356 | 0.000 | 0.165 | 0.066 | -1.070 |
| P52019 | Sqle | 0.002 | 0.027 | 0.000 | 1.740 | -0.441 | 5.210 | 5.320 | 6.150 |
| Q9JHR7 | Ide | 0.004 | 0.036 | 0.000 | 0.341 | -0.020 | -0.764 | -0.819 | -0.576 |
| Q91W53 | Golga7 | 0.087 | 0.224 | 0.117 | 0.000 | -0.062 | 0.932 | 0.100 | 0.924 |
| Q8BUK6 | Hook3 | 0.569 | 0.837 | 0.000 | -0.177 | 0.034 | 0.322 | -0.911 | -0.229 |
| O88447 | Klc1 | 0.910 | 1.000 | 0.000 | -0.262 | 0.051 | 0.542 | -0.641 | 0.015 |

| | | | | | | | | | |
|---|---|---|---|---|---|---|---|---|---|
| Q9D1P0 | Mrpl13 | 0.522 | 0.787 | 0.000 | 0.348 | -0.702 | 0.959 | -0.679 | 0.596 |
| Q8BU31 | Rap2c | 0.351 | 0.592 | 0.427 | -4.150 | 0.000 | 0.508 | -0.089 | 0.526 |
| Q99104 | Myo5a | 0.027 | 0.107 | -0.006 | 0.000 | 0.152 | 0.704 | 1.170 | 1.920 |
| Q9CQ80 | Vps25 | 0.220 | 0.420 | 0.000 | 0.351 | -0.161 | 0.672 | 0.002 | 1.430 |
| Q8K1M6 | Dnm1l | 0.892 | 1.000 | 0.042 | 0.000 | -0.313 | 0.147 | -0.185 | -0.310 |
| Q9D2R0 | Aacs | 0.237 | 0.444 | 1.080 | 0.000 | -0.323 | 1.580 | 0.944 | 0.464 |
| O88271 | Cfdp1 | 0.005 | 0.043 | 0.000 | -0.146 | 0.287 | -1.330 | -2.440 | -2.380 |
| P83882 | Rpl36a | 0.645 | 0.922 | 0.260 | 0.000 | -0.007 | 0.602 | -0.206 | 0.227 |
| P46638 | Rab11b | 0.020 | 0.089 | 0.000 | 0.117 | -0.097 | -0.270 | -0.495 | -0.285 |
| Q9ERG2 | Strn3 | 0.489 | 0.754 | 1.350 | -1.770 | 0.000 | 1.130 | -0.551 | 1.530 |
| Q6ZQL4 | Wdr43 | 0.934 | 1.000 | 0.170 | 0.000 | -0.109 | 0.209 | -0.182 | -0.003 |
| P23249 | Mov10 | 0.371 | 0.618 | 0.000 | 0.062 | -0.014 | 0.456 | -0.062 | 0.118 |
| P00493 | Hprt1 | 0.004 | 0.038 | 0.000 | 0.060 | -0.208 | -1.260 | -1.950 | -1.240 |
| O88351 | Ikbkb | 0.730 | 1.000 | 0.314 | -0.360 | 0.000 | 0.876 | -0.396 | -0.052 |
| Q64191 | Aga | 0.034 | 0.122 | 0.000 | 0.123 | -1.280 | 0.991 | 1.240 | 0.958 |
| Q9D517 | Agpat3 | 0.027 | 0.107 | 0.000 | 0.348 | -0.252 | 0.753 | 0.547 | 0.673 |
| Q8R5H1 | Usp15 | 0.003 | 0.031 | 0.364 | 0.000 | -0.025 | -1.680 | -2.580 | -1.690 |
| Q62193 | Rpa2 | 0.053 | 0.161 | -4.040 | 0.454 | 0.000 | -5.270 | -6.020 | -4.510 |
| Q8K4X7 | Agpat4 | 0.005 | 0.042 | 0.000 | 0.300 | -0.338 | 1.030 | 0.953 | 1.130 |
| P31996 | Cd68 | 0.988 | 1.000 | 0.659 | -0.005 | 0.000 | 0.757 | -0.122 | 0.002 |
| Q6PIP5 | Nudcd1 | 0.042 | 0.139 | 0.000 | -0.718 | 0.138 | -0.872 | -2.330 | -1.830 |
| Q9QYS9 | Qki | 0.249 | 0.462 | -0.069 | 0.000 | 0.024 | -0.109 | -0.787 | -0.085 |
| Q9Z2M7 | Pmm2 | 0.001 | 0.021 | -0.080 | 0.000 | 0.173 | -1.300 | -1.910 | -1.880 |
| Q9CZL5 | Pcbd2 | 0.058 | 0.171 | 0.028 | 0.000 | -0.711 | 0.373 | 0.407 | 0.453 |
| Q9QXA5 | Lsm4 | 0.002 | 0.028 | 0.000 | -0.110 | 0.040 | -0.957 | -1.370 | -0.916 |
| Q9Z266 | Snapin | 0.175 | 0.359 | 0.011 | 0.000 | -0.270 | -0.152 | -0.396 | -0.918 |
| P63166 | Sumo1 | 0.032 | 0.119 | -0.103 | 0.000 | 0.363 | -0.366 | -0.411 | -0.342 |
| P62806 | Hist1h4a | 0.776 | 1.000 | 0.000 | 0.764 | -0.074 | -0.043 | -0.259 | 0.643 |
| Q99KD5 | Unc45a | 0.428 | 0.684 | -0.049 | 0.000 | 0.165 | 0.289 | -0.589 | -0.285 |
| Q9CQR6 | Ppp6c | 0.166 | 0.347 | 0.000 | 0.129 | -1.020 | -1.250 | -0.873 | -0.759 |
| Q8R4K2 | Irak4 | 0.016 | 0.078 | 0.152 | 0.000 | -0.178 | -0.685 | -1.670 | -1.410 |
| P48771 | Cox7a2 | 0.957 | 1.000 | -0.456 | 0.000 | 0.309 | 0.089 | -0.409 | 0.125 |
| Q8R0J7 | Vps37b | 0.293 | 0.520 | 0.183 | -0.042 | 0.000 | 0.116 | -0.393 | -0.175 |
| P47964 | Rpl36 | 0.019 | 0.085 | 0.146 | -0.148 | 0.000 | 0.783 | 0.357 | 0.578 |
| P23298 | Prkch | 0.210 | 0.407 | -0.013 | 0.000 | 0.048 | -0.075 | -0.813 | -0.138 |
| Q8C4Y3 | Nelfb | 0.269 | 0.488 | -0.182 | 0.000 | 0.311 | 0.143 | -1.250 | -0.417 |
| Q9CQC9 | Sar1b | 0.173 | 0.356 | 0.127 | -0.333 | 0.000 | -0.158 | -1.010 | -0.454 |
| Q80U78 | Pum1 | 0.834 | 1.000 | 0.000 | -0.142 | 0.437 | 0.833 | -0.404 | -0.438 |
| Q91ZJ5 | Ugp2 | 0.039 | 0.133 | 0.000 | 0.158 | -0.579 | -0.976 | -1.240 | -0.692 |
| Q9ER64 | Osbpl5 | 0.463 | 0.721 | -0.879 | 0.118 | 0.000 | -0.307 | -0.060 | 0.631 |
| P03911 | Mtnd4 | 0.026 | 0.105 | -1.040 | 0.000 | 0.181 | 0.916 | 1.730 | 1.150 |
| Q9EP72 | Emc7 | 0.060 | 0.175 | 0.022 | 0.000 | -0.211 | 1.430 | 0.205 | 1.070 |
| Q9CY66 | Gar1 | 0.549 | 0.818 | 0.000 | 0.455 | -0.523 | -0.108 | -0.504 | -0.073 |
| Q7TMI3 | Uhrf2 | 0.019 | 0.087 | -0.049 | 0.200 | 0.000 | -0.215 | -0.398 | -0.482 |
| Q8R332 | Nup58 | 0.087 | 0.224 | 0.000 | 0.032 | -0.337 | -0.313 | -1.300 | -0.783 |
| Q9JL56 | Gde1 | 0.140 | 0.306 | -1.780 | 0.755 | 0.000 | 0.971 | 0.895 | 1.310 |
| Q60767 | Ly75 | 0.069 | 0.192 | -0.111 | 0.422 | 0.000 | -0.712 | -1.810 | -0.459 |
| Q9D6L8 | Ppil3 | 0.024 | 0.098 | 0.014 | -0.545 | 0.000 | -0.749 | -0.893 | -0.939 |
| E9PZM4 | Chd2 | 0.002 | 0.022 | 0.000 | 0.221 | -0.104 | -0.784 | -0.978 | -1.070 |
| Q925J9 | Med1 | 0.684 | 0.960 | 0.315 | -0.113 | 0.000 | 0.119 | -0.251 | 0.102 |
| Q7TQ95 | Lnp | 0.459 | 0.719 | 0.255 | -0.105 | 0.000 | 0.500 | -0.269 | 0.679 |
| O70435 | Psma3 | 0.004 | 0.035 | 0.000 | 0.181 | -0.215 | -0.858 | -1.230 | -1.360 |
| Q9QYF9 | Ndrg3 | 0.462 | 0.721 | 0.109 | -0.092 | 0.000 | 0.101 | -2.550 | 0.247 |
| P97427 | Crmp1 | 0.006 | 0.046 | -0.518 | 0.000 | 0.065 | -2.230 | -2.140 | -1.360 |
| P27782 | Lef1 | 0.652 | 0.928 | -1.410 | 0.000 | 1.930 | 0.644 | -0.499 | -1.250 |
| Q6P9R4 | Arhgef18 | 0.035 | 0.124 | -0.095 | 0.000 | 0.552 | -0.603 | -1.540 | -0.742 |
| Q8BL74 | Gtf3c2 | 0.500 | 0.765 | -0.311 | 0.113 | 0.000 | -0.136 | -0.581 | 0.027 |
| Q61207 | Psap | 0.175 | 0.359 | 0.324 | -0.624 | 0.000 | -0.131 | -1.780 | -1.130 |
| P70404 | Idh3g | 0.775 | 1.000 | 0.000 | 0.070 | -0.248 | 0.756 | -0.504 | -0.078 |

| | | | | | | | | | |
|---|---|---|---|---|---|---|---|---|---|
| Q8VE92 | Rbm4b | 0.900 | 1.000 | 0.000 | 0.638 | -0.243 | 0.413 | -0.485 | 0.642 |
| Q9CPU0 | Glo1 | 0.000 | 0.014 | -0.265 | 0.169 | 0.000 | -1.470 | -1.690 | -1.770 |
| Q9CYI4 | Luc7l | 0.092 | 0.233 | 0.092 | 0.000 | -0.204 | -0.138 | -1.260 | -1.150 |
| Q9R1Q9 | Atp6ap1 | 0.509 | 0.773 | 0.000 | -3.800 | 1.510 | 2.510 | -2.850 | 3.560 |
| Q9D3G5 | Stx11 | 0.059 | 0.174 | -0.116 | 0.240 | 0.000 | -0.192 | -0.385 | -0.238 |
| Q6P4S8 | Ints1 | 0.654 | 0.930 | -0.251 | 0.000 | 0.398 | -0.260 | 0.115 | 0.830 |
| Q9WTJ4 | Fiz1 | 0.019 | 0.085 | -0.768 | 0.551 | 0.000 | -2.790 | -4.340 | -2.030 |
| Q8BYU6 | Tor1aip2 | 0.011 | 0.064 | -0.119 | 0.338 | 0.000 | 2.410 | 1.340 | 2.900 |
| Q8VDI9 | Alg9 | 0.565 | 0.834 | 0.000 | 0.345 | -1.230 | 0.923 | -0.370 | -0.251 |
| Q8R4X3 | Rbm12 | 0.025 | 0.101 | 0.114 | -0.295 | 0.000 | -0.754 | -1.930 | -1.300 |
| P97822 | Anp32e | 0.017 | 0.080 | -0.110 | 0.199 | 0.000 | -1.090 | -1.940 | -0.873 |
| Q9D6Y7 | Msra | 0.068 | 0.189 | 0.549 | -0.217 | 0.000 | -0.252 | -0.768 | -0.814 |
| O55013 | Trappc3 | 0.197 | 0.390 | -0.068 | 0.000 | 0.058 | 0.042 | -0.786 | -0.387 |
| Q5DTM8 | Rnf20 | 0.061 | 0.177 | -0.043 | 0.000 | 0.212 | -0.235 | -0.631 | -0.198 |
| P40630 | Tfam | 0.889 | 1.000 | 0.483 | 0.000 | -0.272 | 0.049 | 0.003 | 0.264 |
| O35166 | Gosr2 | 0.289 | 0.514 | 0.000 | -0.790 | 0.687 | 0.295 | 0.517 | 0.710 |
| Q8BVL3 | Snx17 | 0.151 | 0.322 | 0.000 | 1.180 | -1.990 | 1.400 | 1.220 | 1.500 |
| Q9WVE8 | Pacsin2 | 0.460 | 0.719 | 0.260 | 0.000 | -0.091 | 0.564 | -0.113 | 0.262 |
| Q9CR00 | Psmd9 | 0.000 | 0.011 | -0.220 | 0.000 | 0.049 | -1.150 | -1.140 | -1.140 |
| O88508 | Dnmt3a | 0.201 | 0.397 | -0.367 | 0.018 | 0.000 | -0.065 | -1.310 | -0.720 |
| P97386 | Lig3 | 0.578 | 0.849 | 0.068 | 0.000 | -0.173 | 0.026 | -0.596 | 0.058 |
| Q571H0 | Urb1 | 0.375 | 0.623 | 0.847 | 0.000 | -0.023 | 0.911 | -0.805 | -1.810 |
| P57080 | Usp25 | 0.109 | 0.260 | 0.084 | -0.165 | 0.000 | -0.069 | -1.320 | -1.140 |
| P62141 | Ppp1cb | 0.218 | 0.417 | 0.078 | -0.044 | 0.000 | 0.122 | -0.514 | -0.526 |
| Q9R0P6 | Sec11a | 0.066 | 0.186 | 0.000 | 0.620 | -0.132 | 0.704 | 0.707 | 0.867 |
| Q8R422 | Cd109 | 0.002 | 0.025 | 0.000 | -1.010 | 1.720 | 6.180 | 6.070 | 6.910 |
| Q9DBA9 | Gtf2h1 | 0.488 | 0.752 | -0.068 | 1.270 | 0.000 | 0.286 | 0.054 | -0.181 |
| P58058 | Nadk | 0.361 | 0.606 | 1.210 | 0.000 | -0.691 | -0.082 | -0.474 | -0.743 |
| Q4VAE3 | Tmem65 | 0.014 | 0.072 | 0.000 | 0.487 | -0.336 | 1.190 | 1.030 | 1.570 |
| Q91Z38 | Ttc1 | 0.021 | 0.091 | -0.290 | 1.550 | 0.000 | -1.560 | -1.660 | -1.960 |
| P56135 | Atp5j2 | 0.005 | 0.040 | 0.000 | 0.393 | -0.087 | 0.924 | 1.130 | 0.968 |
| Q6P9Q6 | Fkbp15 | 0.205 | 0.402 | 0.000 | 0.346 | -0.021 | 0.306 | -1.450 | -0.962 |
| Q8C0D5 | Efl1 | 0.065 | 0.184 | 0.000 | 0.237 | -0.014 | -0.080 | -0.766 | -0.729 |
| Q8R1N0 | Znf830 | 0.022 | 0.093 | -0.006 | 0.000 | 0.395 | -0.705 | -1.570 | -0.748 |
| Q62348 | Tsn | 0.000 | 0.003 | -0.097 | 0.000 | 0.053 | -1.470 | -1.540 | -1.490 |
| Q6PE01 | Snrnp40 | 0.016 | 0.078 | 0.000 | 0.172 | -0.032 | -0.475 | -0.884 | -0.420 |
| Q9DCM2 | Gstk1 | 0.701 | 0.977 | 0.010 | 0.000 | -0.212 | 0.377 | -0.571 | -0.378 |
| Q9BCZ4 | Vimp | 0.732 | 1.000 | -0.386 | 0.179 | 0.000 | 0.206 | -0.445 | -0.249 |
| P02469 | Lamb1 | 0.000 | 0.010 | 0.000 | -0.970 | 0.049 | 4.590 | 4.580 | 4.770 |
| Q9QUM4 | Slamf1 | 0.023 | 0.096 | 0.042 | -0.099 | 0.000 | -0.350 | -0.725 | -0.965 |
| Q9D0C4 | Trmt5 | 0.109 | 0.260 | -0.090 | 0.320 | 0.000 | -0.850 | -4.670 | -1.540 |
| Q8K4D3 | Slc36a1 | 0.353 | 0.594 | 0.603 | 0.000 | -0.040 | 0.838 | 0.183 | 0.432 |
| Q8BGZ4 | Cdc23 | 0.969 | 1.000 | -0.048 | 0.000 | 0.477 | 0.290 | -0.073 | 0.187 |
| Q8BYA0 | Tbcd | 0.073 | 0.198 | 0.208 | -0.046 | 0.000 | -0.191 | -1.560 | -1.040 |
| Q9CU65 | Zmym2 | 0.023 | 0.097 | 0.139 | 0.000 | -0.153 | -0.563 | -1.080 | -0.512 |
| Q9D6N5 | Drap1 | 0.026 | 0.104 | 0.000 | -0.142 | 0.076 | -0.451 | -1.270 | -0.887 |
| P70268 | Pkn1 | 0.139 | 0.305 | 0.296 | -0.083 | 0.000 | 0.064 | -1.320 | -0.873 |
| Q8K268 | Abcf3 | 0.662 | 0.938 | 0.000 | 0.314 | -0.079 | 0.491 | -0.526 | -0.185 |
| P18181 | Cd48 | 0.140 | 0.306 | -0.550 | 0.111 | 0.000 | -1.040 | -0.896 | -0.252 |
| O35704 | Sptlc1 | 0.322 | 0.557 | 0.000 | 0.610 | -0.610 | 1.960 | -0.450 | 1.180 |
| Q60848 | Hells | 0.176 | 0.360 | 0.617 | -0.364 | 0.000 | 0.051 | -1.150 | -0.971 |
| Q91VU0 | Fam3c | 0.497 | 0.763 | -0.324 | 0.000 | 0.112 | 0.324 | -0.226 | 0.155 |
| Q80UP3 | Dgkz | 0.013 | 0.070 | 0.041 | 0.000 | -0.034 | -0.490 | -1.080 | -0.681 |
| Q9CWK3 | Cd2bp2 | 0.003 | 0.030 | 0.000 | -0.074 | 0.360 | -0.959 | -1.200 | -0.874 |
| O35954 | Pitpnm1 | 0.016 | 0.079 | -0.371 | 0.600 | 0.000 | -0.920 | -1.220 | -1.360 |
| P52432 | Polr1c | 0.009 | 0.058 | 0.073 | -0.049 | 0.000 | -0.297 | -0.468 | -0.254 |
| E9QAM5 | Helz2 | 0.272 | 0.492 | 0.165 | 0.000 | -0.002 | 0.296 | -0.785 | -0.678 |
| Q9R117 | Tyk2 | 0.498 | 0.763 | -0.303 | 1.680 | 0.000 | 1.680 | -1.290 | 5.770 |
| O09126 | Sema4d | 0.389 | 0.638 | 0.000 | -0.504 | 0.016 | -0.443 | -0.502 | -0.137 |

| | | | | | | | | | |
|---|---|---|---|---|---|---|---|---|---|
| Q8C5L6 | Inpp5k | 0.406 | 0.658 | 0.323 | -0.057 | 0.000 | 0.022 | -0.757 | 0.149 |
| Q9D1M4 | Eef1e1 | 0.658 | 0.934 | 0.000 | 0.081 | -0.123 | 0.368 | -0.741 | -0.135 |
| P08556 | Nras | 0.188 | 0.378 | -0.332 | 0.239 | 0.000 | 0.065 | 0.343 | 0.475 |
| Q8R3C0 | Mcmbp | 0.017 | 0.081 | 0.000 | 0.437 | -0.374 | -0.773 | -1.280 | -1.210 |
| P16332 | Mut | 0.015 | 0.076 | 0.000 | 0.403 | -0.314 | 1.300 | 0.791 | 1.140 |
| P98195 | Atp9b | 0.441 | 0.697 | 0.290 | 0.000 | -1.440 | 0.627 | -0.514 | 0.370 |
| Q9WU84 | Ccs | 0.004 | 0.036 | -0.087 | 0.000 | 0.263 | -0.735 | -1.230 | -1.050 |
| Q8BG15 | Ctdspl2 | 0.161 | 0.339 | 0.158 | 0.000 | 0.000 | -0.044 | -1.070 | -0.326 |
| Q7TNP2 | Ppp2r1b | 0.648 | 0.925 | 0.000 | -1.640 | 0.062 | -0.091 | -1.450 | -1.050 |
| Q9QZH3 | Ppie | 0.006 | 0.045 | -0.384 | 0.000 | 0.099 | -1.000 | -1.100 | -0.840 |
| Q9D0B6 | Pbdc1 | 0.040 | 0.134 | -0.101 | 0.317 | 0.000 | -0.847 | -1.480 | -0.430 |
| Q8VDD8 | Wash1 | 0.001 | 0.014 | 0.000 | -0.181 | 0.148 | -2.180 | -2.970 | -2.460 |
| Q9CWF2 | Tubb2b | 0.013 | 0.070 | -1.480 | 0.205 | 0.000 | 2.340 | 1.630 | 2.050 |
| Q924L1 | Letmd1 | 0.672 | 0.950 | -1.210 | 0.847 | 0.000 | -1.630 | 0.966 | -1.040 |
| P58871 | Tnks1bp1 | 0.036 | 0.125 | 0.000 | 0.067 | -0.956 | 0.608 | 0.849 | 0.825 |
| Q9JLR1 | Sec61a2 | 0.001 | 0.015 | 0.743 | -0.345 | 0.000 | 3.980 | 3.380 | 3.910 |
| Q8JZR0 | Acsl5 | 0.110 | 0.261 | -0.081 | 0.000 | 0.085 | 0.658 | 0.086 | 0.323 |
| Q80UU2 | Rpp38 | 0.005 | 0.039 | 0.284 | 0.000 | -0.181 | -0.693 | -0.891 | -0.837 |
| Q9D8S4 | Rexo2 | 0.107 | 0.257 | -0.191 | 0.000 | 0.052 | 0.356 | 0.007 | 0.431 |
| Q7TSQ8 | Pdpr | 0.166 | 0.347 | 0.000 | 0.234 | -0.035 | 1.070 | 0.066 | 0.599 |
| Q99LD4 | Gps1 | 0.116 | 0.271 | 0.000 | -0.608 | 0.075 | -0.392 | -2.690 | -1.650 |
| Q9D7B6 | Acad8 | 0.170 | 0.353 | 0.000 | -1.970 | 0.001 | 1.380 | 0.114 | 0.357 |
| Q9D2E2 | Toe1 | 0.038 | 0.129 | -0.230 | 0.099 | 0.000 | -0.498 | -1.300 | -0.711 |
| Q8BGK6 | Slc7a6 | 0.020 | 0.088 | 0.473 | 0.000 | -0.733 | 1.870 | 1.030 | 1.780 |
| Q3U319 | Rnf40 | 0.939 | 1.000 | -0.282 | 0.018 | 0.000 | -0.232 | -0.673 | 0.745 |
| Q8BWW9 | Pkn2 | 0.014 | 0.073 | 0.067 | -0.265 | 0.000 | 1.650 | 0.718 | 1.050 |
| Q69ZX6 | Morc2a | 0.031 | 0.116 | 0.000 | 0.070 | -0.068 | -0.314 | -1.090 | -0.935 |
| Q8BMG7 | Rab3gap2 | 0.815 | 1.000 | 0.974 | 0.000 | -2.280 | -0.344 | -1.730 | -0.051 |
| P52332 | Jak1 | 0.177 | 0.363 | 0.099 | -0.263 | 0.000 | 0.484 | 0.026 | 0.174 |
| Q8K370 | Acad10 | 0.053 | 0.162 | 0.324 | 0.000 | -0.339 | 0.796 | 0.651 | 0.382 |
| Q9ERG0 | Lima1 | 0.016 | 0.078 | -1.030 | 0.000 | 1.290 | 3.320 | 2.760 | 2.660 |
| P70677 | Casp3 | 0.003 | 0.030 | -0.016 | 0.138 | 0.000 | -1.190 | -2.040 | -1.570 |
| P70303 | Ctps2 | 0.915 | 1.000 | -1.140 | 0.507 | 0.000 | -0.358 | -0.806 | 0.335 |
| Q8C079 | Strip1 | 0.020 | 0.089 | 0.000 | 0.023 | -0.001 | -0.227 | -0.556 | -0.309 |
| O35963 | Rab33b | 0.738 | 1.000 | 0.000 | 0.257 | -0.073 | 0.208 | -0.033 | 0.142 |
| Q91V01 | Lpcat3 | 0.000 | 0.014 | 0.081 | 0.000 | -0.276 | 1.120 | 1.080 | 1.090 |
| Q3U308 | Ctu2 | 0.630 | 0.906 | -0.090 | 0.000 | 0.201 | 0.290 | -0.324 | -0.173 |
| Q9R0N0 | Galk1 | 0.017 | 0.082 | 0.220 | 0.000 | -0.038 | -0.664 | -1.340 | -0.646 |
| Q6TEK5 | Vkorc1l1 | 0.114 | 0.268 | 0.438 | 0.000 | -0.745 | 0.971 | 0.277 | 1.730 |
| P51612 | Xpc | 0.055 | 0.166 | -0.178 | 0.000 | 0.301 | -0.793 | -1.460 | -0.385 |
| Q8VI93 | Oas3 | 0.907 | 1.000 | -1.180 | 1.910 | 0.000 | -0.757 | 1.090 | 0.796 |
| Q9R0L7 | Akap8l | 0.946 | 1.000 | -0.713 | 0.000 | 5.100 | 1.220 | -0.563 | 3.270 |
| A2ALW5 | Dnajc25 | 0.089 | 0.227 | 0.000 | 0.099 | -0.136 | 0.114 | 1.400 | 1.090 |
| Q8BX80 | Engase | 0.040 | 0.135 | 0.000 | 0.000 | -0.421 | -0.479 | -1.130 | -0.991 |
| O35345 | Kpna6 | 0.521 | 0.787 | 0.000 | 0.117 | -0.022 | 0.282 | -0.333 | -0.277 |
| Q6P9R2 | Oxsr1 | 0.403 | 0.653 | 0.000 | -0.051 | 0.401 | 0.108 | -0.789 | 0.103 |
| Q8BX70 | Vps13c | 0.314 | 0.547 | -0.587 | 4.430 | 0.000 | -1.540 | -0.507 | 0.164 |
| Q9D2X5 | Mau2 | 0.343 | 0.582 | 1.700 | -0.299 | 0.000 | 0.721 | -2.370 | -0.480 |
| Q9CQZ6 | Ndufb3 | 0.002 | 0.025 | -0.037 | 0.054 | 0.000 | 0.800 | 0.535 | 0.580 |
| Q8VEH8 | Erlec1 | 0.003 | 0.030 | -0.264 | 0.815 | 0.000 | 2.390 | 2.290 | 2.460 |
| Q9DBS9 | Osbpl3 | 0.028 | 0.108 | -0.747 | 0.622 | 0.000 | -1.900 | -2.520 | -1.220 |
| Q9ET22 | Dpp7 | 0.022 | 0.093 | 0.280 | -0.651 | 0.000 | 1.120 | 0.745 | 1.040 |
| Q7TT00 | Supt20h | 0.529 | 0.796 | -0.259 | 0.000 | 0.989 | -0.214 | -1.420 | 0.814 |
| P55264 | Adk | 0.048 | 0.152 | 0.000 | -0.071 | 0.107 | -0.916 | -2.270 | -0.795 |
| Q99J45 | Nrbp1 | 0.236 | 0.444 | 0.000 | 0.419 | -0.402 | 0.398 | -1.600 | -2.240 |
| Q99MD9 | Nasp | 0.010 | 0.058 | 0.000 | 0.025 | -0.302 | -0.880 | -1.720 | -1.430 |
| Q61183 | Papola | 0.003 | 0.031 | -0.236 | 0.000 | 0.015 | -2.470 | -1.820 | -3.070 |
| Q9JK38 | Gnpnat1 | 0.020 | 0.087 | -0.428 | 0.206 | 0.000 | -1.870 | -4.330 | -2.580 |
| Q9EQN3 | Tsc22d4 | 0.102 | 0.247 | -0.175 | 0.000 | 0.444 | -0.414 | -1.580 | -0.445 |

| | | | | | | | | | |
|---|---|---|---|---|---|---|---|---|---|
| Q64523 | Hist2h2ac | 0.516 | 0.779 | 0.000 | -4.270 | 2.780 | 1.090 | -7.740 | -1.910 |
| Q6GSS7 | Hist2h2aa1 | 0.516 | 0.779 | 0.000 | -4.270 | 2.780 | 1.090 | -7.740 | -1.910 |
| Q8C5L7 | Rbm34 | 0.393 | 0.641 | 0.000 | -0.513 | 0.047 | -0.005 | -1.680 | -0.339 |
| Q61164 | Ctcf | 0.396 | 0.646 | 0.000 | -0.032 | 0.363 | 0.254 | -1.530 | -0.016 |
| Q9EQW7 | Kif13a | 0.113 | 0.266 | 0.000 | -0.297 | 0.102 | -0.141 | -3.210 | -2.580 |
| P45878 | Fkbp2 | 0.153 | 0.325 | 0.467 | -0.204 | 0.000 | 0.711 | 0.217 | 0.684 |
| P05063 | Aldoc | 0.181 | 0.368 | 0.123 | -0.392 | 0.000 | -0.459 | -0.168 | -0.728 |
| Q08122 | Tle3 | 0.005 | 0.040 | -0.384 | 0.000 | 0.107 | -1.320 | -2.140 | -1.610 |
| Q8VEJ4 | Nle1 | 0.656 | 0.932 | 0.581 | 0.000 | -0.057 | 0.782 | -1.120 | 0.012 |
| Q9D6M3 | Slc25a22 | 0.001 | 0.016 | 0.000 | 0.088 | -0.280 | 1.510 | 1.170 | 1.410 |
| Q6PGB6 | Naa50 | 0.083 | 0.216 | 0.000 | 1.470 | -0.508 | -1.190 | -2.400 | -0.760 |
| Q9ERE7 | Mesdc2 | 0.075 | 0.203 | 0.218 | -0.024 | 0.000 | 0.875 | 0.330 | 0.366 |
| P97808 | Fxyd5 | 0.556 | 0.825 | 0.040 | -0.208 | 0.000 | 0.153 | -0.912 | -0.054 |
| Q9D2R6 | Coa3 | 0.017 | 0.080 | 0.000 | 0.304 | -0.248 | 1.120 | 0.611 | 0.904 |
| Q6PDK2 | Kmt2d | 0.835 | 1.000 | -0.075 | 0.000 | 0.729 | 0.995 | -3.110 | 1.750 |
| O35955 | Psmb10 | 0.003 | 0.033 | -0.486 | 0.139 | 0.000 | -1.520 | -1.850 | -1.380 |
| Q9D666 | Sun1 | 0.011 | 0.064 | -0.159 | 0.054 | 0.000 | 0.644 | 0.375 | 0.368 |
| Q69ZQ2 | Isy1 | 0.308 | 0.540 | 0.000 | 0.024 | -0.086 | -0.353 | -4.570 | -0.172 |
| Q9CQM9 | Glrx3 | 0.670 | 0.948 | 0.000 | 0.737 | -0.831 | 0.182 | -0.361 | -0.620 |
| Q3UHD9 | Agap2 | 0.512 | 0.776 | 0.000 | -1.590 | 2.090 | 1.140 | 0.229 | 1.570 |
| P62869 | Tceb2 | 0.005 | 0.040 | -0.160 | 0.000 | 0.361 | -0.924 | -1.230 | -0.867 |
| Q8VE18 | Smg8 | 0.067 | 0.186 | 0.000 | -0.252 | 0.155 | -0.327 | -0.869 | -0.432 |
| P59997 | Kdm2a | 0.046 | 0.148 | -0.142 | 0.000 | 0.321 | -0.327 | -0.528 | -0.298 |
| Q5U4D9 | Thoc6 | 0.253 | 0.467 | -0.007 | 0.107 | 0.000 | 0.133 | -0.430 | -0.299 |
| Q61790 | Lag3 | 0.676 | 0.952 | -1.220 | 1.200 | 0.000 | -1.810 | -0.508 | 0.885 |
| Q8BMA6 | Srp68 | 0.366 | 0.613 | 0.286 | -0.009 | 0.000 | 0.374 | -0.726 | -0.412 |
| P47934 | Crat | 0.395 | 0.644 | 0.000 | -3.600 | 0.145 | 2.180 | -1.640 | 0.743 |
| Q9R0M6 | Rab9a | 0.599 | 0.872 | 0.489 | -0.023 | 0.000 | 1.030 | -0.266 | 0.401 |
| Q91WK1 | Spryd4 | 0.045 | 0.146 | 0.000 | 0.231 | -0.354 | 0.951 | 0.398 | 0.566 |
| Q9CQE1 | Nipsnap3b | 0.179 | 0.365 | 0.000 | 0.874 | -0.677 | 0.769 | 0.963 | 0.692 |
| Q9WVD5 | Slc25a15 | 0.005 | 0.042 | -0.508 | 0.364 | 0.000 | 1.270 | 1.750 | 1.660 |
| Q8BFU2 | Hist3h2a | 0.777 | 1.000 | 0.000 | -4.410 | 1.560 | 0.303 | -3.670 | -1.420 |
| Q8CGP5 | Hist1h2af | 0.777 | 1.000 | 0.000 | -4.410 | 1.560 | 0.303 | -3.670 | -1.420 |
| Q8CGP7 | Hist1h2ak | 0.777 | 1.000 | 0.000 | -4.410 | 1.560 | 0.303 | -3.670 | -1.420 |
| Q6DID3 | Scaf8 | 0.074 | 0.200 | -0.313 | 0.005 | 0.000 | -0.483 | -1.190 | -0.490 |
| Q4FZC9 | Syne3 | 0.324 | 0.559 | 0.000 | -0.003 | 0.647 | 0.218 | -0.099 | -0.456 |
| Q7TQK4 | Exosc3 | 0.740 | 1.000 | 0.915 | 0.000 | -1.500 | 0.287 | -2.560 | 0.408 |
| Q3U3R4 | Lmf1 | 0.048 | 0.151 | 0.000 | -1.190 | 0.317 | 1.070 | 0.891 | 1.090 |
| Q8VCS3 | Fam20b | 0.046 | 0.148 | -0.069 | 0.014 | 0.000 | 0.362 | 0.123 | 0.555 |
| P63328 | Ppp3ca | 0.969 | 1.000 | 0.252 | -1.670 | 0.000 | -0.681 | -0.504 | -0.307 |
| Q99LG2 | Tnpo2 | 0.144 | 0.311 | -0.517 | 0.405 | 0.000 | -0.599 | -0.596 | -0.407 |
| P34902 | Il2rg | 0.090 | 0.230 | -0.469 | 0.000 | 0.113 | -0.901 | -0.440 | -0.531 |
| O70422 | Gtf2h4 | 0.073 | 0.198 | 0.000 | -0.232 | 0.120 | -0.256 | -0.742 | -0.400 |
| Q9R059 | Fhl3 | 0.007 | 0.049 | 0.173 | -0.055 | 0.000 | 1.130 | 0.644 | 0.797 |
| Q99N69 | Lpxn | 0.148 | 0.318 | -1.040 | 0.000 | 0.363 | -0.818 | -1.570 | -0.895 |
| Q921E6 | Eed | 0.268 | 0.487 | 0.000 | -0.754 | 0.004 | -0.297 | -1.060 | -0.684 |
| Q8BVW3 | Trim14 | 0.019 | 0.086 | -0.022 | 0.118 | 0.000 | -0.230 | -0.342 | -0.147 |
| Q5SU73 | Coil | 0.142 | 0.309 | -0.437 | 0.214 | 0.000 | -0.490 | -1.940 | -0.572 |
| P97360 | Etv6 | 0.033 | 0.119 | 0.065 | -0.481 | 0.000 | -0.573 | -0.940 | -1.020 |
| Q8BFR4 | Gns | 0.030 | 0.113 | 0.302 | -0.538 | 0.000 | 1.030 | 0.604 | 0.845 |
| Q91Z53 | Grhpr | 0.060 | 0.176 | 0.000 | -0.087 | 0.077 | -0.125 | -0.652 | -0.656 |
| P11688 | Itga5 | 0.002 | 0.028 | 1.160 | -0.159 | 0.000 | 3.840 | 3.150 | 3.570 |
| P97494 | Gclc | 0.814 | 1.000 | 0.000 | -3.610 | 0.176 | -1.950 | -2.290 | -0.243 |
| Q9DCC4 | Pycrl | 0.014 | 0.073 | -0.043 | 0.149 | 0.000 | -0.894 | -0.823 | -0.369 |
| Q9WTK3 | Gpaa1 | 0.096 | 0.239 | 0.000 | 0.450 | -0.747 | 0.795 | 0.416 | 1.080 |
| Q61165 | Slc9a1 | 0.476 | 0.736 | 0.000 | 3.350 | -0.438 | 1.690 | 2.590 | 1.560 |
| P35285 | Rab22a | 0.166 | 0.347 | 0.077 | -0.430 | 0.000 | -0.291 | -0.396 | -0.563 |
| O08759 | Ube3a | 0.729 | 1.000 | -1.650 | 0.000 | 0.109 | -0.266 | -1.150 | -0.824 |
| P55937 | Golga3 | 0.555 | 0.824 | 0.743 | -0.155 | 0.000 | 0.761 | -0.700 | -0.507 |

| | | | | | | | | | |
|---|---|---|---|---|---|---|---|---|---|
| Q8CGF7 | Tcerg1 | 0.003 | 0.030 | 0.000 | -0.052 | 0.036 | -0.456 | -0.710 | -0.493 |
| Q8R0A0 | Gtf2f2 | 0.054 | 0.164 | 0.000 | 0.383 | -0.040 | -0.234 | -1.020 | -0.550 |
| Q69ZR2 | Hectd1 | 0.209 | 0.406 | 0.000 | -0.236 | 0.376 | 0.342 | -1.490 | -2.030 |
| P01887 | B2m | 0.002 | 0.024 | -0.085 | 0.041 | 0.000 | -0.362 | -0.494 | -0.399 |
| Q3TZX8 | Nol9 | 0.039 | 0.134 | 0.000 | 0.630 | -0.031 | -0.350 | -0.702 | -0.502 |
| P15864 | Hist1h1c | 0.098 | 0.241 | 0.000 | -0.140 | 0.151 | -0.276 | -0.471 | -0.109 |
| P70218 | Map4k1 | 0.001 | 0.015 | -0.001 | 0.000 | 0.020 | -0.901 | -1.290 | -1.110 |
| O55142 | Rpl35a | 0.064 | 0.183 | 0.103 | 0.000 | -0.035 | 0.678 | 0.149 | 0.445 |
| Q9D7H3 | RtcA | 0.331 | 0.569 | 0.000 | 0.509 | -0.242 | -0.137 | -0.464 | 0.003 |
| O08585 | Clta | 0.023 | 0.095 | 0.196 | 0.000 | -0.151 | -0.400 | -0.895 | -0.573 |
| Q00651 | Itga4 | 0.060 | 0.176 | -0.013 | 0.000 | 0.220 | -0.247 | -0.705 | -0.212 |
| Q9JHQ5 | Lztfl1 | 0.249 | 0.461 | 0.000 | 1.550 | -1.860 | -0.883 | -2.190 | -1.500 |
| Q8CI33 | Cwf19l1 | 0.057 | 0.168 | 0.000 | -0.338 | 0.204 | -0.426 | -1.560 | -1.040 |
| Q9DBD5 | Pelp1 | 0.585 | 0.857 | 0.000 | 0.295 | -0.055 | 0.379 | -0.345 | -0.239 |
| Q8BPB0 | Mob1b | 0.065 | 0.184 | 0.002 | -0.022 | 0.000 | -1.070 | -0.971 | -0.160 |
| Q921Y0 | Mob1a | 0.065 | 0.184 | 0.002 | -0.022 | 0.000 | -1.070 | -0.971 | -0.160 |
| Q91VJ4 | Stk38 | 0.126 | 0.288 | 0.000 | 0.140 | -0.080 | -0.206 | -1.220 | -0.366 |
| Q91VA6 | Poldip2 | 0.121 | 0.278 | 0.000 | 0.211 | -1.280 | 1.200 | 0.544 | 0.335 |
| Q8R307 | Vps18 | 0.451 | 0.708 | 0.576 | 0.000 | -2.330 | -5.690 | 0.344 | -1.420 |
| Q9Z1P6 | Ndufa7 | 0.004 | 0.037 | -0.044 | 0.000 | 0.010 | 1.000 | 0.554 | 0.732 |
| O35492 | Clk3 | 0.094 | 0.236 | 0.396 | 0.000 | -0.283 | -0.169 | -1.650 | -1.240 |
| Q9CQY5 | Magt1 | 0.003 | 0.030 | 0.204 | 0.000 | -0.242 | 1.620 | 1.090 | 1.260 |
| Q80UK8 | Ints2 | 0.770 | 1.000 | 0.000 | 0.667 | -0.462 | 0.070 | -0.378 | 0.167 |
| Q9JJA7 | Ccnl2 | 0.068 | 0.188 | -0.146 | 0.098 | 0.000 | -1.860 | -1.480 | -4.980 |
| Q91VT1 | Nsmce2 | 0.108 | 0.258 | 0.022 | 0.000 | -0.060 | -0.181 | -1.100 | -0.457 |
| Q9QZA0 | Ca5b | 0.641 | 0.917 | -0.511 | 1.310 | 0.000 | 0.880 | 0.859 | -3.100 |
| P53798 | Fdft1 | 0.001 | 0.020 | 0.237 | -0.533 | 0.000 | 3.080 | 2.240 | 2.750 |
| Q8CGC6 | Rbm28 | 0.512 | 0.776 | 0.000 | 0.234 | -0.130 | 0.198 | -0.353 | -0.157 |
| Q91XC9 | Pex16 | 0.174 | 0.359 | 0.000 | 0.199 | -0.204 | -0.082 | -0.321 | -0.286 |
| Q8BJU0 | Sgta | 0.034 | 0.122 | 0.181 | -0.136 | 0.000 | -0.501 | -1.020 | -0.422 |
| Q80YQ2 | Med23 | 0.492 | 0.756 | -0.693 | 0.000 | 0.423 | -1.010 | -4.040 | 1.240 |
| P58281 | Opa1 | 0.111 | 0.263 | -0.010 | 0.052 | 0.000 | -0.089 | -0.563 | -0.194 |
| Q9D1K2 | Atp6v1f | 0.511 | 0.776 | 0.318 | 0.000 | -0.373 | 1.120 | -0.116 | -0.082 |
| Q9QY36 | Naa10 | 0.267 | 0.486 | -0.260 | 0.090 | 0.000 | -0.029 | -1.070 | -0.333 |
| Q03526 | Itk | 0.005 | 0.041 | 0.620 | -0.355 | 0.000 | -2.150 | -1.480 | -1.840 |
| Q9D0L8 | Rnmt | 0.050 | 0.155 | -0.425 | 0.000 | 0.004 | -0.704 | -1.880 | -1.010 |
| Q8VDT9 | Mrpl50 | 0.179 | 0.365 | 0.195 | -2.670 | 0.000 | 0.890 | 0.464 | 0.716 |
| Q6ZWY3 | Rps27l | 0.220 | 0.419 | 0.023 | -0.195 | 0.000 | 0.934 | -0.035 | 0.227 |
| P70700 | Polr1b | 0.494 | 0.759 | -2.100 | 0.525 | 0.000 | -0.634 | -1.670 | -1.200 |
| A2RSY6 | Trmt1l | 0.025 | 0.102 | 0.000 | 0.306 | -0.067 | -0.295 | -0.736 | -0.522 |
| Q8K215 | Lyrm4 | 0.019 | 0.086 | 0.000 | 0.224 | -0.231 | 0.718 | 0.439 | 0.602 |
| Q8VE11 | Mtmr6 | 0.246 | 0.457 | 0.271 | -0.215 | 0.000 | 0.096 | -0.464 | -0.672 |
| P35601 | Rfc1 | 0.381 | 0.631 | 0.361 | -0.017 | 0.000 | 0.169 | -0.405 | -0.034 |
| P68134 | Acta1 | 0.003 | 0.033 | 0.940 | -1.600 | 0.000 | 5.180 | 4.230 | 4.750 |
| Q8BTV2 | Cpsf7 | 0.081 | 0.213 | 0.276 | -0.116 | 0.000 | -0.122 | -0.689 | -0.432 |
| Q9R1P0 | Psma4 | 0.005 | 0.039 | 0.000 | -0.149 | 0.029 | -0.825 | -1.490 | -1.300 |
| Q6NXI6 | Rprd2 | 0.317 | 0.551 | -1.120 | 0.219 | 0.000 | -2.430 | -0.277 | -0.826 |
| O88958 | Gnpda1 | 0.039 | 0.133 | -0.415 | 0.245 | 0.000 | -1.350 | -1.430 | -0.531 |
| P97814 | Pstpip1 | 0.021 | 0.091 | 0.000 | -0.470 | 0.035 | -0.725 | -1.330 | -1.030 |
| Q9CR20 | Ier3ip1 | 0.008 | 0.054 | -0.005 | 0.000 | 0.262 | 1.340 | 0.984 | 0.751 |
| P61963 | Dcaf7 | 0.926 | 1.000 | -0.608 | 0.000 | 0.679 | -0.850 | -0.695 | 1.910 |
| Q9JLZ3 | Auh | 0.243 | 0.453 | 0.028 | 0.000 | -0.092 | 0.540 | -0.170 | 0.570 |
| Q9DBC3 | Cmtr1 | 0.022 | 0.093 | -0.160 | 0.062 | 0.000 | -0.408 | -0.824 | -0.464 |
| Q8BGS7 | Cept1 | 0.152 | 0.323 | -0.085 | 0.000 | 0.517 | 0.219 | 1.040 | 0.812 |
| Q9JLV1 | Bag3 | 0.004 | 0.038 | -0.462 | 0.056 | 0.000 | 1.980 | 1.160 | 1.520 |
| Q61462 | Cyba | 0.727 | 1.000 | 0.833 | -0.144 | 0.000 | 0.806 | -0.516 | -0.163 |
| Q8BGA9 | Oxa1l | 0.039 | 0.133 | 0.000 | -1.550 | 0.191 | 1.400 | 1.020 | 1.330 |
| O88286 | Wiz | 0.633 | 0.909 | 0.000 | -0.611 | 0.289 | -0.058 | -1.210 | 0.168 |
| Q91WK2 | Eif3h | 0.199 | 0.394 | 0.008 | 0.000 | -0.211 | -0.065 | -0.983 | -0.428 |

| | | | | | | | | | |
|---|---|---|---|---|---|---|---|---|---|
| Q9CZE3 | Rab32 | 0.582 | 0.853 | 0.695 | 0.000 | -0.235 | 0.973 | 0.064 | 0.146 |
| Q3UMW8 | Cln5 | 0.192 | 0.384 | 0.765 | -0.088 | 0.000 | 0.123 | -0.679 | -0.468 |
| Q8BUY5 | Timmdc1 | 0.012 | 0.067 | 0.000 | 0.616 | -0.099 | 1.080 | 1.290 | 1.470 |
| B2RRE7 | Otud4 | 0.049 | 0.154 | 0.191 | -0.090 | 0.000 | -0.294 | -1.270 | -0.795 |
| Q62018 | Ctr9 | 0.075 | 0.202 | 0.000 | 0.236 | -0.610 | 0.292 | 0.707 | 0.680 |
| Q9CZV5 | Supt7l | 0.575 | 0.845 | -0.216 | 0.417 | 0.000 | -1.000 | 0.236 | 0.159 |
| Q9EQ06 | Hsd17b11 | 0.029 | 0.111 | 0.144 | 0.000 | -0.254 | 0.493 | 0.301 | 0.386 |
| Q9ERB0 | Snap29 | 0.440 | 0.697 | 0.870 | -3.570 | 0.000 | 0.006 | 0.284 | 0.520 |
| Q9WV70 | Noc2l | 0.688 | 0.964 | 1.540 | -3.470 | 0.000 | 0.186 | -0.279 | -4.600 |
| P18572 | Bsg | 0.144 | 0.311 | 0.286 | 0.000 | -0.498 | 1.030 | 0.217 | 0.378 |
| Q8R5L3 | Vps39 | 0.681 | 0.957 | 0.177 | 0.000 | -0.224 | 0.208 | -0.321 | -0.195 |
| Q99JX3 | Gorasp2 | 0.318 | 0.551 | 0.543 | -0.132 | 0.000 | -0.070 | -1.110 | 0.110 |
| Q7TN29 | Smap2 | 0.490 | 0.754 | 0.000 | 0.100 | -0.152 | -0.023 | -1.170 | 0.172 |
| P33215 | Nedd1 | 0.164 | 0.344 | -0.152 | 0.000 | 0.093 | 0.011 | -0.730 | -0.524 |
| O08528 | Hk2 | 0.670 | 0.948 | 0.751 | 0.000 | -0.383 | 0.849 | -0.130 | 0.255 |
| Q8BZ21 | Kat6a | 0.030 | 0.114 | -0.637 | 0.000 | 0.235 | -0.967 | -1.210 | -1.830 |
| O35218 | Cpsf2 | 0.010 | 0.060 | -0.237 | 0.000 | 0.049 | -0.570 | -0.956 | -1.070 |
| O09000 | Ncoa3 | 0.012 | 0.067 | -0.143 | 0.000 | 0.327 | -0.566 | -1.070 | -0.840 |
| Q8K394 | Plcl2 | 0.265 | 0.483 | -1.200 | 0.866 | 0.000 | -2.160 | 0.368 | -3.650 |
| Q9QYF1 | Rdh11 | 0.005 | 0.043 | -0.299 | 0.000 | 0.105 | 1.200 | 1.590 | 0.889 |
| Q68FG3 | Spty2d1 | 0.592 | 0.864 | 0.222 | 0.000 | -0.215 | 0.424 | 0.089 | -1.650 |
| O54918 | Bcl2l11 | 0.580 | 0.851 | 0.000 | 0.199 | -0.024 | 0.457 | -0.129 | 0.176 |
| P70279 | Surf6 | 0.040 | 0.135 | -0.194 | 0.000 | 0.050 | -0.442 | -1.100 | -0.554 |
| P10853 | Hist1h2bf | 0.891 | 1.000 | 0.000 | 1.230 | -0.393 | -0.370 | -0.110 | 1.030 |
| P10854 | Hist1h2bm | 0.891 | 1.000 | 0.000 | 1.230 | -0.393 | -0.370 | -0.110 | 1.030 |
| Q64475 | Hist1h2bb | 0.891 | 1.000 | 0.000 | 1.230 | -0.393 | -0.370 | -0.110 | 1.030 |
| Q64478 | Hist1h2bh | 0.891 | 1.000 | 0.000 | 1.230 | -0.393 | -0.370 | -0.110 | 1.030 |
| Q64525 | Hist2h2bb | 0.891 | 1.000 | 0.000 | 1.230 | -0.393 | -0.370 | -0.110 | 1.030 |
| Q6ZWY9 | Hist1h2bc | 0.891 | 1.000 | 0.000 | 1.230 | -0.393 | -0.370 | -0.110 | 1.030 |
| Q8CGP2 | Hist1h2bp | 0.891 | 1.000 | 0.000 | 1.230 | -0.393 | -0.370 | -0.110 | 1.030 |
| Q3U4G3 | Xxylt1 | 0.129 | 0.292 | -3.840 | 0.000 | 0.665 | 2.220 | 0.974 | 1.950 |
| Q9D8Y1 | Tmem126a | 0.000 | 0.011 | 0.000 | 0.009 | -0.058 | 1.380 | 1.100 | 1.350 |
| Q8VE10 | Naa40 | 0.971 | 1.000 | 0.406 | -0.674 | 0.000 | 0.640 | -0.660 | -0.192 |
| P29352 | Ptpn22 | 0.929 | 1.000 | -0.876 | 0.000 | 0.428 | 0.287 | -0.538 | -0.326 |
| Q9D067 | Mdm1 | 0.213 | 0.409 | -0.765 | 0.076 | 0.000 | -1.780 | -1.030 | -0.213 |
| Q3TL44 | Nlrx1 | 0.072 | 0.196 | 0.000 | 0.222 | -0.158 | 0.479 | 0.242 | 0.303 |
| P61202 | Cops2 | 0.045 | 0.146 | 0.034 | -0.233 | 0.000 | -0.510 | -1.110 | -0.490 |
| Q9CZN8 | Qrsl1 | 0.705 | 0.982 | 0.000 | 0.121 | -0.527 | 0.510 | -0.341 | -0.174 |
| Q9WUD1 | Stub1 | 0.453 | 0.711 | 0.263 | 0.000 | -0.427 | -2.900 | 0.114 | 0.084 |
| Q9CQC6 | Bzw1 | 0.586 | 0.858 | 0.316 | 0.000 | -0.007 | 0.514 | -0.502 | -0.281 |
| P27577 | Ets1 | 0.124 | 0.284 | 0.000 | -0.224 | 0.369 | -0.182 | -1.230 | -0.507 |
| O35239 | Ptpn9 | 0.099 | 0.243 | 1.850 | 0.000 | -0.292 | 1.650 | 2.830 | 1.930 |
| Q8BTY2 | Slc4a7 | 0.880 | 1.000 | -1.360 | 0.085 | 0.000 | 1.850 | -2.210 | -0.310 |
| P62311 | Lsm3 | 0.001 | 0.015 | -0.195 | 0.357 | 0.000 | -1.830 | -1.570 | -1.770 |
| Q8CI95 | Osbpl11 | 0.208 | 0.404 | 0.301 | -0.146 | 0.000 | 0.187 | -3.070 | -1.250 |
| Q9JMG1 | Edf1 | 0.286 | 0.511 | -0.171 | 0.000 | 0.306 | -0.570 | -0.968 | 0.252 |
| Q91VN4 | Chchd6 | 0.012 | 0.068 | 0.000 | 0.253 | -1.110 | 1.520 | 1.500 | 1.590 |
| Q9ESJ0 | Xpo4 | 0.053 | 0.162 | -0.029 | 0.000 | 0.040 | -0.345 | -1.510 | -0.877 |
| Q8BSU7 | Mob3a | 0.407 | 0.658 | 0.000 | 0.097 | -2.220 | -0.040 | -0.204 | 0.256 |
| Q6S5J6 | Krit1 | 0.383 | 0.632 | -0.145 | 0.136 | 0.000 | 0.151 | -5.620 | -0.086 |
| Q61211 | Eif2d | 0.125 | 0.285 | 0.228 | -0.376 | 0.000 | -0.314 | -1.080 | -0.460 |
| Q9WTS2 | Fut8 | 0.130 | 0.293 | 0.405 | 0.000 | -0.479 | 0.706 | 0.666 | 0.242 |
| Q78IS1 | Tmed3 | 0.014 | 0.074 | 0.000 | 0.223 | -1.510 | 1.870 | 1.690 | 2.000 |
| Q8R079 | Bfar | 0.008 | 0.054 | 0.000 | 0.725 | -0.217 | 1.700 | 1.460 | 1.620 |
| Q8VEB4 | Pla2g15 | 0.315 | 0.549 | 0.547 | 0.000 | -0.266 | 0.162 | -0.645 | -0.399 |
| Q9WU56 | Pus1 | 0.903 | 1.000 | 2.900 | -0.702 | 0.000 | 2.620 | 0.251 | -1.290 |
| Q6PHN9 | Rab35 | 0.343 | 0.583 | 0.000 | 0.085 | -0.300 | 0.045 | -0.496 | -0.430 |
| Q8K4I3 | Arhgef6 | 0.010 | 0.058 | -0.259 | 0.000 | 0.183 | -0.812 | -1.510 | -1.100 |
| Q91ZN5 | Slc35b2 | 0.050 | 0.156 | 0.290 | -0.058 | 0.000 | 0.603 | 0.284 | 0.611 |

| | | | | | | | | | |
|---|---|---|---|---|---|---|---|---|---|
| Q9CQT2 | Rbm7 | 0.006 | 0.045 | 0.000 | 0.373 | -0.119 | -1.020 | -0.864 | -0.687 |
| B2RWS6 | Ep300 | 0.670 | 0.948 | 0.000 | 1.170 | -0.194 | 0.000 | 0.248 | 0.134 |
| O35904 | Pik3cd | 0.505 | 0.769 | 0.000 | -0.553 | 0.026 | -0.185 | -0.997 | -0.099 |
| P53569 | Cebpz | 0.047 | 0.150 | -0.954 | 0.335 | 0.000 | -1.100 | -1.420 | -1.770 |
| Q71RI9 | Kyat3 | 0.016 | 0.079 | 0.000 | 0.041 | -0.711 | 0.821 | 0.680 | 0.805 |
| O88587 | Comt | 0.014 | 0.072 | -0.457 | 0.392 | 0.000 | 1.030 | 0.947 | 1.110 |
| Q3U3I9 | Znf865 | 0.618 | 0.892 | -0.629 | 0.000 | 0.065 | -0.096 | -1.410 | 0.093 |
| O55187 | Cbx4 | 0.330 | 0.567 | -0.691 | 0.501 | 0.000 | -0.719 | -0.528 | -0.198 |
| Q03958 | Pfdn6 | 0.085 | 0.220 | 0.000 | 1.140 | -1.340 | -2.000 | -2.560 | -1.220 |
| P50295 | Nat2 | 0.967 | 1.000 | -0.476 | 1.580 | 0.000 | -0.090 | 1.210 | 0.073 |
| Q91YJ2 | Snx4 | 0.196 | 0.389 | -2.760 | 1.530 | 0.000 | 2.230 | 0.355 | 3.230 |
| P10820 | Prf1 | 0.674 | 0.951 | -2.310 | 0.297 | 0.000 | -1.810 | -1.130 | -0.338 |
| Q8CGZ0 | Cherp | 0.085 | 0.220 | -0.172 | 0.165 | 0.000 | -0.178 | -0.463 | -0.254 |
| Q3UFM5 | Nom1 | 0.004 | 0.036 | 0.326 | 0.000 | -0.274 | -1.150 | -1.300 | -0.992 |
| Q9D338 | Mrpl19 | 0.721 | 1.000 | 0.000 | -0.168 | 0.031 | 1.100 | -0.582 | -0.080 |
| Q8R3Q0 | Saraf | 0.011 | 0.063 | 0.000 | -0.070 | 0.063 | 0.953 | 0.424 | 0.822 |
| Q8BIP0 | Dars2 | 0.009 | 0.056 | 0.571 | -0.292 | 0.000 | 1.550 | 1.440 | 2.180 |
| Q9Z1G3 | Atp6v1c1 | 0.811 | 1.000 | 0.655 | -0.150 | 0.000 | 0.660 | -0.397 | -0.062 |
| P63085 | Mapk1 | 0.087 | 0.224 | 0.263 | 0.000 | -0.001 | -0.072 | -0.496 | -0.204 |
| Q6DID7 | Wls | 0.003 | 0.031 | 0.000 | 0.433 | -0.720 | 2.130 | 2.080 | 2.410 |
| A2A4P0 | Dhx8 | 0.062 | 0.178 | 0.141 | -0.124 | 0.000 | -0.336 | -1.520 | -0.833 |
| Q3TB48 | Tmem104 | 0.184 | 0.373 | 0.520 | -0.340 | 0.000 | 0.970 | 0.323 | 0.427 |
| A3KMP2 | Ttc38 | 0.005 | 0.039 | -0.216 | 0.000 | 0.035 | -1.560 | -2.530 | -1.630 |
| Q60974 | Ncor1 | 0.231 | 0.436 | -0.323 | 0.025 | 0.000 | -0.567 | -1.790 | -0.112 |
| Q60855 | Ripk1 | 0.652 | 0.929 | 0.790 | 0.000 | -0.944 | 0.818 | -1.660 | -0.584 |
| Q810V0 | Mphosph10 | 0.560 | 0.829 | 0.000 | -0.034 | 0.392 | 0.108 | -0.121 | 0.077 |
| P42128 | Foxk1 | 0.148 | 0.318 | -0.996 | 0.000 | 0.660 | -1.930 | -3.160 | -0.337 |
| P39688 | Fyn | 0.582 | 0.853 | -0.633 | 0.103 | 0.000 | -0.287 | -0.529 | -0.169 |
| O88842 | Fgd3 | 0.002 | 0.024 | 0.129 | 0.000 | -0.012 | -1.240 | -1.930 | -1.440 |
| Q99LB2 | Dhrs4 | 0.046 | 0.147 | 0.000 | 0.173 | -0.424 | 1.020 | 0.369 | 0.601 |
| Q8C5H8 | Nadk2 | 0.914 | 1.000 | 0.228 | 0.000 | -0.004 | 0.494 | -0.258 | -0.095 |
| Q99JP7 | Ggt7 | 0.000 | 0.008 | -0.382 | 0.079 | 0.000 | 3.770 | 3.880 | 4.330 |
| O54901 | Cd200 | 0.061 | 0.176 | 0.000 | -0.070 | 0.296 | -0.290 | -0.577 | -0.177 |
| Q8R0H9 | Gga1 | 0.483 | 0.746 | 0.000 | -0.245 | 1.100 | 0.656 | -5.080 | 0.722 |
| Q9CYD3 | Crtap | 0.001 | 0.019 | 0.000 | 1.040 | -0.922 | 5.360 | 4.890 | 5.340 |
| Q9DBU0 | Tm9sf1 | 0.008 | 0.051 | 0.000 | 0.173 | -0.179 | 0.623 | 0.620 | 0.936 |
| A2A5R2 | Arfgef2 | 0.338 | 0.577 | 0.091 | 0.000 | -0.236 | 0.211 | -0.703 | -0.690 |
| Q9DBB4 | Naa16 | 0.660 | 0.936 | 0.000 | 1.260 | -0.821 | 0.296 | 1.320 | -0.120 |
| Q8CJG1 | Ago1 | 0.840 | 1.000 | -0.612 | 1.730 | 0.000 | 0.802 | 1.310 | -0.427 |
| Q3TAA7 | Stk11ip | 0.010 | 0.061 | 0.014 | -0.019 | 0.000 | -0.701 | -1.570 | -1.160 |
| Q61749 | Eif2b4 | 0.245 | 0.456 | 0.000 | 0.607 | -0.066 | -0.100 | -0.030 | -1.620 |
| Q9JJ80 | Rpf2 | 0.464 | 0.722 | 0.000 | 2.000 | -0.260 | 0.242 | -1.260 | 0.554 |
| Q6VN19 | Ranbp10 | 0.266 | 0.485 | 0.000 | -1.080 | 0.050 | 0.796 | 0.177 | -0.191 |
| Q64700 | Rbl2 | 0.364 | 0.611 | -0.325 | 0.595 | 0.000 | -0.538 | -0.970 | 0.337 |
| Q64697 | Ptprcap | 0.230 | 0.434 | -0.165 | 0.189 | 0.000 | 0.135 | 0.105 | 0.266 |
| O88738 | Birc6 | 0.391 | 0.640 | 0.401 | 0.000 | -0.399 | 0.604 | -1.420 | -1.120 |
| Q9QXK7 | Cpsf3 | 0.058 | 0.171 | -0.218 | 0.098 | 0.000 | -0.222 | -0.525 | -0.385 |
| O35975 | Art2b | 0.057 | 0.170 | 0.000 | -0.256 | 0.901 | -1.100 | -1.580 | -0.463 |
| Q9R0Q7 | Ptges3 | 0.026 | 0.105 | -0.221 | 0.085 | 0.000 | -2.030 | -0.845 | -1.090 |
| O70496 | Clcn7 | 0.194 | 0.386 | 0.493 | -0.414 | 0.000 | 0.741 | 0.292 | 0.423 |
| Q9CQE3 | Mrps17 | 0.067 | 0.187 | 0.000 | 1.020 | -0.199 | 0.968 | 1.420 | 1.540 |
| Q8BPS4 | Gpr180 | 0.028 | 0.107 | 0.000 | -3.310 | 0.570 | 3.540 | 2.880 | 3.280 |
| Q80TP3 | Ubr5 | 0.617 | 0.891 | 0.000 | -1.080 | 0.287 | 0.572 | -1.580 | 2.050 |
| Q9JKL4 | Ndufaf3 | 0.124 | 0.284 | 0.000 | -1.040 | 0.623 | 0.737 | 0.694 | 1.040 |
| Q8BTY1 | Kyat1 | 0.005 | 0.039 | -0.231 | 0.533 | 0.000 | -1.400 | -2.010 | -2.310 |
| Q8VD04 | Gripap1 | 0.146 | 0.315 | 0.000 | 0.152 | -0.449 | -0.246 | -1.500 | -0.746 |
| Q8BHC4 | Dcakd | 0.017 | 0.081 | 0.466 | 0.000 | -1.560 | 2.210 | 1.810 | 2.370 |
| Q61553 | Fscn1 | 0.003 | 0.030 | 0.000 | -0.057 | 0.058 | 2.340 | 1.390 | 1.810 |
| Q8VDI1 | Epsti1 | 0.617 | 0.891 | 0.000 | -3.110 | 0.484 | -0.654 | -2.470 | -1.520 |

| Race | Sex | Age |
|---|---|---|
| Black | Male | 60 |
| Caucasian | Male | 61 |
| Black | Male | 66 |
| Black | Female | 59 |
| Caucasian | Male | 61 |
| Black | Male | 64 |
| Black | Male | 63 |
| Black | Male | 51 |
| Black | Female | 54 |
| Black | Female | 66 |

**Supplementary Table 12. Race, Sex and Age of Donors**

| | | | | | | | | | |
|---|---|---|---|---|---|---|---|---|---|
| Q99MK8 | Adrbk1 | 0.014 | 0.073 | 0.104 | 0.000 | -0.048 | -0.465 | -1.150 | -0.916 |
| Q9D1I6 | Mrpl14 | 0.431 | 0.687 | 0.279 | 0.000 | -0.206 | 0.925 | -0.016 | 0.044 |
| Q8BM72 | Hspa13 | 0.001 | 0.015 | 0.000 | 0.147 | -0.238 | 1.210 | 1.070 | 1.230 |
| Q8VE19 | Mios | 0.833 | 1.000 | 0.167 | -1.300 | 0.000 | -1.170 | -0.653 | 0.273 |
| Q8R4Z4 | Etv3 | 0.143 | 0.309 | -1.260 | 0.000 | 2.930 | 2.380 | 3.170 | 3.030 |
| P97820 | Map4k4 | 0.129 | 0.291 | 0.077 | -0.561 | 0.000 | 1.420 | 0.039 | 0.619 |
| P41216 | Acsl1 | 0.926 | 1.000 | 0.000 | 0.371 | -0.869 | 0.044 | 0.090 | -0.768 |
| Q5SF07 | Igf2bp2 | 0.047 | 0.150 | -0.112 | 3.740 | 0.000 | 5.410 | 4.380 | 4.900 |
| Q8BUV3 | Gphn | 0.039 | 0.133 | -0.540 | 0.113 | 0.000 | -0.819 | -1.730 | -1.000 |
| Q9DBS5 | Klc4 | 0.089 | 0.227 | 0.147 | -0.698 | 0.000 | -0.502 | -1.210 | -1.180 |
| Q9R020 | Zranb2 | 0.054 | 0.164 | 0.490 | -0.013 | 0.000 | -0.266 | -1.240 | -0.669 |
| P24638 | Acp2 | 0.034 | 0.121 | -0.034 | 0.054 | 0.000 | 0.460 | 0.246 | 0.763 |
| Q9D0D5 | Gtf2e1 | 0.030 | 0.112 | 0.202 | -0.161 | 0.000 | -0.852 | -2.490 | -1.470 |
| P52503 | Ndufs6 | 0.095 | 0.237 | -0.325 | 0.000 | 0.739 | 0.967 | 0.626 | 1.040 |
| Q99LS3 | Psph | 0.014 | 0.072 | -0.560 | 0.141 | 0.000 | -1.410 | -2.990 | -2.800 |
| Q6A028 | Swap70 | 0.926 | 1.000 | 0.163 | -0.424 | 0.000 | 0.391 | -0.367 | -0.378 |
| Q3UMF0 | Cobll1 | 0.420 | 0.674 | -1.450 | 0.298 | 0.000 | -1.040 | -1.100 | -0.545 |
| Q8R5F3 | Oard1 | 0.041 | 0.138 | 0.000 | -0.160 | 0.309 | -0.415 | -1.080 | -1.560 |
| Q6NTA4 | Rragb | 0.839 | 1.000 | 0.315 | 0.000 | -0.024 | 0.559 | -0.365 | -0.095 |
| Q80X95 | Rraga | 0.839 | 1.000 | 0.315 | 0.000 | -0.024 | 0.559 | -0.365 | -0.095 |
| Q8CFT2 | Setd1b | 0.197 | 0.391 | 0.000 | 0.729 | -0.041 | -0.218 | -0.221 | -2.330 |
| Q9CQ56 | Use1 | 0.012 | 0.069 | -0.371 | 0.000 | 0.230 | 1.080 | 0.697 | 0.796 |
| Q3U186 | Rars2 | 0.906 | 1.000 | 0.398 | -0.283 | 0.000 | 1.060 | -0.833 | -0.334 |
| P62488 | Polr2g | 0.001 | 0.022 | -0.119 | 0.000 | 0.092 | -0.560 | -0.762 | -0.686 |
| Q9WVM1 | Racgap1 | 0.320 | 0.555 | 1.770 | -0.913 | 0.000 | 2.990 | 0.689 | 0.859 |
| Q61112 | Sdf4 | 0.541 | 0.808 | -0.105 | 0.540 | 0.000 | 0.492 | -0.149 | 0.767 |
| Q924M7 | Mpi | 0.117 | 0.273 | -0.284 | 0.000 | 0.250 | -0.620 | -2.930 | -0.916 |
| Q8CAY6 | Acat2 | 0.170 | 0.352 | 0.652 | -0.172 | 0.000 | -0.123 | -0.377 | -0.341 |
| Q8R2T8 | Gtf3c5 | 0.023 | 0.096 | -0.196 | 0.161 | 0.000 | -0.379 | -0.708 | -0.482 |
| Q8K4P0 | Wdr33 | 0.948 | 1.000 | -1.110 | 0.000 | 0.822 | -1.840 | 1.850 | -0.538 |
| Q9Z0U1 | Tjp2 | 0.008 | 0.051 | -0.391 | 0.328 | 0.000 | -1.280 | -2.290 | -2.170 |
| Q64516 | Gk | 0.095 | 0.238 | 0.000 | -0.012 | 0.006 | 0.349 | 0.017 | 0.348 |
| Q8K2F0 | Brd3 | 0.367 | 0.614 | 0.328 | -1.040 | 0.000 | -0.693 | -0.824 | -0.484 |
| Q9JHU9 | Isyna1 | 0.002 | 0.029 | 0.000 | 0.098 | -0.167 | -1.140 | -1.850 | -1.610 |
| Q9D1H7 | Get4 | 0.857 | 1.000 | 0.078 | -0.294 | 0.000 | 0.092 | -0.419 | 0.000 |
| Q9CYN9 | Atp6ap2 | 0.094 | 0.236 | 0.000 | 0.055 | -0.400 | 1.080 | 0.129 | 0.492 |
| Q9WUV0 | Orc5 | 0.343 | 0.582 | 0.199 | 0.000 | -0.561 | -0.287 | -0.714 | -0.241 |
| O88712 | Ctbp1 | 0.008 | 0.052 | 0.000 | -0.306 | 0.027 | -1.060 | -1.940 | -1.410 |
| Q8VEK6 | Ing3 | 0.841 | 1.000 | -2.000 | 0.000 | 0.232 | -0.688 | -1.700 | 0.065 |
| Q6Q783 | Kmt5c | 0.331 | 0.568 | 0.000 | -0.746 | 0.263 | -0.306 | -0.727 | -0.531 |
| Q91W96 | Anapc4 | 0.893 | 1.000 | 0.000 | -0.318 | 0.245 | 0.275 | -0.305 | 0.058 |
| P56671 | Maz | 0.246 | 0.457 | -1.300 | 0.426 | 0.000 | 0.187 | 0.273 | 1.070 |
| Q6P1H6 | Ankle2 | 0.334 | 0.571 | 0.000 | -0.919 | 0.452 | 0.030 | 0.201 | 0.880 |
| Q6A065 | Cep170 | 0.998 | 1.000 | 0.069 | -0.070 | 0.000 | 0.013 | -0.627 | 0.616 |
| Q3U5Q7 | Cmpk2 | 0.171 | 0.354 | 0.545 | 0.000 | -0.063 | 0.143 | -0.649 | -0.640 |
| Q9R0A0 | Pex14 | 0.500 | 0.765 | -0.427 | 0.003 | 0.000 | 0.334 | -1.500 | -0.479 |
| A6X919 | Dpy19l1 | 0.008 | 0.055 | -0.026 | 0.177 | 0.000 | 1.360 | 0.686 | 1.070 |
| Q9R0Q9 | Mpdu1 | 0.207 | 0.404 | 0.000 | -0.009 | 0.020 | 0.421 | -0.073 | 0.373 |
| Q8BK08 | Tmem11 | 0.001 | 0.019 | -0.043 | 0.285 | 0.000 | 1.080 | 0.954 | 1.080 |
| Q9WUH1 | Tmem115 | 0.086 | 0.223 | -0.017 | 0.000 | 0.140 | 0.458 | 0.247 | 1.020 |
| Q8K400 | Stxbp5 | 0.503 | 0.767 | -2.540 | 0.000 | 1.020 | -0.396 | -2.860 | -1.110 |
| Q61122 | Nab1 | 0.355 | 0.598 | -0.983 | 0.000 | 0.250 | -0.976 | -1.240 | -0.108 |
| Q99JT2 | Stk26 | 0.019 | 0.087 | -0.010 | 0.035 | 0.000 | -0.926 | -2.380 | -1.490 |
| Q8R409 | Hexim1 | 0.026 | 0.103 | -0.301 | 0.000 | 0.002 | -0.507 | -0.763 | -1.100 |
| Q80ZV0 | Rnaseh2b | 0.816 | 1.000 | 2.290 | 0.000 | -0.370 | 2.040 | -0.713 | -0.300 |
| P55821 | Stmn2 | 0.078 | 0.208 | 0.511 | -0.166 | 0.000 | -0.361 | -2.580 | -1.480 |
| P54227 | Stmn1 | 0.078 | 0.208 | 0.511 | -0.166 | 0.000 | -0.361 | -2.580 | -1.480 |
| Q8BU30 | Iars | 0.239 | 0.448 | 0.356 | 0.000 | -0.013 | 0.658 | 0.088 | 0.450 |
| Q61510 | Trim25 | 0.944 | 1.000 | -0.760 | 2.540 | 0.000 | 0.403 | -0.707 | 1.800 |

| | | | | | | | | | |
|---|---|---|---|---|---|---|---|---|---|
| Q60649 | Clpb | 0.990 | 1.000 | 0.421 | 0.000 | -0.108 | 0.340 | -0.304 | 0.289 |
| P23506 | Pcmt1 | 0.001 | 0.022 | -0.179 | 0.000 | 0.198 | -0.923 | -0.926 | -0.835 |
| Q3TC33 | Ccdc127 | 0.031 | 0.114 | -0.180 | 0.197 | 0.000 | 0.651 | 0.308 | 0.506 |
| Q9CQI6 | Cotl1 | 0.000 | 0.011 | 0.009 | -0.028 | 0.000 | -1.450 | -1.890 | -1.690 |
| Q8BFQ3 | Gpr68 | 0.340 | 0.579 | -0.551 | 0.216 | 0.000 | -1.070 | -1.480 | 0.309 |
| P61514 | Rpl37a | 0.417 | 0.671 | 0.000 | -0.469 | 0.078 | 0.438 | -1.460 | -0.984 |
| P70362 | Ufd1l | 0.819 | 1.000 | 0.000 | 0.014 | -1.410 | -0.501 | -0.022 | -1.310 |
| Q3UVL4 | Vps51 | 0.208 | 0.404 | 0.208 | -1.640 | 0.000 | -0.988 | -2.320 | -1.320 |
| Q6PAQ4 | Rexo4 | 0.491 | 0.755 | 0.000 | 1.990 | -4.510 | 0.958 | 0.732 | 0.192 |
| Q8K3X4 | Irf2bpl | 0.277 | 0.498 | 0.000 | -0.251 | 0.040 | 0.100 | -0.645 | -0.692 |
| P81183 | Ikzf2 | 0.043 | 0.141 | -0.323 | 0.000 | 0.107 | -0.403 | -0.664 | -0.474 |
| Q921E2 | Rab31 | 0.164 | 0.343 | 0.261 | 0.000 | -0.128 | 1.650 | 0.002 | 0.999 |
| P70122 | Sbds | 0.148 | 0.318 | 0.000 | -0.375 | 0.036 | -0.198 | -0.845 | -0.521 |
| Q8BYK6 | Ythdf3 | 0.400 | 0.650 | -0.967 | 1.010 | 0.000 | -0.910 | -0.617 | -0.155 |
| Q8CFE4 | Scyl2 | 0.940 | 1.000 | 0.208 | 0.000 | -0.054 | 0.460 | -0.112 | -0.250 |
| Q8BTT6 | Diexf | 0.150 | 0.320 | 0.268 | -0.494 | 0.000 | -0.240 | -1.120 | -0.670 |
| A2AM29 | Mllt3 | 0.347 | 0.587 | -3.040 | 0.000 | 0.145 | 1.570 | -0.014 | -0.553 |
| P97461 | Rps5 | 0.392 | 0.641 | 0.431 | 0.000 | -0.092 | 0.476 | -0.557 | -0.791 |
| P59016 | Vps33b | 0.021 | 0.090 | 0.000 | 0.285 | -0.108 | -0.524 | -1.370 | -0.946 |
| Q9CR26 | Vta1 | 0.047 | 0.150 | 0.000 | 0.330 | -0.281 | -0.431 | -0.914 | -0.549 |
| O54984 | Asna1 | 0.442 | 0.698 | 0.075 | 0.000 | -0.446 | 0.147 | -0.143 | 0.099 |
| Q8C827 | Zfp62 | 0.293 | 0.520 | -0.534 | 0.344 | 0.000 | 0.573 | 0.335 | 0.005 |
| O88746 | Tom1 | 0.080 | 0.212 | 0.000 | 0.397 | -0.025 | -0.094 | -1.590 | -1.260 |
| Q78PG9 | Ccdc25 | 0.908 | 1.000 | -3.370 | 0.752 | 0.000 | -0.726 | -1.210 | -1.150 |
| P48453 | Ppp3cb | 0.034 | 0.122 | -0.837 | 0.064 | 0.000 | -2.010 | -3.260 | -1.390 |
| Q9CY73 | Mrpl44 | 0.009 | 0.057 | 0.000 | 0.358 | -0.175 | 0.827 | 0.770 | 0.815 |
| Q3UIA2 | Arhgap17 | 0.173 | 0.356 | 0.521 | -0.634 | 0.000 | 0.359 | 0.548 | 1.870 |
| Q80U63 | Mfn2 | 1.000 | 1.000 | 0.000 | 0.057 | -0.064 | -0.674 | 0.298 | 0.369 |
| Q9CX00 | Ist1 | 0.541 | 0.807 | 0.000 | -0.229 | 0.048 | 0.269 | -0.707 | -0.339 |
| Q6PDL0 | Dync1li2 | 0.091 | 0.231 | -1.050 | 0.729 | 0.000 | 1.510 | 0.945 | 0.902 |
| Q6PCM2 | Ints6 | 0.035 | 0.124 | -0.670 | 0.000 | 0.128 | -1.040 | -1.070 | -0.853 |
| Q8VE96 | Slc35f6 | 0.285 | 0.509 | 0.589 | -0.450 | 0.000 | 0.607 | -1.970 | -1.740 |
| Q8VDQ8 | Sirt2 | 0.284 | 0.508 | -0.577 | 0.764 | 0.000 | -0.309 | -2.220 | -0.144 |
| Q8VDM1 | Zgpat | 0.055 | 0.166 | 0.000 | -0.755 | 0.028 | -0.768 | -1.080 | -1.150 |
| Q8BX09 | Rbbp5 | 0.060 | 0.175 | -0.133 | 0.000 | 0.042 | -0.266 | -0.743 | -0.324 |
| Q9D2C7 | Tmbim6 | 0.209 | 0.406 | -0.632 | 0.134 | 0.000 | -0.118 | 0.420 | 0.780 |
| Q8K2C8 | Gpat4 | 0.002 | 0.023 | 0.456 | 0.000 | -0.135 | 2.420 | 1.830 | 1.920 |
| Q9DC71 | Mrps15 | 0.711 | 0.987 | -0.194 | 0.000 | 0.044 | 0.841 | -0.657 | 0.193 |
| Q3TCH7 | Cul4a | 0.019 | 0.085 | 0.019 | -0.448 | 0.000 | 1.410 | 0.531 | 1.040 |
| Q8R0K4 | Ccdc137 | 0.576 | 0.846 | -3.380 | 0.000 | 0.605 | -0.132 | -0.475 | 0.118 |
| E9Q784 | Zc3h13 | 0.413 | 0.665 | 0.000 | -0.003 | 1.120 | 0.228 | 1.240 | 0.968 |
| Q920A5 | Scpep1 | 0.166 | 0.347 | 0.029 | -0.044 | 0.000 | 0.724 | -0.059 | 0.508 |
| Q9CR59 | Gadd45gip1 | 0.130 | 0.293 | 0.000 | 0.151 | -0.194 | 0.895 | 0.031 | 0.578 |
| Q64131 | Runx3 | 0.617 | 0.891 | 0.000 | 2.480 | -1.840 | 0.041 | 1.500 | 1.260 |
| P70195 | Psmb7 | 0.075 | 0.202 | 0.000 | 0.109 | -0.848 | -0.791 | -1.820 | -1.190 |
| Q9WTL7 | Lypla2 | 0.069 | 0.191 | -0.604 | 1.470 | 0.000 | -1.450 | -2.490 | -0.908 |
| Q8BVU5 | Nudt9 | 0.155 | 0.330 | 0.000 | -0.351 | 0.224 | 0.896 | 2.140 | 0.089 |
| O70378 | Emc8 | 0.013 | 0.070 | -0.015 | 0.398 | 0.000 | 1.360 | 0.752 | 1.150 |
| Q8BVA5 | Ldah | 0.407 | 0.658 | 0.000 | 0.014 | -0.054 | 0.285 | -1.310 | -0.310 |
| P39428 | Traf1 | 0.026 | 0.103 | 0.000 | 0.232 | -0.135 | -0.817 | -1.800 | -0.841 |
| Q9ES74 | Nek7 | 0.578 | 0.849 | 0.000 | -0.083 | 0.235 | 0.231 | -0.367 | -0.069 |
| P70428 | Ext2 | 0.130 | 0.293 | 0.004 | -4.570 | 0.000 | 1.530 | 1.060 | 1.580 |
| Q8BFQ6 | Dirc2 | 0.209 | 0.405 | 0.419 | 0.000 | -0.130 | 0.891 | 0.085 | 0.610 |
| Q2TPA8 | Hsdl2 | 0.861 | 1.000 | 0.190 | -0.464 | 0.000 | 0.705 | -0.463 | -0.284 |
| P48725 | Pcnt | 0.954 | 1.000 | 0.634 | -3.730 | 0.000 | 0.667 | -2.740 | -1.330 |
| Q8R344 | Ccdc12 | 0.022 | 0.094 | 0.045 | -0.401 | 0.000 | -0.707 | -1.280 | -1.590 |
| Q9Z172 | Sumo3 | 0.504 | 0.768 | 0.000 | 0.965 | -0.168 | -0.170 | -3.010 | 1.160 |
| Q80ZD3 | Slc26a11 | 0.066 | 0.185 | -0.440 | 0.256 | 0.000 | 0.382 | 0.561 | 1.090 |
| Q9WU28 | Pfdn5 | 0.026 | 0.104 | 0.337 | 0.000 | -0.011 | -0.630 | -0.847 | -0.279 |

| | | | | | | | | | |
|---|---|---|---|---|---|---|---|---|---|
| Q9CQE7 | Ergic3 | 0.027 | 0.105 | 0.103 | -0.420 | 0.000 | 0.997 | 0.398 | 0.714 |
| P70269 | Ctse | 0.222 | 0.423 | 0.292 | 0.000 | -0.683 | -0.227 | -1.010 | -0.757 |
| Q4QRL3 | Ccdc88b | 0.044 | 0.142 | 0.000 | 0.221 | -0.969 | -2.650 | -4.350 | -1.550 |
| P18052 | Ptpra | 0.075 | 0.202 | -0.163 | 0.203 | 0.000 | 0.660 | 0.156 | 0.735 |
| Q8BIH0 | Sap130 | 0.032 | 0.117 | 0.000 | -0.146 | 1.120 | 1.420 | 1.820 | 1.790 |
| Q99MU3 | Adar | 0.004 | 0.038 | -0.077 | 0.000 | 0.110 | -1.330 | -1.330 | -0.753 |
| Q811M1 | Arhgap15 | 0.024 | 0.097 | 0.063 | -0.056 | 0.000 | -0.320 | -0.940 | -0.689 |
| Q14AX6 | Cdk12 | 0.354 | 0.596 | 0.044 | -0.085 | 0.000 | 0.080 | -0.202 | -0.290 |
| Q3TC72 | Fahd2 | 0.460 | 0.719 | 0.000 | 0.309 | -0.221 | 0.143 | 0.026 | 0.366 |
| Q921I2 | Klhdc4 | 0.441 | 0.697 | 0.550 | -0.537 | 0.000 | 1.260 | -0.103 | 0.190 |
| Q9D0I8 | Mrto4 | 0.098 | 0.242 | 0.240 | 0.000 | -0.243 | -0.157 | -0.805 | -0.552 |
| Q8CIC2 | Nupl2 | 0.671 | 0.949 | -1.300 | 0.382 | 0.000 | -4.180 | 2.690 | 4.170 |
| Q8C092 | Taf5 | 0.864 | 1.000 | -0.202 | 2.780 | 0.000 | 1.600 | 0.516 | -0.136 |
| P97492 | Rgs14 | 0.001 | 0.019 | -0.396 | 0.195 | 0.000 | -2.050 | -1.910 | -1.620 |
| Q9R1C6 | Dgke | 0.653 | 0.929 | 0.000 | 2.230 | -1.630 | 0.861 | 0.474 | 0.905 |
| Q61205 | Pafah1b3 | 0.001 | 0.019 | -0.071 | 0.230 | 0.000 | -0.968 | -1.320 | -1.130 |
| Q9D8S9 | Bola1 | 0.460 | 0.719 | 0.000 | -2.370 | 0.187 | 1.050 | -0.214 | -0.646 |
| Q9R1S3 | Pign | 0.869 | 1.000 | -2.550 | 0.000 | 5.880 | -3.330 | 7.380 | -3.010 |
| Q9CR68 | Uqcrfs1 | 0.007 | 0.048 | 0.000 | 0.074 | -0.106 | 0.656 | 0.355 | 0.524 |
| Q8C7Q4 | Rbm4 | 0.306 | 0.538 | -0.094 | 0.211 | 0.000 | -0.147 | -0.129 | 0.023 |
| Q80TN4 | Dnajc16 | 0.527 | 0.793 | 0.000 | 3.370 | -3.120 | 0.741 | 1.600 | 1.870 |
| Q8CJ40 | Crocc | 0.148 | 0.318 | 0.071 | 0.000 | -0.008 | 0.321 | 0.014 | 0.498 |
| Q9JJY4 | Ddx20 | 0.299 | 0.528 | -0.178 | 0.000 | 0.117 | 0.198 | -0.676 | -0.663 |
| Q9CQZ5 | Ndufa6 | 0.233 | 0.438 | 0.000 | 0.678 | -0.132 | 0.410 | 0.479 | 0.879 |
| Q6PAR5 | Gapvd1 | 0.538 | 0.806 | 0.400 | 0.000 | -0.331 | 0.219 | -0.246 | -0.508 |
| Q8CE46 | Pus7l | 0.019 | 0.087 | 0.615 | -0.188 | 0.000 | -0.920 | -2.350 | -2.310 |
| Q922H4 | Gmppa | 0.029 | 0.112 | 0.060 | -0.305 | 0.000 | -0.535 | -1.460 | -1.280 |
| Q64455 | Ptprj | 0.728 | 1.000 | 0.025 | 0.000 | -1.110 | -0.320 | -0.571 | -0.618 |
| P63323 | Rps12 | 0.343 | 0.582 | -1.410 | 0.000 | 0.032 | -1.460 | -0.606 | -1.040 |
| Q9Z148 | Ehmt2 | 0.654 | 0.930 | -0.141 | 0.000 | 0.802 | -0.151 | -0.571 | 0.696 |
| Q9JI78 | Ngly1 | 0.004 | 0.037 | -0.505 | 0.000 | 0.109 | -1.300 | -1.710 | -1.850 |
| Q99N89 | Mrpl43 | 0.003 | 0.030 | 0.000 | 0.185 | -0.319 | 1.180 | 0.974 | 1.330 |
| O88879 | Apaf1 | 0.004 | 0.033 | 0.100 | -0.012 | 0.000 | -0.683 | -1.230 | -1.050 |
| P56382 | Atp5e | 0.000 | 0.013 | 0.000 | 0.115 | -0.081 | 0.914 | 0.754 | 0.858 |
| Q8BK58 | Hspbap1 | 0.857 | 1.000 | -0.469 | 0.577 | 0.000 | 1.280 | -0.737 | -0.052 |
| P30355 | Alox5ap | 0.424 | 0.678 | 0.786 | 0.000 | -0.301 | 1.160 | 0.291 | 0.218 |
| P51829 | Adcy7 | 0.555 | 0.824 | -0.535 | 0.000 | 0.058 | -0.200 | -0.585 | -0.144 |
| P61294 | Rab6b | 0.031 | 0.116 | -0.459 | 0.121 | 0.000 | -1.130 | -0.840 | -0.616 |
| Q9D4H8 | Cul2 | 0.495 | 0.760 | -1.430 | 1.200 | 0.000 | 0.568 | -1.510 | -1.640 |
| Q8CES0 | Naa30 | 0.115 | 0.270 | -0.122 | 0.000 | 0.086 | -0.325 | -1.500 | -0.472 |
| Q9D659 | Vsir | 0.049 | 0.154 | -0.084 | 0.110 | 0.000 | -0.152 | -0.651 | -0.493 |
| Q9Z129 | Recql | 0.224 | 0.426 | 0.000 | -0.274 | 0.511 | -0.249 | -1.960 | -0.223 |
| P62274 | Rps29 | 0.691 | 0.967 | 0.000 | 0.231 | -0.784 | -0.246 | -0.324 | -0.381 |
| Q91YJ3 | Thyn1 | 0.253 | 0.466 | 0.000 | -0.137 | 0.320 | -0.121 | -0.330 | -0.020 |
| Q61508 | Ecm1 | 0.050 | 0.155 | 0.000 | 0.399 | -0.581 | 0.993 | 0.600 | 0.775 |
| Q3UHD6 | Snx27 | 0.771 | 1.000 | -0.204 | 0.000 | 0.423 | -1.050 | -0.625 | 1.220 |
| Q9JKN1 | Slc30a7 | 0.061 | 0.176 | 0.000 | 0.548 | -0.468 | 1.350 | 0.531 | 1.200 |
| P47930 | Fosl2 | 0.051 | 0.158 | -0.388 | 0.000 | 0.238 | 0.409 | 0.428 | 0.671 |
| Q9WV84 | Nme4 | 0.027 | 0.105 | 1.300 | -1.640 | 0.000 | 3.340 | 2.630 | 2.720 |
| B1AVZ0 | Uprt | 0.350 | 0.591 | 0.000 | -0.111 | 0.166 | 0.131 | -0.627 | -0.192 |
| Q00422 | Gabpa | 0.266 | 0.485 | -0.698 | 0.165 | 0.000 | -0.673 | -0.617 | -0.342 |
| Q9CWN7 | Cnot11 | 0.150 | 0.320 | 0.000 | -0.054 | 0.281 | 0.098 | -2.610 | -1.490 |
| Q9D1P2 | Kat8 | 0.635 | 0.912 | 0.000 | 1.430 | -0.154 | 0.253 | -0.149 | 0.359 |
| Q9DAK9 | Phpt1 | 0.001 | 0.014 | -0.073 | 0.450 | 0.000 | -2.300 | -2.910 | -3.060 |
| Q8BYC6 | Taok3 | 0.583 | 0.855 | 0.000 | 0.347 | -0.177 | 0.237 | -0.245 | -0.214 |
| Q9CTH6 | Fcf1 | 0.015 | 0.076 | 0.162 | -0.174 | 0.000 | -0.675 | -0.546 | -0.377 |
| Q9CXK9 | Rbm33 | 0.775 | 1.000 | 0.000 | -0.130 | 1.580 | 0.673 | 0.060 | 1.320 |
| Q9D1B9 | Mrpl28 | 0.304 | 0.534 | 0.000 | 0.671 | -1.410 | 0.836 | 0.095 | 0.635 |
| P57787 | Slc16a3 | 0.600 | 0.873 | 0.578 | 0.000 | -3.850 | -0.543 | 0.643 | -0.876 |

| | | | | | | | | | |
|---|---|---|---|---|---|---|---|---|---|
| Q91W86 | Vps11 | 0.178 | 0.364 | 0.000 | 0.651 | -0.315 | -0.712 | -0.050 | -0.625 |
| O70469 | Dok2 | 0.058 | 0.172 | -0.026 | 0.000 | 0.092 | -0.602 | -1.430 | -2.690 |
| Q61179 | Irf9 | 0.154 | 0.327 | 0.021 | -0.204 | 0.000 | -0.140 | -1.020 | -0.437 |
| Q80U87 | Usp8 | 0.729 | 1.000 | 0.000 | -0.253 | 0.161 | 0.513 | -0.883 | -0.191 |
| Q9CWP6 | Mospd2 | 0.067 | 0.187 | -1.080 | 0.277 | 0.000 | 1.090 | 0.599 | 2.170 |
| Q6ZWM4 | Lsm8 | 0.003 | 0.032 | -0.155 | 0.000 | 0.168 | -0.915 | -1.470 | -1.420 |
| Q9CQZ0 | Ormdl2 | 0.062 | 0.179 | 0.000 | 0.413 | -0.174 | 1.130 | 0.665 | 0.468 |
| O54988 | Slk | 0.686 | 0.963 | -0.072 | 0.488 | 0.000 | -0.441 | 0.770 | 0.635 |
| Q8BGQ1 | Vipas39 | 0.027 | 0.107 | 0.014 | -0.411 | 0.000 | -1.310 | -1.870 | -0.768 |
| Q60692 | Psmb6 | 0.528 | 0.795 | 0.114 | 0.000 | -0.437 | -0.196 | -0.763 | 0.035 |
| Q3UXZ9 | Kdm5a | 0.036 | 0.126 | -1.040 | 0.237 | 0.000 | -1.650 | -3.230 | -1.830 |
| Q922R5 | Ppp4r3b | 0.120 | 0.276 | -0.578 | 0.000 | 0.820 | -0.544 | -3.400 | -1.380 |
| Q8R4E9 | Cdt1 | 0.525 | 0.791 | -0.052 | 0.000 | 1.510 | 0.633 | 0.311 | -0.962 |
| Q8BIG7 | Comtd1 | 0.937 | 1.000 | 0.000 | 0.166 | -0.276 | 0.673 | -0.609 | -0.276 |
| P28843 | Dpp4 | 0.067 | 0.186 | -0.165 | 0.000 | 0.192 | -0.310 | -0.587 | -0.216 |
| Q8K2D3 | Edc3 | 0.186 | 0.376 | 0.000 | 0.370 | -0.732 | -0.284 | -2.170 | -0.966 |
| Q5IRJ6 | Slc30a9 | 0.003 | 0.030 | 0.000 | 0.577 | -0.728 | 2.360 | 2.540 | 2.810 |
| Q9CXY9 | Pigk | 0.948 | 1.000 | 0.000 | 0.302 | -1.030 | 1.300 | -1.050 | -0.800 |
| Q8QZY6 | Tspan14 | 0.105 | 0.252 | 0.000 | 0.194 | -0.577 | -0.743 | -0.701 | -0.480 |
| Q9D2D7 | Znf687 | 0.027 | 0.107 | -0.869 | 0.000 | 0.043 | -1.140 | -1.420 | -1.520 |
| Q32NY4 | Cnnm3 | 0.062 | 0.179 | 0.143 | 0.000 | -0.041 | 0.185 | 0.199 | 0.396 |
| Q63810 | Ppp3r1 | 0.027 | 0.106 | 0.029 | 0.000 | -0.303 | -0.823 | -0.787 | -1.610 |
| Q9DB40 | Med27 | 0.112 | 0.265 | -0.150 | 0.455 | 0.000 | 0.392 | 0.886 | 0.476 |
| Q8R480 | Nup85 | 0.020 | 0.089 | 0.000 | -0.337 | 0.052 | -0.995 | -2.360 | -1.570 |
| Q8K339 | Kin | 0.040 | 0.134 | 0.000 | 0.000 | 0.258 | -0.190 | -0.860 | -0.618 |
| Q8BJW5 | Nol11 | 0.203 | 0.399 | 0.000 | 0.243 | -0.456 | -0.152 | -1.030 | -0.521 |
| Q9CVD2 | Atxn3 | 0.128 | 0.290 | 0.000 | 2.050 | -1.220 | -2.050 | -2.400 | -0.837 |
| P12399 | Ctla2a | 0.017 | 0.081 | 0.000 | 0.559 | -1.130 | -2.010 | -3.210 | -2.490 |
| Q924D0 | Rtn4ip1 | 0.540 | 0.807 | 0.000 | 0.155 | -0.927 | 0.162 | -2.980 | -0.090 |
| P15535 | B4galt1 | 0.213 | 0.410 | -0.367 | 0.178 | 0.000 | 0.336 | 0.153 | 0.102 |
| Q8CBC4 | Cnst | 0.009 | 0.057 | -0.306 | 0.000 | 0.270 | -0.789 | -1.300 | -1.290 |
| Q8VDY9 | Caap1 | 0.372 | 0.621 | -0.225 | 0.000 | 0.447 | -0.044 | -0.844 | 0.064 |
| Q8R2Q8 | Bst2 | 0.003 | 0.033 | 0.189 | -0.754 | 0.000 | 2.100 | 1.580 | 1.960 |
| Q5U3K5 | Rabl6 | 0.406 | 0.658 | 1.260 | -2.210 | 0.000 | 1.200 | -3.130 | -4.630 |
| Q8BZ98 | Dnm3 | 0.116 | 0.271 | 0.100 | 0.000 | -0.235 | 0.384 | 0.055 | 0.252 |
| Q9D023 | Mpc2 | 0.034 | 0.121 | 0.000 | 0.091 | -0.908 | 0.954 | 0.558 | 0.976 |
| O70493 | Snx12 | 0.150 | 0.321 | -0.260 | 0.065 | 0.000 | -0.143 | -0.807 | -0.403 |
| Q9DBY1 | Syvn1 | 0.020 | 0.087 | 0.000 | 0.682 | -0.142 | 1.310 | 1.110 | 1.100 |
| Q5H8C4 | Vps13a | 0.282 | 0.505 | -0.287 | 3.230 | 0.000 | -3.080 | -0.289 | 0.484 |
| Q99KX1 | Mlf2 | 0.315 | 0.549 | 0.000 | -0.307 | 0.193 | -0.085 | -0.336 | -0.256 |
| Q60596 | Xrcc1 | 0.290 | 0.517 | 0.023 | -0.284 | 0.000 | 0.104 | -0.792 | -0.651 |
| P59481 | Lman2l | 0.552 | 0.821 | 0.000 | 0.129 | -0.521 | 0.946 | -8.880 | 1.110 |
| P97789 | Xrn1 | 0.074 | 0.201 | 0.000 | -0.143 | 0.334 | -0.210 | -0.476 | -0.909 |
| Q99K01 | Pdxdc1 | 0.918 | 1.000 | 0.000 | -0.172 | 0.089 | 1.040 | -0.335 | -0.985 |
| Q8CIG3 | Kdm1b | 0.705 | 0.982 | 0.120 | -1.320 | 0.000 | -1.960 | -1.000 | 0.665 |
| Q9CZT6 | Cmss1 | 0.064 | 0.183 | 0.000 | -1.570 | 0.227 | -1.610 | -2.350 | -1.960 |
| Q99KK1 | Reep3 | 0.256 | 0.470 | 0.000 | -2.660 | 0.078 | 0.631 | 0.147 | 0.263 |
| Q6P5G6 | Ubxn7 | 0.109 | 0.260 | 0.000 | 1.210 | -0.071 | -1.190 | -0.345 | -0.395 |
| Q8K2Y7 | Mrpl47 | 0.048 | 0.152 | 0.000 | 0.273 | -0.157 | 1.070 | 0.895 | 0.325 |
| Q9CPQ1 | Cox6c | 0.008 | 0.052 | 0.000 | 0.148 | -0.102 | 0.358 | 0.542 | 0.504 |
| O54833 | Csnk2a2 | 0.725 | 1.000 | 0.164 | -0.263 | 0.000 | 0.721 | -0.602 | -0.770 |
| Q9CRG1 | Tm7sf3 | 0.667 | 0.945 | 0.418 | -0.002 | 0.000 | 0.609 | -0.354 | -0.319 |
| Q9CQA5 | Med4 | 0.999 | 1.000 | 0.000 | -2.040 | 1.460 | 0.168 | -1.730 | 0.990 |
| Q60676 | Ppp5c | 0.129 | 0.291 | 0.000 | -1.090 | 0.502 | -0.225 | -4.320 | -4.210 |
| P62743 | Ap2s1 | 0.350 | 0.592 | 0.421 | -1.680 | 0.000 | 0.710 | 0.072 | 0.099 |
| Q99N85 | Mrps18a | 0.010 | 0.060 | 0.176 | -0.091 | 0.000 | 0.951 | 0.506 | 0.708 |
| Q9CR11 | Yeats4 | 0.614 | 0.888 | 0.000 | 2.770 | -1.020 | -0.268 | 1.670 | 2.690 |
| Q9D8X2 | Ccdc124 | 0.201 | 0.397 | 0.000 | -0.531 | 0.050 | -0.129 | -1.780 | -0.916 |
| Q9D832 | Dnajb4 | 0.901 | 1.000 | 0.150 | 0.000 | -0.565 | 0.480 | -0.352 | -0.401 |

| | | | | | | | | | |
|---|---|---|---|---|---|---|---|---|---|
| A2A8Z1 | Osbpl9 | 0.215 | 0.412 | 0.344 | 0.000 | -0.424 | 0.033 | -0.872 | -0.935 |
| P14069 | S100a6 | 0.187 | 0.376 | -0.046 | 0.238 | 0.000 | -0.128 | -0.390 | 0.011 |
| O55100 | Syngr1 | 0.000 | 0.013 | 0.248 | -0.546 | 0.000 | 4.430 | 3.640 | 4.470 |
| Q6PB44 | Ptpn23 | 0.645 | 0.922 | 0.000 | 0.015 | -0.813 | 0.790 | -0.854 | 0.086 |
| P63030 | Mpc1 | 0.002 | 0.028 | 0.158 | -0.024 | 0.000 | 1.540 | 0.979 | 1.210 |
| Q921N7 | Tmem70 | 0.029 | 0.111 | -0.295 | 3.890 | 0.000 | 5.300 | 5.780 | 6.340 |
| Q9D920 | Borcs5 | 0.162 | 0.340 | 0.264 | -0.348 | 0.000 | -0.244 | -0.500 | -0.331 |
| Q9CRA4 | Msmo1 | 0.001 | 0.019 | 0.000 | -1.340 | 0.329 | 4.080 | 4.320 | 4.870 |
| P97346 | Nxn | 0.333 | 0.570 | 0.333 | 0.000 | -1.090 | -2.170 | 0.304 | -1.790 |
| Q91VX2 | Ubap2 | 0.246 | 0.457 | 0.280 | -0.451 | 0.000 | 0.798 | -0.235 | 1.300 |
| Q9R0Q6 | Arpc1a | 0.701 | 0.977 | 0.000 | 0.404 | -0.663 | -0.086 | -0.302 | -0.265 |
| Q9CYK1 | Wars2 | 0.461 | 0.720 | -0.361 | 0.518 | 0.000 | 0.617 | -0.291 | 0.913 |
| Q7TMQ7 | Wdr91 | 0.770 | 1.000 | 0.000 | 0.261 | -0.766 | -0.437 | -1.500 | 0.756 |
| Q6DFX2 | Antxr2 | 0.938 | 1.000 | 0.081 | -4.330 | 0.000 | 0.051 | -0.084 | -3.740 |
| Q6PCN7 | Hltf | 0.161 | 0.339 | -0.167 | 0.000 | 0.167 | -0.166 | -2.240 | -0.792 |
| Q99LI8 | Hgs | 0.616 | 0.891 | 0.000 | -3.760 | 1.220 | 0.693 | -0.749 | 0.054 |
| O70456 | Sfn | 0.035 | 0.123 | 0.000 | -0.144 | 0.061 | -0.606 | -2.040 | -1.400 |
| Q6NVF4 | Helb | 0.030 | 0.113 | 0.000 | 0.432 | -0.446 | -3.290 | -2.260 | -1.130 |
| Q61586 | Gpam | 0.000 | 0.012 | 0.000 | -0.259 | 0.263 | 2.000 | 1.940 | 2.100 |
| Q8VC70 | Rbms2 | 0.050 | 0.155 | 0.875 | -3.560 | 0.000 | 4.120 | 2.420 | 2.850 |
| Q8VDV8 | Mitd1 | 0.674 | 0.950 | 0.863 | 0.000 | -0.130 | -0.222 | 0.264 | 0.219 |
| A2BDX3 | Mocs3 | 0.016 | 0.079 | -0.131 | 0.015 | 0.000 | -0.574 | -0.554 | -0.274 |
| P63073 | Eif4e | 0.110 | 0.261 | 0.056 | 0.000 | -0.079 | -0.067 | -0.746 | -0.443 |
| Q8BYL4 | Yars2 | 0.539 | 0.807 | 0.319 | -1.240 | 0.000 | 0.125 | -1.890 | -0.671 |
| Q9JHI9 | Slc40a1 | 0.774 | 1.000 | 1.360 | -0.984 | 0.000 | 1.010 | -0.349 | 0.438 |
| P41233 | Abca1 | 0.053 | 0.162 | 2.580 | -1.450 | 0.000 | 3.720 | 3.300 | 3.800 |
| Q8C052 | Map1s | 0.324 | 0.559 | 1.440 | 0.000 | -0.235 | -2.860 | 0.557 | -0.363 |
| Q80TA6 | Mtmr12 | 0.017 | 0.081 | -0.216 | 0.000 | 0.039 | -0.710 | -1.700 | -1.270 |
| Q920L1 | Fads1 | 0.002 | 0.022 | 0.753 | 0.000 | -1.200 | 4.390 | 4.140 | 4.320 |
| Q6NZQ4 | Paxip1 | 0.002 | 0.024 | -0.702 | 0.000 | 0.105 | -2.740 | -2.460 | -2.170 |
| Q9D786 | Haus5 | 0.097 | 0.240 | -0.007 | 0.000 | 0.031 | -0.178 | -1.330 | -0.640 |
| A2AGH6 | Med12 | 0.669 | 0.947 | -0.883 | 0.364 | 0.000 | -0.436 | -0.859 | 0.129 |
| Q8VDG5 | Ppcs | 0.001 | 0.015 | 0.000 | -0.088 | 0.459 | -1.780 | -2.250 | -2.160 |
| Q8BX17 | Gemin5 | 0.136 | 0.300 | 0.000 | -0.197 | 0.340 | -0.034 | -0.557 | -0.653 |
| Q60949 | Tbc1d1 | 0.929 | 1.000 | -2.370 | 0.000 | 0.522 | -0.861 | 0.527 | -1.210 |
| P34152 | Ptk2 | 0.297 | 0.524 | 0.207 | 0.000 | -0.153 | 0.088 | -0.575 | -0.244 |
| Q8VCH6 | Dhcr24 | 0.003 | 0.030 | -1.970 | 0.475 | 0.000 | 4.270 | 4.540 | 4.840 |
| Q3UFY7 | Nt5c3b | 0.132 | 0.296 | 1.580 | 0.000 | -1.680 | 2.400 | 1.420 | 1.670 |
| P36536 | Sar1a | 0.972 | 1.000 | 0.038 | -1.740 | 0.000 | -0.594 | -0.758 | -0.420 |
| O70433 | Fhl2 | 0.300 | 0.529 | 0.301 | -2.800 | 0.000 | 1.230 | -0.207 | 0.303 |
| P47811 | Mapk14 | 0.232 | 0.436 | 0.000 | 0.568 | -0.624 | -3.770 | -0.803 | -0.291 |
| Q9ERI6 | Rdh14 | 0.189 | 0.379 | 0.000 | 0.132 | -0.126 | -0.072 | -0.191 | -0.680 |
| O88561 | Slc27a3 | 0.002 | 0.025 | 0.000 | 1.250 | -0.059 | 3.450 | 3.940 | 4.190 |
| Q8C1E7 | Tmem120a | 0.021 | 0.090 | 0.000 | 1.190 | -0.322 | 3.090 | 2.270 | 1.920 |
| P97765 | Wbp2 | 0.002 | 0.024 | -0.004 | 0.131 | 0.000 | -1.010 | -1.590 | -1.550 |
| Q3U0M1 | Trappc9 | 0.817 | 1.000 | -0.454 | 0.109 | 0.000 | -0.300 | -0.560 | 0.289 |
| Q8R313 | Exoc6 | 0.164 | 0.343 | 0.374 | 0.000 | -0.064 | -5.790 | -1.000 | -1.050 |
| A6X8Z5 | Arhgap31 | 0.879 | 1.000 | 1.430 | -1.790 | 0.000 | 0.384 | -2.660 | 2.810 |
| Q9D6U8 | Fam162a | 0.000 | 0.013 | -0.044 | 0.195 | 0.000 | 1.030 | 0.901 | 0.963 |
| Q8K3C3 | Lzic | 0.143 | 0.310 | 0.371 | -1.190 | 0.000 | -0.852 | -1.480 | -1.240 |
| Q9D125 | Mrps25 | 0.125 | 0.285 | 5.570 | -3.410 | 0.000 | 6.160 | 5.650 | 5.610 |
| Q9R049 | Amfr | 0.388 | 0.636 | -0.100 | 0.870 | 0.000 | 0.513 | 0.788 | 0.419 |
| Q9DB34 | Chmp2a | 0.798 | 1.000 | 0.119 | -0.014 | 0.000 | 0.221 | -0.174 | 0.165 |
| Q5U4C3 | Scaf1 | 0.618 | 0.892 | 0.000 | -0.994 | 0.360 | 0.667 | -1.780 | -0.847 |
| Q8BKX6 | Smg1 | 0.474 | 0.733 | 0.353 | -1.190 | 0.000 | -0.243 | -1.940 | -0.363 |
| P49586 | Pcyt1a | 0.554 | 0.822 | 0.423 | 0.000 | -0.798 | 0.117 | -0.722 | -0.634 |
| Q9DC29 | Abcb6 | 0.117 | 0.272 | 0.022 | -0.201 | 0.000 | -0.174 | -0.821 | -0.399 |
| O54998 | Fkbp7 | 0.022 | 0.095 | -0.509 | 2.610 | 0.000 | 4.180 | 3.980 | 4.750 |
| Q8VDZ4 | Zdhhc5 | 0.251 | 0.463 | 0.000 | -0.624 | 1.340 | 0.439 | 1.390 | 1.660 |

| | | | | | | | | | |
|---|---|---|---|---|---|---|---|---|---|
| Q61037 | Tsc2 | 0.494 | 0.759 | -0.314 | 0.000 | 0.272 | 0.238 | -0.069 | 0.232 |
| Q9CZ57 | Nsun4 | 0.430 | 0.686 | 0.129 | -1.190 | 0.000 | 0.559 | -0.895 | 1.310 |
| P62309 | Snrpg | 0.091 | 0.232 | 0.000 | 0.621 | -0.692 | -0.743 | -1.190 | -0.818 |
| Q80U70 | Suz12 | 0.113 | 0.266 | 0.000 | 0.065 | -0.310 | -0.990 | -2.960 | -0.684 |
| O35344 | Kpna3 | 0.032 | 0.118 | 0.000 | 0.111 | -0.203 | -0.415 | -1.000 | -0.587 |
| Q9DAT5 | Trmu | 0.965 | 1.000 | 0.330 | -0.121 | 0.000 | 0.746 | -0.391 | -0.095 |
| O88811 | Stam2 | 0.844 | 1.000 | 0.793 | -0.066 | 0.000 | -2.650 | 4.530 | 0.183 |
| P52623 | Uck1 | 0.106 | 0.254 | 0.000 | -0.181 | 0.131 | -0.078 | -0.618 | -0.720 |
| Q60575 | Kif1b | 0.549 | 0.817 | 0.000 | 0.476 | -0.013 | 1.190 | -2.150 | -0.500 |
| Q8BZQ7 | Anapc2 | 0.858 | 1.000 | 0.000 | 0.388 | -1.980 | 1.800 | -1.690 | -2.560 |
| P26011 | Itgb7 | 0.206 | 0.403 | -0.370 | 1.190 | 0.000 | -0.580 | -0.272 | -0.497 |
| B2RX14 | Zcchc11 | 0.288 | 0.513 | 0.813 | -0.294 | 0.000 | 0.265 | -2.670 | -0.521 |
| O35130 | Emg1 | 0.066 | 0.186 | 0.220 | -0.200 | 0.000 | -0.494 | -2.150 | -1.090 |
| Q8CD26 | Slc35e1 | 0.070 | 0.193 | 1.180 | -0.035 | 0.000 | 2.010 | 1.630 | 1.070 |
| Q69ZC8 | Gpalpp1 | 0.047 | 0.150 | -0.158 | 1.170 | 0.000 | 3.410 | 1.640 | 1.870 |
| Q9JM13 | Rabgef1 | 0.270 | 0.489 | 0.000 | 0.936 | -0.963 | 0.109 | -2.890 | -1.190 |
| Q5SS80 | Dhrs13 | 0.001 | 0.019 | -0.233 | 0.000 | 0.067 | 1.520 | 1.110 | 1.520 |
| Q3UE37 | Ube2z | 0.134 | 0.298 | 0.000 | -0.126 | 0.232 | 0.110 | -2.060 | -2.380 |
| Q7TN58 | Tmc8 | 0.075 | 0.202 | -0.585 | 0.000 | 0.174 | -1.240 | -1.290 | -0.445 |
| Q8BLR5 | Psd4 | 0.060 | 0.175 | 0.000 | -0.228 | 0.469 | -1.850 | -2.990 | -0.583 |
| Q3UL36 | Arglu1 | 0.210 | 0.407 | 0.098 | -0.107 | 0.000 | 0.196 | -0.943 | -1.200 |
| P34884 | Mif | 0.193 | 0.385 | -0.437 | 0.000 | 0.174 | -0.562 | -4.920 | -1.210 |
| Q8VC65 | Nrm | 0.086 | 0.222 | 0.000 | 0.145 | -0.254 | 0.254 | 0.325 | 0.165 |
| Q03141 | Mark3 | 0.872 | 1.000 | 0.347 | 0.000 | -0.830 | 0.273 | -0.380 | -0.172 |
| Q3TWF6 | Wdr70 | 0.840 | 1.000 | 0.000 | 2.870 | -1.380 | 0.191 | 1.450 | -1.090 |
| Q99NH0 | Ankrd17 | 0.978 | 1.000 | 0.000 | -0.078 | 0.040 | 0.641 | -0.560 | -0.088 |
| Q9D1R1 | Tmem126b | 0.441 | 0.698 | 0.146 | -0.371 | 0.000 | -0.573 | 0.779 | 0.798 |
| Q99LI9 | Clp1 | 0.049 | 0.154 | 0.000 | -0.466 | 0.106 | -0.628 | -0.539 | -0.741 |
| Q8BKI2 | Tnrc6b | 0.181 | 0.368 | 0.385 | -0.531 | 0.000 | -0.390 | -0.386 | -0.939 |
| O35216 | Cenpa | 0.114 | 0.268 | 0.001 | -0.082 | 0.000 | -0.253 | -0.531 | -0.092 |
| Q6PIJ4 | Nfrkb | 0.972 | 1.000 | -1.190 | 0.727 | 0.000 | 0.003 | -0.075 | -0.451 |
| Q3U487 | Hectd3 | 0.509 | 0.773 | -0.887 | 0.876 | 0.000 | 0.088 | -1.320 | -0.223 |
| P59178 | L3mbtl2 | 0.800 | 1.000 | 0.000 | 0.358 | -1.860 | 0.594 | -0.726 | -2.230 |
| A6H630 | Armt1 | 0.012 | 0.067 | 0.101 | -0.391 | 0.000 | -1.820 | -4.060 | -3.390 |
| P83887 | Tubg1 | 0.204 | 0.400 | -0.306 | 0.000 | 0.031 | -0.359 | -0.367 | -0.142 |
| Q7TPN9 | Prr14 | 0.091 | 0.231 | -0.071 | 0.000 | 0.554 | -0.152 | -0.582 | -0.336 |
| Q6ZQE4 | Nemp1 | 0.975 | 1.000 | 0.000 | -0.209 | 0.239 | 0.125 | -0.173 | 0.093 |
| Q9QYR6 | Map1a | 0.277 | 0.498 | -1.020 | 0.047 | 0.000 | -0.594 | -0.990 | -0.769 |
| Q8K296 | Mtmr3 | 0.128 | 0.290 | 0.000 | 0.462 | -0.686 | -0.643 | -0.845 | -0.685 |
| Q68FE8 | Znf280d | 0.869 | 1.000 | 0.000 | -3.330 | 0.418 | -1.590 | -1.570 | -0.412 |
| Q3TQB2 | Foxred1 | 0.796 | 1.000 | -0.086 | 0.930 | 0.000 | 0.212 | 0.095 | 0.873 |
| Q9WU62 | Incenp | 0.140 | 0.306 | 0.524 | 0.000 | -0.282 | -0.029 | -0.934 | -0.856 |
| Q9CQ91 | Ndufa3 | 0.595 | 0.868 | -0.286 | 0.062 | 0.000 | 0.320 | -0.326 | 0.166 |
| Q8CIG8 | Prmt5 | 0.035 | 0.124 | 0.000 | 0.259 | -0.546 | -0.723 | -1.530 | -1.800 |
| P30677 | Gna14 | 0.352 | 0.593 | 0.000 | 2.050 | -0.325 | 3.440 | 0.286 | 1.720 |
| Q8C5P5 | Nt5dc1 | 0.002 | 0.025 | -0.291 | 0.000 | 0.247 | -1.180 | -1.430 | -1.520 |
| P30285 | Cdk4 | 0.927 | 1.000 | -1.000 | 0.000 | 0.348 | 0.457 | -0.400 | -0.875 |
| Q99JW2 | Acy1 | 0.219 | 0.418 | -1.060 | 3.310 | 0.000 | -0.115 | -1.590 | -2.610 |
| Q9CS42 | Prps2 | 0.013 | 0.070 | 0.016 | -0.356 | 0.000 | -0.738 | -1.420 | -1.340 |
| P23772 | Gata3 | 0.502 | 0.767 | -6.200 | 0.000 | 0.195 | -0.520 | -0.462 | -0.382 |
| Q8BGR9 | Ublcp1 | 0.647 | 0.923 | 0.155 | -1.020 | 0.000 | -0.287 | -0.813 | -0.360 |
| Q9EQG9 | Col4a3bp | 0.061 | 0.177 | 0.743 | 0.000 | -0.010 | -0.270 | -1.100 | -0.586 |
| Q99KW3 | Triobp | 0.447 | 0.704 | 1.350 | 0.000 | -0.569 | 1.560 | 0.174 | 0.802 |
| P49446 | Ptpre | 0.361 | 0.607 | 0.248 | 0.000 | -0.421 | 0.026 | -0.348 | -0.818 |
| P81069 | Gabpb2 | 0.101 | 0.247 | -0.582 | 0.000 | 0.001 | -0.633 | -0.904 | -0.497 |
| Q9CPR8 | Nsmce3 | 0.214 | 0.411 | 0.219 | -0.997 | 0.000 | -0.521 | -1.170 | -0.944 |
| Q99N95 | Mrpl3 | 0.067 | 0.187 | 0.000 | 0.558 | -0.395 | 1.000 | 0.576 | 0.857 |
| Q9R1C0 | Taf7 | 0.309 | 0.542 | 0.000 | 0.218 | -0.124 | -0.012 | -0.048 | -0.424 |
| Q9QZS3 | Numb | 0.846 | 1.000 | 0.000 | -2.960 | 1.490 | -1.000 | -0.066 | 0.457 |

| | | | | | | | | | |
|---|---|---|---|---|---|---|---|---|---|
| Q9JIB4 | Gtf2h2 | 0.243 | 0.453 | 0.000 | -0.382 | 0.134 | -0.180 | -1.860 | -0.442 |
| Q9DB10 | Smdt1 | 0.087 | 0.224 | -0.402 | 0.296 | 0.000 | 0.541 | 0.458 | 0.326 |
| A2A6Q5 | Cdc27 | 0.004 | 0.038 | 0.037 | 0.000 | -0.024 | 1.120 | 0.633 | 0.785 |
| Q8CG50 | Rab43 | 0.799 | 1.000 | -3.230 | 0.000 | 0.170 | 0.428 | -0.190 | -4.960 |
| O88796 | Rpp30 | 0.185 | 0.374 | 0.000 | 0.657 | -0.507 | 1.050 | 0.269 | 0.787 |

**Supplementary Table 2.** T cell exhaustion associated proteins detected in CD4+ T cells from old mice with or without mito-transfer. CD4+ T cells from young and old mice, and from old mice after mito-transfer were cultured for 4 h before processing for mass spectrometry analysis. Data expressed as median protein Log2 fold change of CD4+ T cells from old mice, from 3 individual old mice (paired experiment).

| GO | Category | Description | Count | % | Log10(P) | Log10(q) |
|---|---|---|---|---|---|---|
| GO:0045333 | GO BP | cellular respiration | 37 | 12.71 | -31.24 | -26.97 |
| mmu04141 | KEGG | Protein processing in endoplasmic reticulum - Mus musculus (house mouse) | 29 | 9.97 | -22.64 | -19.77 |
| mmu00100 | KEGG | Steroid biosynthesis - Mus musculus (house mouse) | 9 | 3.09 | -11.64 | -9.19 |
| R-MMU-1592230 | Reactome | Mitochondrial biogenesis | 9 | 3.09 | -10.23 | -7.82 |
| GO:0006637 | GO BP | acyl-CoA metabolic process | 13 | 4.47 | -9.97 | -7.57 |
| GO:0061024 | GO BP | membrane organization | 34 | 11.68 | -9.92 | -7.53 |
| GO:0006457 | GO BP | protein folding | 17 | 5.84 | -9.74 | -7.36 |
| GO:0043603 | GO BP | amide metabolic process | 35 | 12.03 | -8.81 | -6.49 |
| GO:0090150 | GO BP | establishment of protein localization to membrane | 18 | 6.19 | -8.81 | -6.49 |
| mmu01200 | KEGG | Carbon metabolism - Mus musculus (house mouse) | 13 | 4.47 | -7.9 | -5.62 |
| GO:0018126 | GO BP | protein hydroxylation | 7 | 2.41 | -7.1 | -4.84 |
| WP662 | WikiPathwa | Amino acid metabolism | 11 | 3.78 | -7.06 | -4.81 |
| CORUM:413 | CORUM | (ER)-localized multiprotein complex, Ig heavy chains associated | 5 | 1.72 | -6.95 | -4.7 |
| GO:0007007 | GO BP | inner mitochondrial membrane organization | 7 | 2.41 | -6.66 | -4.44 |
| GO:0098660 | GO BP | inorganic ion transmembrane transport | 26 | 8.93 | -6.59 | -4.37 |
| GO:0034976 | GO BP | response to endoplasmic reticulum stress | 15 | 5.15 | -6.12 | -3.96 |
| R-MMU-8949664 | Reactome | Processing of SMDT1 | 5 | 1.72 | -5.74 | -3.6 |
| GO:0031647 | GO BP | regulation of protein stability | 17 | 5.84 | -5.7 | -3.57 |
| GO:0006851 | GO BP | mitochondrial calcium ion transmembrane transport | 5 | 1.72 | -5.59 | -3.48 |
| GO:1902930 | GO BP | regulation of alcohol biosynthetic process | 7 | 2.41 | -5.11 | -3.03 |

**Supplementary Table 3.** The Top 20 processes of significantly upregulated DEPs in CD4+ T cells from old mice, with or without mito-transfer in relation to T Cell exhaustion. Data generated through Pathway & Process Enrichment (PPE) analysis using the Metascape analysis platform.

| GO | Description | Log10(P) |
|---|---|---|
| GO:0045333 | cellular respiration | -33.8 |
| R-MMU-1428517 | The citric acid (TCA) cycle and respiratory electron transport | -33.3 |
| GO:0009060 | aerobic respiration | -32.8 |

**Supplementary Table 4.** The Top 3 Protein-Protein Interaction (PPI) networks in CD4+ T cells from old mice, with or without mito-transfer, in relation to T Cell exhaustion. Data generated using the Metascape analysis platform.

| MCODE | GO | Description | Log10(P) |
|---|---|---|---|
| MCODE_1 | R-MMU-163200 | Respiratory electron transport, ATP synthesis by chemiosmotic coupling, and heat production by uncoupling proteins. | -48.1 |
| MCODE_1 | mmu00190 | Oxidative phosphorylation - Mus musculus (house mouse) | -46.8 |
| MCODE_1 | WP295 | Electron transport chain | -46.8 |
| MCODE_2 | GO:0032543 | mitochondrial translation | -22.7 |
| MCODE_2 | R-MMU-5389840 | Mitochondrial translation elongation | -21.6 |
| MCODE_2 | R-MMU-5419276 | Mitochondrial translation termination | -21.4 |
| MCODE_3 | mmu04141 | Protein processing in endoplasmic reticulum - Mus musculus (house mouse) | -21 |
| MCODE_3 | mmu00513 | Various types of N-glycan biosynthesis - Mus musculus (house mouse) | -17 |
| MCODE_3 | GO:0018279 | protein N-linked glycosylation via asparagine | -16.5 |
| MCODE_4 | GO:0071681 | cellular response to indole-3-methanol | -9.1 |
| MCODE_4 | GO:0071680 | response to indole-3-methanol | -9.1 |
| MCODE_4 | mmu05200 | Pathways in cancer - Mus musculus (house mouse) | -9.1 |
| MCODE_5 | GO:0030036 | actin cytoskeleton organization | -4.6 |
| MCODE_5 | GO:0030029 | actin filament-based process | -4.4 |
| MCODE_5 | mmu04510 | Focal adhesion - Mus musculus (house mouse) | -4.3 |
| MCODE_6 | mmu00280 | Valine, leucine and isoleucine degradation - Mus musculus (house mouse) | -8.8 |
| MCODE_7 | mmu04141 | Protein processing in endoplasmic reticulum - Mus musculus (house mouse) | -12.6 |
| MCODE_7 | GO:0006457 | protein folding | -7.2 |
| MCODE_7 | GO:0034976 | response to endoplasmic reticulum stress | -6.7 |
| MCODE_8 | mmu00100 | Steroid biosynthesis - Mus musculus (house mouse) | -18.5 |
| MCODE_8 | R-MMU-191273 | Cholesterol biosynthesis | -17.5 |
| MCODE_8 | GO:1902653 | secondary alcohol biosynthetic process | -16.4 |
| MCODE_9 | GO:0009060 | aerobic respiration | -10.7 |
| MCODE_9 | R-MMU-1428517 | The citric acid (TCA) cycle and respiratory electron transport | -10.6 |
| MCODE_9 | GO:0009152 | purine ribonucleotide biosynthetic process | -10.5 |
| MCODE_10 | mmu04141 | Protein processing in endoplasmic reticulum - Mus musculus (house mouse) | -5.7 |
| MCODE_11 | WP103 | Cholesterol biosynthesis | -8.9 |
| MCODE_11 | mmu00100 | Steroid biosynthesis - Mus musculus (house mouse) | -8.5 |
| MCODE_11 | R-MMU-191273 | Cholesterol biosynthesis | -8.1 |
| MCODE_12 | R-MMU-1650814 | Collagen biosynthesis and modifying enzymes | -7.7 |
| MCODE_12 | mmu00310 | Lysine degradation - Mus musculus (house mouse) | -7.6 |
| MCODE_12 | R-MMU-1474290 | Collagen formation | -7.4 |
| MCODE_13 | GO:0010817 | regulation of hormone levels | -4.6 |
| MCODE_13 | GO:0014070 | response to organic cyclic compound | -4.3 |
| MCODE_14 | GO:0006457 | protein folding | -6.3 |
| MCODE_14 | GO:0051604 | protein maturation | -5 |

**Supplementary Table 5.** The MCODE networks identified for significantly upregulated DEPs in CD4+ T cells from old mice, with or without mito-transfer, in relation to T Cell exhaustion. Data generated using the Metascape analysis platform.

| GO | Category | Description | Count | % | Log10(P) | Log10(q) |
|---|---|---|---|---|---|---|
| R-MMU-8953854 | Reactome | Metabolism of RNA | 55 | 19.57 | -29.98 | -25.71 |
| R-MMU-450531 | Reactome | Regulation of mRNA stability by proteins that bind AU-rich elements | 23 | 8.19 | -23.34 | -19.38 |
| mmu00010 | KEGG | Glycolysis / Gluconeogenesis - Mus musculus (house mouse) | 11 | 3.91 | -8.91 | -6.72 |
| R-MMU-6798695 | Reactome | Neutrophil degranulation | 27 | 9.61 | -8.71 | -6.53 |
| GO:0030029 | GO BP | actin filament-based process | 28 | 9.96 | -8.37 | -6.2 |
| GO:0034655 | GO BP | nucleobase-containing compound catabolic process | 18 | 6.41 | -7.59 | -5.44 |
| mmu00270 | KEGG | Cysteine and methionine metabolism - Mus musculus (house mouse) | 9 | 3.2 | -7.38 | -5.24 |
| GO:0050684 | GO BP | regulation of mRNA processing | 13 | 4.63 | -7.35 | -5.22 |
| GO:0098761 | GO BP | cellular response to interleukin-7 | 6 | 2.14 | -7.25 | -5.13 |
| WP572 | WikiPathwa | EGFR1 signaling pathway | 14 | 4.98 | -7.01 | -4.92 |
| R-MMU-194315 | Reactome | Signaling by Rho GTPases | 25 | 8.9 | -6.68 | -4.6 |
| GO:0010563 | GO BP | negative regulation of phosphorus metabolic process | 21 | 7.47 | -6.47 | -4.4 |
| GO:0052548 | GO BP | regulation of endopeptidase activity | 19 | 6.76 | -6.41 | -4.34 |
| GO:0044283 | GO BP | small molecule biosynthetic process | 20 | 7.12 | -5.57 | -3.56 |
| GO:0031123 | GO BP | RNA 3'-end processing | 9 | 3.2 | -5.51 | -3.51 |
| WP2185 | WikiPathwa | Purine metabolism | 12 | 4.27 | -5.47 | -3.47 |
| WP387 | WikiPathwa | IL-6 signaling pathway | 9 | 3.2 | -5.16 | -3.19 |
| R-MMU-109606 | Reactome | Intrinsic Pathway for Apoptosis | 6 | 2.14 | -5 | -3.04 |
| GO:0071364 | GO BP | cellular response to epidermal growth factor stimulus | 6 | 2.14 | -5 | -3.04 |
| GO:0022613 | GO BP | ribonucleoprotein complex biogenesis | 18 | 6.41 | -4.75 | -2.82 |

**Supplementary Table 6.** The Top 20 processes of significantly downregulated DEPs in CD4+ T cells from old mice, with or without mito-transfer, in relation to T Cell exhaustion. Data generated through Pathway & Process Enrichment (PPE) analysis using the Metascape analysis platform.

| GO | Description | Log10(P) |
|---|---|---|
| R-MMU-8953854 | Metabolism of RNA | -30.6 |
| R-MMU-450531 | Regulation of mRNA stability by proteins that bind AU-rich elements | -24.5 |
| R-MMU-1236978 | Cross-presentation of soluble exogenous antigens (endosomes) | -22 |

**Supplementary Table 7.** The Top 3 Protein-Protein Interaction (PPI) networks of significantly downregulated DEPs in CD4+ T cells from old mice, with or without mito-transfer, in relation to T Cell exhaustion. Data generated using the Metascape analysis platform.

| MCODE | GO | Description | Log10(P) |
|---|---|---|---|
| MCODE_1 | R-MMU-1169091 | Activation of NF-kappaB in B cells | -47 |
| MCODE_1 | R-MMU-1234176 | hydroxylation of Hypoxia-inducible Factor Alpha | -46.5 |
| MCODE_1 | R-MMU-1234174 | Cellular response to hypoxia | -46 |
| MCODE_2 | mmu00010 | Glycolysis / Gluconeogenesis - Mus musculus (house mouse) | -13.4 |
| MCODE_2 | WP157 | Glycolysis and gluconeogenesis | -9.5 |
| MCODE_2 | GO:0006007 | glucose catabolic process | -8.5 |
| MCODE_3 | R-MMU-75035 | Chk1/Chk2(Cds1) mediated inactivation of Cyclin B:Cdk1 complex | -10 |
| MCODE_3 | R-MMU-111447 | mitochondria | -10 |
| MCODE_3 | R-MMU-114452 | Activation of BH3-only proteins | -9.2 |
| MCODE_4 | mmu03040 | mouse) | -17 |
| MCODE_4 | R-MMU-72163 | mRNA Splicing - Major Pathway | -16.7 |
| MCODE_4 | R-MMU-72172 | mRNA Splicing | -16.6 |
| MCODE_5 | GO:1902115 | regulation of organelle assembly | -3.7 |
| MCODE_5 | GO:0032535 | regulation of cellular component size | -3 |
| MCODE_5 | GO:0043254 | complex assembly | -2.9 |
| MCODE_6 | GO:0016051 | carbohydrate biosynthetic process | -4.6 |
| MCODE_6 | GO:0006006 | glucose metabolic process | -4.5 |
| MCODE_6 | GO:0019318 | hexose metabolic process | -4.2 |
| MCODE_7 | WP407 | Kit receptor signaling pathway | -8.9 |
| MCODE_7 | mmu04658 | Th1 and Th2 cell differentiation - Mus musculus (house mouse) | -8.4 |
| MCODE_7 | R-MMU-8854691 | Interleukin-20 family signaling | -8.3 |
| MCODE_8 | R-MMU-6807505 | genes | -9.2 |
| MCODE_8 | GO:0006366 | transcription by RNA polymerase II | -6.5 |
| MCODE_8 | GO:0006351 | DNA-templated transcription | -5.9 |
| MCODE_9 | GO:0097435 | supramolecular fiber organization | -4.1 |
| MCODE_10 | R-MMU-6791226 | Major pathway of rRNA processing in the nucleolus and cytosol | -6.2 |
| MCODE_10 | R-MMU-72312 | rRNA processing | -6.2 |
| MCODE_10 | R-MMU-8868773 | cytosol | -6.2 |

**Supplementary Table 8.** The MCODE networks identified significantly downregulated DEPs in CD4+ T cells from old mice, with or without mito-transfer, in relation to T Cell exhaustion. Data generated using the Metascape analysis platform.

| Uniprot ID | Gene Symbol | t-test | FDR (adj. P-val.) | O1 | O2 | O3 | OM1 | OM2 | OM3 |
|---|---|---|---|---|---|---|---|---|---|
| P16125 | Ldhb | 0.000 | 0.013 | -0.212 | 0.132 | 0.000 | -1.330 | -1.570 | -1.550 |
| P08228 | Sod1 | 0.000 | 0.013 | -0.224 | 0.010 | 0.000 | -1.080 | -1.290 | -1.170 |
| Q922W5 | Pycr1 | 0.000 | 0.014 | 0.208 | 0.000 | -0.219 | 2.510 | 1.920 | 2.420 |
| P63101 | Ywhaz | 0.001 | 0.015 | 0.000 | -0.021 | 0.113 | -1.140 | -1.610 | -1.370 |
| P06151 | Ldha | 0.001 | 0.018 | -0.165 | 0.115 | 0.000 | -0.980 | -1.190 | -0.934 |
| Q9DB20 | Atp5o | 0.001 | 0.019 | 0.000 | 0.129 | -0.067 | 0.904 | 0.673 | 0.901 |
| P23506 | Pcmt1 | 0.001 | 0.022 | -0.179 | 0.000 | 0.198 | -0.923 | -0.926 | -0.835 |
| Q8R0F3 | Sumf1 | 0.002 | 0.024 | 0.000 | 0.232 | -0.568 | 1.860 | 1.650 | 1.660 |
| Q01853 | Vcp | 0.002 | 0.025 | 0.000 | 0.056 | -0.134 | -0.465 | -0.616 | -0.590 |
| P17182 | Eno1 | 0.002 | 0.028 | 0.055 | -0.141 | 0.000 | -0.948 | -1.530 | -1.280 |
| P48678 | Lmna | 0.002 | 0.028 | 0.000 | 0.046 | -0.348 | 0.967 | 0.745 | 0.972 |
| P35235 | Ptpn11 | 0.002 | 0.029 | -0.286 | 0.010 | 0.000 | -0.906 | -1.190 | -0.916 |
| Q04207 | Rela | 0.003 | 0.030 | 0.000 | -0.006 | 0.032 | -0.840 | -1.440 | -1.170 |
| P27641 | Xrcc5 | 0.003 | 0.030 | -0.102 | 0.000 | 0.052 | -1.050 | -1.770 | -1.530 |
| P28474 | Adh5 | 0.003 | 0.032 | -0.238 | 0.142 | 0.000 | -1.150 | -1.810 | -1.400 |
| Q62077 | Plcg1 | 0.003 | 0.033 | -0.125 | 0.000 | 0.471 | -1.190 | -1.370 | -1.750 |
| P19157 | Gstp1 | 0.004 | 0.036 | -0.138 | 0.000 | 0.106 | -1.050 | -1.380 | -0.809 |
| P00493 | Hprt1 | 0.004 | 0.038 | 0.000 | 0.060 | -0.208 | -1.260 | -1.950 | -1.240 |
| Q9QXS6 | Dbn1 | 0.005 | 0.039 | -1.190 | 2.450 | 0.000 | 7.040 | 6.310 | 6.710 |
| Q9R1E0 | Foxo1 | 0.005 | 0.039 | -0.131 | 0.075 | 0.000 | -0.360 | -0.460 | -0.388 |
| Q60631 | Grb2 | 0.005 | 0.041 | -0.091 | 0.403 | 0.000 | -1.080 | -1.510 | -0.944 |
| P52431 | Pold1 | 0.005 | 0.043 | -0.206 | 0.235 | 0.000 | -0.797 | -1.220 | -0.903 |
| P63038 | Hspd1 | 0.006 | 0.044 | 0.000 | 0.119 | -0.240 | 1.090 | 0.650 | 0.822 |
| P42232 | Stat5b | 0.007 | 0.048 | -0.226 | 0.000 | 0.150 | -0.984 | -1.490 | -0.914 |
| P63017 | Hspa8 | 0.008 | 0.054 | 0.046 | 0.000 | -0.090 | -0.386 | -0.759 | -0.718 |
| P23475 | Xrcc6 | 0.009 | 0.056 | 0.040 | 0.000 | -0.446 | -0.978 | -1.630 | -1.300 |
| O08734 | Bak1 | 0.010 | 0.060 | 0.000 | 0.028 | -0.547 | 0.687 | 0.764 | 0.655 |
| Q9D666 | Sun1 | 0.011 | 0.064 | -0.159 | 0.054 | 0.000 | 0.644 | 0.375 | 0.368 |
| Q06890 | Clu | 0.014 | 0.073 | 0.000 | 0.199 | -0.686 | -1.550 | -2.880 | -1.990 |
| P42230 | Stat5a | 0.015 | 0.076 | -0.331 | 0.072 | 0.000 | -0.731 | -1.550 | -1.370 |
| Q8R4K2 | Irak4 | 0.016 | 0.078 | 0.152 | 0.000 | -0.178 | -0.685 | -1.670 | -1.410 |
| P10639 | Txn | 0.016 | 0.079 | 0.196 | -0.365 | 0.000 | -0.914 | -2.060 | -1.810 |
| P25799 | Nfkb1 | 0.017 | 0.080 | 0.000 | -0.065 | 0.139 | -0.430 | -1.120 | -0.909 |
| P47791 | Gsr | 0.018 | 0.084 | 0.341 | 0.000 | -0.118 | -0.466 | -1.080 | -1.080 |
| P07901 | Hsp90aa1 | 0.021 | 0.090 | 0.000 | 0.010 | -0.335 | -0.557 | -1.180 | -1.010 |
| P09671 | Sod2 | 0.023 | 0.096 | -0.061 | 0.348 | 0.000 | 0.715 | 0.476 | 0.682 |
| P70371 | Terf1 | 0.025 | 0.102 | 0.000 | 0.395 | -0.619 | -1.120 | -1.050 | -1.160 |
| P39428 | Traf1 | 0.026 | 0.103 | 0.000 | 0.232 | -0.135 | -0.817 | -1.800 | -0.841 |
| P28352 | Apex1 | 0.028 | 0.107 | 0.000 | -0.288 | 0.047 | -0.452 | -0.881 | -0.582 |
| Q03147 | Cdk7 | 0.029 | 0.110 | 0.000 | -0.008 | 0.017 | -0.257 | -0.564 | -0.237 |
| P20444 | Prkca | 0.030 | 0.113 | -0.268 | 0.109 | 0.000 | -0.390 | -0.460 | -0.434 |
| P35821 | Ptpn1 | 0.031 | 0.116 | -0.040 | 0.235 | 0.000 | -0.162 | -0.284 | -0.302 |
| P12023 | App | 0.032 | 0.117 | 4.640 | -1.820 | 0.000 | 7.180 | 7.130 | 7.230 |
| P63166 | Sumo1 | 0.032 | 0.119 | -0.103 | 0.000 | 0.363 | -0.366 | -0.411 | -0.342 |
| Q9QVP9 | Ptk2b | 0.038 | 0.131 | -1.180 | 0.133 | 0.000 | -1.480 | -1.840 | -1.650 |
| P43404 | Zap70 | 0.038 | 0.131 | -0.465 | 0.041 | 0.000 | -0.669 | -0.946 | -0.589 |
| Q9Z2A7 | Dgat1 | 0.041 | 0.136 | 0.000 | 0.214 | -0.404 | 0.617 | 0.361 | 0.650 |
| P06537 | Nr3c1 | 0.043 | 0.141 | -0.291 | 0.000 | 0.306 | -0.491 | -0.996 | -0.556 |
| P16858 | Gapdh | 0.043 | 0.142 | -0.051 | 0.026 | 0.000 | -0.153 | -0.557 | -0.357 |
| P24270 | Cat | 0.046 | 0.148 | 0.177 | 0.000 | -0.253 | 1.040 | 0.403 | 0.494 |
| Q8BKJ9 | Sirt7 | 0.048 | 0.152 | 0.000 | -0.263 | 0.089 | -0.314 | -0.839 | -0.582 |
| Q60520 | Sin3a | 0.049 | 0.154 | -0.146 | 0.000 | 0.086 | -0.240 | -0.604 | -0.309 |
| P04202 | Tgfb1 | 0.054 | 0.163 | 0.000 | 0.044 | -0.363 | -0.731 | -0.344 | -0.794 |
| P14733 | Lmnb1 | 0.060 | 0.175 | -0.138 | 0.000 | 0.107 | -0.348 | -0.608 | -0.184 |
| P33609 | Pola1 | 0.060 | 0.176 | 0.058 | 0.000 | -0.834 | -1.570 | -1.450 | -0.752 |

| | | | | | | | | | |
|---|---|---|---|---|---|---|---|---|---|
| O09172 | Gclm | 0.514 | 0.778 | 0.497 | 0.000 | -0.914 | 0.147 | -0.892 | -0.812 |
| Q9QXS6 | Dbn1 | 0.005 | 0.039 | -1.190 | 2.450 | 0.000 | 7.040 | 6.310 | 6.710 |
| Q80W54 | Zmpste24 | 0.080 | 0.211 | 0.027 | 0.000 | -0.220 | 0.715 | 0.060 | 0.482 |
| P11103 | Parp1 | 0.506 | 0.770 | -0.125 | 0.040 | 0.000 | -0.190 | -0.321 | 0.120 |
| P11352 | Gpx1 | 0.131 | 0.294 | 0.457 | 0.000 | -0.235 | -0.028 | -0.804 | -0.877 |
| Q04750 | Top1 | 0.149 | 0.319 | 0.118 | -0.057 | 0.000 | -0.028 | -0.407 | -0.160 |
| P39749 | Fen1 | 0.169 | 0.351 | 0.002 | -0.215 | 0.000 | -0.243 | -0.778 | -0.197 |
| Q9CQB5 | Cisd2 | 0.179 | 0.365 | 0.000 | 0.161 | -0.086 | 0.224 | 0.101 | 0.145 |
| P07901 | Hsp90aa1 | 0.021 | 0.090 | 0.000 | 0.010 | -0.335 | -0.557 | -1.180 | -1.010 |
| Q02111 | Prkcq | 0.722 | 1.000 | 0.000 | 1.430 | -0.213 | -0.229 | 0.752 | 0.016 |
| Q60631 | Grb2 | 0.005 | 0.041 | -0.091 | 0.403 | 0.000 | -1.080 | -1.510 | -0.944 |
| P42230 | Stat5a | 0.015 | 0.076 | -0.331 | 0.072 | 0.000 | -0.731 | -1.550 | -1.370 |
| P17918 | Pcna | 0.142 | 0.309 | 0.516 | 0.000 | -0.203 | -0.004 | -0.794 | -0.676 |
| Q8BKJ9 | Sirt7 | 0.048 | 0.152 | 0.000 | -0.263 | 0.089 | -0.314 | -0.839 | -0.582 |
| P09671 | Sod2 | 0.023 | 0.096 | -0.061 | 0.348 | 0.000 | 0.715 | 0.476 | 0.682 |
| P52431 | Pold1 | 0.005 | 0.043 | -0.206 | 0.235 | 0.000 | -0.797 | -1.220 | -0.903 |
| P23475 | Xrcc6 | 0.009 | 0.056 | 0.040 | 0.000 | -0.446 | -0.978 | -1.630 | -1.300 |
| P98083 | Shc1 | 0.721 | 1.000 | 0.042 | -0.011 | 0.000 | 1.180 | -0.727 | 0.209 |
| P35700 | Prdx1 | 0.130 | 0.293 | 0.028 | 0.000 | -0.030 | 0.421 | -0.006 | 0.325 |
| P70399 | Tp53bp1 | 0.062 | 0.178 | 0.267 | 0.000 | -0.013 | -0.418 | -1.440 | -0.510 |
| Q9Z2A7 | Dgat1 | 0.041 | 0.136 | 0.000 | 0.214 | -0.404 | 0.617 | 0.361 | 0.650 |
| P63280 | Ube2i | 0.764 | 1.000 | 0.000 | 0.245 | -0.255 | -0.134 | -0.038 | 0.016 |
| Q04207 | Rela | 0.003 | 0.030 | 0.000 | -0.006 | 0.032 | -0.840 | -1.440 | -1.170 |
| P16125 | Ldhb | 0.000 | 0.013 | -0.212 | 0.132 | 0.000 | -1.330 | -1.570 | -1.550 |
| Q99J62 | Rfc4 | 0.246 | 0.457 | 0.532 | -0.054 | 0.000 | 0.414 | -0.865 | -1.300 |
| Q08481 | Pecam1 | 0.631 | 0.907 | 0.000 | 1.060 | -0.637 | -0.841 | 0.355 | -0.038 |
| Q6GQT1 | A2m | 0.349 | 0.590 | 0.000 | -0.058 | 0.442 | 0.387 | 0.048 | 0.814 |
| Q8K3H0 | Appl1 | 0.122 | 0.280 | -0.019 | 0.000 | 0.087 | 0.005 | -0.899 | -0.625 |
| P42232 | Stat5b | 0.007 | 0.048 | -0.226 | 0.000 | 0.150 | -0.984 | -1.490 | -0.914 |
| Q9QVP9 | Ptk2b | 0.038 | 0.131 | -1.180 | 0.133 | 0.000 | -1.480 | -1.840 | -1.650 |
| Q9R1E0 | Foxo1 | 0.005 | 0.039 | -0.131 | 0.075 | 0.000 | -0.360 | -0.460 | -0.388 |
| P01901 | H2-K1 | 0.918 | 1.000 | 0.000 | 1.100 | -0.068 | -0.245 | 0.777 | 0.664 |
| P29533 | Vcam1 | 0.146 | 0.315 | 0.746 | 0.000 | -0.120 | 0.114 | -1.040 | -0.893 |
| Q62388 | Atm | 0.107 | 0.257 | -0.265 | 0.116 | 0.000 | -0.451 | -1.290 | -0.368 |
| P62137 | Ppp1ca | 0.822 | 1.000 | 0.237 | -0.916 | 0.000 | -0.030 | -0.612 | -0.318 |
| P20444 | Prkca | 0.030 | 0.113 | -0.268 | 0.109 | 0.000 | -0.390 | -0.460 | -0.434 |
| P41245 | Mmp9 | 0.231 | 0.435 | 2.280 | 0.000 | -0.046 | 0.785 | -1.710 | -1.570 |
| Q03147 | Cdk7 | 0.029 | 0.110 | 0.000 | -0.008 | 0.017 | -0.257 | -0.564 | -0.237 |
| P10126 | Eef1a1 | 0.463 | 0.721 | 0.222 | -0.135 | 0.000 | 0.343 | -0.593 | -0.404 |
| P28474 | Adh5 | 0.003 | 0.032 | -0.238 | 0.142 | 0.000 | -1.150 | -1.810 | -1.400 |
| P10639 | Txn | 0.016 | 0.079 | 0.196 | -0.365 | 0.000 | -0.914 | -2.060 | -1.810 |
| P47791 | Gsr | 0.018 | 0.084 | 0.341 | 0.000 | -0.118 | -0.466 | -1.080 | -1.080 |
| P68181 | Prkacb | 0.979 | 1.000 | -1.250 | 0.245 | 0.000 | -0.353 | -0.234 | -0.378 |
| Q8R0F3 | Sumf1 | 0.002 | 0.024 | 0.000 | 0.232 | -0.568 | 1.860 | 1.650 | 1.660 |
| Q8VEE4 | Rpa1 | 0.072 | 0.197 | 0.054 | -0.271 | 0.000 | -0.678 | -1.690 | -0.560 |
| P97313 | Prkdc | 0.549 | 0.818 | 0.000 | -0.329 | 0.216 | -0.441 | -0.410 | 0.217 |
| P04202 | Tgfb1 | 0.054 | 0.163 | 0.000 | 0.044 | -0.363 | -0.731 | -0.344 | -0.794 |
| P12023 | App | 0.032 | 0.117 | 4.640 | -1.820 | 0.000 | 7.180 | 7.130 | 7.230 |
| O35144 | Terf2 | 0.061 | 0.176 | -0.147 | 0.408 | 0.000 | -0.504 | -0.507 | -0.217 |
| P60766 | Cdc42 | 0.652 | 0.928 | 0.377 | -0.109 | 0.000 | 0.126 | -0.409 | 0.199 |
| Q9JLN9 | Mtor | 0.116 | 0.271 | 0.095 | 0.000 | -0.092 | -0.022 | -0.391 | -0.444 |
| P28798 | Grn | 0.144 | 0.311 | 0.111 | 0.000 | -1.300 | -0.647 | -1.980 | -2.100 |
| Q5DW34 | Ehmt1 | 0.432 | 0.687 | 0.000 | -0.729 | 0.184 | -0.148 | -1.100 | -0.351 |
| P17879 | Hspa1b | 0.401 | 0.652 | 0.000 | -1.290 | 0.818 | 0.709 | 0.226 | 0.366 |
| P08226 | Apoe | 0.508 | 0.772 | 0.861 | -0.299 | 0.000 | 0.960 | 0.161 | 0.360 |
| Q63844 | Mapk3 | 0.097 | 0.239 | 0.625 | -0.043 | 0.000 | -0.104 | -0.713 | -0.406 |
| Q8K3K7 | Agpat2 | 0.136 | 0.300 | 0.673 | -0.697 | 0.000 | 0.782 | 0.680 | 0.732 |
| Q62077 | Plcg1 | 0.003 | 0.033 | -0.125 | 0.000 | 0.471 | -1.190 | -1.370 | -1.750 |
| P33609 | Pola1 | 0.060 | 0.176 | 0.058 | 0.000 | -0.834 | -1.570 | -1.450 | -0.752 |

| | | | | | | | | | |
|---|---|---|---|---|---|---|---|---|---|
| P70371 | Terf1 | 0.025 | 0.102 | 0.000 | 0.395 | -0.619 | -1.120 | -1.050 | -1.160 |
| P25799 | Nfkb1 | 0.017 | 0.080 | 0.000 | -0.065 | 0.139 | -0.430 | -1.120 | -0.909 |
| P28867 | Prkcd | 0.874 | 1.000 | 0.163 | 0.000 | -1.170 | -0.357 | 0.299 | -1.260 |
| P11440 | Cdk1 | 0.243 | 0.453 | 0.534 | 0.000 | -1.540 | 1.200 | -0.031 | 0.779 |
| P00493 | Hprt1 | 0.004 | 0.038 | 0.000 | 0.060 | -0.208 | -1.260 | -1.950 | -1.240 |
| O88351 | Ikbkb | 0.730 | 1.000 | 0.314 | -0.360 | 0.000 | 0.876 | -0.396 | -0.052 |
| P63166 | Sumo1 | 0.032 | 0.119 | -0.103 | 0.000 | 0.363 | -0.366 | -0.411 | -0.342 |
| Q8R4K2 | Irak4 | 0.016 | 0.078 | 0.152 | 0.000 | -0.178 | -0.685 | -1.670 | -1.410 |
| P35991 | Btk | 0.317 | 0.551 | 0.632 | 0.000 | -0.712 | -0.088 | -0.620 | -0.963 |
| Q925J9 | Med1 | 0.684 | 0.960 | 0.315 | -0.113 | 0.000 | 0.119 | -0.251 | 0.102 |
| Q9D6Y7 | Msra | 0.068 | 0.189 | 0.549 | -0.217 | 0.000 | -0.252 | -0.768 | -0.814 |
| Q60848 | Hells | 0.176 | 0.360 | 0.617 | -0.364 | 0.000 | 0.051 | -1.150 | -0.971 |
| Q9D1M4 | Eef1e1 | 0.658 | 0.934 | 0.000 | 0.081 | -0.123 | 0.368 | -0.741 | -0.135 |
| P08556 | Nras | 0.188 | 0.378 | -0.332 | 0.239 | 0.000 | 0.065 | 0.343 | 0.475 |
| Q9D666 | Sun1 | 0.011 | 0.064 | -0.159 | 0.054 | 0.000 | 0.644 | 0.375 | 0.368 |
| P34902 | Il2rg | 0.090 | 0.230 | -0.469 | 0.000 | 0.113 | -0.901 | -0.440 | -0.531 |
| P97494 | Gclc | 0.814 | 1.000 | 0.000 | -3.610 | 0.176 | -1.950 | -2.290 | -0.243 |
| Q9WUD1 | Stub1 | 0.453 | 0.711 | 0.263 | 0.000 | -0.427 | -2.900 | 0.114 | 0.084 |
| B2RWS6 | Ep300 | 0.670 | 0.948 | 0.000 | 1.170 | -0.194 | 0.000 | 0.248 | 0.134 |
| O35904 | Pik3cd | 0.505 | 0.769 | 0.000 | -0.553 | 0.026 | -0.185 | -0.997 | -0.099 |
| Q60974 | Ncor1 | 0.231 | 0.436 | -0.323 | 0.025 | 0.000 | -0.567 | -1.790 | -0.112 |
| Q60855 | Ripk1 | 0.652 | 0.929 | 0.790 | 0.000 | -0.944 | 0.818 | -1.660 | -0.584 |
| Q61510 | Trim25 | 0.944 | 1.000 | -0.760 | 2.540 | 0.000 | 0.403 | -0.707 | 1.800 |
| P23506 | Pcmt1 | 0.001 | 0.022 | -0.179 | 0.000 | 0.198 | -0.923 | -0.926 | -0.835 |
| P39428 | Traf1 | 0.026 | 0.103 | 0.000 | 0.232 | -0.135 | -0.817 | -1.800 | -0.841 |
| Q9Z148 | Ehmt2 | 0.654 | 0.930 | -0.141 | 0.000 | 0.802 | -0.151 | -0.571 | 0.696 |
| P51829 | Adcy7 | 0.555 | 0.824 | -0.535 | 0.000 | 0.058 | -0.200 | -0.585 | -0.144 |
| P63073 | Eif4e | 0.110 | 0.261 | 0.056 | 0.000 | -0.079 | -0.067 | -0.746 | -0.443 |
| P34152 | Ptk2 | 0.297 | 0.524 | 0.207 | 0.000 | -0.153 | 0.088 | -0.575 | -0.244 |
| O70433 | Fhl2 | 0.300 | 0.529 | 0.301 | -2.800 | 0.000 | 1.230 | -0.207 | 0.303 |
| P47811 | Mapk14 | 0.232 | 0.436 | 0.000 | 0.568 | -0.624 | -3.770 | -0.803 | -0.291 |
| Q61037 | Tsc2 | 0.494 | 0.759 | -0.314 | 0.000 | 0.272 | 0.238 | -0.069 | 0.232 |
| P34884 | Mif | 0.193 | 0.385 | -0.437 | 0.000 | 0.174 | -0.562 | -4.920 | -1.210 |
| Q06890 | Clu | 0.014 | 0.073 | 0.000 | 0.199 | -0.686 | -1.550 | -2.880 | -1.990 |
| Q9JIB4 | Gtf2h2 | 0.243 | 0.453 | 0.000 | -0.382 | 0.134 | -0.180 | -1.860 | -0.442 |

**Supplementary Table 9.** Senescence associated proteins detected in CD4+ T cells from old mice with or without mito-transfer. CD4+ T cells from young and old mice, and from old mice after mito-transfer were cultured for 4 h before processing for mass spectrometry analysis. Data expressed as median protein Log2 fold change of CD4+ T cells from old mice, from 3 individual old mice (paired experiment). Differentially expressed proteins (DEPs) with adj. p value > 0.5 are highlighted in orange

| GO | Category | Description | Count | % | Log10(P) | Log10(q) |
|---|---|---|---|---|---|---|
| R-MMU-186763 | Reactome | Downstream signal transduction | 4 | 22.22 | -8.2 | -3.99 |
| mmu04066 | KEGG | HIF-1 signaling pathway - Mus musculus (house mouse) | 5 | 27.78 | -7.47 | -3.91 |
| GO:2000377 | GO BP | regulation of reactive oxygen species metabolic process | 4 | 22.22 | -4.97 | -2.31 |
| GO:0006974 | GO BP | DNA damage response | 6 | 33.33 | -4.62 | -2.02 |
| GO:1903828 | GO BP | negative regulation of protein localization | 4 | 22.22 | -4.33 | -1.88 |
| GO:0062197 | GO BP | cellular response to chemical stress | 4 | 22.22 | -4.11 | -1.72 |
| GO:0046434 | GO BP | organophosphate catabolic process | 3 | 16.67 | -3.7 | -1.39 |
| GO:0032496 | GO BP | response to lipopolysaccharide | 4 | 22.22 | -3.58 | -1.3 |
| GO:0062012 | GO BP | regulation of small molecule metabolic process | 4 | 22.22 | -3.55 | -1.27 |
| GO:0001819 | GO BP | positive regulation of cytokine production | 4 | 22.22 | -3 | -0.85 |
| GO:0002443 | GO BP | leukocyte mediated immunity | 3 | 16.67 | -2.41 | -0.37 |

**Supplementary Table 10.** The Top 20 processes of significantly downregulated DEPs in CD4+ T cells with or without mitochondrial transfer, in relation to T Cell senescence. Data generated using the Metascape analysis platform.

| GO | Description | Log10(P) |
|---|---|---|
| R-MMU-186763 | Downstream signal transduction | -8.7 |
| WP373 | IL-3 signaling pathway | -8.4 |
| WP387 | IL-6 signaling pathway | -8.4 |

**Supplementary Table 11.** The Top 3 Protein-Protein Interaction (PPI) networks of significantly downregulated DEPs in CD4+ T cells, with or without mito-transfer, in relation to T Cell senescence. Data generated using the Metascape analysis platform.

| MCODE | GO | Description | Log10(P) |
|---|---|---|---|
| MCODE_1 | mmu00010 | Glycolysis / Gluconeogenesis - Mus musculus (house mouse) | -9.3 |
| MCODE_1 | GO:0032787 | monocarboxylic acid metabolic process | -7.8 |
| MCODE_1 | mmu00620 | Pyruvate metabolism - Mus musculus (house mouse) | -7.1 |
| MCODE_2 | R-MMU-186763 | Downstream signal transduction | -11.7 |
| MCODE_2 | R-MMU-186797 | Signaling by PDGF | -10.6 |
| MCODE_2 | WP407 | Kit receptor signaling pathway | -10 |

**Supplementary Table 12.** The MCODE networks identified significantly downregulated DEPs in CD4+ T cells, with or without mito-transfer, in relation to T Cell senescence. Data generated using the Metascape analysis platform.
